# Integrative Transcriptomic and miRNA Analysis Reveals Immune Suppression and Metabolic Reprogramming in FGFR3–TACC3 Fusion-Positive versus Fusion-Negative Bladder Cancer

**DOI:** 10.1101/2025.06.05.658144

**Authors:** Divya Mishra, Shivangi Agrawal, Divya Malik, Ekta Pathak, Rajeev Mishra

## Abstract

FGFR3–TACC3 gene fusion drives a molecularly distinct subset of bladder cancer characterized by post-transcriptional immune evasion and metabolic rewiring. To elucidate the transcriptomic and regulatory consequences of this fusion, we performed an integrative analysis of mRNA and miRNA expression profiles in fusion-positive versus fusion-negative bladder tumors using TCGA-derived RNA-Seq and miRNA-Seq data. Fusion-positive tumors exhibited upregulation of mitochondrial oxidative phosphorylation and ribosomal genes (e.g., COX5B, ATP5F1D, RPS15) and marked downregulation of immune regulatory genes (e.g., CD4, STAT3, IL6, CD86, PTPRC). Pathway enrichment revealed suppression of JAK/STAT, PD-L1 signaling, and cytokine networks, consistent with a fusion-driven immunosuppressive phenotype. We identified fusion-specific miRNA–mRNA regulatory interactions, including upregulated miRNAs (miR-1226-3p, miR-149-5p, miR-301b-3p) that target immune transcripts and downregulated miRNAs (miR-199b-5p, miR-214-3p) that normally suppress mitochondrial and ribosomal genes. In contrast, fusion-negative tumors displayed an immunoreactive signature marked by elevated IFNG, CXCL10, and IRF1 expression and downregulation of immune-suppressive miRNAs (miR-125a-5p, miR-23b-3p). Minimum free energy analysis and AlphaFold3-based AGO2-RISC docking confirmed structural feasibility of 3D duplex formation for 13 miRNA–target pairs, supporting active silencing complexes. Immune deconvolution revealed reduced infiltration of cytotoxic T cells, NK cells, and dendritic cells in fusion-positive tumors, while fusion-negative tumors showed robust immune cell presence. Key gene–immune cell correlations further confirmed differential immune architecture: SRC and CD4 were strongly linked to suppressed infiltration in fusion-positive tumors, while IFNG and CXCL10 positively correlated with T and NK cell abundance in fusion-negative cases. ROC analysis highlighted COMT, RPS15, and miR-187-3p as robust classifiers of fusion-positive tumors (AUC > 0.9), and CXCL10, CD74, and miR-142 for fusion-negative tumors. Connectivity Map analysis highlighted Lofexidine and Lenalidomide as candidate compounds predicted to reverse fusion-specific transcriptional programs. Collectively, our findings define immune suppression and metabolic reprogramming as core features of FGFR3–TACC3 fusion-positive bladder cancer and nominate fusion-specific miRNA–mRNA interactions as candidate biomarkers and therapeutic targets.

## Introduction

Gene fusions are well-established oncogenic drivers across a broad spectrum of human malignancies. Historically, their tumorigenic potential has been largely attributed to the formation of chimeric proteins with constitutive or deregulated activity [1, 2]. However, emerging evidence has illuminated an additional, critical consequence of gene fusions: disruption of post-transcriptional gene regulation, particularly via microRNAs (miRNAs) [3, 4]. These small, non-coding RNAs regulate gene expression by binding to complementary sequences in the 3′ untranslated regions (3′UTRs) of target mRNAs, leading to translational repression or transcript degradation [5].

Fusion events commonly interfere with miRNA-mediated gene regulation through two principal mechanisms. First, many fusions result in truncation or complete loss of the 3′UTR of the 5′ partner gene, eliminating key miRNA binding sites and allowing oncogenic transcripts to evade post-transcriptional suppression [4]. Second, promoter-hijacking fusions may juxtapose the coding region of a proto-oncogene with a highly active or tissue-specific promoter, resulting in overexpression of the 3′ gene partner that overwhelms endogenous miRNA-based control, even if the 3′UTR remains intact [6, 7]. In both scenarios, these alterations disrupt endogenous miRNA–mRNA feedback loops, permitting sustained oncogene expression and signaling [3].

Large-scale transcriptomic and miRNA profiling efforts, including analyses from The Cancer Genome Atlas (TCGA), have consistently demonstrated that fusion-positive tumors exhibit distinct miRNA expression profiles compared to their fusion-negative counterparts [7]. These differences correlate with critical oncogenic processes such as enhanced proliferation, survival, immune evasion, and metastatic potential. A paradigmatic example is the FGFR3– TACC3 (F3–T3) fusion, initially characterized in glioblastoma and subsequently identified in multiple solid tumors, including bladder cancer (BLCA) [8, 9]. This fusion contributes to tumorigenesis through dual mechanisms: constitutive activation of receptor tyrosine kinase signaling and the loss of miRNA-mediated post-transcriptional repression due to deletion of the FGFR3 3′UTR, thereby circumventing inhibition by tumor-suppressive miRNAs such as miR-99a and miR-100 [10].

In BLCA, Fibroblast growth factor receptors (FGFRs) fusions are observed in approximately 5% of urothelial carcinoma [11]. (FGFRs) fusions including FGFR3– TACC3, FGFR3–JAKMIP1, FGFR3–TNIP2, and FGFR3–BAIAP2L1—have been detected in approximately 40% of non-muscle-invasive bladder cancer (NMIBC) cases and 15% of muscle-invasive bladder cancer (MIBC) cases [12, 13]. The FGFR3–TACC3 fusion arises from an in-frame tandem duplication at chromosome 4p16.3 that fuses the FGFR3 tyrosine kinase domain with the coiled-coil domain of TACC3 (transforming acidic coiled coil-containing protein 3) [14]. This fusion promotes ligand-independent dimerization and constitutive receptor activation, leading to persistent stimulation of downstream oncogenic signaling pathways such as PI3K/AKT, RAS/MAPK, and STAT3 [14, 15]. Simultaneously, the TACC3 component of the fusion interferes with mitotic spindle assembly by displacing endogenous TACC3 from centrosomes, contributing to chromosomal instability and tumor progression [16].

Importantly, the fusion transcript lacks the native FGFR3 3′UTR, abolishing binding sites for miR-99a/100 and miR-181a-5p—miRNAs that are significantly downregulated in BLCA and inversely correlated with FGFR3 expression [17] [18, 19]. This 3′UTR loss enables the fusion transcript to escape miRNA-mediated suppression, resulting in sustained FGFR3 overexpression and hyperactivation of downstream pathways. Additional dysregulated miRNAs, such as let-7a and miR-451—which regulate key oncogenes like MYC—further indicate widespread perturbation of post-transcriptional regulatory networks in the FGFR3– TACC3-positive context.

Clinically, FGFR3–TACC3 fusion-positive tumors exhibit more aggressive behavior, higher recurrence rates, reduced responsiveness to standard therapies, and poorer overall prognosis [13, 20, 21]. In NMIBC, this fusion is associated with progression to MIBC, necessitating frequent invasive surveillance procedures such as cystoscopy, which impose considerable physical and financial burdens. Although FGFR-targeted tyrosine kinase inhibitors are available, resistance mechanisms and immune escape frequently compromise therapeutic efficacy, highlighting the need for novel, personalized therapeutic strategies.

A critical but underexplored dimension of FGFR3–TACC3-driven oncogenesis lies in its impact on the tumor immune microenvironment (TIME). The TIME comprises a dynamic ecosystem of tumor cells, stromal components, and immune populations—including macrophages, CD4⁺ T cells, and natural killer (NK) cells—that collectively influence tumor initiation, immune evasion, and metastatic dissemination [22, 23]. Notably, fusion-positive MIBC tumors exhibit markedly reduced immune cell infiltration relative to fusion-negative or NMIBC counterparts, suggesting an immunosuppressive microenvironment driven by altered signaling and potentially miRNA dysregulation [4, 24]. This immune alteration may reflect fusion-driven remodeling of cytokine and chemokine networks, mediated in part by downstream effects on miRNA regulatory circuits.

Given their role as both effectors and indicators of gene fusion status, miRNAs offer dual utility as mechanistic drivers and clinically actionable biomarkers. Deciphering how FGFR3– TACC3 fusions reshape miRNA–mRNA interactions and modulate immune landscapes in BLCA could yield novel diagnostic tools and uncover therapeutic vulnerabilities, particularly for tumors resistant to current treatment paradigms.

In this study, we comprehensively characterize the molecular and immunological distinctions between FGFR3–TACC3 fusion-positive and fusion-negative bladder cancers. By integrating transcriptomic profiling, miRNA–mRNA regulatory network analysis, and immune deconvolution using algorithms such as ESTIMATE and MCP-counter, we identify key miRNA–hub gene axes and TIME-associated gene modules linked to fusion status. In particular, we highlight a subset of differentially expressed, FGFR3-targeting miRNAs with potential utility as diagnostic markers and therapeutic targets. Our findings shed light on the complex post-transcriptional and immunological alterations associated with FGFR3–TACC3-driven bladder cancer and support the development of miRNA-based strategies for precision oncology in this molecularly defined disease subset.

## Materials and Methods

### Data Retrieval and Preprocessing

RNA-sequencing (RNA-Seq) and microRNA-sequencing (miRNA-Seq) data for bladder cancer (BLCA) and corresponding adjacent normal tissue samples were obtained from The Cancer Genome Atlas (TCGA) through the Genomic Data Commons (GDC) online portal (https://portal.gdc.cancer.gov/). RNA-Seq data were specifically acquired from STAR-count files, while miRNA-Seq data were obtained from BCGSC miRNA profiling files. Utilizing the ChimerSeq module within the ChimerDB 4.0 database, we extracted the list of FGFR3-TACC3 (F3-T3) fusion-positive BLCA samples [25]. Our selection process identified eight unique barcode IDs with confirmed in-frame fusion events; these IDs are detailed in Supplementary Table S1.

To dissect the transcriptional impact of the FGFR3–TACC3 (F3-T3) gene fusion in bladder cancer (BLCA), we stratified the dataset into three comparison groups: (i) fusion-positive vs. fusion-negative BLCA, (ii) fusion-positive vs. normal bladder tissue, and (iii) fusion-negative vs. normal bladder tissue. The dataset initially included RNA-Seq data from 412 primary BLCA tumors and 19 adjacent normal tissues. For miRNA-Seq, a total of 406 samples were retrieved, encompassing eight F3-T3 fusion-positive BLCA, 398 F3-T3 fusion-negative BLCA, and 19 normal samples. To enhance reliability and comparability in subsequent analyses, Principal Component Analysis (PCA) using prcomp() and voom methods from LIMMA-voom algorithm were applied to refine the dataset selection by removing overlapping samples and samples with low gene counts [26, 27]. Post PCA, our refined sample sets included seven fusion-positive and 16 fusion-negative BLCA samples for the differential expression analysis in fusion-positive versus fusion-negative cases, six fusion-positive and 12 adjacent normal samples, and 12 fusion-negative and 11 adjacent normal samples for other comparative analyses (Table 1).

**Table 1.**
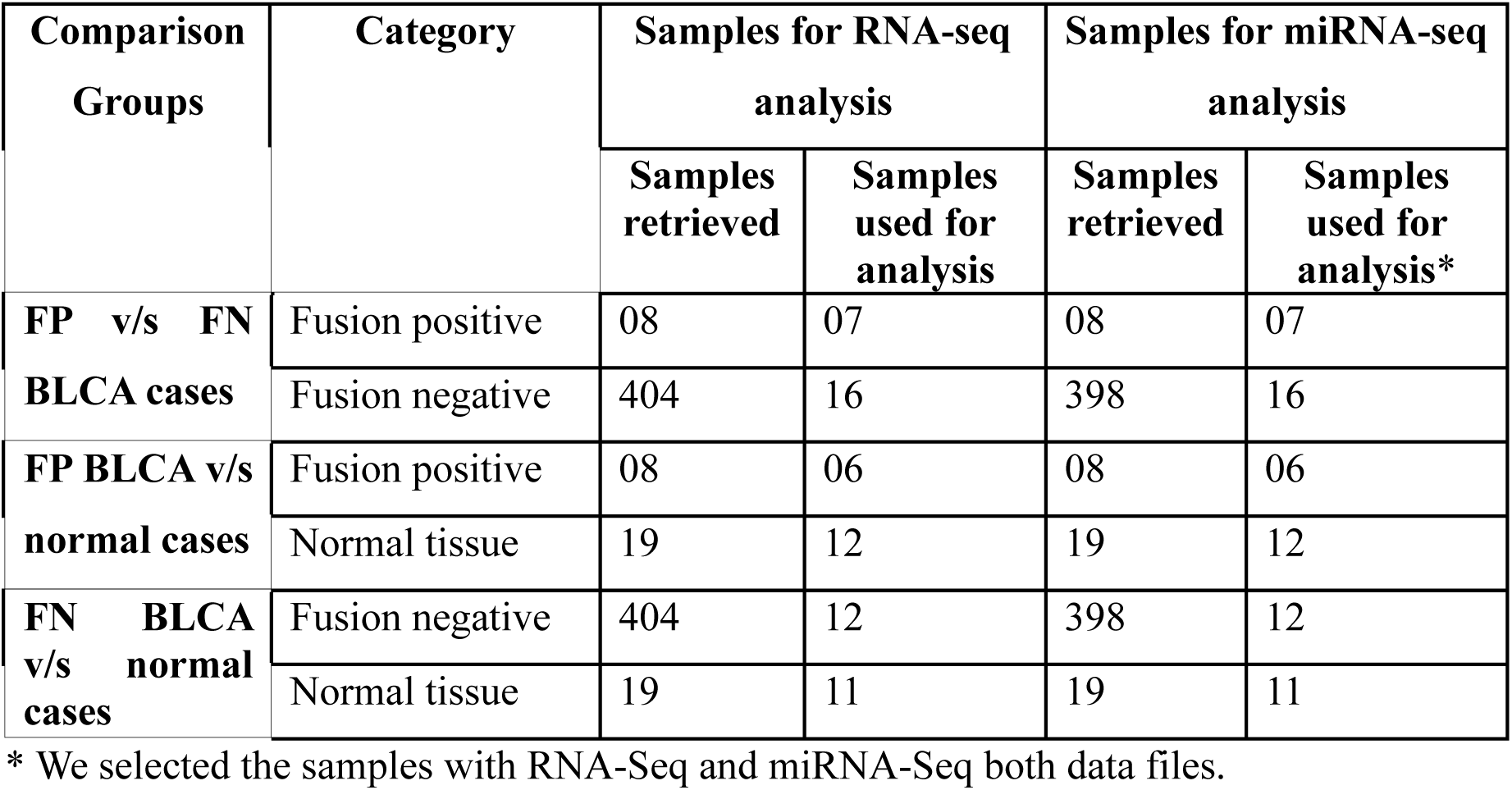
Details of the samples retrieved and used for data analysis in this study.

### Differential Gene and miRNA Expression Analyses

Differential gene and miRNA expression analyses were performed using DESeq2 [28] and LIMMA-voom methods on R software [29]. DESeq2 employs a Wald test for differential expression analysis, whereas LIMMA-voom normalizes count data into log2 counts per million (logCPM) and applies a linear mixed-effects model [28] [29]. Genes and miRNAs were considered significantly differentially expressed if the adjusted adjusted p-value < 0.05 and absolute log2 fold change| ≥ 1. Results from both methods were visualized using volcano plots via the EnhancedVolcano R package [30]. Robust differentially expressed genes (DEGs) and miRNAs (DEMs) identified by intersecting the results from both methods were used for further analysis, employing Venn diagrams through the InteractiVenn online tool [31].

### Protein-protein interaction (PPI) network

Protein-protein interaction networks were constructed using the STRING app within Cytoscape software [32, 33], to explore interactions among exclusive sets of differentially expressed genes for fusion-positive and fusion-negative BLCA cases. Separate networks were created for upregulated and downregulated DEGs. Influential "hub" genes (top 10) were determined using node degree distribution from Network Analyzer and the Maximum Clique Centrality (MCC) algorithm from cytoHubba [34]. Dense modules within each network were identified using the MCODE app, applying a degree cutoff of 2, node score cutoff of 0.2, K-core of 2, and a maximum depth of 100 [35]. Hub genes were further examined for their presence within these densely connected modules.

### Functional and Pathway Enrichment Analyses

Functional implications of the identified DEGs exclusive to fusion-positive and fusion-negative BLCA were assessed through Gene Ontology (GO) terms covering biological processes, molecular functions, cellular components, and KEGG pathways using DAVID bioinformatics resources [36]. Significantly enriched GO terms and pathways were identified with a p-value cutoff of 0.05. Bubble plots generated using ggplot2 facilitated visual interpretation of enrichment results [37]. We performed the GO and KEGG enrichment analyses for the top three statistically significant modules obtained from each of the PPI networks.

### Gene Set Enrichment Analysis (GSEA)

Gene set enrichment analysis (GSEA) was conducted to determine the biological relevance of differential expression results using gene sets from c5.all.v7.5.1.symbols.gmt and c2.cp.kegg.v2023.1.Hs.symbols.gmt databases within MSigDB. Phenotypic labels for GSEA were assigned as fusion-positive versus fusion-negative, and analyses were performed with 1,000 permutations. Default parameters in GSEA software version 4.3.2 were maintained to ensure standardized comparative results [38].

### miRNA-Gene Target Prediction and Regulatory Network Construction

To predict miRNA-gene interactions, mature miRNAs were retrieved from the miRBase database [39]. These were queried against the mirDIP 4.1 database to obtain both predicted and experimentally validated targets [40]. A stringent threshold was applied—only high confidence targets predicted by at least three independent databases were retained. Additionally, only miRNA-target interactions mapping to the 3’ untranslated region (3’UTR) were considered.

To identify biologically relevant targets, predicted gene targets were cross-referenced with DEGs to derive differentially expressed targets (DETs) exclusive to fusion-positive and fusion-negative BLCA samples. FGFR3- and TACC3-targeting miRNAs and their downstream DETs were of particular interest. These gene targets were used to construct a regulatory network.

Regulatory networks were built in Cytoscape v3.9.1 [33], linking upregulated DEMs to downregulated DETs and vice versa. Using network degree centrality, we identified the top 10 hub DEMs from each interaction network. These hub miRNAs were then mapped to hub DEGs identified in the PPI networks to prioritize the most influential regulatory relationships. The final refined networks highlight these hub DEM–hub DEG interactions within each BLCA subtype.

### Minimum Free Energy calculations for miRNA-Gene Targets Interaction

To validate the miRNA-target interactions, mature miRNA sequences were downloaded from the miRBase database, and 3’UTR sequences of candidate mRNAs were obtained from Ensembl BioMart in FASTA format. RNAhybrid was employed to calculate the minimum free energy (MFE) of duplex formation [41]. An MFE threshold of ≤ -17 kcal/mol was set to define high-affinity interactions. miRNA-mRNA pairs were visualized using heatmaps created with the pheatmap R package [42]. Interactions passing the MFE threshold were prioritized for further biological interpretation and experimental follow-up.

### miRNA-mRNA Duplex Structure Prediction using AlphaFold 3

To further characterize the physical interactions between miRNAs and their mRNA targets, we predicted both two-dimensional (2D) and three-dimensional (3D) structures of selected miRNA-mRNA duplexes. Initial 2D structural predictions were generated using RNAhybrid [43]. For 3D modeling, we utilized the AlphaFold 3 server to simulate the complex formed by the miRNA-mRNA duplex bound to human Argonaute 2 (HsAGO2) protein [44]. The protein sequence of HsAGO2 was obtained from the UniProt database [45]. Argonaute proteins associate with guide RNAs to form complexes that slice transcripts that pair to the guide [46].

Model accuracy was assessed based on interface predicted template modeling (ipTM) and predicted template modeling (pTM) scores. Structures with ipTM ≥ 0.8 and pTM ≥ 0.9 were deemed reliable. We visualized the interactions between the miRNA-mRNA duplex and AGO2 domains (N, PAZ, MID, PIWI) using UCSF Chimera v1.18 [47]. miRNA-mRNA pairs showing spatial proximity to AGO2’s positively charged residues in the central channel were considered splicing-competent, reflecting potential for post-transcriptional silencing [46] [44].

### Tumor Microenvironment Analysis

To evaluate the tumor immune microenvironment (TIME) and stromal infiltration, we computed ESTIMATE scores using the ESTIMATE R package [48]. These scores, derived from normalized expression matrices, quantify tumor purity (ESTIMATE score), stromal content (Stromal score), and immune cell infiltration (Immune score).

To deconvolute the immune landscape, the Microenvironment Cell Populations-counter (MCP-counter) method implemented in the Immunedeconv R package was applied [49]. This allowed estimation of the abundance of 11 immune and stromal cell populations, including CD8+ T cells, NK cells, B cells, macrophages, dendritic cells, neutrophils, endothelial cells, and cancer-associated fibroblasts (CAFs). Data were visualized using boxplots to compare infiltration patterns across fusion-positive, fusion-negative, and normal tissues.

### Immune-Related Gene Retrieval and Correlation Analysis

We compiled a reference list of immune-related genes from the Immunome and IMMPORT databases [50, 51]. Differentially expressed immune-related genes were identified by comparing this list with the DEGs exclusive to fusion-positive and fusion-negative BLCA TIME.

To uncover associations between immune gene expression, ESTIMATE-derived scores, and immune cell infiltration, we performed correlation analyses using Pearson’s correlation coefficient [52]. Correlation values of R ≥ 0.7 with p-value ≤ 0.05 were considered statistically significant. Heatmaps were generated using the corrplot R package, and scatter plots were constructed to assess linear associations. This analysis enabled identification of key immune regulatory axes specific to the tumor microenvironment of fusion-positive and fusion-negative BLCA cases.

### ROC Curve Analysis

To evaluate the diagnostic potential of candidate genes and miRNAs identified from the PPI network and enrichment analyses, we performed Receiver Operating Characteristic (ROC) curve analysis. Initially, we calculated the variance across samples and excluded low-variance genes and miRNAs to focus on informative features. Data were then log-transformed to obtain normalized expression values. ROC curves and Area Under the Curve (AUC) metrics were computed using the pROC package in R [53]. Genes and miRNAs achieving AUC > 0.7 were considered to have significant discriminatory power between fusion-positive, fusion-negative, and normal bladder cancer samples. Visualization of ROC curves was carried out using the plotROC function from the ggplot2 package [54].

### Connectivity Map (CMap) Analysis for Therapeutic Candidate Identification

To identify small molecules with potential therapeutic efficacy against fusion-positive and fusion-negative bladder cancer (BLCA), we employed the Connectivity Map (CMap) database query platform [55]. CMap analysis was designed to match disease-associated transcriptional signatures with gene expression profiles modulated by various small molecules, thereby facilitating identification of compounds with likely inhibitory effects on the disease phenotype [55].

For fusion-positive BLCA, the transcriptional signature was constructed by merging the upregulated differentially expressed genes (DEGs) from significant modules 1, 2, and 3, as identified through the protein-protein interaction (PPI) network clustering. These merged DEGs formed a consolidated gene signature, which was used as a query input. In contrast, the fusion-negative BLCA condition exhibited sufficient DEG representation in each individual module; thus, upregulated DEGs from each module were used separately to construct independent gene signatures. Each of these individual gene sets was submitted independently as a query. All gene expression queries were run against the CMap Touchstone reference dataset, which contains L1000 transcriptional profiles across a wide range of small molecules [55].

CMap employs a connectivity mapping algorithm to calculate the similarity between the user-submitted gene expression signature and reference perturbagen profiles. The result is reported as a normalized connectivity score (NCS). A positive NCS indicates a similar transcriptional response between the disease state and the perturbagen, suggesting a potential mimetic effect. Conversely, a negative NCS indicates an antagonistic or inhibitory effect of the compound on the submitted gene expression profile [55]. The magnitude of the NCS reflects the degree of similarity or dissimilarity, with larger absolute values indicating stronger relationships.

For therapeutic discovery, we prioritized small molecules with negative normalized connectivity scores, as these represent candidates capable of reversing disease-associated gene expression signatures. From each fusion condition query (fusion-positive merged query and fusion-negative module-specific queries), the top 20 small molecules with the most negative NCS values were selected for downstream analysis.

Only compounds with known mechanisms of action (MOA) were retained for final consideration. Molecules lacking annotated MOAs were excluded to ensure biological interpretability and translational potential. Furthermore, for each selected compound, its reported gene targets were extracted and cross-referenced against our DEG dataset. Compounds were prioritized if their targets were significantly upregulated in our study or if the molecular pathways associated with these targets were significantly dysregulated based on prior enrichment analyses.

This approach allowed us to identify pharmacologically actionable compounds that are not only predicted to reverse disease-specific gene expression signatures but also exhibit mechanistic relevance to the underlying molecular pathophysiology of FGFR3-TACC3 fusion-driven and fusion-negative BLCA subtypes.

## Results

### Transcriptomic Profiling of F3-T3 Fusion-Positive and Fusion-Negative BLCA Samples

To elucidate the transcriptomic differences associated with FGFR3-TACC3 (F3-T3) fusion status in bladder cancer (BLCA), we conducted differential gene expression (DEG) analysis across three comparison groups: F3-T3 fusion-positive vs. fusion-negative, F3-T3 fusion-positive vs. normal adjacent bladder tissue, and F3-T3 fusion-negative vs. normal adjacent bladder tissue.

Prior to DEG analysis, we applied unsupervised Multi-Dimensional Scaling (MDS) and Principal Component Analysis (PCA) to assess sample distribution and remove outliers. These dimensionality reduction approaches allowed identification of well-separated clusters, facilitating the selection of high-confidence samples for downstream comparisons (Table 1). The PCA plots, constructed using normalized expression data, revealed distinct stratification between sample groups in all comparisons. The first two principal components (PCs), PC1 and PC2, captured the majority of variance within the datasets. Specifically, in the fusion-positive vs. fusion-negative cohort, PC1 and PC2 accounted for 34.9% and 30.2% of the variance, respectively. In the fusion-positive vs. normal comparison, PC1 explained 31.0% and PC2 21.9% of the variance. For the fusion-negative vs. normal comparison, PC1 and PC2 contributed 20.8% and 8.5% of the variance, respectively. These results underscore the transcriptional divergence among the three groups (Figure 1). After removing overlapping or low-quality samples identified in PCA and MDS analyses, we assessed the quality of expression data using mean-variance plots, confirming a linear mean-variance relationship across three comparison groups—a prerequisite for reliable DEG analysis using DESeq2 and LIMMA.

**Figure 1.**
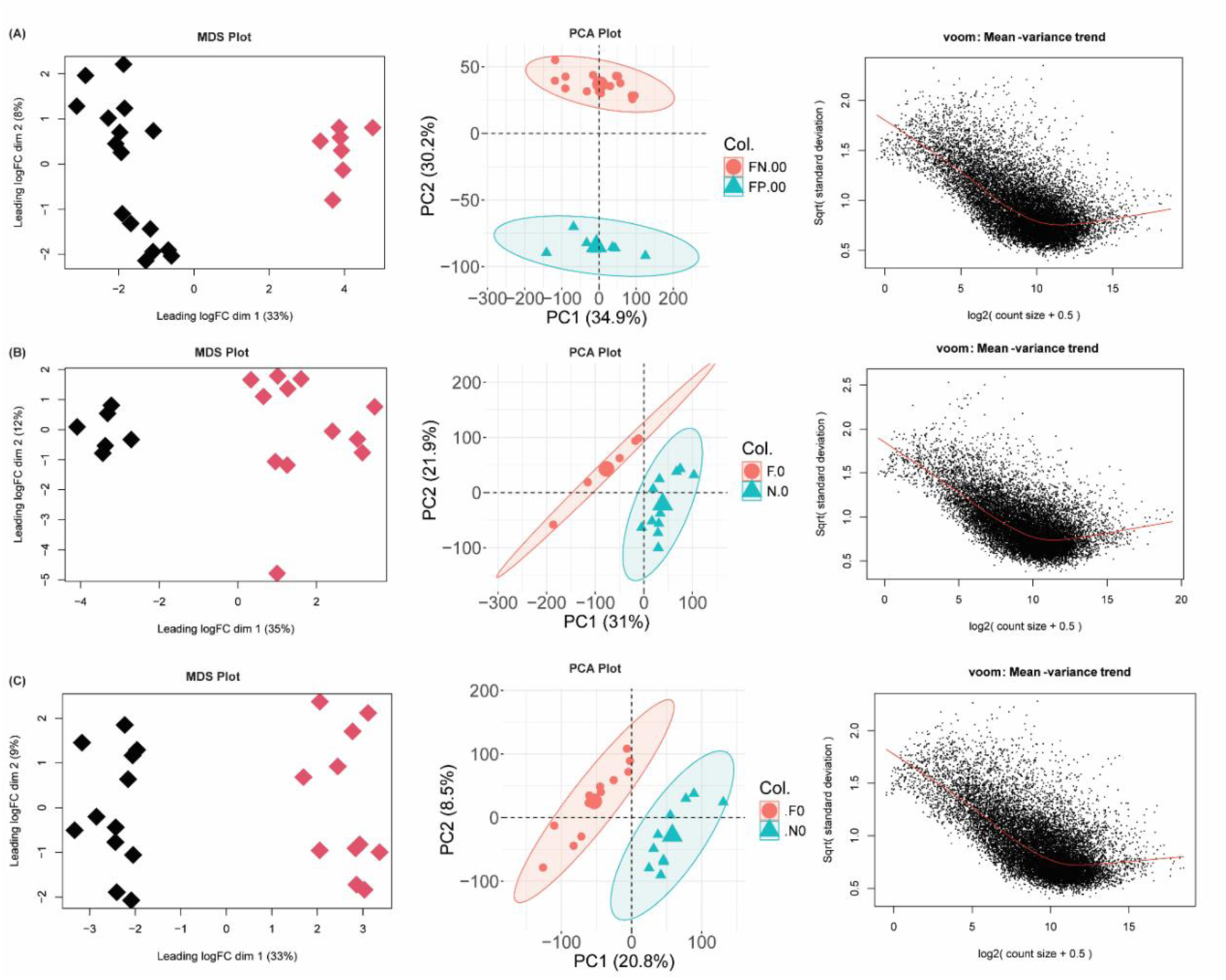
Multi-dimensional scaling (MDS) plot of the paired samples of all three BLCA datasets, including tumor (red dots) and normal (black dots) RNA-Seq data. (A) F3-T3 fusion-positive and fusion-negative BLCA, (B) F3-T3 fusion-positive and normal samples, and (C) F3-T3 fusion-negative BLCA and normal samples. Euclidean distances between samples were calculated based on genes with the largest standard deviations. Principal Component Analysis (PCA) plots using normalized counts are shown to visualize clustering patterns. Smoothed mean-variance plots validate data suitability for differential analysis.

Differential expression analysis was performed using both DESeq2 and LIMMA algorithms, ensuring robustness and reproducibility of identified DEGs. The total number of DEGs in each comparison is summarized in Table 2. Venn diagram analysis revealed 4,479 DEGs (1,509 upregulated and 2,970 downregulated) in the fusion-positive vs. fusion-negative cohort, 4,070 DEGs (1,496 upregulated and 2,574 downregulated) in the fusion-positive vs. normal cohort, and 4,520 DEGs (2,028 upregulated and 2,492 downregulated) in the fusion-negative vs. normal cohort (Figure 2C). A detailed list of unique DEGs identified exclusively in fusion-positive and fusion-negative groups is provided in Supplementary Table S2 (Figure 2D). The LIMMA-based analysis of the fusion-positive vs. fusion-negative dataset identified the top ten upregulated protein-coding genes as ANXA10, BTBD16, HMGCS2, C10orf99, CYP4F8, MSMB, PHGR1, CLCA4, GPX2, and MYBPC1. The most significantly downregulated genes included SFRP2, SFRP4, GREM1, CCL21, LRRC15, FNDC1, DPT, DES, CXCL13, and OGN. Corresponding volcano plots for the remaining two comparisons are presented in Figure 2A and 2B. A comparative evaluation between LIMMA and DESeq2 outputs demonstrated substantial overlap among the top-ranking DEGs. Notably, in the fusion-positive vs. fusion-negative group, three upregulated genes (PHGR1, MYBPC1, MSMB) and two downregulated genes (SFRP2, OGN) were consistently identified in both methods. In the fusion-positive vs. normal cohort, four upregulated genes—CST4, HSD3B1, SLC1A6, and CST1—were common to both approaches. For the fusion-negative vs. normal comparison, seven upregulated DEGs (KRT6B, KRT14, CASP14, S100A7, KRT6C, KRT6A, and S100A7A) and six downregulated DEGs (CRTAC1, SLC14A1, BMP5, SLC5A7, TMEM132C, and SHH) were recurrent across the two analytical platforms.

**Figure 2.**
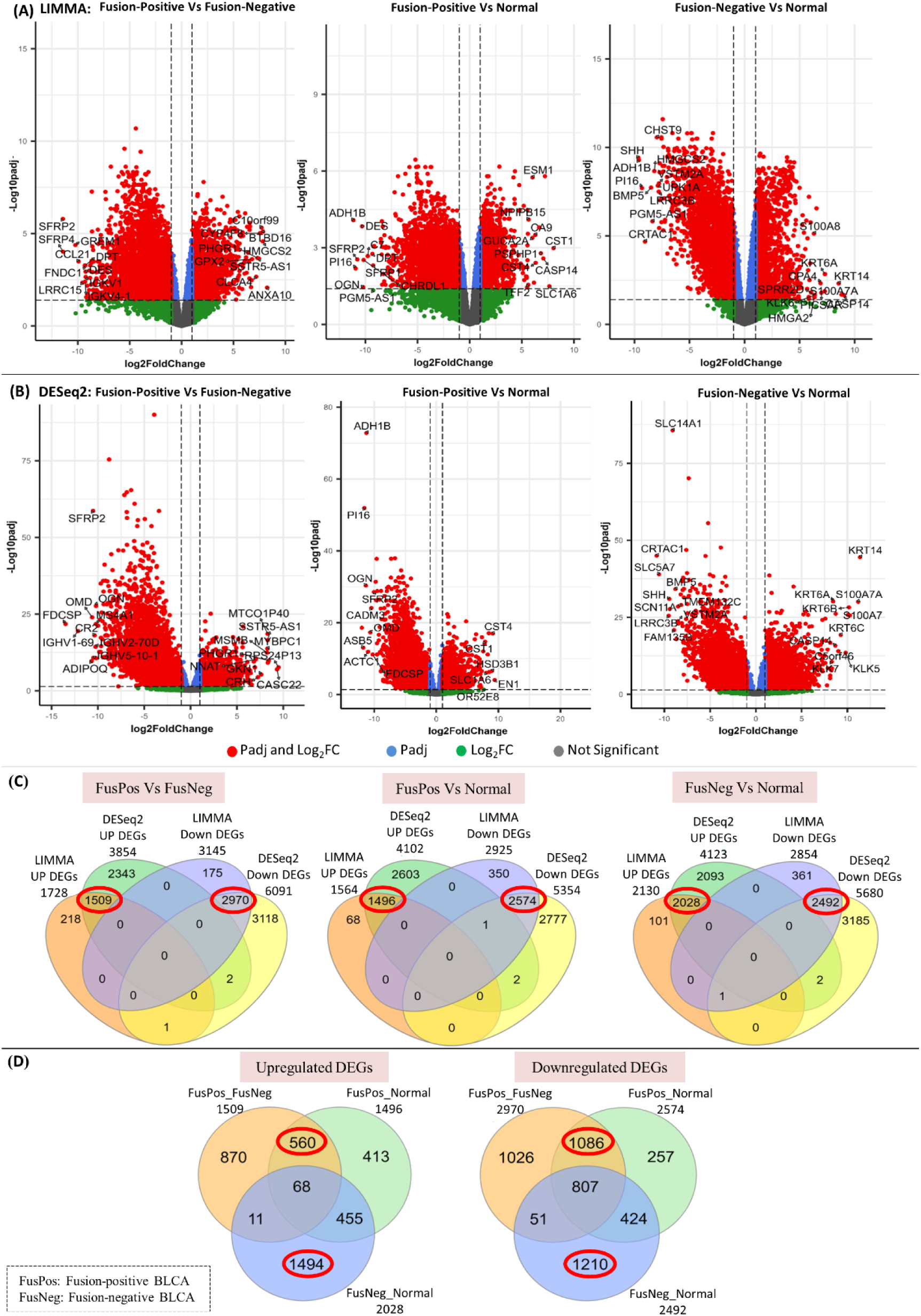
Analysis of differentially expressed genes in BLCA. (A) Volcano plots showing DEGs identified by LIMMA. (B) Volcano plots showing DEGs identified by DESeq2. (C) Venn diagrams displaying intersecting DEGs (highlighted in red) from both methods across the three comparisons. (D) Venn diagram analysis showing upregulated and downregulated genes exclusive to fusion-positive (FusPos) and fusion-negative (FusNeg) BLCA samples.

**Table 2.**
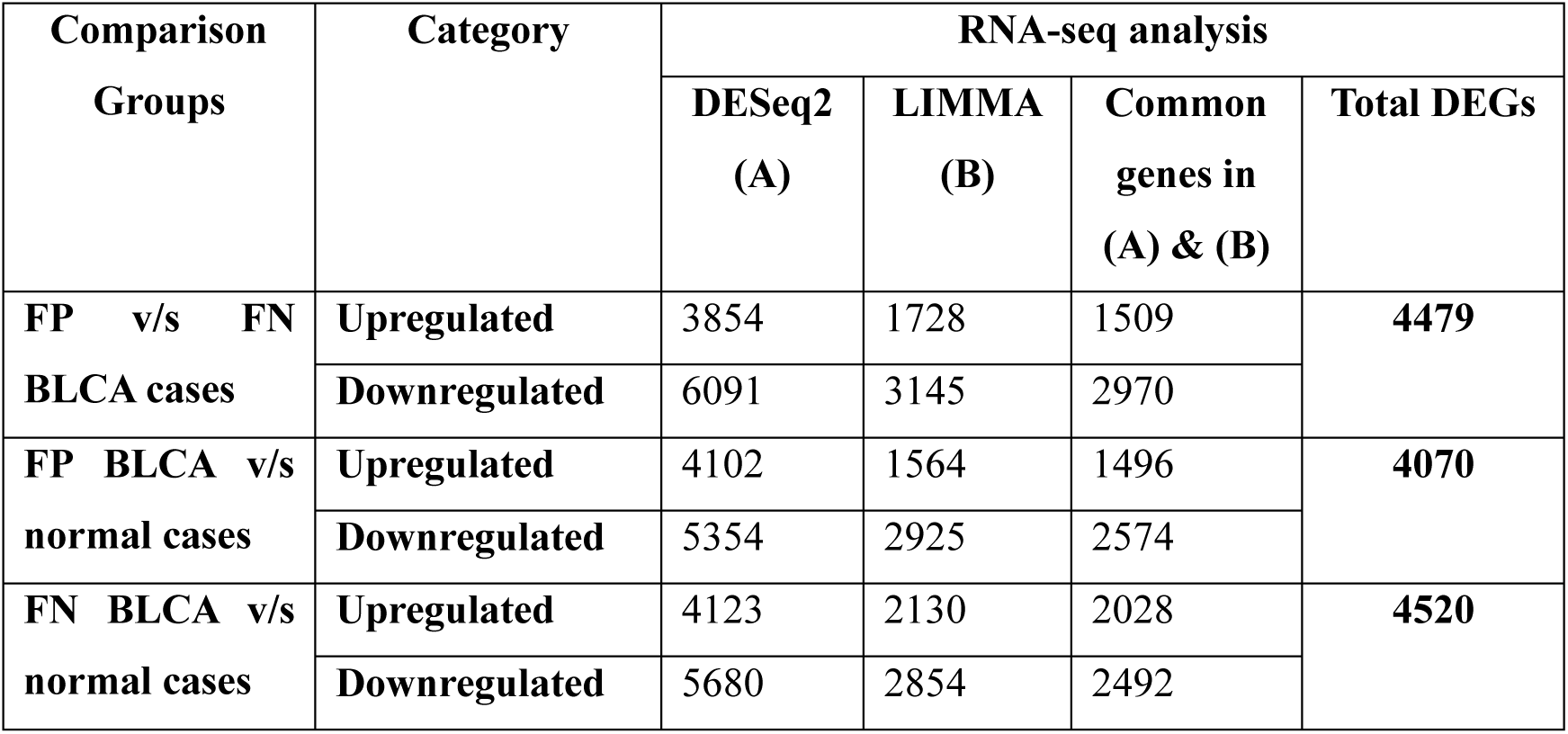
DEGs identified via DESeq2 and LIMMA tools.

Despite using consistent thresholds (adjusted p-value < 0.05 and |log2 fold change| ≥ 1), variations in DEG composition were observed between DESeq2 and LIMMA outputs, highlighting the influence of algorithm-specific statistical modeling (Figure 2A–B). To mitigate such discrepancies, only DEGs identified by both methods were considered for downstream analyses.

### PPI Network Analysis Reveals Distinct Sets of Hub Genes in Fusion-Positive and Fusion-Negative BLCA

To elucidate key PPI interactions in F3–T3 fusion-associated bladder cancer (BLCA), we constructed protein–protein interaction (PPI) networks for differentially expressed genes (DEGs) from both F3–T3 fusion-positive and fusion-negative cases.

### Fusion-Positive BLCA: PPI Modules and Hub Gene Identification

We performed the PPI network analysis using 422 upregulated and 1048 downregulated protein-coding DEGs identified in fusion-positive BLCA samples. From these networks, three significant modules (module scores >4.0) were extracted using MCODE clustering. Degree centrality was applied to prioritize influential nodes. Ten high-degree genes were classified as hub genes from the upregulated subset: COMT, COX5B, NDUFA13, RPS15, COX6B1, ATP5F1D, FASN, SRC, NDUFS5, and UQCR10 (Figure 3A; Table 3). Of these, seven genes—COX5B, NDUFA13, RPS15, COX6B1, ATP5F1D, NDUFS5, and UQCR10—were not only high-degree interactors but also constituents of the significant modules, reinforcing their central role in the fusion-positive PPI architecture. Conversely, among the downregulated DEGs, ten hub genes were discerned: IL6, PTPRC, STAT3, ITGAM, CCL5, CD4, CD8A, FCGR3A, STAT1, and CD86. All of these genes were components of the top-ranked modules, particularly modules 1 and 2 (Figure 3A). These genes are prominently linked to immune modulation and inflammatory signaling, suggesting functional attenuation in the fusion-positive tumor microenvironment.

**Figure 3.**
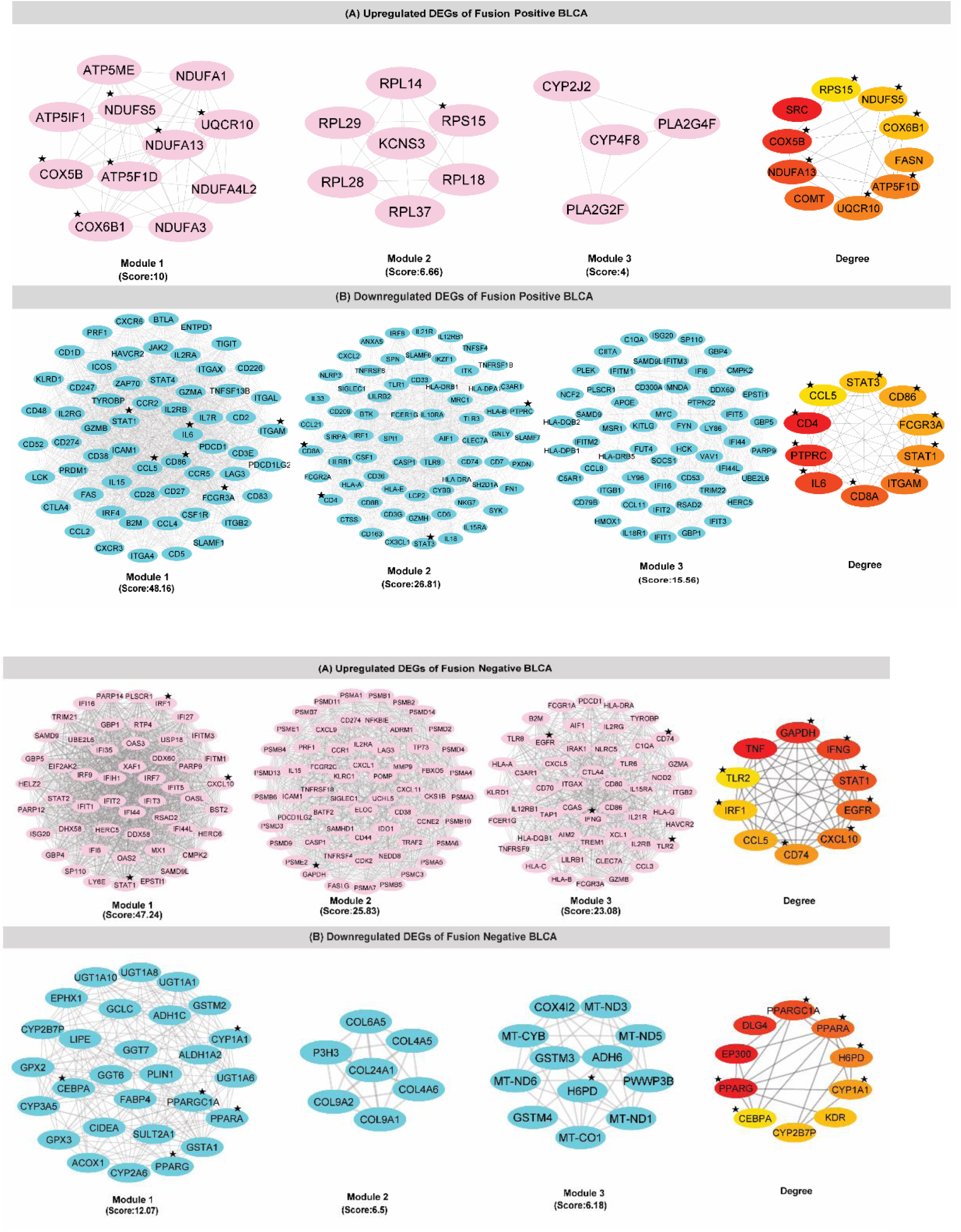
Figure 3. Protein–Protein Interaction (PPI) Network Analysis, Module Extraction, and Hub Gene Identification. A) (Top three significant modules and top ten hub genes identified from the upregulated DEGs in F3–T3 fusion-positive bladder cancer (BLCA) samples. B) Top three significant modules and top ten hub genes from the downregulated DEGs in the same samples. Bottom Panel: A) PPI modules and hub gene identification for upregulated DEGs in fusion-negative BLCA samples compared to normal tissue. B) Corresponding PPI network and hub genes from the downregulated DEGs in fusion-negative cases. Hub genes were prioritized based on degree centrality using CytoHubba. The color gradient from red to yellow represents the hub score intensity. Asterisks indicate hub genes that overlapped with those found within the top-ranking network modules.

**Table 3.**
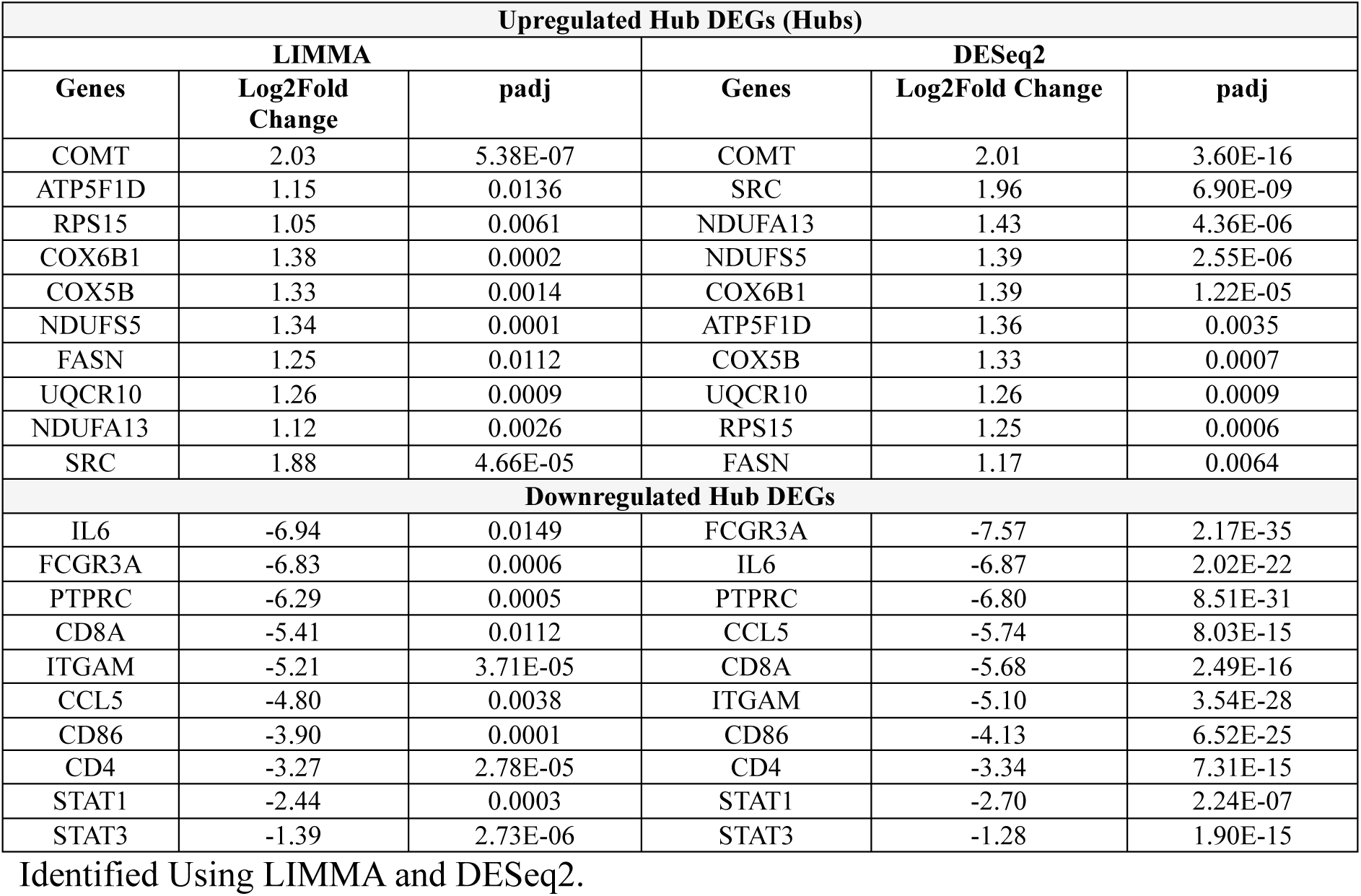
Differentially Expressed Hub Genes in Fusion-Positive vs Fusion-Negative BLCA.

### Fusion-Negative BLCA: PPI Modules and Hub Gene Identification

An analogous PPI analysis was extended to 1494 upregulated and 1210 downregulated exclusive DEGs derived from fusion-negative vs. normal BLCA comparisons. For both upregulated and downregulated networks, three top-scoring MCODE modules were extracted. Degree centrality metrics revealed the following ten upregulated hub genes: IFNG, CCL5, TLR2, IRF1, TNF, CD74, STAT1, EGFR, GAPDH, and CXCL10 (Figure 3B; Table 4). Notably, eight of these genes—excluding CCL5 and TNF—were embedded within the core interaction modules, highlighting their regulatory relevance. For the downregulated subset, ten hub genes were delineated: PPARG, KIT, KDR, DLG4, CEBPA, EP300, H6PD, PPARGC1A, PPARA, and CYP1A1. Among these, six genes (PPARG, CEBPA, H6PD, PPARGC1A, PPARA, CYP1A1) overlapped with nodes in the extracted modules, supporting their potential role in metabolic and epigenetic deregulation in fusion-negative BLCA cases (Figure 3B).

**Table 4.** Differentially Expressed Hub Genes in Fusion-Negative vs Normal BLCA Identified Using LIMMA and DESeq2.

| Upregulated DEGs (Hubs) |  |  |  |  |  |
| --- | --- | --- | --- | --- | --- |
| LIMMA |  |  | DESeq2 |  |  |
| Genes | Log2Fold Change | padj | Genes | Log2Fold Change | padj |
| CXCL10 | 5.20 | 3.43E-05 | CXCL10 | 5.03 | 1.06E-10 |
| IFNG | 3.96 | 0.0002 | IFNG | 3.85 | 9.24E-09 |
| STAT1 | 2.15 | 6.47E-06 | CCL5 | 2.29 | 1.98E-06 |
| TLR2 | 2.14 | 9.49E-05 | TLR2 | 2.20 | 4.99E-09 |
| IRF1 | 1.82 | 0.0001 | STAT1 | 1.92 | 2.33E-07 |
| CCL5 | 1.81 | 0.00097 | TNF | 1.75 | 0.0005 |
| TNF | 1.66 | 0.00691 | IRF1 | 1.69 | 6.77E-05 |
| GAPDH | 1.34 | 1.82E-06 | EGFR | 1.53 | 0.0003 |
| CD74 | 1.31 | 0.0105 | CD74 | 1.39 | 0.0020 |
| EGFR | 1.13 | 0.0058 | GAPDH | 1.35 | 2.13E-12 |
| Downregulated Hub DEGs |  |  |  |  |  |
| CYP1A1 | -6.55 | 0.0055 | CYP1A1 | -5.59 | 5.00E-07 |
| KIT | -4.69 | 3.77E-08 | KIT | -4.22 | 3.85E-22 |
| PPARGC1A | -4.50 | 6.53E-05 | PPARGC1A | -3.33 | 3.57E-05 |
| PPARG | -2.84 | 0.0006 | PPARG | -2.10 | 0.0020 |
| KDR | -1.57 | 0.0009 | KDR | -1.66 | 0.0001 |
| DLG4 | -1.27 | 0.0007 | H6PD | -1.16 | 1.09E-08 |
| CEBPA | -1.25 | 0.0086 | DLG4 | -1.13 | 0.0014 |
| PPARA | -1.19 | 0.0006 | PPARA | -1.05 | 0.0004 |
| H6PD | -1.17 | 1.75E-05 | CEBPA | -1.02 | 0.0259 |
| EP300 | -1.02 | 8.28E-05 | EP300 | -1.01 | 1.63E-06 |

The comprehensive list of hub DEGs, along with their respective log₂ fold-change values and statistical significance (padj), as identified using both LIMMA and DESeq2 pipelines, are provided in Table 3 (fusion-positive vs. fusion-negative) and Table 4 (fusion-negative vs. normal BLCA).

### Functional Enrichment of DEGs Reveals Distinct Molecular Signatures in in fusion-positive and fusion-negative BLCA cohorts

To delineate the biological implications of differentially expressed genes (DEGs) in F3–T3 fusion-positive and fusion-negative bladder cancer (BLCA), we conducted functional enrichment analyses of genes from the most significant modules derived from PPI networks.

### Fusion-Positive BLCA: Functional Annotation of Module-Specific DEGs

Enrichment analysis of upregulated DEGs in fusion-positive BLCA revealed significant involvement in mitochondrial and metabolic functions, including ATP synthesis, oxidative phosphorylation, thermogenesis, and reactive oxygen species (ROS) detoxification. These genes were also enriched in ribosomal biogenesis and rRNA processing, indicating elevated biosynthetic and metabolic activity (Figure 4A). Corresponding KEGG pathway enrichment confirmed their participation in oxidative phosphorylation, thermogenesis, chemical carcinogenesis via ROS, and cardiac muscle contraction pathways. Notably, several hub genes—COX5B, NDUFA13, RPS15, COX6B1, ATP5F1D, NDUFS5, and UQCR10—were consistently represented across these enriched pathways (Table 5). In contrast, downregulated DEGs were strongly associated with immune regulation, including both innate and adaptive responses. Functional annotations implicated these genes in T cell activation, B cell proliferation, Th17 cell differentiation, cytokine production, and apoptotic signaling. KEGG analysis reinforced these findings by highlighting enrichment in cytokine–cytokine receptor interaction, cell adhesion molecules, Th1 and Th2 differentiation, TNF signaling, and the JAK-STAT axis (Figure 4B; Table 5). All ten downregulated hub genes (IL6, PTPRC, STAT3, ITGAM, CCL5, CD4, CD8A, FCGR3A, STAT1, and CD86) were found within these immune-associated pathways.

**Figure 4.**
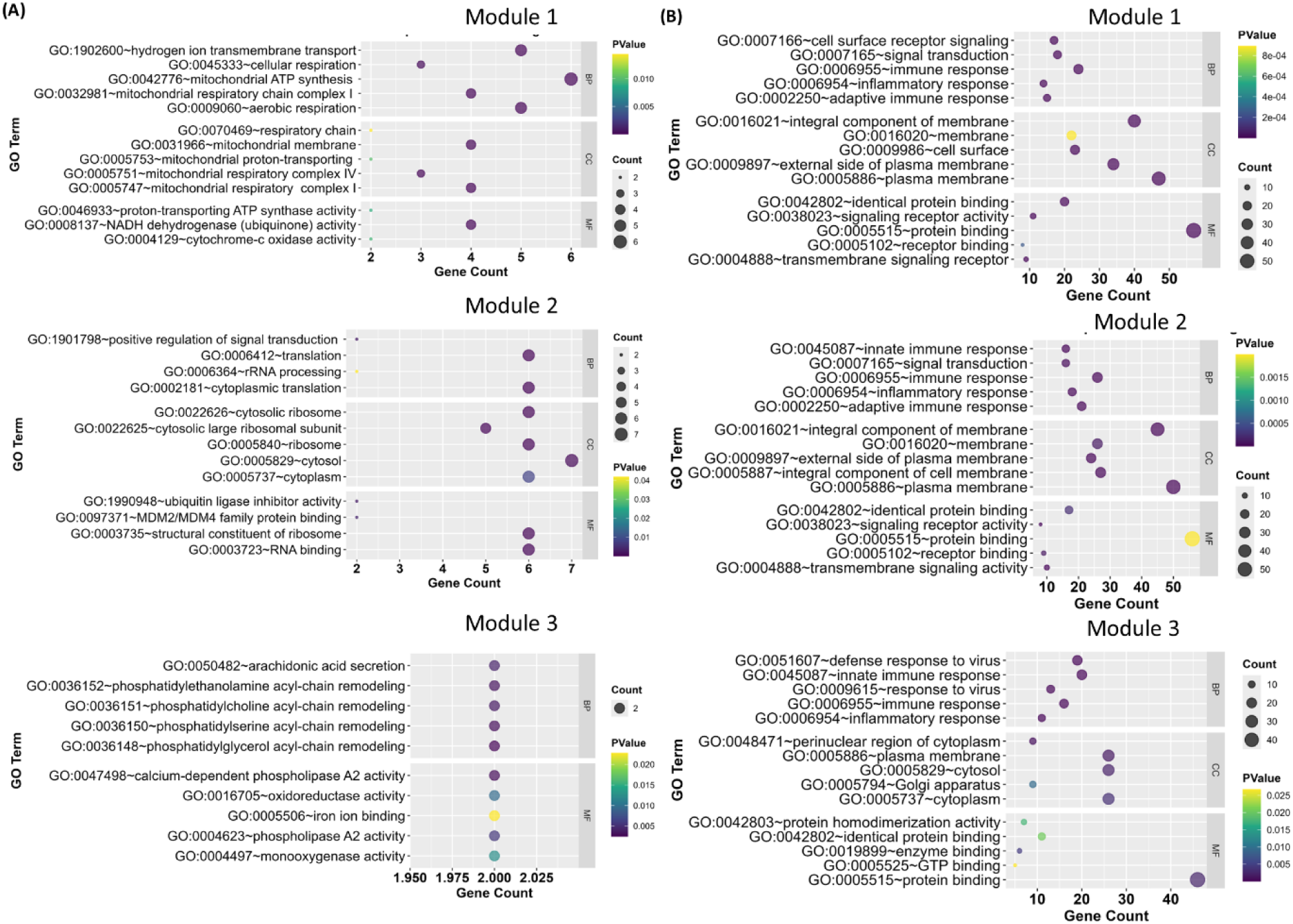
Bubble plots showing enriched biological processes and pathways for module-derived DEGs in fusion-positive BLCA. (A) Functional enrichment for upregulated DEGs modules. (B) Functional enrichment for downregulated DEGs modules. Node size represents gene count per term; color gradient indicates statistical significance.

**Table 5.**
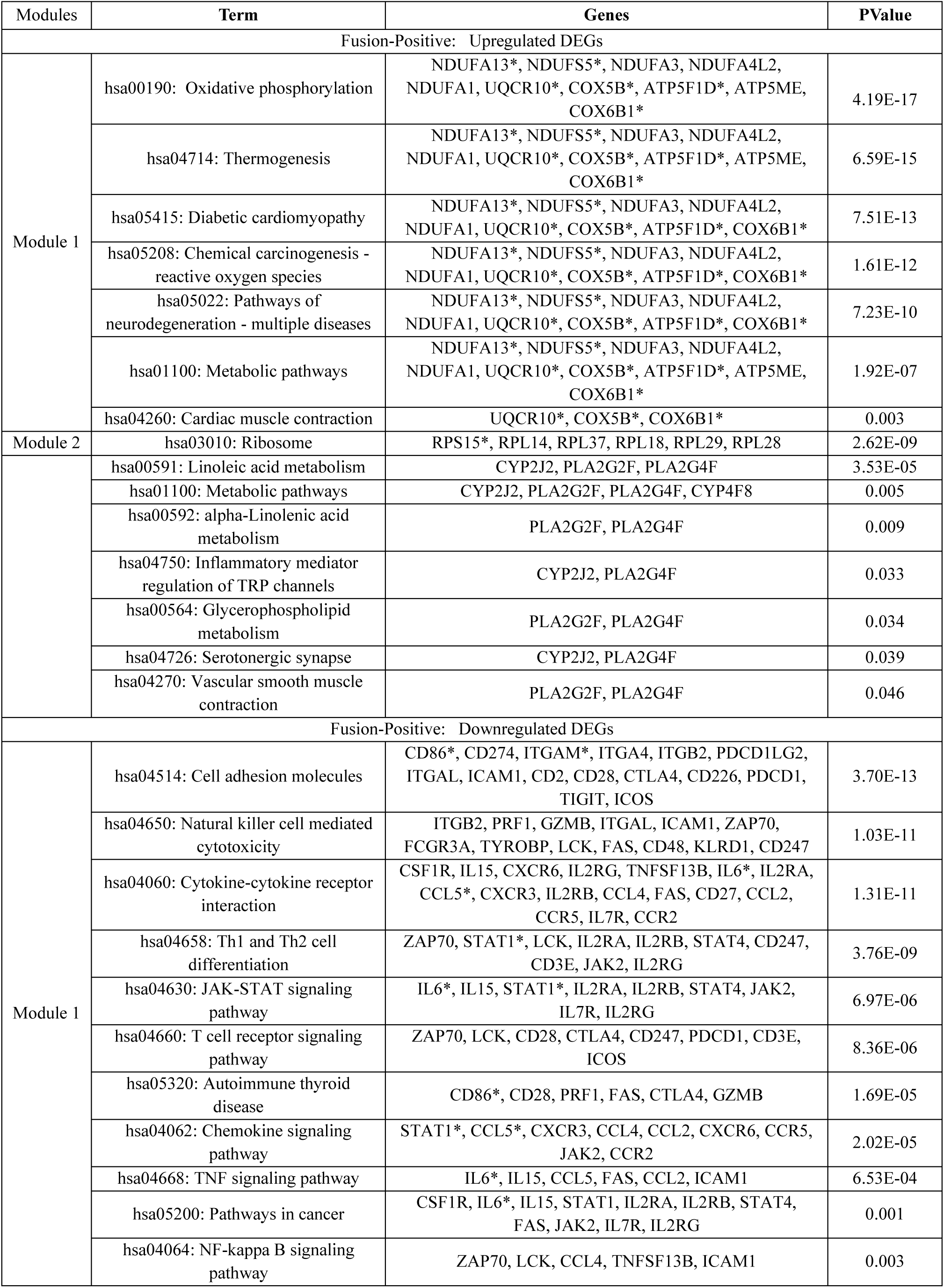
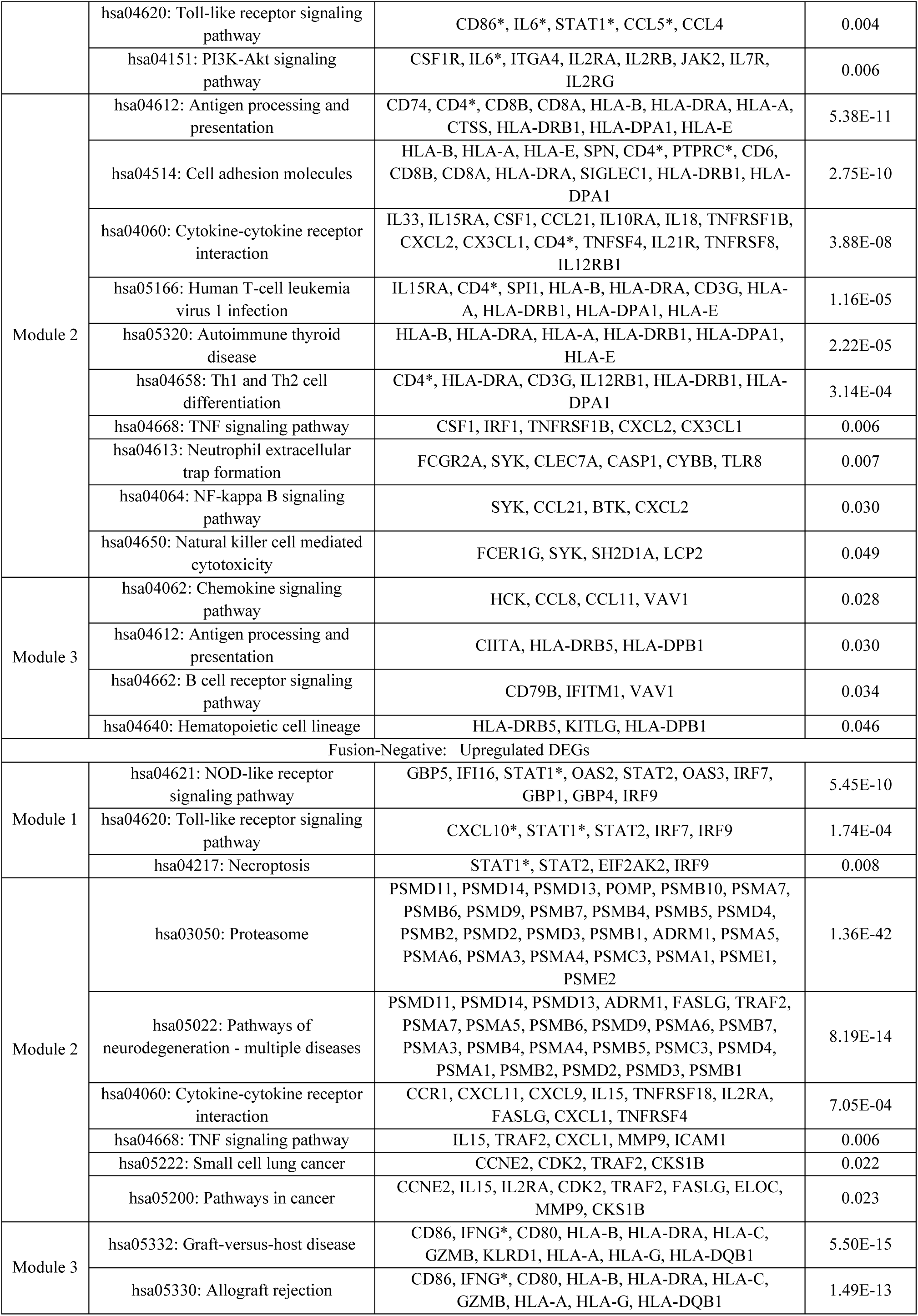
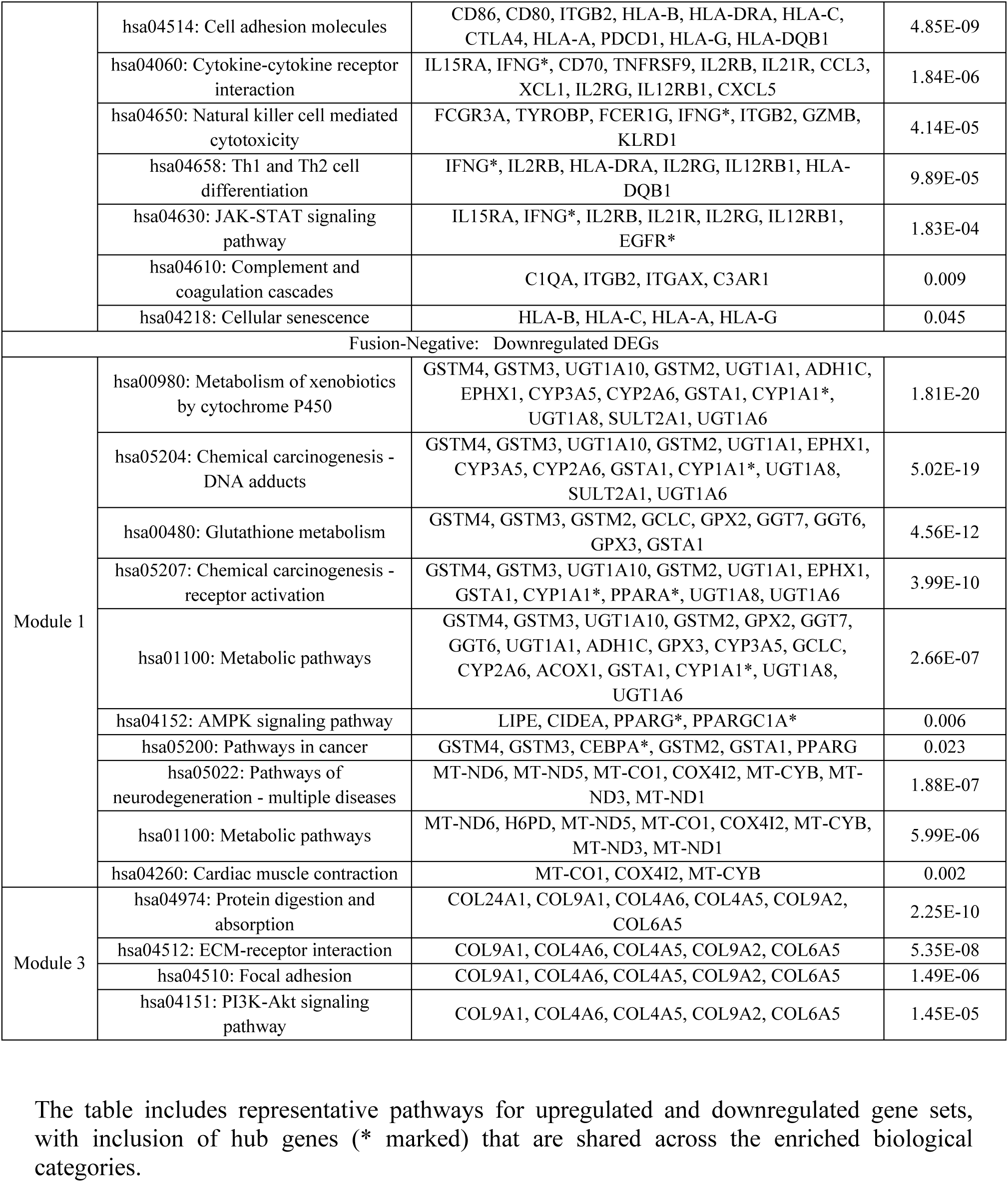
KEGG pathway enrichment analysis of DEGs from significant modules identified in the PPI networks of fusion-positive and fusion-negative BLCA.

### Fusion-Negative BLCA: Enrichment of Immunometabolic Signatures

In the fusion-negative vs. normal comparison, upregulated DEGs were enriched for immune-related functions, including cytokine activity, protein binding, MAPK signaling, canonical Wnt regulation, and JAK-STAT signaling (Supplementary Table S3). The KEGG pathway landscape highlighted necroptosis, cytokine–cytokine receptor interactions, and cellular senescence, along with TNF signaling, multiple neurodegeneration-related pathways, and Th1/Th2 differentiation. Among the ten upregulated hub genes, eight—IFNG, TLR2, IRF1, CD74, STAT1, EGFR, GAPDH, and CXCL10—were present in these enriched networks (Table 5). Downregulated DEGs in this group were predominantly linked to extracellular matrix organization, lipid metabolism, and mitochondrial electron transport. Pathways significantly enriched included metabolic processes, PI3K-Akt signaling, ECM–receptor interaction, cardiac muscle contraction, and focal adhesion. Notably, six of the ten hub genes (PPARG, CEBPA, H6PD, PPARGC1A, PPARA, and CYP1A1) were embedded within these biological networks, further confirming their mechanistic relevance in tumor suppression and metabolic regulation in the fusion-negative BLCA cohort.

The table includes representative pathways for upregulated and downregulated gene sets, with inclusion of hub genes (* marked) that are shared across the enriched biological categories.

### Gene Set Enrichment Analysis Highlights Oncogenic Pathways in Fusion-Positive BLCA

To further investigate dysregulated signaling transduction pathways, we applied Gene Set Enrichment Analysis (GSEA) to transcriptomic data from fusion-positive and fusion-negative samples. In fusion-positive BLCA, the upregulated gene sets were significantly enriched for pathways related to energy metabolism and biosynthesis, including oxidative phosphorylation, ribosome, spliceosome, and steroid hormone biosynthesis, as well as tyrosine, glutathione, and retinol metabolism (Figure 5A; Supplementary Table 4). Leading-edge analysis indicated critical contributors to these pathways. Specifically, COX5B, COX6B1, ATP5F1D, NDUFS5, and UQCR10 were mapped to oxidative phosphorylation; RPS15 and COX5B contributed to ribosomal activity; and COMT was linked to tyrosine metabolism (Supplementary Table 5A). In parallel, downregulated DEGs were significantly associated with immunoregulatory and cell adhesion-related pathways, including complement and coagulation cascades, cytokine–cytokine receptor interaction, chemokine signaling, ECM–receptor interaction, focal adhesion, allograft rejection, and autoimmune thyroid disease (Figure 5B; Supplementary Table 4). These pathways were directly associated with downregulated hub genes such as PTPRC, ITGAM, CD4, CD8A, and CD86 through the CAMs signaling axis. Additional associations included IL6 and CCL5 in cytokine–cytokine interaction, STAT3, CCL5, and STAT1 in chemokine signaling, and IL6, STAT1, and STAT3 in JAK-STAT signaling (Supplementary Table 5B).

**Figure 5.**
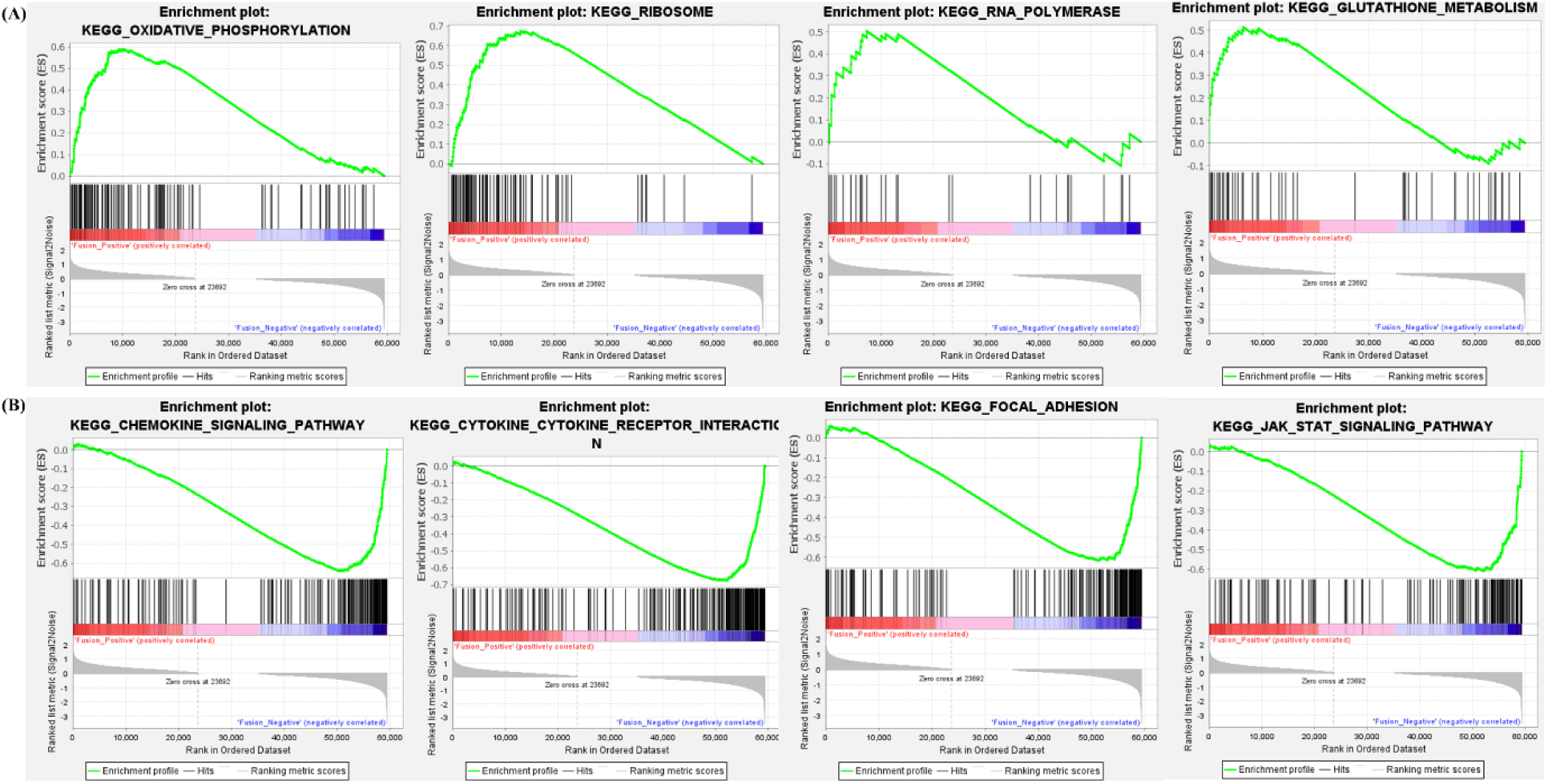
Gene Set Enrichment Analysis (GSEA) plots highlighting significantly enriched pathways in fusion-positive vs fusion-negative BLCA. (A) Pathways associated with upregulated genes. (B) Pathways associated with downregulated genes. Ranked gene list is plotted from right (most upregulated) to left (most downregulated). Enriched pathways are shown at a cutoff of |NES| ≥ 1.

### Differential Expression Analysis of miRNAs Reveals Distinct Regulatory Signatures in Fusion-Positive and Fusion-Negative BLCA

To investigate the post-transcriptional regulatory system in BLCA, we performed differential expression analysis of miRNAs (DEMs) across F3–T3 fusion-positive, fusion-negative, and normal bladder tissue groups. Comparative analysis using both LIMMA and DESeq2 pipelines yielded consistent patterns of miRNA dysregulation. The differential expression profiles are visualized using volcano plots (Figure 6A, B), and the number of DEMs identified in each contrast is summarized in **Table 6**.

**Figure 6.**
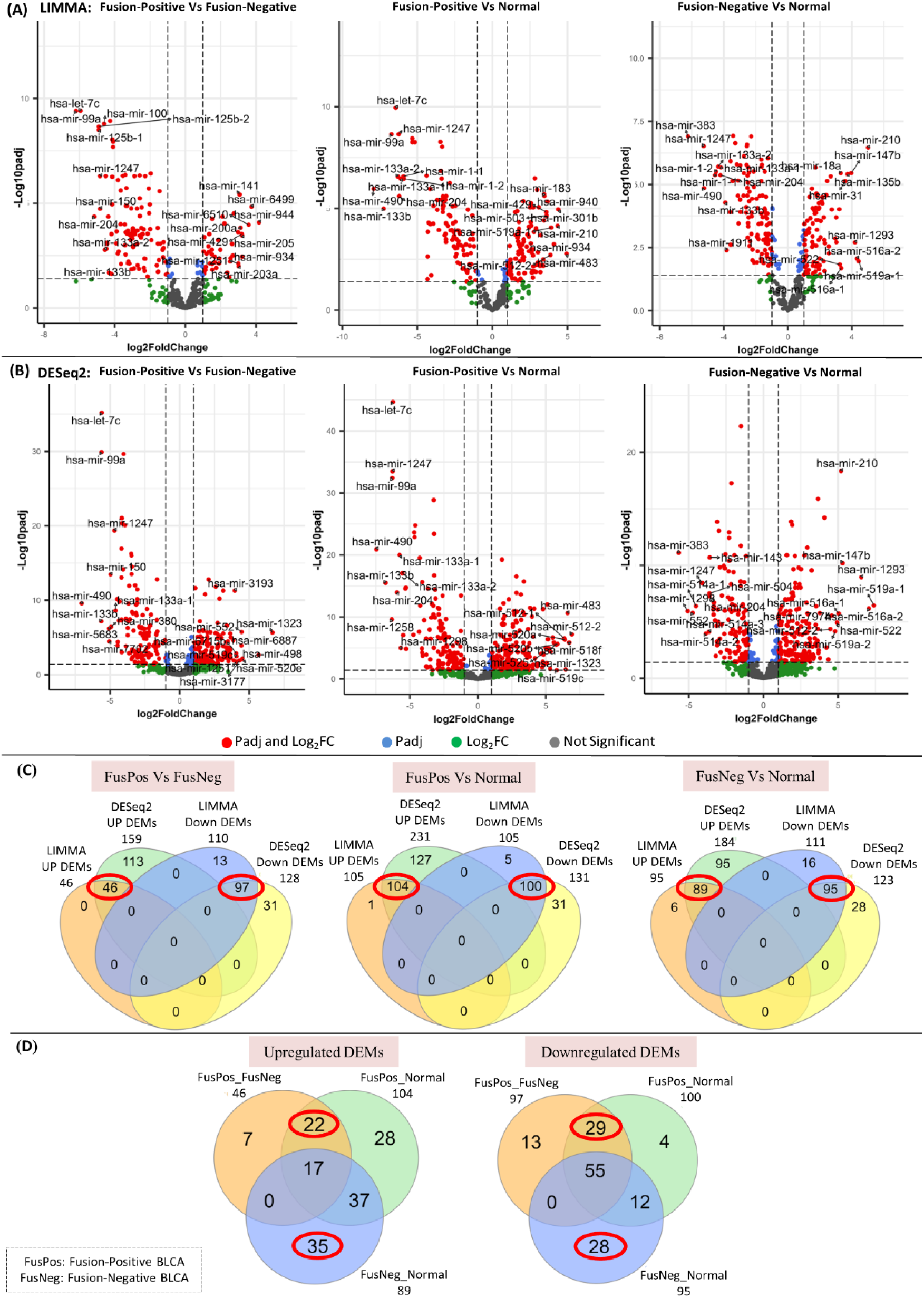
Differential expression of miRNAs in fusion-positive and fusion-negative BLCA cohorts. (A–B) Volcano plots showing DEMs identified using LIMMA and DESeq2, respectively. (C) Venn diagram displaying intersecting DEMs between methods. (D) Venn diagram highlighting exclusive DEMs for fusion-positive and fusion-negative cohorts.

**Table 6.**
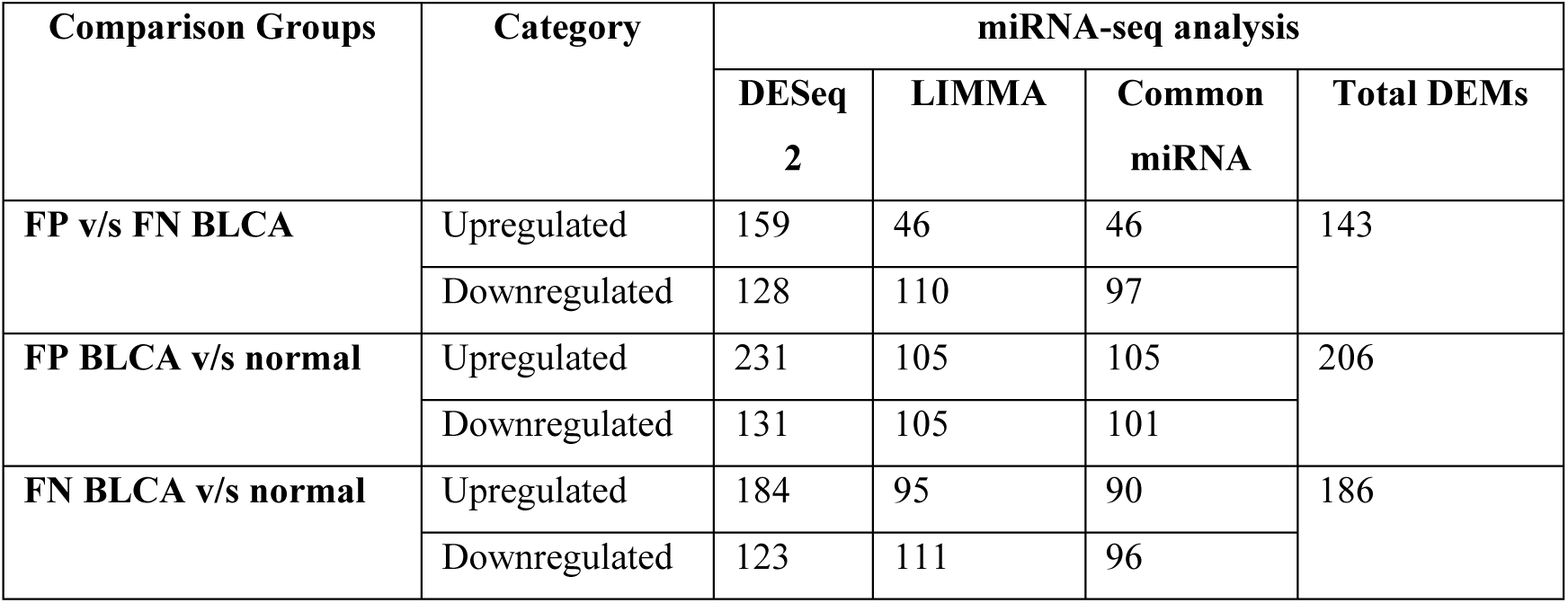
Number of differentially expressed miRNAs (DEMs) identified using LIMMA and DESeq2 in each group comparison (fusion-positive vs fusion-negative, fusion-positive vs normal, fusion-negative vs normal).

Intersecting DEMs obtained via both methods were selected using Venn analysis for robustness (Figure 6C). In total, we identified 47 upregulated and 97 downregulated DEMs in fusion-positive vs. fusion-negative BLCA. Similarly, fusion-positive vs. normal BLCA revealed 105 upregulated and 101 downregulated DEMs, whereas fusion-negative vs. normal BLCA yielded 90 upregulated and 96 downregulated DEMs. Specifically, 22 overexpressed and 29 underexpressed miRNAs were uniquely associated with the fusion-positive cohort, while 35 upregulated and 28 downregulated miRNAs were exclusive to the fusion-negative group (Figure 6D; Supplementary Table S6).

### miRNA Target Prediction and Construction of Regulatory Networks in Fusion-Positive BLCA

To understand the functional implications of DEMs, we predicted their mRNA targets and constructed miRNA–mRNA regulatory networks. For the fusion-positive group, we identified 1013 downregulated target genes corresponding to 22 upregulated DEMs, and 400 upregulated targets for 29 downregulated DEMs (Supplementary Table S7). The resulting regulatory networks are shown in Figure 7A and 7B.

**Figure 7.**
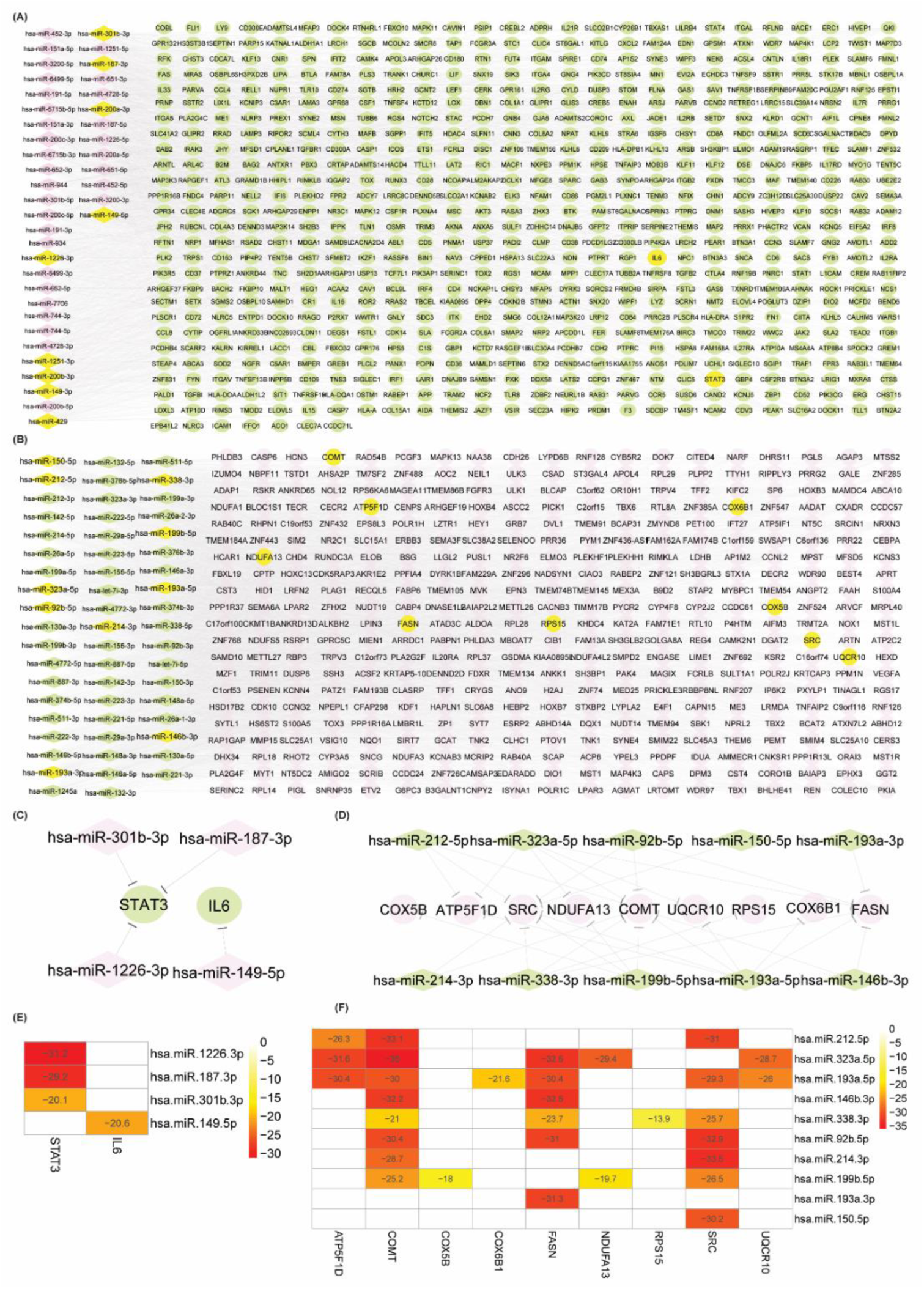
miRNA–mRNA regulatory networks in fusion-positive BLCA. (A–B) Networks of upregulated and downregulated DEMs with their respective targets. (C–D) Hub miRNAs targeting hub genes from PPI analysis. (E–F) Heatmaps showing minimum free energy (MFE) of validated interactions. Node shapes represent molecules: circles = genes; diamonds = miRNAs. Color scheme: pink = upregulated; green = downregulated.

Hub miRNAs were identified using degree centrality. Notably, four upregulated miRNAs— hsa-miR-149-5p, hsa-miR-187-3p, hsa-miR-301b-3p, and hsa-miR-1226-3p—were found to target two key downregulated hub genes (STAT3, IL6) previously identified in the PPI network (Figure 7C). Conversely, ten downregulated hub miRNAs—including hsa-miR-323a-5p, hsa-miR-199b-5p, hsa-miR-338-3p, hsa-miR-214-3p, hsa-miR-193a-5p, hsa-miR-193a-3p, hsa-miR-150-5p, hsa-miR-146b-3p, hsa-miR-212-5p and hsa-miR-92b-5p— targeted nine upregulated hub genes (COMT, COX6B1, NDUFA13, COX5B, ATP5F1D, RPS15, FASN, SRC, and UQCR10) (Figure 7D).

Minimum free energy (MFE) validation using RNAhybrid confirmed strong miRNA–mRNA interactions, excluding only RPS15 and COX5B, which did not meet the -17 kcal/mol threshold. The interaction affinities are visualized via heatmaps in Figure 7E and 7F, indicating robust miRNA-mediated regulatory potential within the fusion-positive cohort.

### Regulatory Network of DEMs in Fusion-Negative BLCA and Target Validation

For fusion-negative BLCA, we identified 767 downregulated targets of 35 upregulated miRNAs and 688 upregulated targets of 28 downregulated miRNAs (Supplementary Table S7; Figure 8A, D). Ten upregulated miRNAs were prioritized as hubs based on network centrality, including hsa-miR-21-5p, hsa-miR-1270, hsa-miR-130a-3p, hsa-miR-1293, hsa-miR-92a-3p, hsa-miR-142-3p, hsa-miR-137-3p, hsa-miR-147b-3p, hsa-miR-522-3p and hsa-miR-766-5p. These targeted key downregulated hub genes such as PPARGC1A, PPARG, PPARA, CEBPA, and KIT (Figure 8B).

**Figure 8.**
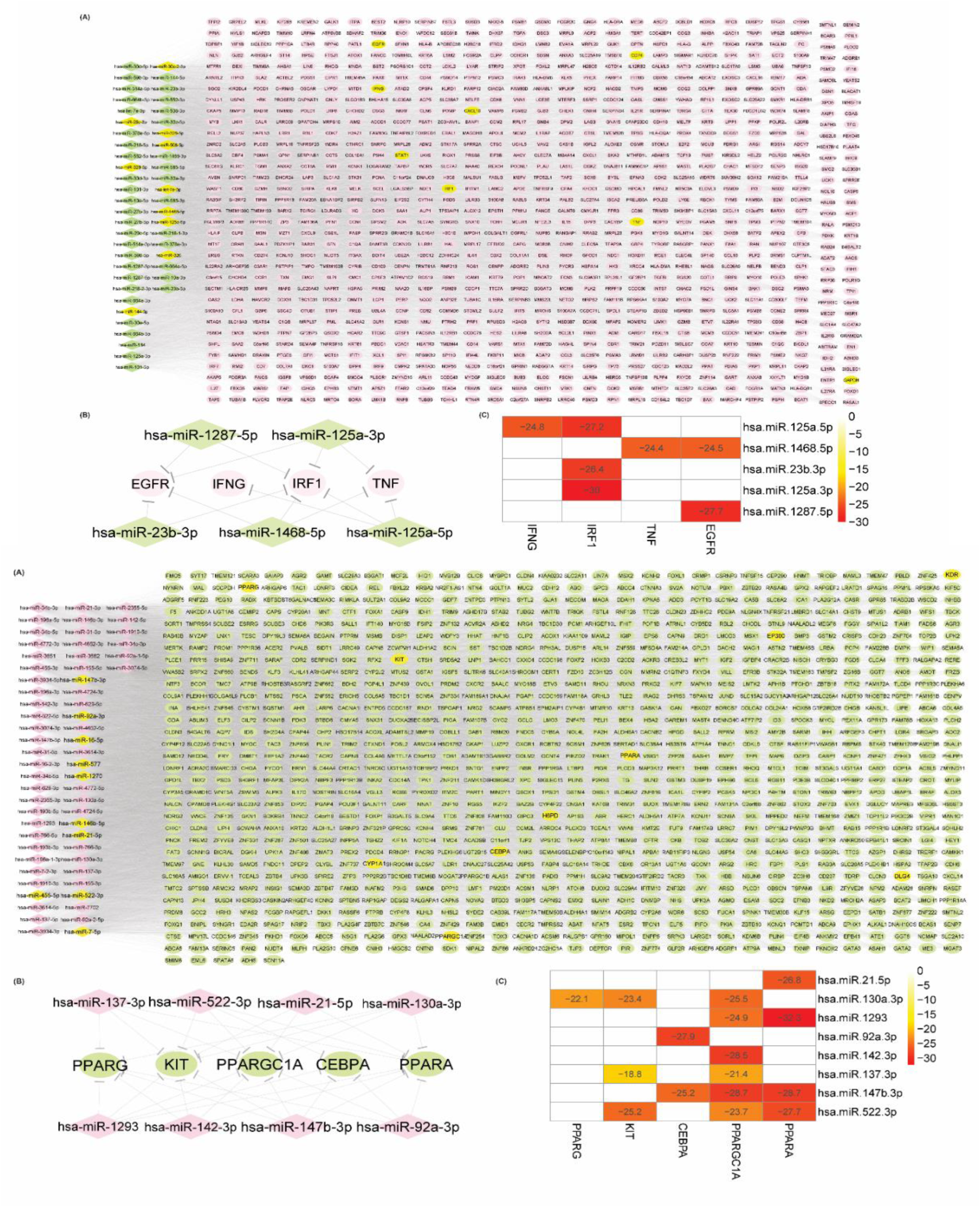
miRNA–Gene Regulatory Network Analysis in Fusion-Negative BLCA **Upper Panel:**(A) Network depicting interactions between upregulated differentially expressed miRNAs (DEMs) and their downregulated differentially expressed target genes (DETs). (B) Hub miRNAs (upregulated) targeting downregulated hub genes identified in the PPI network, illustrating key regulatory relationships. (C) Heatmap representing the binding affinity of upregulated miRNAs to their downregulated targets, quantified by minimum free energy (MFE, kcal/mol). Lower MFE values indicate stronger binding. **Lower Panel:** (A) Network illustrating interactions between downregulated DEMs and their upregulated DETs. (B) Hub miRNAs (downregulated) targeting upregulated hub genes from the PPI network (C) Heatmap showing MFE-based binding affinities between downregulated miRNAs and their upregulated gene targets. In all panels, elliptical nodes represent genes, and rectangular nodes represent miRNAs. Color coding: pink indicates upregulation, and green denotes downregulation.

In contrast, ten downregulated hub miRNAs—including hsa-miR-125a-5p, hsa-miR-144-3p, hsa-miR-1468-5p, hsa-miR-23b-3p, hsa-miR-218-5p, hsa-miR-508-3p, hsa-let-7e-5p, hsa-miR-125a-3p, hsa-miR-10a-5p and hsa-miR-1287-5p —targeted four upregulated hub genes: IFNG, IRF1, TNF, and EGFR (Figure 8E). MFE-based validation revealed particularly strong binding affinities. For instance, PPARGC1A was targeted by six hub miRNAs, and PPARG by hsa-miR-130a-3p with an MFE of –22.1 kcal/mol. IRF1 was targeted by three miRNAs, and IFNG by one, all showing high-affinity interactions (Figure 8C, F).

### Regulatory Network and Target Mapping of miRNAs Interacting with FGFR3 and TACC3 in Fusion-Positive BLCA

Given the biological relevance of the FGFR3–TACC3 fusion in BLCA, we investigated miRNAs directly targeting FGFR3 and TACC3. A total of 205 miRNAs were predicted to target FGFR3 and 22 for TACC3. After validating their expression in the fusion-positive cohort via miRNA-Seq data, three FGFR3-targeting miRNAs (hsa-miR-1226-3p, hsa-miR-6715b-5p, hsa-miR-6499-5p) were found to be overexpressed exclusively in fusion-positive BLCA.These miRNAs collectively targeted 165 downregulated genes, including two PPI-derived hub genes—STAT3 and CD4. Meanwhile, six TACC3-targeting miRNAs were associated with 68 upregulated gene targets, including the upregulated hub gene SRC, targeted by hsa-miR-483-5p (Figure 9).

**Figure 9.**
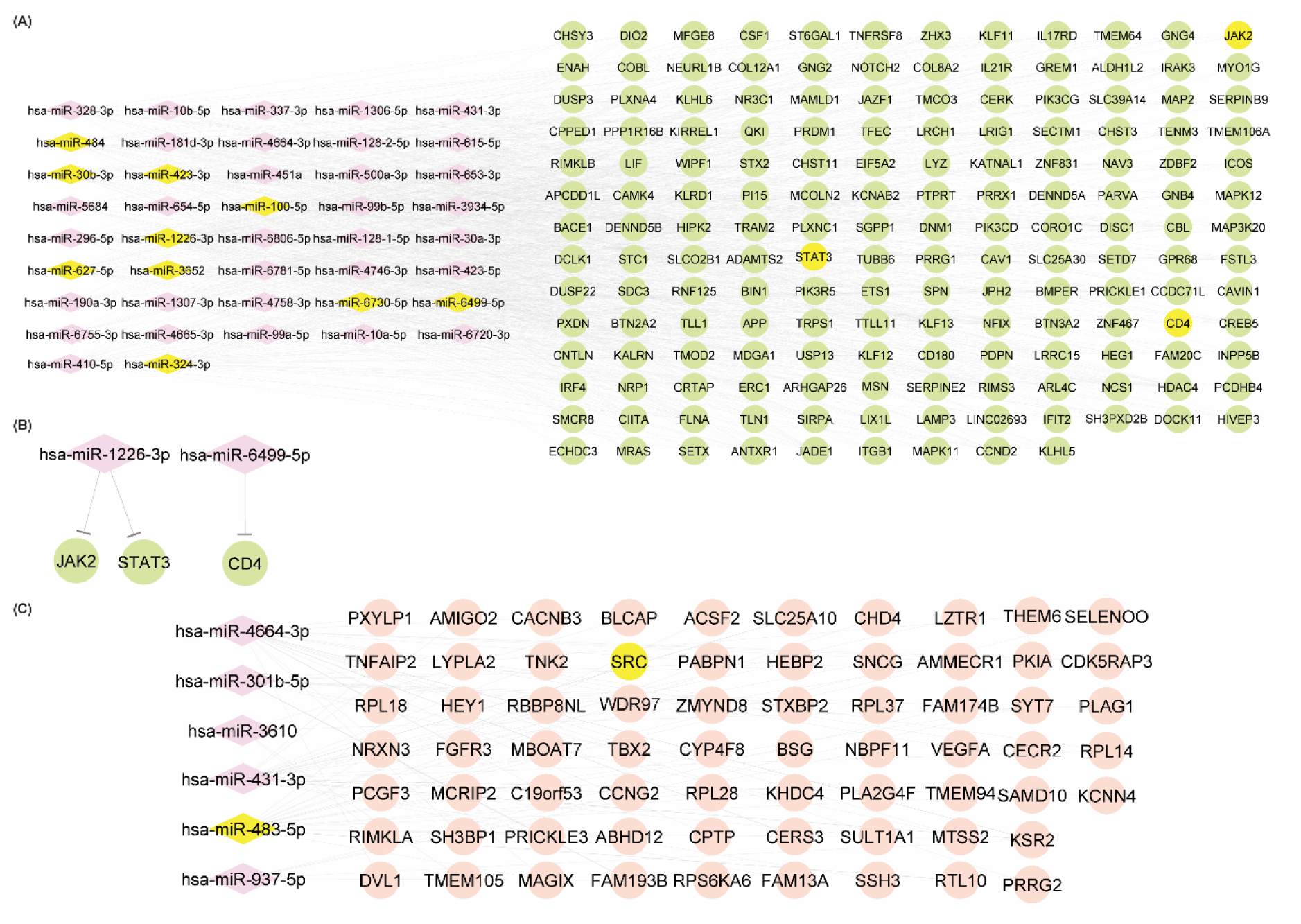
miRNA–Gene Regulatory Network Targeting FGFR3 and TACC3 in Fusion-Positive BLCA.(A) Regulatory network showing FGFR3-targeting miRNAs and their predicted downregulated gene targets in fusion-positive bladder cancer. (B) Subnetwork highlighting FGFR3-targeting hub miRNAs that interact with three key downregulated hub genes (STAT3, CD4, and IL6) identified from the PPI analysis. (C) Network of TACC3-targeting miRNAs and their associated upregulated gene targets. The miRNA hsa-miR-483-5p and its upregulated hub target SRC are highlighted in yellow to emphasize their functional significance in the fusion-positive regulatory context.

Pathway enrichment analysis of these gene targets showed that FGFR3-related targets were significantly involved in T cell receptor signaling, JAK-STAT signaling, PI3K-Akt, focal adhesion, and pathways in cancer, while TACC3-associated gene targets were enriched in tyrosine kinase inhibitor resistance and bladder cancer pathways (Table 7). Using RNAhybrid, we validated miRNA–mRNA duplexes based on MFE. Notably, hsa-miR-1226-3p–STAT3 and hsa-miR-6499-5p–CD4 interactions demonstrated strong thermodynamic affinity (MFE < –17 kcal/mol), highlighting their potential functional relevance in modulating oncogenic signaling in FGFR3–TACC3 fusion-positive BLCA.

**Table 7.**
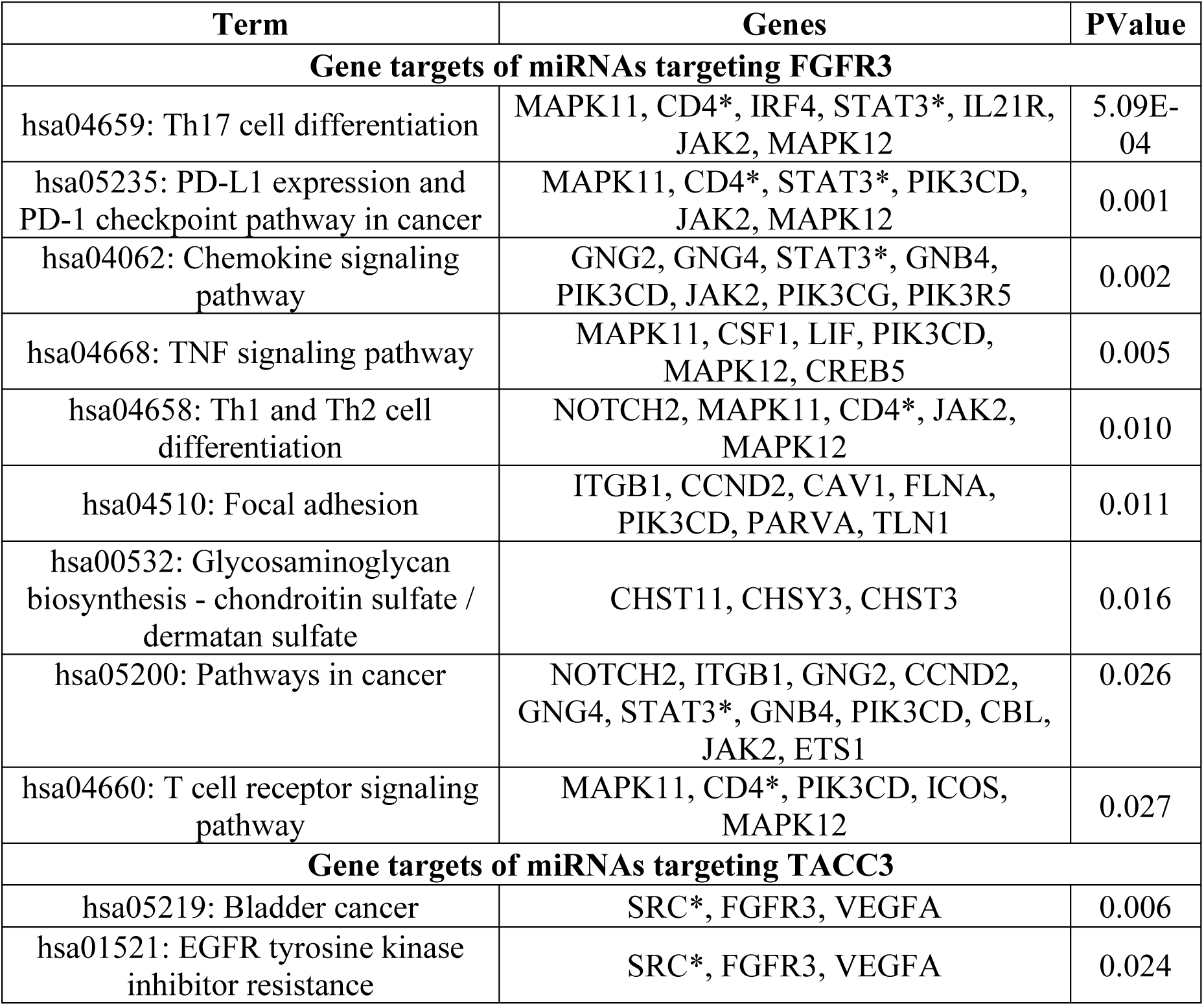
Pathway enrichment analysis of predicted gene targets of miRNAs targeting FGFR3 and TACC3. Includes significantly enriched KEGG pathways for downregulated and upregulated gene targets, respectively.

Following the construction of miRNA–gene regulatory networks, we identified key differentially expressed miRNAs (DEMs) that directly targeted hub genes in both fusion-positive and fusion-negative BLCA contexts. These miRNAs demonstrated strong potential as central post-transcriptional regulators within the dysregulated molecular landscape. The curated lists of these miRNA–hub gene pairs are detailed in Table 8 (fusion-positive vs fusion-negative) and Table 9 (fusion-negative vs normal).

**Table 8.**
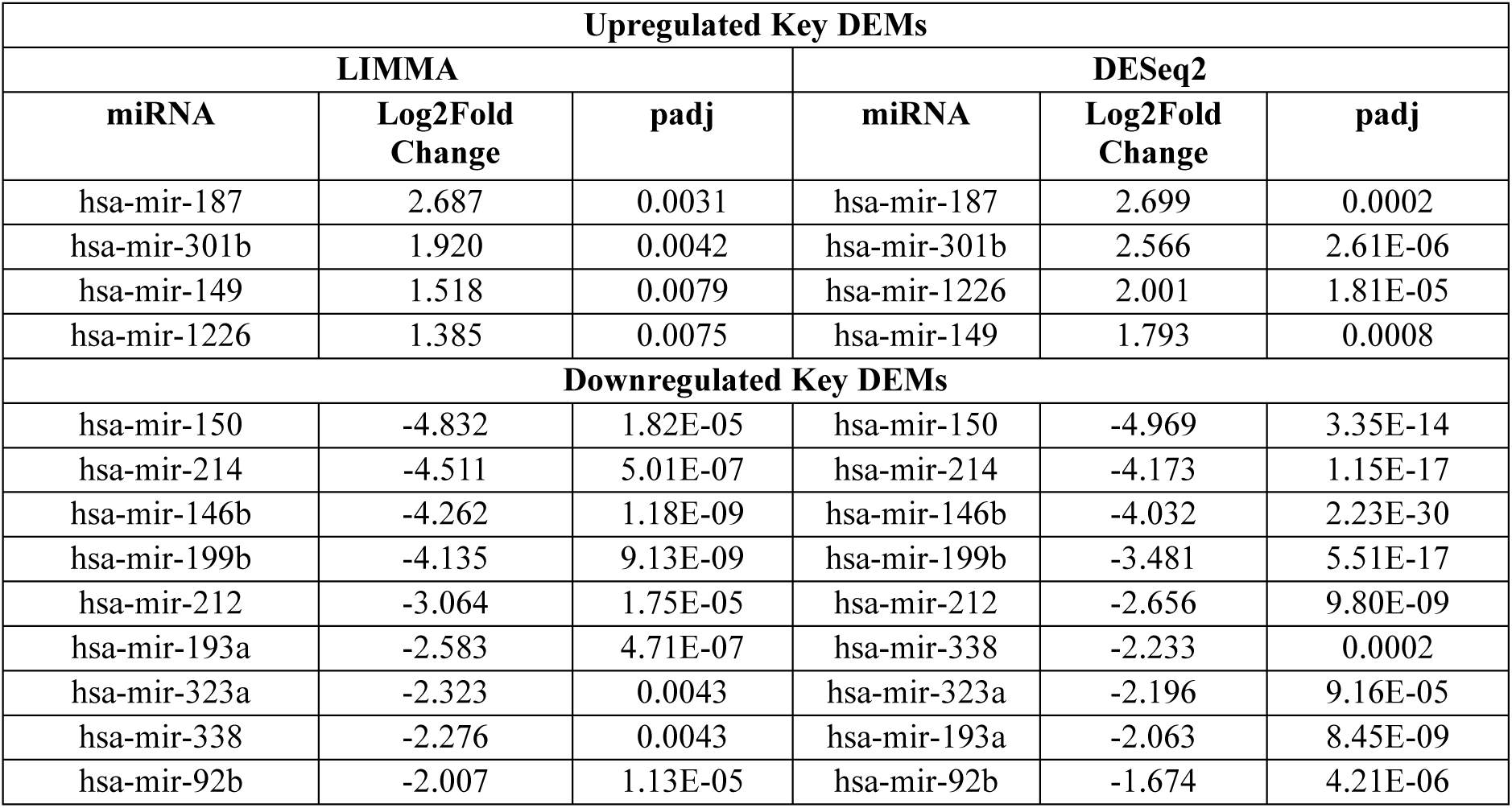
List of key differentially expressed miRNAs (DEMs) targeting hub genes in F3–T3 fusion-positive vs fusion-negative bladder cancer.

| Upregulated Key DEMs |  |  |  |  |  |
| --- | --- | --- | --- | --- | --- |
| LIMMA |  |  | DESeq2 |  |  |
| miRNA | Log2Fold Change | padj | miRNA | Log2Fold Change | padj |
| hsa-mir-187 | 2.687 | 0.0031 | hsa-mir-187 | 2.699 | 0.0002 |
| hsa-mir-301b | 1.920 | 0.0042 | hsa-mir-301b | 2.566 | 2.61E-06 |
| hsa-mir-149 | 1.518 | 0.0079 | hsa-mir-1226 | 2.001 | 1.81E-05 |
| hsa-mir-1226 | 1.385 | 0.0075 | hsa-mir-149 | 1.793 | 0.0008 |
| Downregulated Key DEMs |  |  |  |  |  |
| hsa-mir-150 | -4.832 | 1.82E-05 | hsa-mir-150 | -4.969 | 3.35E-14 |
| hsa-mir-214 | -4.511 | 5.01E-07 | hsa-mir-214 | -4.173 | 1.15E-17 |
| hsa-mir-146b | -4.262 | 1.18E-09 | hsa-mir-146b | -4.032 | 2.23E-30 |
| hsa-mir-199b | -4.135 | 9.13E-09 | hsa-mir-199b | -3.481 | 5.51E-17 |
| hsa-mir-212 | -3.064 | 1.75E-05 | hsa-mir-212 | -2.656 | 9.80E-09 |
| hsa-mir-193a | -2.583 | 4.71E-07 | hsa-mir-338 | -2.233 | 0.0002 |
| hsa-mir-323a | -2.323 | 0.0043 | hsa-mir-323a | -2.196 | 9.16E-05 |
| hsa-mir-338 | -2.276 | 0.0043 | hsa-mir-193a | -2.063 | 8.45E-09 |
| hsa-mir-92b | -2.007 | 1.13E-05 | hsa-mir-92b | -1.674 | 4.21E-06 |

**Table 9.** List of key differentially expressed miRNAs (DEMs) targeting hub genes in fusion-negative vs normal bladder cancer.

| Upregulated Key DEMs |  |  |  |  |  |
| --- | --- | --- | --- | --- | --- |
| LIMMA |  |  | DESeq2 |  |  |
| miRNA | Log2Fold Change | padj | miRNA | Log2Fold Change | padj |
| hsa-mir-1293 | 4.195 | 0.0019 | hsa-mir-1293 | 6.554 | 1.13E-09 |
| hsa-mir-147b | 3.960 | 3.54E-06 | hsa-mir-147b | 5.301 | 6.37E-11 |
| hsa-mir-522 | 3.237 | 0.0148 | hsa-mir-522 | 4.932 | 1.18E-05 |
| hsa-mir-142 | 2.142 | 3.12E-05 | hsa-mir-137 | 3.129 | 0.0007 |
| hsa-mir-21 | 1.722 | 2.13E-06 | hsa-mir-142 | 2.364 | 1.63E-10 |
| hsa-mir-137 | 1.668 | 0.0492 | hsa-mir-21 | 1.854 | 1.37E-14 |
| hsa-mir-130a | 1.536 | 0.0011 | hsa-mir-130a | 1.797 | 1.10E-06 |
| hsa-mir-92a | 1.090 | 0.0008 | hsa-mir-92a | 1.355 | 1.76E-06 |
| Downregulated Key DEMs |  |  |  |  |  |
| hsa-mir-23b | -2.290538471 | 1.24E-07 | hsa-mir-23b | -2.142837575 | 5.59E-18 |
| hsa-mir-1468 | -1.766475028 | 2.88E-05 | hsa-mir-1468 | -1.670925202 | 1.10E-06 |
| hsa-mir-125a | -1.367948555 | 3.09E-05 | hsa-mir-1287 | -1.174763159 | 9.82E-07 |
| hsa-mir-1287 | -1.287600406 | 0.000143222 | hsa-mir-125a | -1.164072813 | 1.50E-05 |

### miRNA–mRNA Complexes Suggests Splicing-Competent Conformations

To gain mechanistic insights into the functional interactions of DEMs with their target mRNAs, we performed secondary and tertiary structure modeling of selected miRNA–mRNA pairs. The aim was to evaluate their structural compatibility with the RNA-induced silencing complex (RISC), particularly their capacity to be recognized and cleaved by human Argonaute 2 (HsAGO2).

### Fusion-Positive BLCA: Structural Modeling of High-Priority Interactions

Using RNAhybrid, we predicted the 2D structures and binding affinities for 31 influential miRNA–mRNA pairs (Supplementary Figure S1). Among these, 9 pairs displayed strong structural compatibility based on AlphaFold 3-derived scores: ipTM ≥ 0.8 and pTM ≥ 0.9, signifying confident inter-chain and model accuracy predictions, respectively (Table 10). Further protein-RNA docking revealed that seven miRNA–mRNA duplexes—ATP5F1D–hsa-miR-193a-5p, COMT–hsa-miR-338-3p, STAT3–hsa-miR-301b-3p, STAT3–hsa-miR-187-3p, JAK2–hsa-miR-1226-3p, COMT–hsa-miR-193a-5p, and CD4–hsa-miR-6499-5p—were spatially localized within the catalytic cleft of HsAGO2. These duplexes engaged with key HsAGO2 domains (N, L1, PAZ, L2, MID, and PIWI) and occupied the central groove of the protein, forming a splicing-competent conformation [56] (Figure 10). Analysis of the 6 Å region of the duplex revealed precise electrostatic interactions between the RNA phosphate backbone and positively charged residues in the N domain of AGO2—namely, Lys62, Lys65, Arg68, Arg69, Arg72, Arg97, Lys98, Lys124, and Arg126. These interactions likely facilitate anchoring and alignment within the catalytic site, thereby enabling endonucleolytic cleavage.

**Figure 10.**
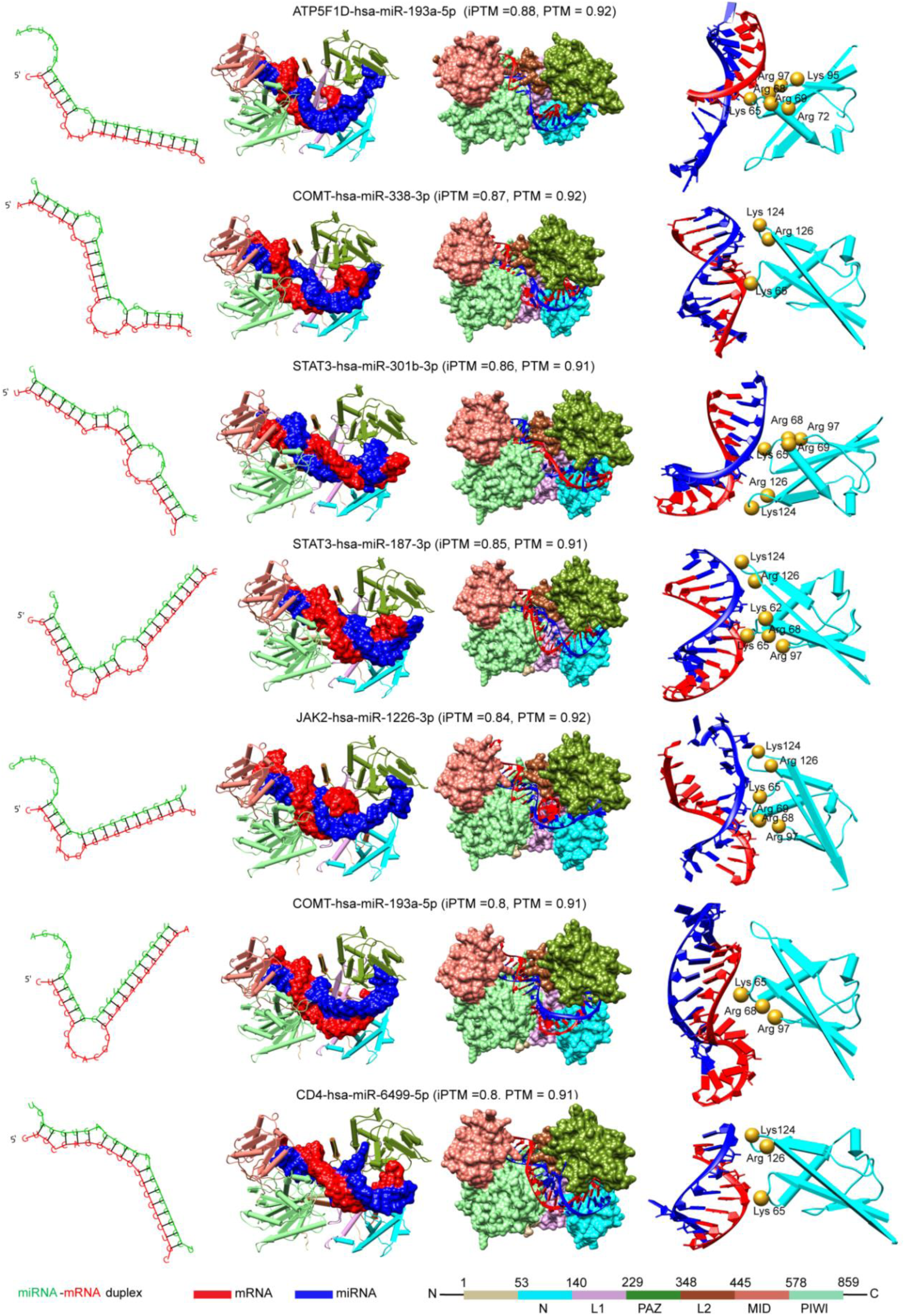
Structure prediction of critical miRNA-mRNA pairs in fusion positive BLCA cases. The figure depicts 2D secondary structure of each miRNA–mRNA duplex predicted by RNAhybrid, 3D structure of the duplex complexed with HsAGO2 in a fully paired conformation, Close-up visualization of interactions with the N-domain of HsAGO2, showing engagement with residues critical for slicing.

**Table 10.** Prediction scores (ipTM and pTM) for 3D structures of high-confidence miRNA–mRNA complexes in fusion-positive BLCA cases. Only interactions with ipTM ≥ 0.8 and pTM ≥ 0.9 are reported.

| mRNA | miRNA | ipTM score | PTM score |
| --- | --- | --- | --- |
| ATP5F1D | hsa-miR-193a-5p | 0.88 | 0.92 |
| UQCR10 | hsa-miR-193a-5p | 0.88 | 0.92 |
| COX6B1 | hsa-miR-193a-5p | 0.87 | 0.91 |
| COMT | hsa-miR-338-3p | 0.87 | 0.92 |
| STAT3 | hsa-miR-301b-3p | 0.86 | 0.91 |
| STAT3 | hsa-miR-187-3p | 0.85 | 0.91 |
| JAK2 | hsa-miR-1226-3p | 0.84 | 0.92 |
| COMT | hsa-miR-193a-5p | 0.8 | 0.91 |
| CD4 | hsa-miR-6499-5p | 0.8 | 0.91 |
| COX5B | hsa-miR-199b-5p | 0.78 | 0.91 |
| COMT | hsa-miR-323a-5p | 0.77 | 0.91 |
| COMT | hsa-miR-92b-5p | 0.75 | 0.91 |
| SRC | hsa-miR-338-3p | 0.73 | 0.91 |
| ATP5F1D | hsa-miR-212-5p | 0.72 | 0.9 |
| COMT | hsa-miR-214-3p | 0.7 | 0.9 |
| SRC | hsa-miR-199b-5p | 0.64 | 0.9 |
| COMT | hsa-miR-199b-5p | 0.62 | 0.9 |
| IL6 | hsa-miR-149-5p | 0.61 | 0.87 |
| NDUFA13 | hsa-miR-323a-5p | 0.6 | 0.9 |
| STAT3 | hsa-miR-1226-3p | 0.59 | 0.89 |
| SRC | hsa-miR-214-3p | 0.57 | 0.89 |
| SRC | hsa-miR-150-5p | 0.56 | 0.9 |
| ATP5F1D | hsa-miR-323a-5p | 0.54 | 0.89 |
| SRC | hsa-miR-483-5p | 0.53 | 0.89 |
| SRC | hsa-miR-212-5p | 0.51 | 0.89 |
| COMT | hsa-miR-212-5p | 0.47 | 0.87 |
| SRC | hsa-miR-193a-5p | 0.45 | 0.88 |
| SRC | hsa-miR-92b-5p | 0.42 | 0.87 |
| NDUFA13 | hsa-miR-199b-5p | 0.4 | 0.88 |
| UQCR10 | hsa-miR-323a-5p | 0.39 | 0.87 |
| COMT | hsa-miR-146b-3p | 0.28 | 0.85 |

### Fusion-Negative BLCA: Structural Analysis of miRNA–mRNA Complexes

A similar computational framework was applied to the fusion-negative group, where 12 critical miRNA–mRNA interactions were modeled for 2D structure and binding affinity (Supplementary Figure S2). Of these, 50% of the pairs met the high-confidence structural threshold (ipTM ≥ 0.8, pTM ≥ 0.9) (Table 11). Docking results showed that four miRNA– mRNA duplexes—PPARGC1A–hsa-miR-130a-3p, PPARGC1A–hsa-miR-522-3p, PPARG– hsa-miR-130a-3p, and PPARGC1A–hsa-miR-142-3p—formed stable, splicing-competent configurations upon binding to HsAGO2 (Figure 11). These complexes engaged in electrostatic interactions with the same N-domain residues implicated in the fusion-positive group, supporting a conserved mechanism of RISC-mediated slicing. Notably, PPARGC1A emerged as a key hub target for multiple miRNAs across the fusion-negative cohort, reinforcing its regulatory vulnerability and potential role in the suppression of metabolic reprogramming.

**Figure 11.**
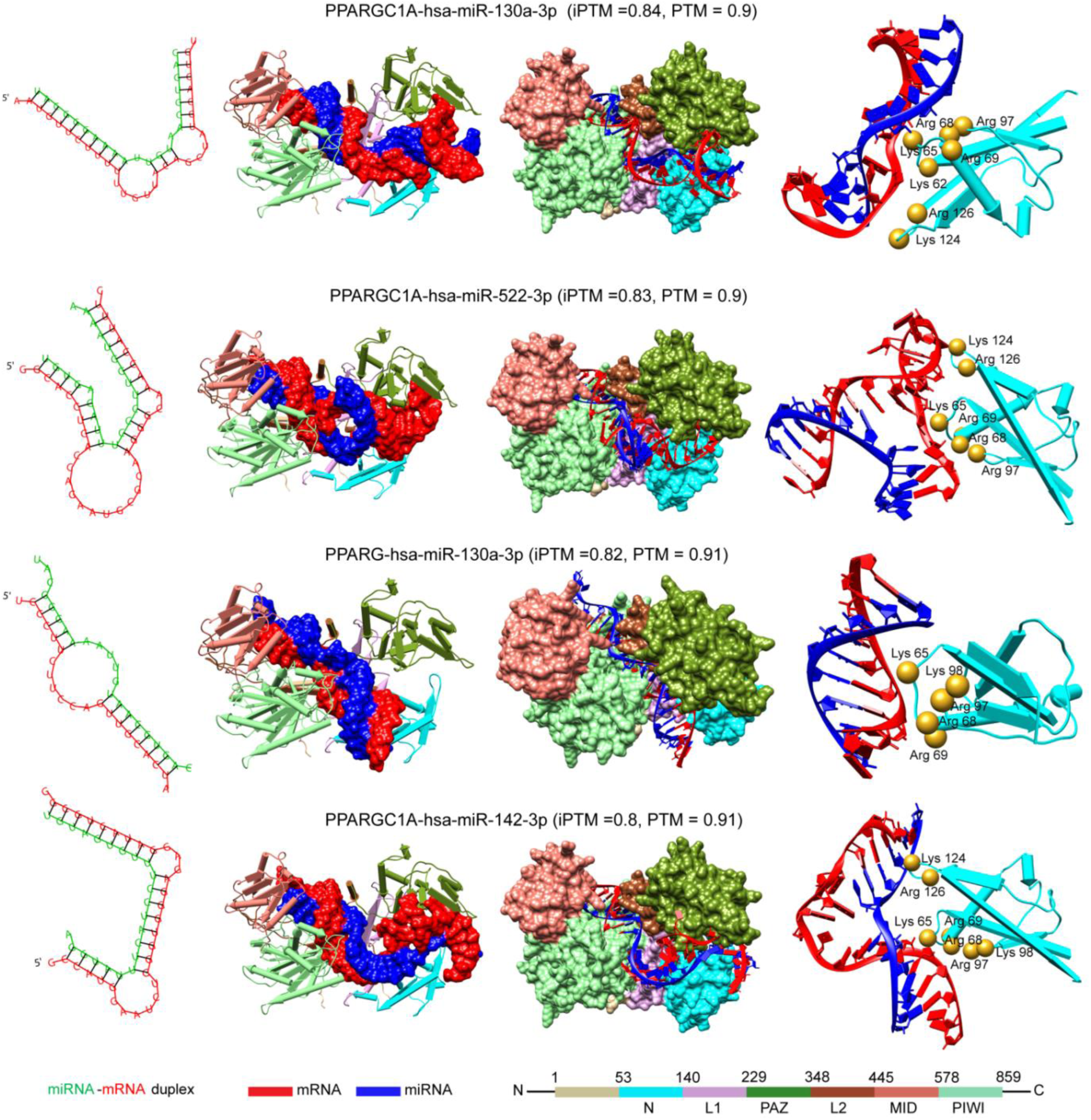
Structural modeling of miRNA–mRNA duplexes in fusion-negative BLCA. Displays analogous information to Figure 10, illustrating duplex formation, 3D AGO2 docking, and key residue interactions in the N-domain that mediate catalytic slicing.

**Table 11.** Prediction scores for 3D structures of influential miRNA–mRNA interactions in fusion-negative BLCA. Included are models predicted to be structurally competent for AGO2-mediated slicing [56].

| mRNA | miRNA | ipTM score | PTM score |
| --- | --- | --- | --- |
| IFNG | hsa-miR-125a-5p | 0.87 | 0.91 |
| CEBPA | hsa-miR-92a-3p | 0.85 | 0.88 |
| PPARGC1A | hsa-miR-130a-3p | 0.84 | 0.9 |
| PPARGC1A | hsa-miR-522-3p | 0.83 | 0.9 |
| PPARG | hsa-miR-130a-3p | 0.82 | 0.91 |
| PPARGC1A | hsa-miR-142-3p | 0.8 | 0.91 |
| PPARGC1A | hsa-miR-137-3p | 0.74 | 0.88 |
| CEBPA | hsa-miR-147b-3p | 0.68 | 0.9 |
| PPARGC1A | hsa-miR-1293 | 0.68 | 0.9 |
| PPARGC1A | hsa-miR-147b-3p | 0.62 | 0.9 |
| EGFR | hsa-miR-1287-5p | 0.56 | 0.87 |
| EGFR | hsa-miR-1468-5p | 0.49 | 0.88 |

### Quantitative Profiling of the Tumor Immune Microenvironment (TIME) in Fusion-Positive and Fusion-Negative BLCA

To explore the tumor immune microenvironment (TIME) in FGFR3–TACC3 (F3–T3) fusion-defined bladder cancer (BLCA), we quantitatively assessed stromal and immune cell infiltration in both fusion-positive and fusion-negative patient groups. ESTIMATE analysis was used to compute stromal scores, immune scores, and composite ESTIMATE scores across all BLCA samples.

Fusion-positive BLCA samples exhibited significantly lower stromal, immune, and ESTIMATE scores compared to both fusion-negative cases and normal bladder tissue. Interestingly, while fusion-negative cases also showed lower stromal and ESTIMATE scores relative to normal samples, their immune scores were markedly elevated (Figure 12). These observations suggest a divergence in immune microenvironment composition driven by F3– T3 fusion status.

**Figure 12.**
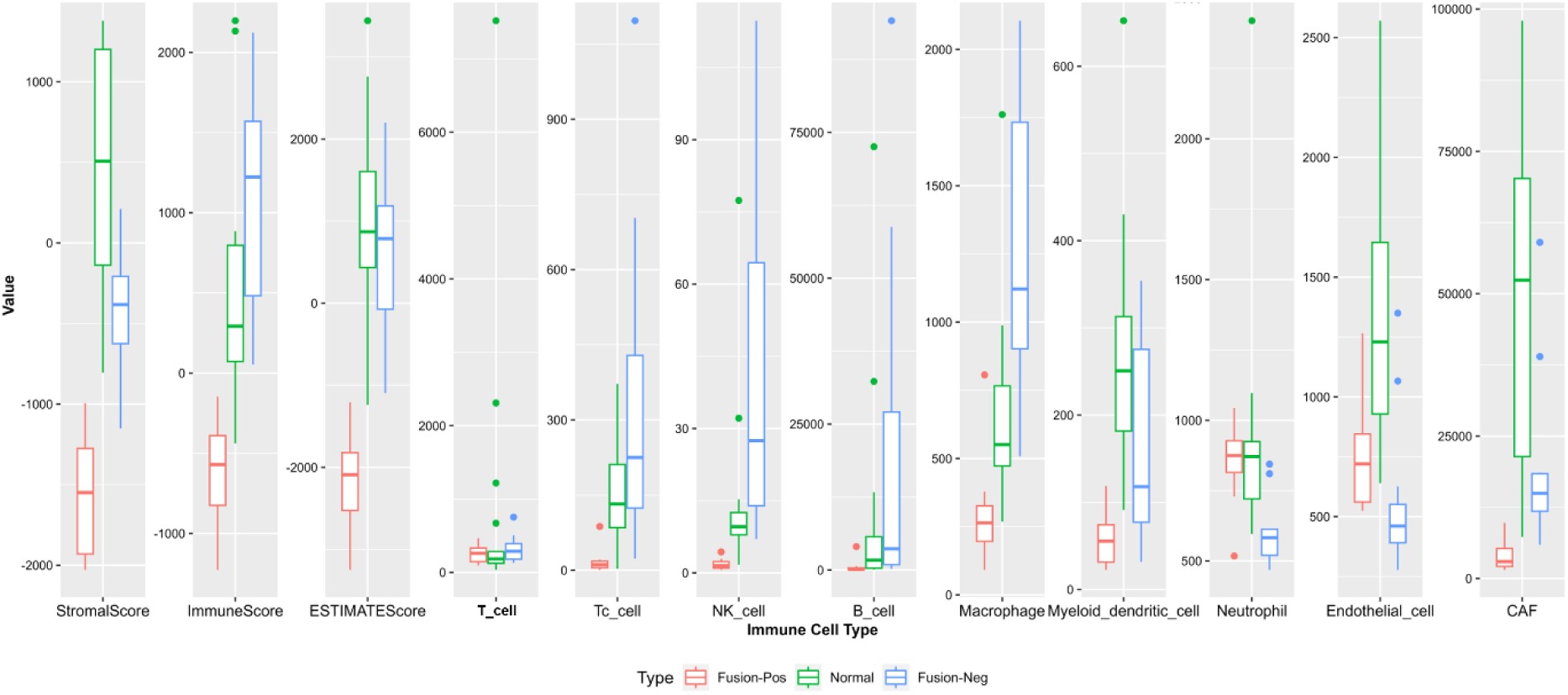
Boxplot showing the distribution of the ESTIMATE scores and tumor infiltration levels of nine immune cells infiltration status in the datasets. The data used to produce the boxplots is the normalized RNA expression data. Each boxplot is represented by the median and two hinges (representing the first and third quartiles).

### Immune Cell Composition Analysis Using MCP Counter Algorithm

To deconvolute the cellular makeup of the tumor stroma and immune compartments, we employed the MCP-counter algorithm, quantifying the relative abundance of nine major cell types: T cells, cytotoxic T cells (Tc), B cells, NK cells, macrophages, myeloid dendritic cells (mDCs), neutrophils, endothelial cells, and cancer-associated fibroblasts (CAFs).

Our results indicated a marked reduction in Tc cells, NK cells, B cells, macrophages, mDCs, and CAFs in fusion-positive samples relative to fusion-negative BLCA (Figure 12). Conversely, fusion-positive tumors showed elevated infiltration of neutrophils and endothelial cells. T cells showed no statistically significant difference across the fusion groups. This differential infiltration pattern underscores a distinct immunological landscape in the fusion-positive TIME, potentially reflective of an immune-evasive phenotype.

### Correlation Between ESTIMATE Scores and Immune Cell Types

To further understand the immune–stromal interplay, we conducted correlation analyses between the ESTIMATE-derived scores and the infiltration levels of immune cell types in each cohort. In fusion-positive BLCA, stromal, immune, and ESTIMATE scores showed moderate to strong inter-correlations (Pearson’s R = 0.55–0.90), while even stronger correlations were observed in fusion-negative BLCA (R = 0.76–0.93) (Figure 13A & Figure 14A).

**Figure 13.**
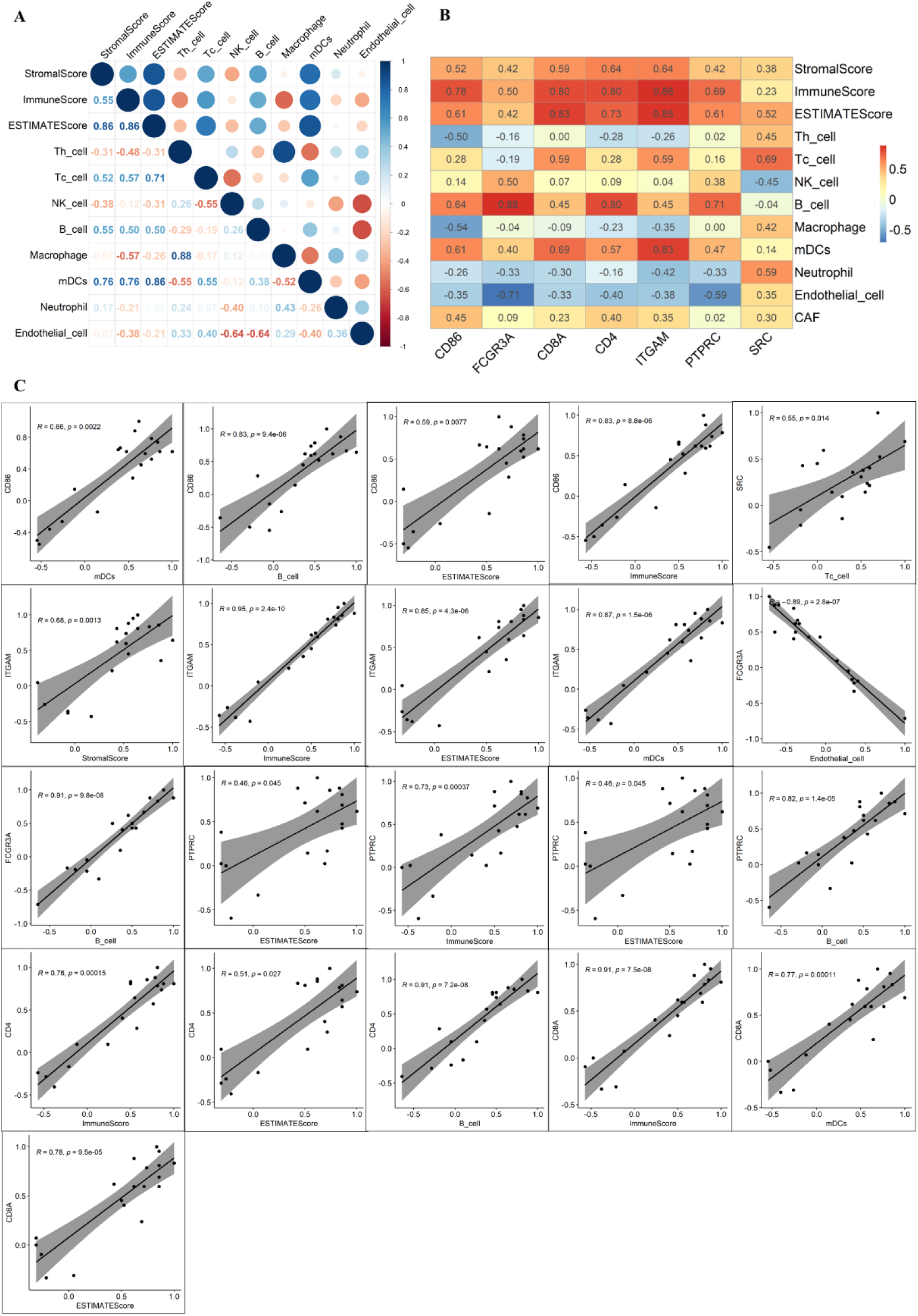
(A) Correlation analysis among the three ESTIMATE immune scores and infiltrated immunocytes for F3-T3 fusion positive BLCA. (B) Pheatmap showing correlation coefficients between the immunocytes and immune related influential genes. Red and blue color demonstrates positive and negative association, and darker and lighter shade shows higher and lower value of coefficient, respectively. (C) Scatter plots showing linear relationship between immune cells infiltration level versus immune cell associated gene expression in fusion-positive BLCA. X-axis represents cell infiltration level and Y-axis represents the gene expression for the associated immune cell.

**Figure 14.**
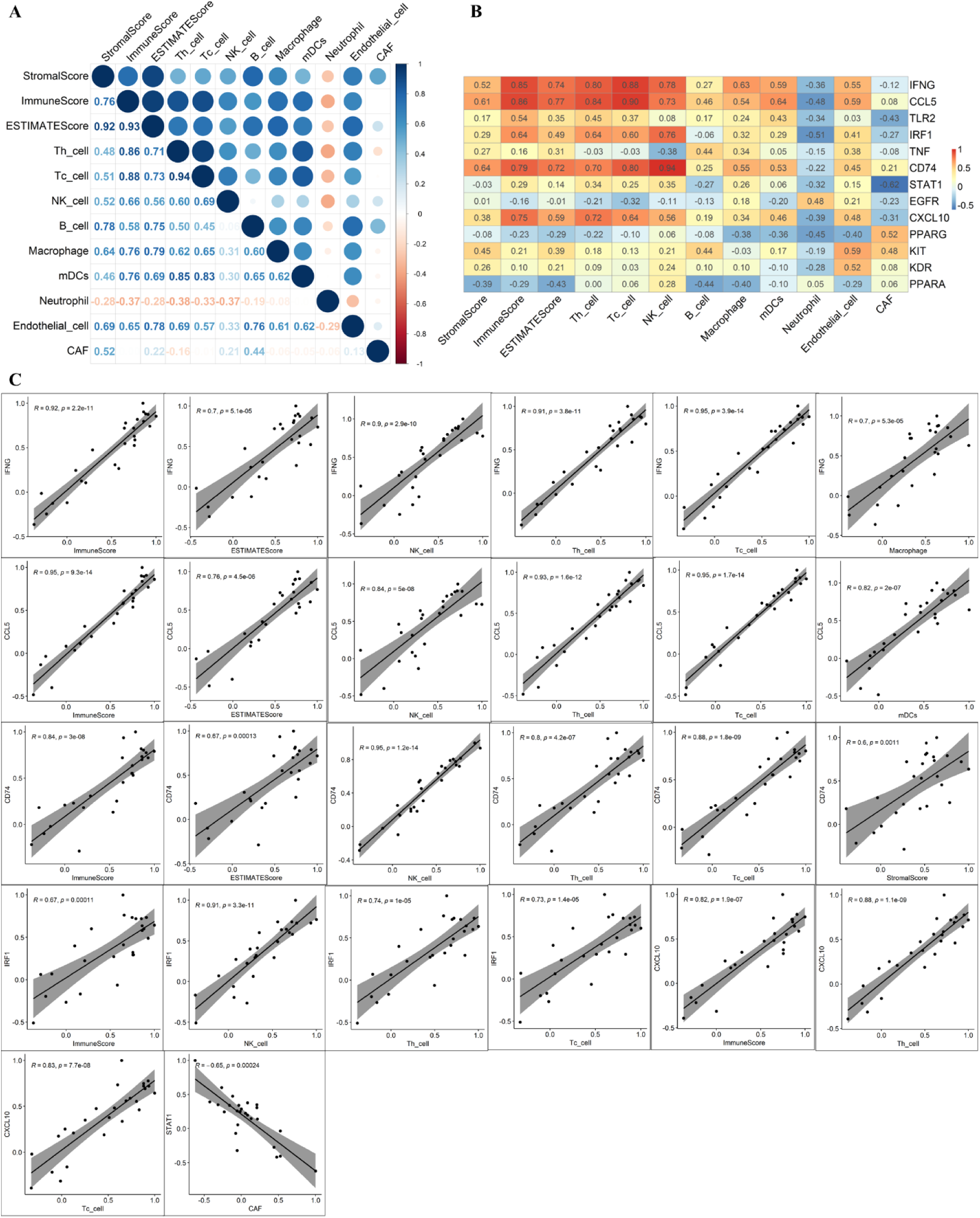
(A) Correlation analysis among the three ESTIMATE immune scores and infiltrated immunocytes for fusion-negative BLCA. (B) Pheatmap showing correlation coefficients between the immunocytes and immune related influential genes. Red and blue color demonstrates positive and negative association, and darker and lighter shade shows higher and lower value of coefficient, respectively. (C) Scatter plots showing linear relationship between immune cells infiltration level versus immune cell associated gene expression in fusion-negative BLCA. X-axis represents cell infiltration level and Y-axis represents the gene expression for the associated immune cell.

We next assessed inter-cellular correlations among immune cell subsets. Weak to moderate correlations were observed for most immune cell types in both fusion cohorts. Notably, T cells positively correlated with macrophages, mDCs were positively associated with CAFs, and endothelial cells negatively correlated with Tc and NK cells (Figure 13B & Figure 14B). These relationships further reflect the divergent immunoarchitectures of fusion-positive and fusion-negative tumors.

### Immune Gene Correlation and Functional Integration of TIME Signatures

To investigate immune gene involvement in the TIME, we retrieved 2,143 immune-related genes and cross-referenced these with our identified differentially expressed hub genes. Seven immune-related hub DEGs were detected in the fusion-positive BLCA cohort, and 13 in the fusion-negative cohort (Supplementary Table S8). In fusion-positive BLCA, ESTIMATE scores showed strong positive correlations with key immune genes such as CD4, ITGAM, CD86, CD8A, and PTPRC. Myeloid cell infiltration was significantly correlated with CD86, CD8A, and ITGAM, Tc cell infiltration was associated with SRC, and B cells showed positive correlations with CD86, FCGR3A, CD4, and PTPRC. Conversely, FCGR3A was negatively associated with endothelial cell infiltration. Upon scatter plot visualization, only gene–cell pairs with |R| ≥ 0.6 and p-value ≤ 0.05 were retained, excluding combinations such as ESTIMATEScore–CD86, –PTPRC, –CD4, and Tc cell–SRC due to lower significance (Figure 12C & Figure 13C). In contrast, fusion-negative BLCA revealed a more strongly correlated immune environment. T cells, Tc cells, NK cells, macrophages, mDCs, and endothelial cells all showed high mutual correlation, while neutrophils and CAFs displayed weaker associations. Importantly, several gene–cell pairs displayed strong positive correlations (R ≥ 0.7), including:

- StromalScore with CCL5, CD74
- ImmuneScore and ESTIMATEScore with IFNG, CD74, CCL5
- T and Tc cells with CXCL10, CD74, IRF1, CCL5, IFNG
- NK cells with CD74, IRF1, CCL5, IFNG
- Macrophages with IFNG
- mDCs with CCL5

Additionally, STAT1 exhibited a strong negative correlation with CAFs (R = –0.62). All retained gene–cell pairs demonstrated significant and linear associations on scatter plots, further supporting their biological relevance in TIME remodeling in fusion-negative BLCA.

### Integrative Network Modeling of Immune–Gene–miRNA Interactions in Fusion-Defined BLCA

To elucidate the interplay between immune regulation and gene expression in F3–T3 fusion-defined bladder cancer (BLCA), we developed integrative regulatory networks for both fusion-positive and fusion-negative cohorts. These networks link infiltrating immune/stromal cells with key tumor immune microenvironment (TIME)-associated hub genes and their regulatory microRNAs (miRNAs). Only those differentially expressed miRNAs (DEMs) that specifically target TIME-related differentially expressed genes (DEGs) were incorporated, with interaction confidence substantiated by high mirDIP prediction scores—asterisked to denote high-confidence miRNA–target pairs (Figure 15). In the fusion-positive network (Figure 15, left panel), we observed upregulated microRNAs including hsa-miR-1226-3p, hsa-miR-1251-5p, hsa-miR-1226-3p, hsa-miR-149-5p, hsa-miR-187-3p, hsa-miR-301b-3p, hsa-miR-6499-5p, hsa-miR-744-5p, hsa-miR-452-5p were predicted to target downregulated hub genes FCGR3A, CD4, PTPRC, CD8A, ITGAM. These immune-related genes are associated with elevated endothelial cells, and reduced infiltration of B cells, and myeloid dendritic cells (mDCs) suggesting that miRNA-mediated silencing contributes to an immunosuppressive TIME in fusion-positive BLCA. In contrast, the fusion-negative network (Figure 15, right panel) is characterized by the upregulation of immune-stimulatory genes including IRF1 and IFNG, which are negatively regulated by downregulated miRNAs such as hsa-miR-125a-5p, and has-miR-23-3p. These genes are linked to enhanced infiltration of Tc cells, and NK cells consistent with a more active and immunoreactive TIME. Altogether, these models emphasize distinct regulatory circuits underlying the TIME in fusion-positive versus fusion-negative BLCA. These findings highlight two distinct miRNA–gene–cellular interaction frameworks: one suppressive in fusion-positive BLCA, and one permissive to immune engagement in fusion-negative BLCA. Such context-specific regulatory architectures could inform the design of fusion-aware therapeutic strategies.

**Figure 15.**
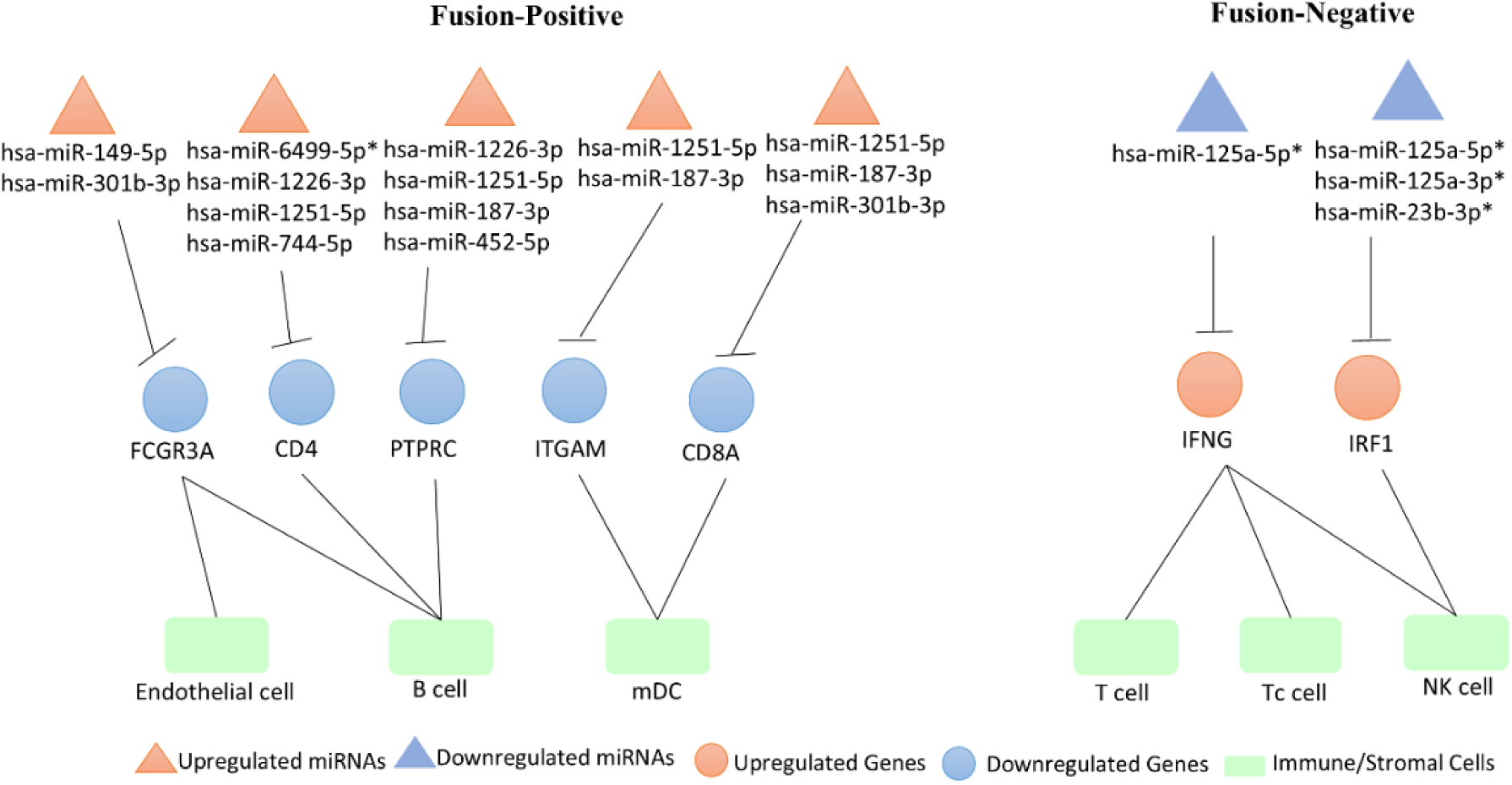
Integrative regulatory networks linking immune/stromal cells, TIME-associated hub genes, and their regulatory miRNAs in F3–T3 fusion-positive and fusion-negative BLCA.Only differentially expressed miRNAs (DEMs) targeting TIME-related hub DEGs are shown. High-confidence miRNA–target interactions predicted by miRDIP are marked with asterisks (*).

In an effort to unravel cross-pathway interactions, we focused on four key dysregulated functional pathways in fusion-positive BLCA: oxidative phosphorylation, cytokine signaling, chemokine signaling, and autoimmune response (Figure 16). To identify convergence points among these pathways, we reconstructed a protein–protein interaction (PPI) network using genes enriched in our GSEA (Figure 14). Remarkably, IFNGR2 (interferon gamma receptor 2) emerged as the sole connecting node bridging the upregulated oxidative phosphorylation module with the downregulated immune-related pathways. Expression analysis revealed a non-significant change in IFNGR2 levels but a log2 fold-change of -1.01 for IFNGR1, its receptor partner. Given that interferon gamma receptor (IFNGR) signaling is critical for JAK-STAT pathway activation, the observed downregulation likely contributes to deficient immune recruitment in fusion-positive tumors [129, 130]. This pattern was absent in fusion-negative BLCA, further reinforcing the subtype-specific immunological divergence.

**Figure 16.**
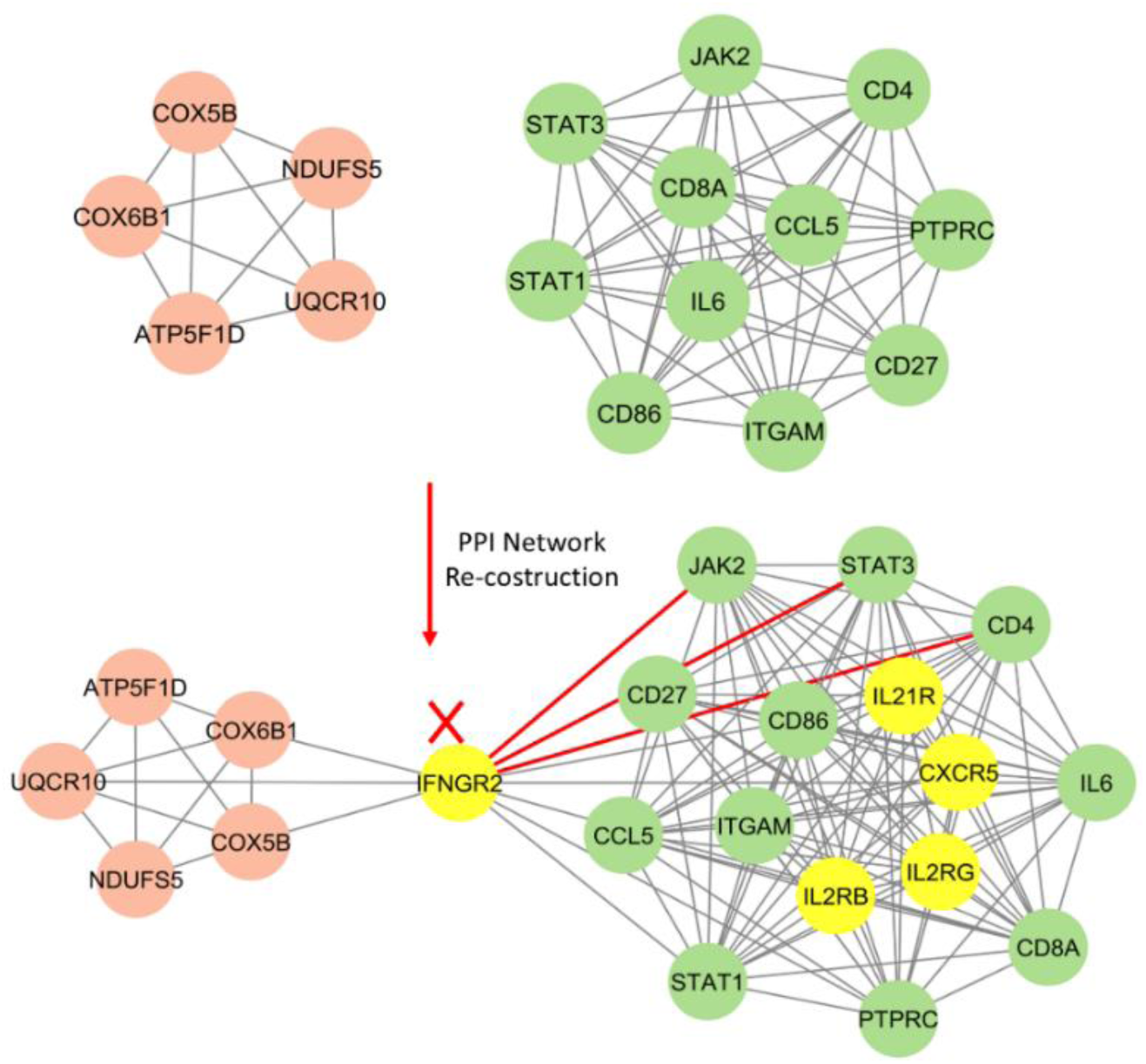
Protein-protein interaction network reconstruction of DEGs present in oxidative phosphorylation, cytokine signaling, chemokine signaling and autoimmune responses in fusion-positive BLCA. Upregulated DEGs are represented using red color nodes and downregulated DEGs are represented using green color nodes. Yellow nodes are the extra nodes added in the network using STRING database. Highlighted red color edges are showing the direct interaction between missing IFNGR2 and the downregulated JAK2, STAT3 and CD4 genes.

### Diagnostic Evaluation of Hub Genes and miRNAs via ROC Curve Analysis

To evaluate the diagnostic efficacy of fusion-specific gene signatures, ROC (Receiver Operating Characteristic) curve analysis was conducted using high-variance genes exclusive to the F3-T3 fusion-positive BLCA group. From the shortlisted set of 20 exclusive genes, 10 genes displayed an AUC (Area Under the Curve) value greater than 0.7 in the fusion-positive versus fusion-negative comparison, thereby indicating robust discriminative capability (Figure 17A). These included six upregulated genes—COMT, RPS15, COX6B1, FASN, SRC, and NDUFS5—and four downregulated genes—PTPRC, FCGR3A, CCL5, and STAT1.

**Figure 17.**
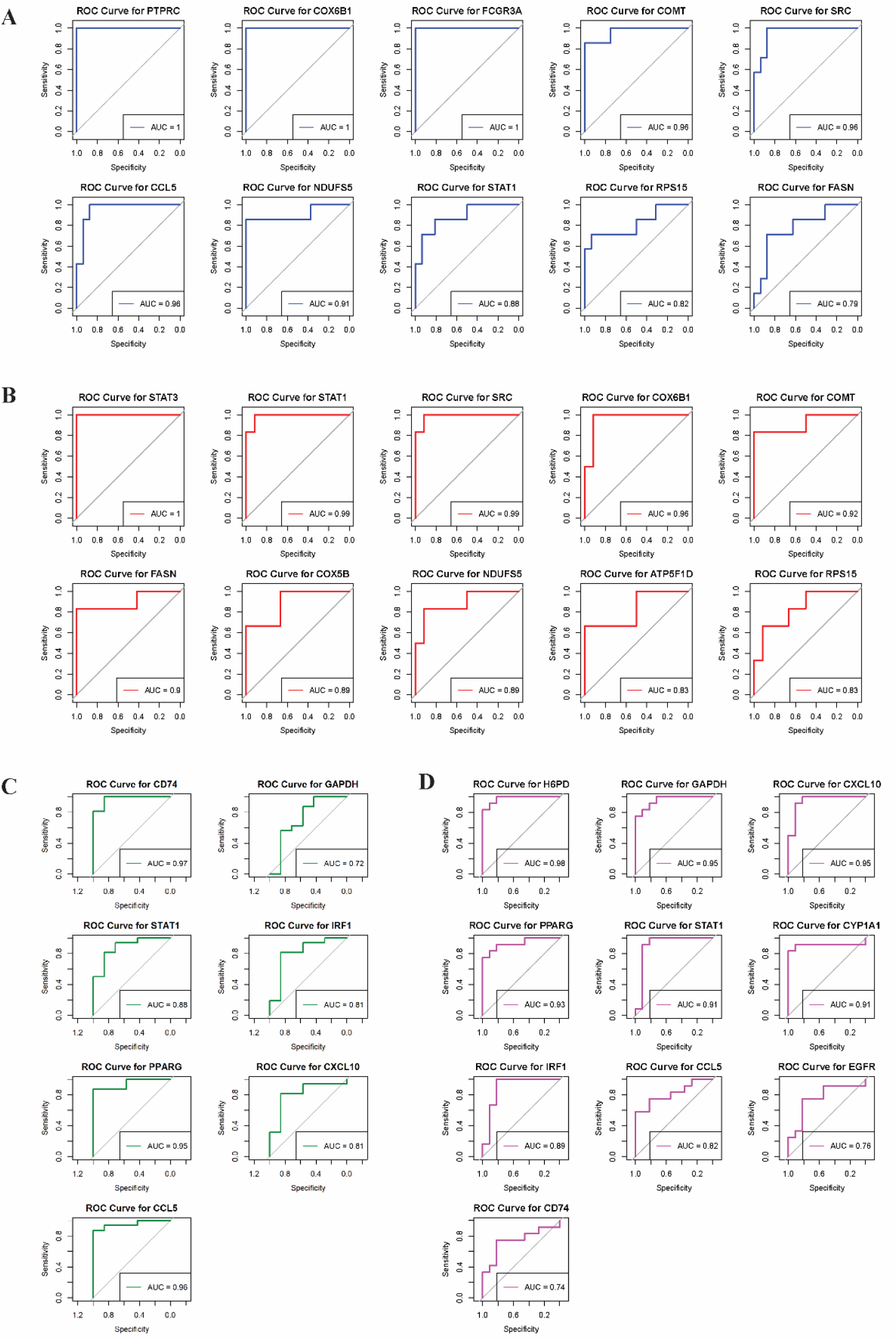
Receiver Operating Characteristic (ROC) curves and Area Under the Curve (AUC) values for identified hub genes in BLCA. This figure evaluates the diagnostic performance of hub genes with AUC > 0.7 as potential biomarkers in bladder cancer. (A–B) ROC curves for fusion-positive exclusive hub genes in fusion-positive vs. fusion-negative and fusion-positive vs. normal comparisons, respectively. (C–D) ROC curves for fusion-negative exclusive hub genes in fusion-negative vs. fusion-positive and fusion-negative vs. normal comparisons, respectively.

Notably, CCL5 and STAT1, which are known mediators of Th17 cell differentiation and chemokine signaling pathways, demonstrated exceptional predictive power, with AUC values of 0.96 and 0.88, respectively, for differentiating fusion-positive from fusion-negative cases. These results reinforce the role of immune dysregulation as a distinguishing molecular feature in fusion-driven BLCA. Moreover, comparison of fusion-positive samples against normal bladder tissue revealed that COMT, RPS15, COX6B1, FASN, SRC, NDUFS5, and STAT1 retained consistent AUC performance, suggesting their potential as diagnostic markers across both fusion-negative and healthy contexts (Figure 17B).

In parallel, fusion-negative exclusive DEGs were also evaluated. Among these, six upregulated genes—CD74, CCL5, IRF1, STAT1, and CXCL10—along with one downregulated gene, PPARG, exhibited high diagnostic potential (Figure 16C–D). CD74, in particular, demonstrated the strongest discriminative performance with an AUC of 0.97, followed by STAT1, PPARG, and other immune regulators.

### Diagnostic Performance of Fusion-Specific miRNAs

Next, we extended the ROC curve analysis to evaluate the diagnostic capability of differentially expressed miRNAs exclusive to fusion-positive and fusion-negative cohorts. In total, 14 fusion-positive exclusive miRNAs were analyzed, of which seven miRNAs exhibited high variance and AUC values exceeding 0.7 in the fusion-positive vs. fusion-negative group (Figure 18A). This panel included two upregulated miRNAs—hsa-miR-187 and hsa-miR-149—and five downregulated miRNAs—hsa-miR-193a, hsa-miR-146b, hsa-miR-338, hsa-miR-92b, hsa-miR-199b, and hsa-miR-150. All seven miRNAs also maintained high diagnostic accuracy (AUC > 0.7) in the fusion-positive versus normal comparison (Figure 18B), underscoring their robust classification ability in distinguishing fusion-positive patients from both fusion-negative and healthy cohorts.

**Figure 18.**
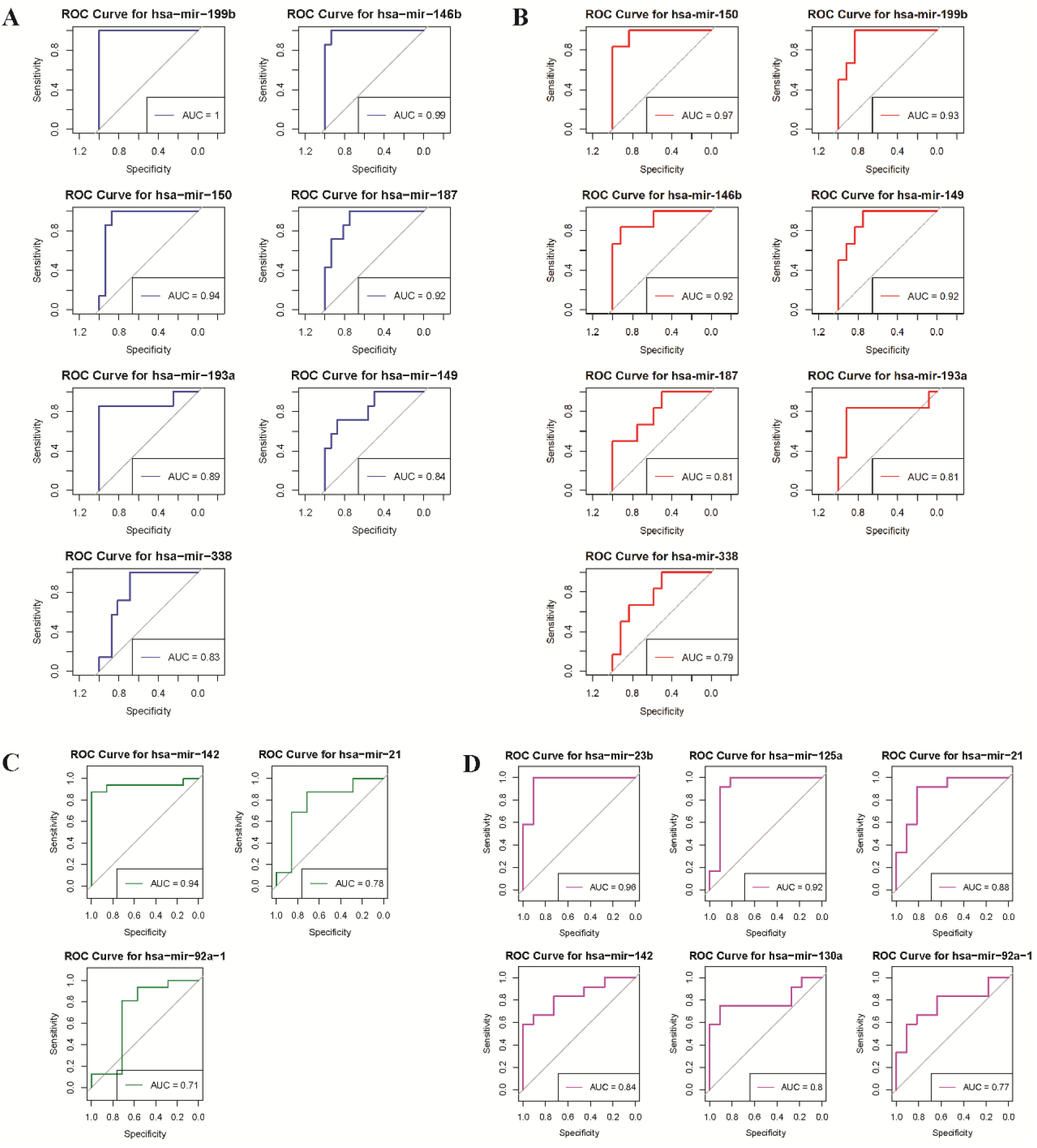
ROC curves and AUC values for identified key miRNAs as potential biomarkers in BLCA. This analysis evaluates the predictive utility of DEMs with AUC > 0.7. (A–B) Fusion-positive exclusive miRNAs in fusion-positive vs. fusion-negative and fusion-positive vs. normal comparisons. (C–D) Fusion-negative exclusive miRNAs in fusion-negative vs. fusion-positive and fusion-negative vs. normal comparisons.

Similarly, for fusion-negative exclusive DEMs, we identified hsa-miR-142, hsa-miR-21, and hsa-miR-92a as potential biomarkers with good predictive performance across both fusion-positive and normal comparisons (Figure 18C–D). These miRNAs have previously been implicated in immune regulation and tumor proliferation, providing additional biological support for their diagnostic relevance.

### Identification of Candidate Therapeutics through Connectivity Map (CMap) Analysis

To explore pharmacological molecules capable of reversing the disease-associated gene expression signatures, we employed the CMap query platform against upregulated DEGs from fusion-positive and fusion-negative BLCA modules. Molecules were prioritized based on negative normalized connectivity scores, indicating potential antagonistic effects. In the fusion-positive context, Lofexidine, an α2-adrenergic receptor agonist, was identified as a top-ranked molecule with inhibitory signature. Notably, the related cardiac muscle contraction pathway, associated with adrenergic signaling, was significantly dysregulated in the fusion-positive transcriptome, further supporting its mechanistic relevance.

For fusion-negative BLCA, two distinct compounds were identified: Lenalidomide, which targets TNF, and BX-795, an inhibitor of CDK2 and CHEK1—both genes being upregulated in fusion-negative samples, with CDK2 also localized in one of the top PPI network modules. The list of these candidate drugs, their mechanisms of action, molecular targets, and connectivity scores are summarized in Table 12.

**Table 12.**
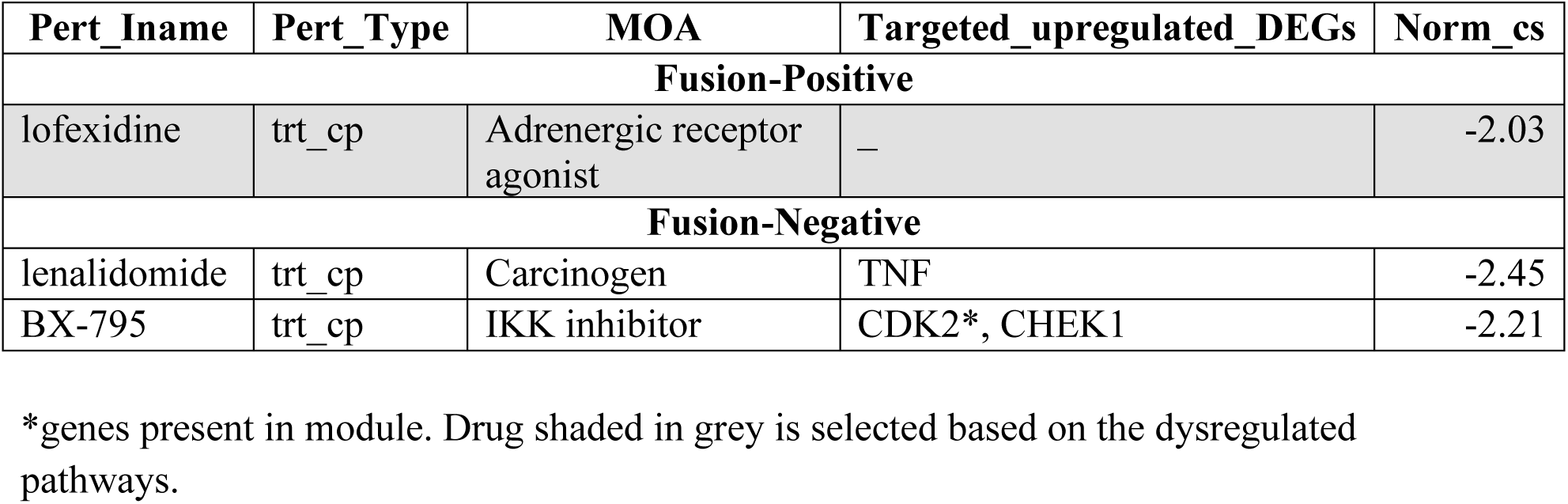
List of drugs obtained from CMap analysis based on normalised connectivity score, mechanism of action and drug targets.

| Pert_Iname | Pert_Type | MOA | Targeted_upregulated_DEGs | Norm_cs |
| --- | --- | --- | --- | --- |
| **Fusion-Positive** |  |  |  |  |
| lofexidine | trt_cp | Adrenergic receptor agonist | — | -2.03 |
| **Fusion-Negative** |  |  |  |  |
| lenalidomide | trt_cp | Carcinogen | TNF | -2.45 |
| BX-795 | trt_cp | IKK inhibitor | CDK2*, CHEK1 | -2.21 |

\*genes present in module. Drug shaded in grey is selected based on the dysregulated pathways.

## Discussion

In this study, we implemented an integrative comparative approach to better understand the unique transcriptional landscape associated with F3-T3 fusion-positive bladder cancer (BLCA). Unlike traditional analyses that contrast tumors with normal tissues alone, we independently compared F3-T3 fusion-positive tumors against both normal urothelium and fusion-negative BLCA. This dual comparison strategy allowed for a more precise delineation of transcriptional programs specific to the F3-T3 fusion event, disentangling them from broader tumorigenic alterations. Importantly, this approach enabled the identification of molecular features and potential therapeutic targets exclusive to the F3-T3 fusion subtype— features that are often obscured in standard tumor-versus-normal comparisons.

Bladder cancer (BLCA) is broadly classified into two histological categories: muscle-invasive bladder cancer (MIBC) and non-muscle-invasive bladder cancer (NMIBC), both of which exhibit distinct biological behaviors and treatment outcomes. These classifications are further supported by transcriptomic analyses [57, 58]. Additionally, molecular stratification has identified two major expression subtypes—luminal and basal. Notably, F3-T3 fusion events are predominantly associated with the luminal subtype of BLCA [14, 59]. In our analysis, we specifically addressed critical challenges in BLCA treatment, including poor prognosis, resistance to kinase inhibition, immune response variability, and tumor progression from low- to high-grade states—all of which are known to contribute to clinical heterogeneity and therapeutic failure.

Our findings revealed that genes upregulated in F3-T3 fusion-positive tumors were significantly enriched in oxidative phosphorylation and metabolic pathways, suggesting mitochondrial dysregulation. Mitochondria are central to ATP generation and thermogenesis, and their altered function may reflect the metabolic reprogramming characteristic of certain malignancies [60]. Gene Ontology (GO) enrichment analysis highlighted dysregulated mitochondrial metabolism in fusion-positive tumors, implying that targeted modulation of mitochondrial genes or miRNAs involved in oxidative phosphorylation may offer therapeutic benefit. We identified six upregulated hub differentially expressed genes (DEGs)— NDUFA13, NDUFS5, UQCR10, COX5B, ATP5F1D, and COX6B1—implicated in both oxidative phosphorylation and thermogenic processes in these tumors.

Furthermore, KEGG pathway enrichment analysis linked these same genes to cardiac-related pathways, including diabetic cardiomyopathy and cardiac muscle contraction. This is particularly relevant given that cardiac disease has been identified as the leading cause of long-term mortality in BLCA patients, surpassing the primary tumor in impact [61]. However, the biomechanical consequences of the F3-T3 fusion—especially in the context of cardiac comorbidities—remain insufficiently characterized and warrant further investigation.

Another compelling finding was the prominence of the ribosome pathway among the top dysregulated KEGG pathways in F3-T3 fusion-positive BLCA. Ribosomes are critical for protein synthesis, and alterations in ribosomal protein expression may reflect profound changes in cellular function and tumor progression. Specifically, RPS15, a component of the 40S ribosomal subunit, was found to be overexpressed in fusion-positive tumors. Elevated RPS15 expression has been linked to aggressive tumor phenotypes and poor prognosis in multiple cancers, including esophageal squamous cell carcinoma [62], colorectal cancer [63], and chronic lymphocytic leukemia [64]. Mechanistically, this may be mediated via the MDM2/MDM4 axis and dysregulation of TP53, which can exacerbate DNA damage responses and influence cell cycle control [65]. These results position RPS15 as a potentially actionable biomarker in the context of fusion-driven tumor heterogeneity in BLCA.

In parallel, we observed significant downregulation of several immune-related hub DEGs— CD86, IL6, STAT1, CCL5, and ITGAM—in fusion-positive tumors. These genes are enriched across a range of immune and cell adhesion-related pathways, including cell adhesion molecules, cytokine-cytokine receptor interaction, JAK-STAT signaling, chemokine signaling, TNF signaling, and Toll-like receptor signaling. Prior studies have shown that overexpression of FGFR3 in BLCA leads to resistance to cell–cell contact inhibition [66], a phenomenon that might be compounded by the downregulation of adhesion molecules such as CD86 and ITGAM observed in our dataset.

Immune modulatory pathways—particularly the cytokine-cytokine receptor axis and chemokine signaling—also featured prominently in our results. DEGs such as CCL5, IL6, STAT1, and CD4 are known to influence the tumor immune microenvironment (TIME), with established links to both prognosis and immunotherapeutic responses in BLCA [67, 68]. The dysregulation of these genes underscores the potential immune evasion mechanisms in F3-T3 fusion-positive tumors.

Interestingly, despite the oncogenic role often attributed to the JAK-STAT pathway, downregulation of this signaling cascade may also impair anti-tumor immunity, particularly by diminishing natural killer (NK) cell activity and enhancing metastatic potential [69]. In our study, four key genes involved in this pathway—CD86, IL6, STAT1, and CCL5—were downregulated, highlighting a possible immunosuppressive phenotype in fusion-positive BLCA. Additionally, Toll-like receptors (TLRs), which act as membrane-bound sensors of pro-inflammatory signals and mediate intracellular signaling leading to cytokine and co-stimulatory molecule production, are also implicated in the altered immune landscape of these tumors [70]. The complex interplay between TLR signaling and cytokine expression may act as a double-edged sword, either promoting or inhibiting tumor progression depending on the context.

Our construction of miRNA–gene regulatory network provided insights into the post-transcriptional landscape specific to F3-T3 fusion-positive BLCA. This network highlighted the interactions between several upregulated hub genes—ATP5F1D, COMT, COX5B, COX6B1, FASN, NDUFA13, RPS15, SRC, and UQCR10—and downregulated differentially expressed miRNAs (DEMs), including hsa-miR-323a-5p, hsa-miR-199b-5p, hsa-miR-338-3p, hsa-miR-193a-5p, hsa-miR-214-3p, hsa-miR-193a-3p, hsa-miR-150-5p, hsa-miR-146b-3p, hsa-miR-212-5p, and hsa-miR-92b-5p. These interactions underscore the potential regulatory complexity underlying the unique transcriptomic signature of fusion-positive BLCA.

In addition to these interactions, our study identified notable miRNA–gene regulatory pairs involving upregulated miRNAs and downregulated gene targets, including hsa-miR-1226-3p, hsa-miR-187-3p, and hsa-miR-301b-3p targeting STAT3, as well as hsa-miR-149-5p targeting IL6. These specific interactions point to functionally relevant regulatory circuits with potential translational implications. Notably, miR-149-5p has been shown to function as a tumor suppressor in BLCA, and its elevated expression correlates with poor overall survival, suggesting its utility as a non-invasive prognostic biomarker [71]. It also modulates IL6 expression through the LncRNA-LINC00460 axis in nasopharyngeal carcinoma [72].

Similarly, miR-1226-3p is implicated in oncogenic mechanisms in bladder urothelial carcinoma, nasopharyngeal carcinoma, and endometrial cancer [73-75], while miR-301b-3p, a member of the miR-130 family, promotes proliferation across various cancers including BLCA [76-78]. MiR-187-3p regulates the MAPK signaling pathway, which governs cell proliferation, survival, and is associated with poor BLCA outcomes [79, 80]. Meanwhile, miR-193a-3p—first identified as a tumor suppressor silenced via methylation during oral carcinogenesis [81]—has been reported as dysregulated [82], in lung cancer [83] and prostate cancer [84].

Other miRNAs in our network also display well-established tumor suppressor functions. MiR-338-3p, miR-323a-5p, and miR-212-5p inhibit proliferation and induce apoptosis, making them attractive prognostic markers in BLCA [85-87]. MiR-150-5p limits cancer aggressiveness in bladder, prostate, and head and neck squamous cell carcinomas [88-90], while downregulation of miR-146b-3p has been associated with BLCA progression [91]. MiR-199b-5p plays a tumor-suppressive role in renal and oral cancers and has been proposed as a prognostic marker in BLCA [92] [93], MiR-214-3p is also noteworthy for its association with improved survival outcomes in muscle-invasive BLCA [94]. Lastly, miR-92b-5p dysregulation has been reported in BLCA [95]. Despite substantial evidence of their oncogenic or tumor-suppressive roles, the contribution of these miRNAs to gene fusion-associated molecular mechanisms in BLCA remains underexplored.

To strengthen the credibility of our predicted miRNA–mRNA interactions, we incorporated minimum free energy (MFE) calculations, which confirmed the thermodynamic stability of several miRNA–target gene pairings. Collectively, these findings provide compelling evidence that dysregulated DEM–DEG pairs may serve as promising therapeutic targets, particularly for patients with F3-T3 fusion-positive tumors.

To contrast the fusion-driven molecular alterations, we also analyzed dysregulated DEGs and enriched pathways in fusion-negative BLCA. Interestingly, an inverse expression pattern was observed: DEGs upregulated in fusion-negative tumors were often enriched in pathways where those same genes were downregulated in fusion-positive tumors, and vice versa. For instance, pathways such as the NOD-like receptor signaling, allograft rejection, and Toll-like receptor signaling were enriched for upregulated DEGs STAT1, CXCL10, and IFNG in the fusion-negative cohort. Conversely, pathways such as chemical carcinogenesis–DNA adduct formation, xenobiotic metabolism via cytochrome P450, AMPK signaling, and broader cancer-related pathways were enriched among downregulated DEGs CYP1A1, PPARA, PPARG, PPARGC1A, and CEBPA.

The NOD-like receptor signaling pathway, when aberrantly activated, promotes immune evasion, angiogenesis, and chemoresistance in various cancers [88]. Toll-like receptors (TLRs) function as pattern recognition receptors, triggering innate immune responses against microbial and tumor-associated stimuli [96]. In our study, STAT1 was downregulated and associated with TLR signaling in fusion-positive tumors, yet upregulated and enriched in the same pathway in fusion-negative tumors. This suggests a dual, context-dependent role for STAT1. In the fusion-negative context, STAT1 may promote invasion and metastasis via JAK1/STAT1 activation.

CXCL10 also emerged as a potential immunomodulatory target in fusion-negative tumors. Known to influence angiogenesis and site-specific metastasis [97, 98], CXCL10 and IFNG upregulation may enhance T cell infiltration within the tumor immune microenvironment (TIME), aligning with our observations of increased cytotoxic T cell (Tc) infiltration in the fusion-negative cohort [99, 100].

Extending our analysis to miRNA–gene networks in fusion-negative BLCA, we identified several regulatory pairs involving upregulated DEMs and downregulated DEGs. Notable examples include hsa-miR-522-3p, miR-147b-3p, miR-1293, miR-21-5p targeting PPARA, and hsa-miR-522-3p, miR-147b-3p, miR-1293, miR-137-3p, miR-142-3p, miR-130a-3p targeting PPARGC1A. Other pairs included hsa-miR-147b-3p/92a-3p, hsa-miR-522-3p, miR-137-3p targeting KIT, and hsa-miR-130a-3p regulating PPARG. Conversely, several downregulated DEMs were predicted to target upregulated DEGs, such as hsa-miR-125a-5p/IFNG, hsa-miR-125a-3p, miR-23b-3p, miR-125a-5p/IRF1, hsa-miR-1287-5p, miR-1468-5p/EGFR, and miR-1468-5p/TNF. These interactions offer new insights into the differential miRNA–mRNA regulatory architecture distinguishing fusion-positive and fusion-negative BLCA subtypes.

Our analysis of the miRNA–gene regulatory network further identified several miRNAs with prognostic or functional relevance in fusion-negative BLCA. For instance, miR-522-3p overexpression is associated with poor survival and enhanced cell proliferation in multiple cancers, suggesting an oncogenic role [101]. In contrast, miR-147b-3p has been described as a pro-apoptotic miRNA, contributing positively to patient survival [102]. Notably, in our data, miR-147b-3p targets PPARGC1A, which was downregulated in fusion-negative BLCA. This gene plays a pivotal role in the AMPK signaling pathway, a pathway known to be suppressed via macrophage-mediated mechanisms, leading to mTORC1 activation and promoting tumor progression in BLCA [103]. In addition, hsa-miR-1293 has emerged as a potential predictive biomarker in cancers such as papillary renal cell carcinoma, lung adenocarcinoma, and pancreatic adenocarcinoma [104-106], whereas miR-21-5p has been proposed as a diagnostic tool specific for NMIBC in BLCA [107]. Moreover, miR-142-3p promotes the stemness of bladder cancer stem cells via PI3K/AKT pathway activation [108], and miR-130a-3p, a known serum diagnostic biomarker for BLCA [109], was upregulated in the fusion-negative cohort, indicating its potential role in defining subtype-specific miRNA signatures.

Tumor-suppressive miRNAs such as miR-125a-5p and miR-23b-3p are reported to inhibit tumorigenesis in hepatocellular carcinoma [110] and cervical cancer [111], respectively. Similarly, miR-1287-5p downregulation is associated with breast cancer progression [112], while miR-1468-5p is overexpressed in cervical cancer exosomes, impairing T-cell immunity by targeting lymphatic endothelial cells [113]. These findings reflect how dysregulated miRNA expression may orchestrate subtype-specific immune landscapes in BLCA.

We further examined PPARG, a nuclear receptor that shapes BLCA subtype and modulates immune exclusion. PPARG is typically downregulated in basal-type BLCA, which is often muscle-invasive [114]. Upon ligand binding, the PPARG/RXR heterodimer recruits co-activators such as PPARGC1A [115]. Differences in immune infiltration across BLCA subtypes—where basal and stroma-rich tumors exhibit high immune infiltration while luminal tumors show immune exclusion—reflect underlying transcriptional regulation and TIME architecture [116], PPARG, through NF-κB suppression and AP1 interaction, influences immune dynamics [117], and its downregulation may be associated with high immune infiltration in fusion-negative tumors.

Interestingly, EGFR overexpression, a hallmark of basal subtype BLCA, was also observed in our fusion-negative cases [118]. IRF1, a transcription factor expressed in all immune cells, and IFNG, a central cytokine in antitumor immunity, showed a strong correlation with NK, Tc, and T cell infiltration in fusion-negative tumors, consistent with previous findings [119] [120].

We then turned our attention to FGFR3 and TACC3-targeting miRNAs. In the context of FGFR3–TACC3 gene fusion, the 3’ untranslated region (3’UTR) of FGFR3 is disrupted, preventing canonical miRNA binding. Consequently, miRNAs previously targeting FGFR3 may now regulate alternative mRNA targets more effectively in fusion-positive BLCA. We identified three such FGFR3-targeting miRNAs—hsa-miR-1226-3p, hsa-miR-6715b-5p, and hsa-miR-6499-5p—that were upregulated in fusion-positive tumors. Among them, hsa-miR-1226-3p and hsa-miR-6499-5p targeted STAT3 and CD4, respectively, both of which were downregulated in fusion-positive BLCA. Furthermore, we found that TACC3-targeting miRNAs such as hsa-miR-483-3p regulated SRC, an upregulated hub gene in this subtype. These miRNA–gene axes were embedded within key pathways including PD-L1/PD-1 checkpoint, T cell receptor signaling, JAK-STAT, Th1/Th2/Th17 differentiation, EGFR inhibitor resistance, and primary immunodeficiency, suggesting that FGFR3 and TACC3 fusion events orchestrate broad transcriptional shifts in immune and oncogenic signaling.

Additionally, we noted other targets of hsa-miR-99a, a miRNA known to suppress FGFR3 by binding its 3’UTR. FGFR3 fusion renders this site unavailable, contributing to unchecked FGFR3 signaling and enhanced tumor progression, as previously reported [121]. Although immune escape in MIBC has been acknowledged [13, 122], the mechanistic basis remains elusive. Our findings suggest that deregulated miRNA–gene networks, particularly those involving FGFR3 and TACC3 fusions, may underlie this phenomenon. Specifically, the overexpression of miR-1226-3p and miR-6499-5p, in the absence of FGFR3 3’UTR binding sites, likely shifts their regulatory focus to alternate targets such as CD4, STAT3, and JAK2—genes involved in immune checkpoint signaling, Th17 cell differentiation, and the JAK-STAT pathway. Downregulation of these genes corresponds with reduced infiltration of CD8+ T cells, B cells, NK cells, macrophages, and myeloid dendritic cells in fusion-positive tumors, suggesting an immune-suppressive TIME. We propose a plausible molecular mechanism (see Figure 19) in which CD4 and STAT3 downregulation enhances PD-1 pathway activity, promoting T cell exhaustion and diminishing antitumor immunity—a protective mechanism against autoimmunity but detrimental in the cancer context [123, 124]. Additionally, STAT3 plays a crucial role in inducing IL-17 expression during Th17 differentiation. Its downregulation, alongside JAK2, may thus suppress Th cell lineage commitment, reinforcing immune escape in fusion-positive BLCA [125, 126].

**Figure 19.**
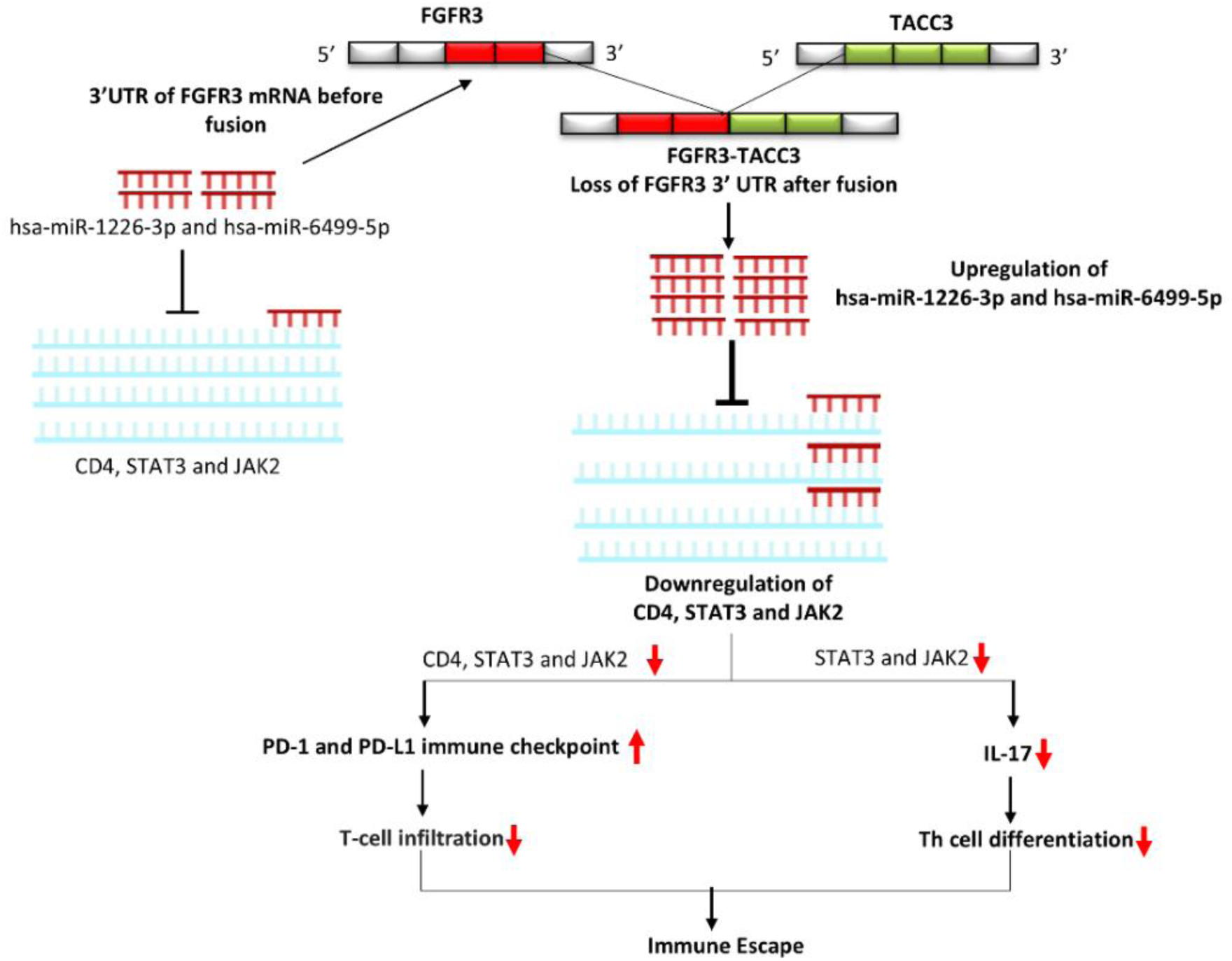
Proposed molecular mechanism of immune escape in F3–T3 fusion-positive BLCA. The figure illustrates how the deregulation of key immune genes and miRNA–mRNA interactions contribute to immune evasion, reduced immune cell infiltration, and T cell exhaustion in fusion-positive tumors.

Collectively, our analysis underscore the biological divergence between F3-T3 fusion-driven and non-fusion-driven BLCA, as reflected in the contrasting expression profiles, enriched signaling pathways, and miRNA–gene regulatory networks. These findings emphasize the necessity of subtype-specific therapeutic strategies and underscore the value of miRNA-mediated regulation as a potential avenue for personalized treatment in bladder cancer.

Furthermore, our analysis reveals a clear immune-excluded phenotype in F3-T3 fusion-positive BLCA, characterized by reduced infiltration of both immune and stromal cells relative to fusion-negative tumors and adjacent normal tissues. These findings offer a framework for understanding how gene fusions modulate the tumor immune landscape and potentially explain the variability in immune checkpoint blockade (ICB) responsiveness between molecular subtypes. This work lays the foundation for future efforts aimed at miRNA–mRNA interactions to stratify patients and design precision immunotherapeutic strategies tailored to fusion status in bladder cancer.

In line with previous findings, we corroborate that lower immune infiltration significantly contributes to poor prognosis in patients with F3-T3 fusion-positive BLCA [4]. This immunologically "cold" phenotype suggests an impaired anti-tumor immune response and underscores the necessity for immune profiling as a guiding framework for immunotherapeutic interventions. We identified six downregulated immune-related hub DEGs—FCGR3A, CD86, CD8A, CD4, ITGAM, and PTPRC—in fusion-positive BLCA. In contrast, fusion-negative tumors displayed upregulation of IFNG, CCL5, IRF1, CD74, CXCL10, and STAT1, collectively reflecting a more active tumor immune microenvironment (TIME).

The FCGR3A gene, encoding the Fc fragment of IgG receptor IIIa, has been proposed as a prognostic marker in prostate cancer and low-grade glioma. It plays a pivotal role in natural killer (NK) cell survival and proliferation by preventing progenitor cell apoptosis [127, 128]. In our study, the observed downregulation of FCGR3A was associated with reduced NK cell infiltration in fusion-positive BLCA, suggesting that FCGR3A may serve as a diagnostic biomarker reflecting immune depletion in this specific subtype.

To validate the immune relevance of other dysregulated DEGs, we explored the Human Protein Atlas, confirming associations between CD86, CD8A, CD4, PTPRC, and ITGAM with various immune cell types. These genes have established prognostic utility in BLCA. For example, CD86, CD4, and CD8A are frequently associated with better survival outcomes [131-133]. Moreover, CD86 was previously found to be downregulated in NMIBC, aligning with our current findings in fusion-positive BLCA [134]. PTPRC (also known as CD45) shows progressive upregulation with tumor stage, suggesting a dynamic role in BLCA progression [135], while ITGAM downregulation is implicated in impaired immune infiltration across multiple cancers, including osteosarcoma [136]. These patterns collectively point toward a functionally suppressed immune checkpoint activity in early or less inflamed fusion-positive tumors. Beyond transcriptomic profiling, we compiled experimental evidence supporting the biomarker potential of the identified DEGs and DEMs across multiple cancer types, including BLCA (see Table 13, Table 14, and Table 15). This evidence enhances the translational relevance of our findings and provides a curated resource for prioritizing targets for further functional studies.

**Table 13.**
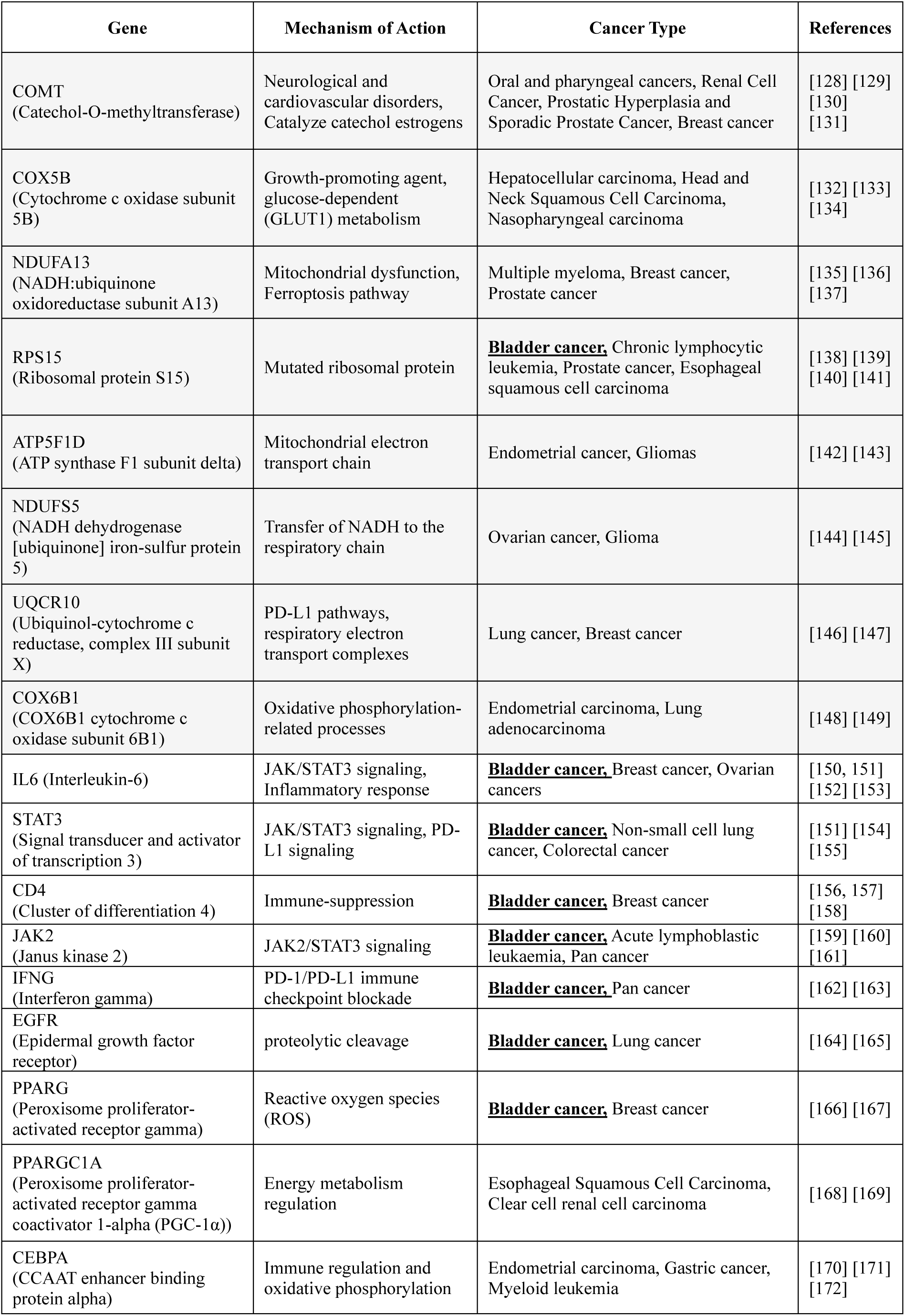
Influential genes as diagnostic/prognostic biomarkers in different cancer types.

**Table 14.**
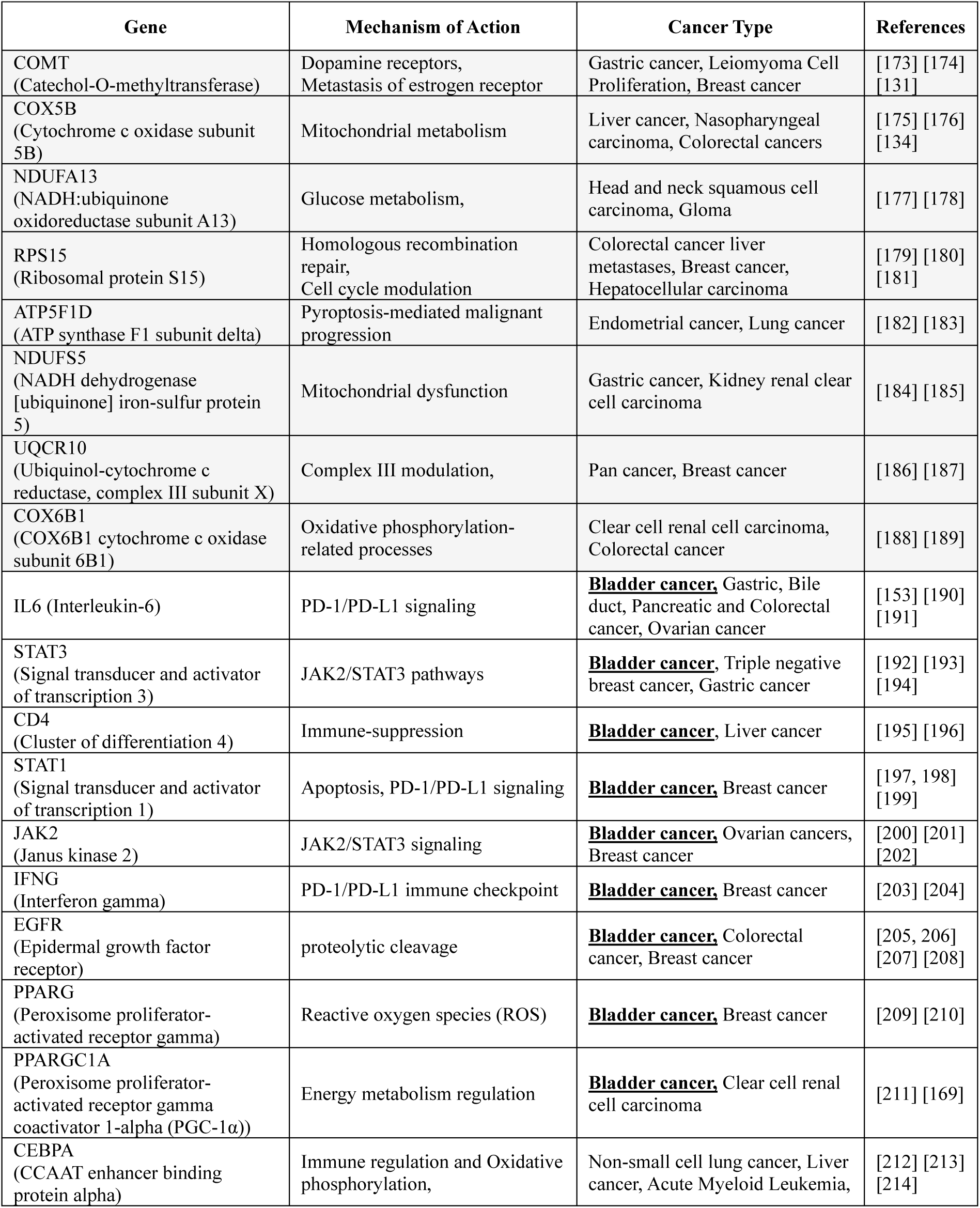
Influential genes as therapeutic targets in different cancer types.

**Table 15.** Influential miRNAs as diagnostic/prognostic biomarkers in different cancer types.

| miRNA | Cancer Type | References |
| --- | --- | --- |
| hsa-miR-323a | **<u>Bladder cancer</u>**, Glioblastoma, Esophageal squamous cell carcinoma | [215] [216] [217] |
| hsa-miR-212 | Breast cancer, Acute myeloid leukaemia (AML), Gastric cancer | [218] [219] [220] |
| hsa-miR-193a | **<u>Bladder cancer</u>**, Lung cancer, Thyroid cancer | [221] [222] [223] |
| hsa-miR-146b | **<u>Bladder cancer</u>**, Papillary thyroid cancer, Gallbladder carcinoma | [224, 225] [226] [227] |
| hsa-miR-338 | Renal cell carcinoma, Colorectal cancer, Glioma | [228] [229] [230] |
| hsa-miR-92b | Colorectal cancer, **<u>Bladder cancer</u>**, Hepatocellular carcinoma | [231] [232] [233] |
| hsa-miR-214 | **<u>Bladder cancer</u>**, Gastric cancer, Prostate cancer | [234, 235] [236] [237] |
| hsa-miR-199b | **<u>Bladder cancer</u>**, Parathyroid tumors, Hepatocellular carcinoma | [238] [239] [240] |
| hsa-miR-1226 | Pancreatic ductal adenocarcinoma, Colorectal cancer | [241] [242] |
| hsa-miR-187 | Prostate cancer, **<u>Bladder cancer</u>**, Oral carcinoma | [243] [244] [245] |
| hsa-miR-301b | **<u>Bladder cancer</u>**, Non-small cell lung cancer, Breast cancer | [246] [247] [248] |
| hsa-miR-149 | Corpus Endometrial Carcinoma, **<u>Bladder Cancer</u>**, Glioma | [249] [250] [251] |
| hsa-miR-1226 | Pancreatic ductal adenocarcinoma, Non-small cell lung cancer | [241] [252] |
| hsa-miR-125a | Cervical cancer, Urothelial Carcinoma, Non-small cell lung cancer, Prostate cancer | [253] [254] [255] [256] |
| hsa-miR-23b | **<u>Bladder cancer</u>**, Head and neck squamous cell, Colorectal cancer | [257] [258] [259] |
| hsa-miR-125a | Cervical cancer, Colon cancer, Breast cancer | [253] [260] [261] |
| hsa-miR-1287 | Colorectal cancer, Breast cancer | [262] [263] |
| hsa-miR-1468 | Non-small cell lung cancer, Hepatocellular carcinoma | [264] [265] |
| hsa-miR-130a | **<u>Bladder cancer</u>**, Colorectal cancer, Gastric cancer | [246] [266] [267] |
| hsa-miR-92a | **<u>Bladder cancer</u>**, Cholangiocarcinoma, Gastric cancer | [268] [269] [270] |
| hsa-miR-147b | Ovarian cancer, Hepatocellular carcinoma, Colorectal cancer | [271] [272] [273] |
| hsa-miR-1293 | Clear cell renal cell carcinoma, Lung adenocarcinoma | [274] [275] |
| hsa-miR-142 | Renal cell carcinoma, Prostate cancer, Colon cancer | [276] [277] [278] |
| hsa-miR-137 | Glioblastoma, Lung cancer, Pancreatic Cancer | [279] [280, 281] |
| hsa-miR-522 | Triple-negative breast cancer, Hepatocellular carcinoma, Gastric cancers | [282] [283] [284] |

To explore therapeutic implications, we employed Connectivity Map (CMap) analysis, which yielded distinct drug repurposing candidates for the two fusion-based BLCA subtypes. For fusion-positive tumors, we propose Lofexidine, an α2-adrenergic receptor agonist. Lofexidine lowers sympathetic stress responses, blood pressure, and heart rate [137]. intersecting with the cardiac muscle contraction pathway—a dysregulated axis in fusion-positive tumors identified via KEGG analysis. For fusion-negative cases, we suggest Lenalidomide and BX-795. Lenalidomide, known for its immunomodulatory, anti-inflammatory, and anti-angiogenic properties, has been shown to enhance the efficacy of BCG immunotherapy in NMIBC [138], [139, 140]. Notably, it targets TNF, a cytokine found upregulated in fusion-negative tumors. BX-795, a potent inhibitor of PDK1, CDK2, and CHEK1, also inhibits IKK and JNK/p38 pathways, showing anti-tumor activity in models such as neuroblastoma [141, 142]. Given that CDK2 and CHEK1 were upregulated in our fusion-negative BLCA cohort, BX-795 could represent a precision-targeted adjuvant therapy.

In conclusion, our study provides the first comprehensive transcriptomic and regulatory landscape distinguishing F3-T3 fusion-positive from fusion-negative BLCA, focusing on immune modulation, miRNA–mRNA interactions, and subtype-specific pathway dysregulation. We identified several miRNAs (e.g., hsa-miR-1226-3p, hsa-miR-6499-5p, hsa-miR-99a) and their respective gene targets (CD4, STAT3, JAK2, FGFR3, TACC3) that may serve as both mechanistic markers and therapeutic candidates. Particularly, the dysregulation of the PD-1/PD-L1 axis and JAK-STAT pathway in fusion-positive tumors appears to drive immune escape via reduced immune cell infiltration and T cell exhaustion. Finally, we emphasize that the distinct immune landscapes of fusion-positive and fusion-negative tumors should guide immunotherapy design, including the use of immune checkpoint inhibitors or combination regimens. While our study provides strong computational and literature-based support, experimental validation—both in vitro and in vivo—is imperative for clinical translation. Furthermore, future work may explore tumor heterogeneity and stage-specific analysis to refine these insights and enable precision oncology in bladder cancer.

## Conclusion

This study presents a comprehensive transcriptomic and miRNA regulatory dissection of FGFR3–TACC3 fusion-positive versus fusion-negative bladder cancer using integrative TCGA-derived RNA-Seq and miRNA-Seq data. We uncover two fundamentally distinct molecular subtypes: fusion-positive tumors exhibit pronounced metabolic reprogramming through the upregulation of oxidative phosphorylation and ribosomal genes (e.g., COX5B, ATP5F1D, RPS15), and simultaneous immune suppression, evidenced by the downregulation of core immune regulators (CD4, STAT3, IL6, CCL5) and depleted infiltration of cytotoxic T cells, NK cells, and antigen-presenting cells.

Mechanistically, these alterations are linked to fusion-specific miRNA–mRNA regulatory axes, with upregulated miRNAs such as miR-1226-3p and miR-149-5p targeting immune transcripts, and downregulated miRNAs (e.g., miR-199b-5p, miR-214-3p) losing repression over metabolic genes. Structural modeling confirmed the AGO2-compatible binding of 13 high-confidence miRNA–mRNA duplexes, validating their potential for post-transcriptional silencing.

In contrast, fusion-negative tumors show an immune-inflamed phenotype, characterized by elevated IFNG, CXCL10, IRF1 expression and robust immune cell infiltration, regulated by a distinct miRNA network. Immune cell–gene correlation analyses further demonstrated that immune architecture and functional polarization are tightly coupled to fusion status.

Receiver operating characteristic (ROC) analysis identified fusion-stratified diagnostic biomarkers with high discriminatory power (AUC > 0.9), including COMT, RPS15, and miR-187-3p for fusion-positive tumors, and CXCL10, CD74, and miR-142 for fusion-negative cases. Finally, Connectivity Map analysis suggested candidate therapeutics— including Lofexidine for fusion-positive and Lenalidomide/BX-795 for fusion-negative tumors—highlighting the translational relevance of the identified molecular programs.

Altogether, this study establishes that FGFR3–TACC3 fusion orchestrates a dual axis of immune evasion and metabolic activation in bladder cancer, mediated in part by context-specific miRNA–mRNA dysregulation. These findings advance our understanding of fusion-driven tumor biology and nominate novel, fusion-aware biomarkers and therapeutic targets for molecularly guided bladder cancer management. Future experimental validation and clinical correlation will be critical to translating these insights into personalized therapeutic strategies.

### Comparative Summary of Molecular, Immunological, and Regulatory Features in FGFR3– TACC3 Fusion-Positive Versus Fusion-Negative Bladder Cancer

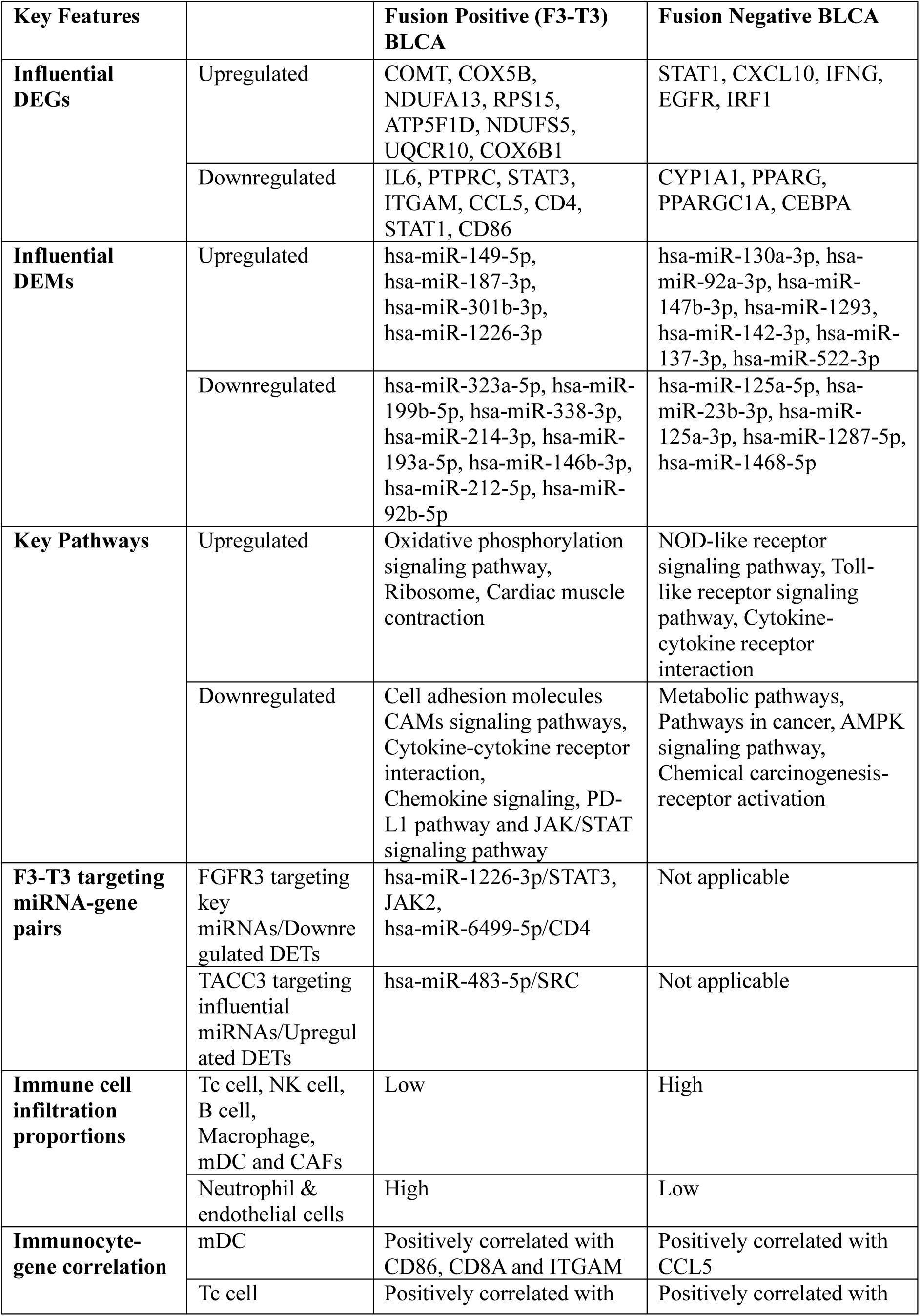
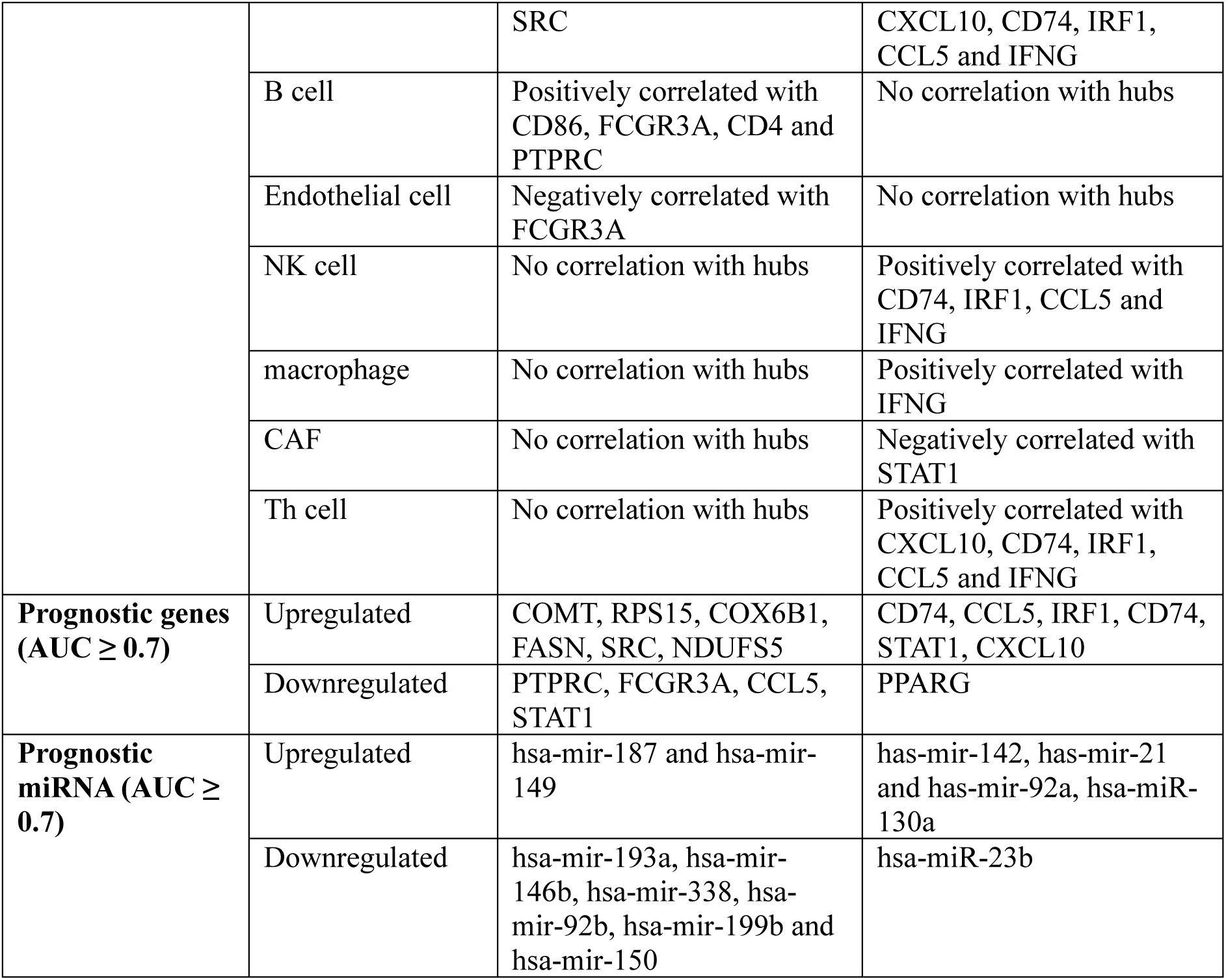

