## Supplemental file for "Integrative Transcriptomic and miRNA Analysis Reveals Immune Suppression and Metabolic Reprogramming in FGFR3–TACC3 Fusion-Positive versus Fusion-Negative Bladder Cancer"

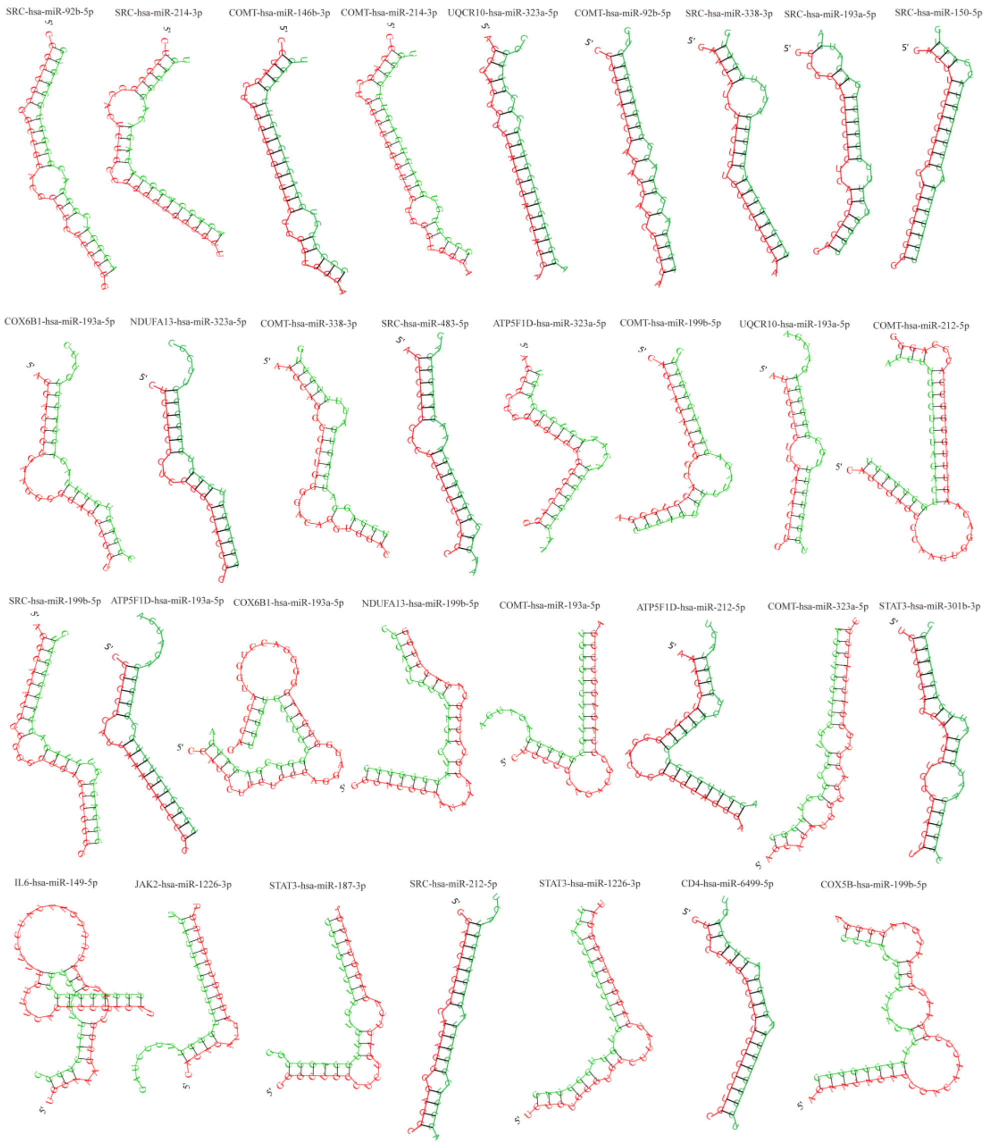


**Supplementary figure S1.** 2D structures of influential miRNA-mRNA pairs in fusion positive BLCA cases.

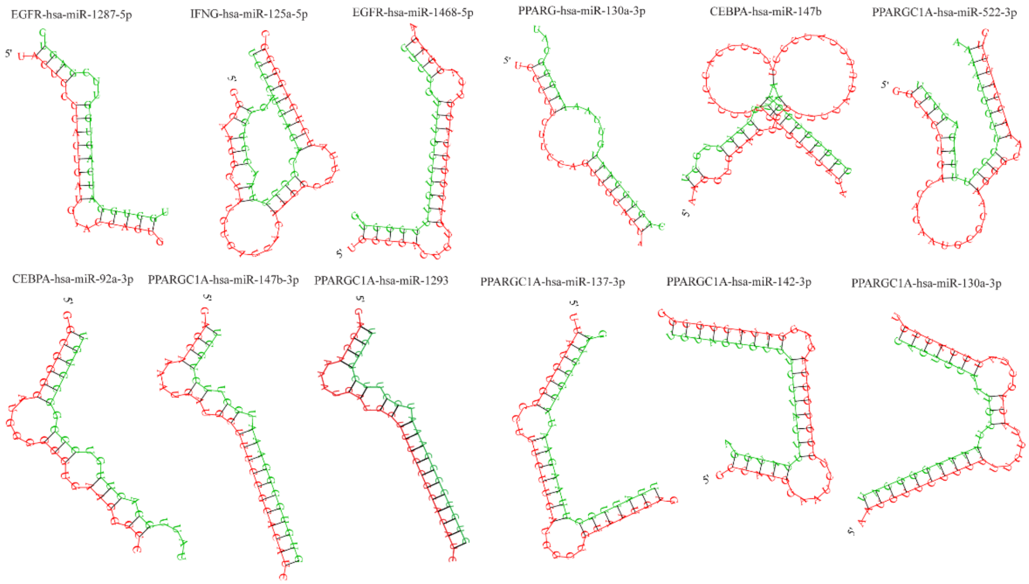


**Supplementary figure S2.** 2D structures of influential miRNA-mRNA pairs in fusion negative BLCA cases.

**Supplementary Table S1. Sample IDs used in BLCA study.**

| **Fusion Positive Case IDs** | | | | |
|---|---|---|---|---|
| TCGA-CF-A47S | TCGA-CF-A3MF | TCGA-CF-A3MG | TCGA-E7-A5KE | TCGA-CF-A47T |
| TCGA-CF-A3MH | TCGA-CF-A8HY | TCGA-XF-AAMZ | | |
| **Fusion Negative Case IDs** | | | | |
| TCGA-HQ-A5ND | TCGA-DK-A6B2 | TCGA-GD-A3OQ | TCGA-E5-A2PC | TCGA-FD-A3B8 |
| TCGA-FD-A6TC | TCGA-CF-A47W | TCGA-FD-A3SQ | TCGA-BT-A20Q | TCGA-GC-A6I1 |
| TCGA-DK-AA74 | TCGA-FD-A43U | TCGA-YF-AA3M | TCGA-DK-A3IM | TCGA-BL-A13J |
| TCGA-DK-A6AW | TCGA-FD-A3NA | TCGA-G2-A2EL | TCGA-FD-A43P | TCGA-DK-A1AA |
| TCGA-GV-A3JV | TCGA-DK-A3IL | TCGA-XF-A9T8 | TCGA-SY-A9G0 | TCGA-UY-A78N |
| TCGA-XF-A8HH | TCGA-XF-A8HC | TCGA-GD-A3OP | TCGA-XF-A9SU | TCGA-DK-A3IT |
| TCGA-DK-A2I1 | TCGA-ZF-AA4T | TCGA-XF-AAML | TCGA-UY-A78O | TCGA-ZF-AA4U |
| TCGA-XF-A9SK | TCGA-E7-A97P | TCGA-4Z-AA82 | TCGA-CF-A8HX | TCGA-UY-A8OC |
| TCGA-ZF-AA5H | TCGA-XF-AAMW | TCGA-K4-A5RI | TCGA-FD-A3SL | TCGA-K4-A5RJ |
| TCGA-E7-A7PW | TCGA-E5-A4U1 | TCGA-XF-A9ST | TCGA-4Z-AA87 | TCGA-ZF-A9R0 |
| TCGA-PQ-A6FI | TCGA-E7-A8O7 | TCGA-GV-A3QH | TCGA-DK-AA6S | TCGA-DK-A2I4 |
| TCGA-C4-A0EZ | TCGA-ZF-A9RG | TCGA-HQ-A2OE | TCGA-UY-A9PA | TCGA-FD-A6TE |
| TCGA-FD-A62S | TCGA-4Z-AA7M | TCGA-4Z-AA81 | TCGA-FD-A3B5 | TCGA-E7-A5KF |
| TCGA-FD-A5C0 | TCGA-GC-A3OO | TCGA-FD-A3B6 | TCGA-FD-A62O | TCGA-GD-A2C5 |
| TCGA-GU-A42R | TCGA-K4-A4AC | TCGA-DK-A1AC | TCGA-FD-A3SM | TCGA-G2-A2EK |
| TCGA-UY-A78M | TCGA-GU-A764 | TCGA-ZF-A9R4 | TCGA-GV-A3QF | TCGA-DK-AA6R |
| TCGA-GV-A3QI | TCGA-G2-AA3B | TCGA-K4-AAQO | TCGA-DK-A1AE | TCGA-4Z-AA80 |
| TCGA-FD-A5C1 | TCGA-BT-A0YX | TCGA-ZF-A9RL | TCGA-4Z-AA7Y | TCGA-UY-A78K |
| TCGA-DK-AA6X | TCGA-DK-AA71 | TCGA-FD-A6TB | TCGA-CU-A0YO | TCGA-XF-A9T0 |
| TCGA-CU-A3YL | TCGA-DK-A2I2 | TCGA-XF-AAMG | TCGA-FD-A3B3 | TCGA-XF-A9T3 |
| TCGA-FD-A6TH | TCGA-XF-AAN3 | TCGA-CF-A1HR | TCGA-CF-A5U8 | TCGA-FD-A3SP |
| TCGA-BT-A20T | TCGA-ZF-AA5P | TCGA-LC-A66R | TCGA-GC-A3YS | TCGA-DK-AA6Q |
| TCGA-FD-A6TG | TCGA-BT-A20V | TCGA-GU-AATP | TCGA-BT-A20W | TCGA-4Z-AA89 |
| TCGA-DK-A6B1 | TCGA-CF-A7I0 | TCGA-2F-A9KW | TCGA-DK-AA6P | TCGA-K4-A6FZ |
| TCGA-ZF-AA58 | TCGA-2F-A9KQ | TCGA-UY-A9PF | TCGA-ZF-A9RC | TCGA-FD-A6TF |
| TCGA-XF-AAMH | TCGA-ZF-A9RE | TCGA-FD-A43N | TCGA-FD-A5BV | TCGA-ZF-AA4W |
| TCGA-BL-A3JM | TCGA-E7-A7XN | TCGA-GV-A3JW | TCGA-5N-A9KI | TCGA-DK-AA6W |
| TCGA-FD-A6TI | TCGA-G2-A2EC | TCGA-DK-A1A7 | TCGA-FD-A43X | TCGA-FD-A6TK |
| TCGA-FD-A3SN | TCGA-E7-A678 | TCGA-E7-A7DV | TCGA-C4-A0F6 | TCGA-2F-A9KP |
| TCGA-ZF-AA51 | TCGA-G2-A2EF | TCGA-DK-A3IK | TCGA-XF-A9SL | TCGA-YF-AA3L |
| TCGA-GV-A40G | TCGA-GD-A3OS | TCGA-C4-A0F0 | TCGA-K4-A54R | TCGA-K4-A6MB |
| TCGA-DK-AA6U | TCGA-DK-A3WW | TCGA-DK-A3X2 | TCGA-XF-AAN1 | TCGA-CU-A0YN |
| TCGA-4Z-AA84 | TCGA-CU-A0YR | TCGA-XF-A9SP | TCGA-YC-A9TC | TCGA-KQ-A41N |
| TCGA-K4-A4AB | TCGA-XF-AAMR | TCGA-S5-A6DX | TCGA-XF-A9T6 | TCGA-K4-A3WS |
| TCGA-FD-A3N6 | TCGA-4Z-AA7W | TCGA-FD-A43S | TCGA-E7-A7DU | TCGA-BT-A0S7 |
| TCGA-BT-A42E | TCGA-CF-A47X | TCGA-GC-A3BM | TCGA-FD-A3SR | TCGA-HQ-A2OF |

| | | | | |
|---|---|---|---|---|
| TCGA-FD-A5BY | TCGA-XF-AAMQ | TCGA-4Z-AA7N | TCGA-G2-A2ES | TCGA-R3-A69X |
| TCGA-XF-AAMJ | TCGA-4Z-AA7O | TCGA-4Z-AA83 | TCGA-4Z-AA7R | TCGA-GU-A762 |
| TCGA-DK-A1A5 | TCGA-CF-A9FH | TCGA-BT-A2LA | TCGA-FJ-A3Z9 | TCGA-GC-A3RD |
| TCGA-XF-A8HB | TCGA-BT-A20U | TCGA-GV-A3QG | TCGA-BT-A3PK | TCGA-LT-A8JT |
| TCGA-FJ-A3ZE | TCGA-XF-A8HE | TCGA-CU-A5W6 | TCGA-CF-A9FF | TCGA-XF-A9SI |
| TCGA-E7-A4XJ | TCGA-E7-A8O8 | TCGA-FD-A3SS | TCGA-DK-A6B0 | TCGA-FJ-A3Z7 |
| TCGA-ZF-AA53 | TCGA-XF-A8HF | TCGA-FT-A61P | TCGA-ZF-A9RN | TCGA-FD-A5BX |
| TCGA-E7-A6MD | TCGA-H4-A2HO | TCGA-SY-A9G5 | TCGA-FD-A62P | TCGA-DK-A3X1 |
| TCGA-BT-A3PH | TCGA-DK-A1AD | TCGA-ZF-A9RM | TCGA-E7-A519 | TCGA-GD-A76B |
| TCGA-4Z-AA86 | TCGA-FD-A3B4 | TCGA-CF-A27C | TCGA-BT-A20J | TCGA-ZF-AA52 |
| TCGA-XF-AAMF | TCGA-XF-A9SY | TCGA-XF-A9SG | TCGA-GC-A3I6 | TCGA-FD-A3SJ |
| TCGA-FD-A3SO | TCGA-S5-AA26 | TCGA-XF-A9SJ | TCGA-FD-A6TD | TCGA-K4-A3WV |
| TCGA-YC-A89H | TCGA-XF-A9SX | TCGA-ZF-AA4R | TCGA-XF-AAME | TCGA-DK-A6B5 |
| TCGA-CF-A9FL | TCGA-FD-A3N5 | TCGA-DK-A6AV | TCGA-E7-A3X6 | TCGA-DK-A3WX |
| TCGA-5N-A9KM | TCGA-CF-A1HS | TCGA-CF-A5UA | TCGA-E7-A85H | TCGA-KQ-A41P |
| TCGA-YC-A8S6 | TCGA-CU-A3KJ | TCGA-FD-A5BS | TCGA-C4-A0F7 | TCGA-DK-AA6M |
| TCGA-FD-A5BU | TCGA-GC-A4ZW | TCGA-XF-AAN4 | TCGA-E5-A4TZ | TCGA-GV-A3JX |
| TCGA-XF-AAN2 | TCGA-XF-AAN8 | TCGA-FD-A62N | TCGA-BT-A3PJ | TCGA-BT-A42F |
| TCGA-G2-AA3F | TCGA-K4-A83P | TCGA-XF-A9SH | TCGA-DK-AA6L | TCGA-GC-A3WC |
| TCGA-G2-A2EO | TCGA-GV-A40E | TCGA-UY-A78P | TCGA-FD-A5BT | TCGA-BT-A42B |
| TCGA-K4-A3WU | TCGA-XF-A9T2 | TCGA-BT-A42C | TCGA-KQ-A41S | TCGA-XF-A9SZ |
| TCGA-DK-A3WY | TCGA-ZF-A9R5 | TCGA-XF-A8HG | TCGA-ZF-AA4V | TCGA-2F-A9KO |
| TCGA-CF-A47Y | TCGA-CF-A9FM | TCGA-FJ-A871 | TCGA-FT-A3EE | TCGA-DK-A2I6 |
| TCGA-DK-A1A6 | TCGA-GD-A6C6 | TCGA-2F-A9KT | TCGA-UY-A9PH | TCGA-XF-A8HD |
| TCGA-GU-A42Q | TCGA-C4-A0F1 | TCGA-ZF-AA54 | TCGA-GU-AATO | TCGA-FD-A5BR |
| TCGA-GV-A3JZ | TCGA-DK-A6B6 | TCGA-GC-A3RC | TCGA-G2-A3VY | TCGA-HQ-A5NE |
| TCGA-XF-AAMY | TCGA-ZF-A9R3 | TCGA-BT-A20X | TCGA-XF-A9SV | TCGA-G2-AA3D |
| TCGA-XF-AAN7 | TCGA-FD-A5BZ | TCGA-GU-A767 | TCGA-ZF-A9R7 | TCGA-DK-A3IQ |
| TCGA-KQ-A41O | TCGA-GC-A6I3 | TCGA-G2-AA3C | TCGA-DK-AA75 | TCGA-KQ-A41R |
| TCGA-BT-A2LD | TCGA-E7-A97Q | TCGA-BT-A20O | TCGA-H4-A2HQ | TCGA-CF-A47V |
| TCGA-GV-A3QK | TCGA-XF-AAMT | TCGA-XF-AAN5 | TCGA-BT-A2LB | TCGA-ZF-AA4X |
| TCGA-UY-A9PD | TCGA-CU-A72E | TCGA-BL-A5ZZ | TCGA-MV-A51V | TCGA-GU-A766 |
| TCGA-E7-A6ME | TCGA-CU-A3QU | TCGA-DK-A1A3 | TCGA-FD-A43Y | TCGA-XF-AAN0 |
| TCGA-DK-A3IS | TCGA-ZF-A9R2 | TCGA-G2-A2EJ | TCGA-DK-AA76 | TCGA-ZF-AA56 |
| TCGA-GC-A3RB | TCGA-G2-A3IB | TCGA-G2-A3IE | TCGA-UY-A78L | TCGA-DK-AA77 |
| TCGA-KQ-A41Q | TCGA-GU-AATQ | TCGA-ZF-AA4N | TCGA-E7-A677 | TCGA-ZF-A9R9 |
| TCGA-PQ-A6FN | TCGA-XF-A9SM | TCGA-DK-A2HX | TCGA-E7-A541 | TCGA-BL-A13I |
| TCGA-UY-A8OD | TCGA-E7-A6MF | TCGA-BT-A20R | TCGA-CF-A3MI | TCGA-BT-A20N |
| TCGA-FJ-A3ZF | TCGA-E7-A4IJ | TCGA-DK-A1AG | TCGA-DK-A1AF | TCGA-LT-A5Z6 |
| TCGA-ZF-AA5N | TCGA-DK-A3IN | TCGA-UY-A9PB | TCGA-BT-A20P | TCGA-XF-AAMX |
| TCGA-E7-A3Y1 | TCGA-FD-A6TA | TCGA-DK-A3IV | TCGA-ZF-A9RD | TCGA-K4-A5RH |

| | | | | |
|---|---|---|---|---|
| TCGA-DK-A3IU | TCGA-UY-A8OB | TCGA-XF-A9T4 | TCGA-GV-A6ZA | TCGA-XF-A8HI |
| TCGA-ZF-A9R1 | TCGA-UY-A9PE | TCGA-XF-A9T5 | TCGA-XF-A9SW | TCGA-FD-A3B7 |
| TCGA-DK-A1AB | TCGA-GU-A42P | TCGA-DK-AA6T | TCGA-GU-A763 | |
| TCGA-4Z-AA7S | TCGA-4Z-AA7Q | TCGA-ZF-A9RF | TCGA-2F-A9KR | |
| **Normal Samples' IDs** | | | | |
| TCGA.BT.A20N | TCGA.BT.A20R | TCGA.BT.A2LA | TCGA.GC.A3BM | TCGA.BT.A20Q |
| TCGA.BT.A2LB | TCGA.GD.A2C5 | TCGA.BT.A20U | TCGA.GC.A6I3 | TCGA.BT.A20W |
| TCGA.GC.A3WC | TCGA.BL.A13J | TCGA.CU.A0YN | TCGA.CU.A0YR | TCGA.K4.A54R |
| TCGA.K4.A3WV | TCGA.K4.A5RI | TCGA.GD.A3OP | TCGA.GD.A3OQ | |

**Supplementary Table S2. The identified Significant set of DEGs in BLCA datasets.**

| **560 exclusive upregulated DEGs for F3-T3 fusion-positive BLCA** | | | | | | | | |
|---|---|---|---|---|---|---|---|---|
| MCRIP2 | SEMA3F | RGS17 | TOX3 | AOC2 | SCAP | ARHGEF 19 | RHPN1 | TMEM145 |
| RAB40A | POLR2J | ANGPT2 | KCNN4 | C1orf159 | RPS15 | SH3BGR L3 | RNF207 | KIFC2 |
| TMEM134 | KRT33A | SEMA6A | CLASRP | MTSS2 | AGMAT | CNKSR1 | SLC45A3 | TRPV3 |
| CORO1B | BAIAP3 | COMT | PPP1R37 | KHDC4 | GALE | SYTL1 | PLA2G2F | HID1 |
| SSH3 | NOX1 | MZF1 | PPP1R13L | LPIN3 | ARTN | TINAGL1 | CFAP298 | GSDMA |
| NADSYN1 | LYPLA2 | FBXL19 | PTOV1 | PEMT | FAAH | BEST4 | CCDC24 | E4F1 |
| ANKRD13 D | SYT7 | ATP5F1D | MED25 | RNF128 | RSRP1 | HCN3 | SIM2 | DNASE1L 2 |
| AHSA2P | MAP4K3 | TECR | C19orf53 | MACROD1 | HOXB3 | PYCR2 | NR2F6 | NT5DC2 |
| SNCG | PRICKLE 3 | IZUMO4 | CCDC61 | GGT2 | NR2C1 | REN | TFF2 | SMIM4 |
| MST1 | IL20RA | ARVCF | EPHX3 | AGAP3 | TBX2 | PPFIA4 | TFF1 | SERINC2 |
| SWSAP1 | NRXN3 | TRMT2A | DYRK1B | REG4 | TMEM54 | DQX1 | C9orf116 | NDUFS5 |
| TNK1 | EPN3 | LZTR1 | CAPS | SLC38A2 | BHLHE41 | RPL37 | PPP1R16A | PLA2G4F |
| PHLDA3 | HEBP2 | CECR2 | BCAT2 | CYP2J2 | HOXC13 | HAPLN1 | FDXR | ATP5ME |
| GOLGA8A | PLEKHH1 | SLC25A1 | ISYNA1 | DHX34 | B9D2 | FAM193B | ZNF385A | B3GALN T1 |
| CENPS | SPAG4 | SH3BP1 | ETV2 | BLOC1S1 | CDH26 | WASH2P | WDR90 | HEXD |
| CABP4 | TNK2 | PATZ1 | ADAP1 | SMPD2 | NDUFA1 | AC004453 .1 | RPL29 | FASN |
| KDF1 | DGAT2 | GCAT | STX1A | COX5B | NT5C | SH3GLB2 | CAMK2N1 | DCXR |
| DOK7 | RPL18 | PICK1 | CYP3A5 | ZFHX2 | MBOAT7 | LRMDA | ATXN7L2 | ZNF768 |
| ORAI3 | ATP2C2 | ASCC2 | DVL1 | FAM13A | PPDPF | ZP1 | FCRLB | PUSL1 |
| CCDC57 | LPAR2 | APOL4 | RPL28 | CCNG2 | TMEM74 B | TRPT1 | DENND2D | KCNAB3 |
| PHLDB3 | ERBB3 | IFT27 | RUNDC3A | CASP6 | COX6B1 | TM7SF2 | ACP6 | ANKK1 |
| VSIG10 | POLR1H | PABPN1 | CDK5RAP3 | AMIGO2 | PRRG2 | TBX6 | CLHC1 | FABP6 |
| ULK1 | FGFR3 | GSS | RECQL5 | DUSP6 | TIMM17B | ALDOA | MST1R | GPRC5C |
| RIMKLA | IP6K2 | ABHD12 | PIGL | CSAD | IDUA | LYPD6B | HEY1 | PYM1 |
| RABEP2 | RNF126 | ZMYND8 | KAT2A | LMBR1L | BAIAP2L 2 | ME3 | TMEM184 A | ZNF296 |
| TMEM94 | RPS6KA6 | CST3 | AADAT | NEIL1 | MPST | ZNF547 | METTL27 | KCNS3 |
| MAMDC4 | SELENOO | CST4 | ST3GAL4 | ULK3 | AP1M2 | C16orf74 | AKR1E2 | NDUFA3 |
| STAP2 | LLGL2 | AMMECR 1 | MVK | RHOT2 | THEM6 | CERS3 | PLEKHF1 | HS6ST2 |
| P4HTM | CAMSAP 3 | ELMO3 | TRPV4 | G6PC3 | PGLS | ABCA10 | CYB5R2 | PKIA |
| DPM3 | RAP1GAP | MMP15 | CHD4 | NARF | SAMD10 | TRIM11 | BLCAP | ZNF692 |
| WDR97 | STXBP2 | ESRP2 | LDHB | GRB7 | PAK4 | CXADR | ACSF2 | KSR2 |
| CITED4 | HSD17B2 | CIAO3 | AC009533.1 | MIEN1 | RBBP8NL | PXYLP1 | ENGASE | ZNF524 |
| TMEM86B | FER1L4 | RHBDL1 | VEGFA | PLPP2 | METTL26 | LRFN2 | RSKR | POLR1C |

| | | | | | | | | |
|---|---|---|---|---|---|---|---|---|
| ZNF443 | SLC15A1 | CAPN15 | FAM162A | TMEM91 | ATP5IF1 | MAPK13 | CACNB3 | LPAR3 |
| SCRIB | YPEL3 | ELOB | NPRL2 | FAM71E1 | SLC6A8 | KRTCAP3 | TTYH1 | BSG |
| ZNF121 | AC246787.1 | SULT1A1 | SMIM22 | AC108136.1 | LMNTD2 | OR7E62P | NQO1 | C2orf15 |
| TEX45 | MKRN2OS | PDXDC2P-NPIPB14P | LINC02560 | ZNF432 | WASH3P | ANKRD65 | SYNE4 | AC092171.5 |
| EPS8L3 | FAM229A | HCAR1 | AC011503.2 | SBNO1-AS1 | MRPL40 | MCF2L-AS1 | PLAG1 | LINC02604 |
| MUC2 | AL354836.1 | ARRDC1 | CYP4F23P | AC005840.2 | INKA1 | RPL6P27 | TP53TG1 | NOL12 |
| APRT | DANCR | SRC | AC010326.3 | KLRA1P | PCGF3 | DICER1-AS1 | WASH6P | AC015912.3 |
| SNORD104 | SLC8A1-AS1 | RAD54B | AL390719.2 | C17orf100 | NDUFA4L2 | TMEM147-AS1 | MFSD5 | TBC1D3L |
| IQANK1 | AC107983.1 | GATA3-AS1 | AL024508.1 | MIR200CHG | BCAP31 | RPS28P4 | SNORC | PCCA-DT |
| C1orf53 | LINC02672 | RAB40C | AL645608.7 | AC126178.1 | NDUFA13 | CKMT1B | HOXB4 | AC021491.4 |
| LIME1 | LINC01341 | S100A6 | GTF2IP20 | FAHD2P1 | EDARADD | LINC01637 | NAA38 | AC244153.1 |
| RTL8A | GAPDHP1 | AC069360.1 | UCA1 | CNPY2 | MST1P2 | GOLGA2P5 | SLC25A10 | HERC2P2 |
| C6orf136 | PCAT6 | AC109439.1 | TUBA5P | PRANCR | CYP4F8 | BX284668.5 | RIPPLY3 | AC084781.1 |
| KRTAP5-10 | MIR34AHG | AC003072.1 | RTL10 | AC018904.1 | MST1L | KMT2E-AS1 | PRR36 | SRCIN1 |
| C12orf73 | AC026403.1 | SNHG18 | NPEPL1 | AL390728.6 | OR10H1 | RPL37P6 | PI4KAP2 | AL161669.3 |
| PSENEN | PET100 | ZFPM2-AS1 | AATBC | HOXB7 | FAM183A | DECR2 | AIFM3 | DHRS11 |
| TMEM256 | MAP4K3-DT | LINC00967 | ARHGAP27P1-BPTFP1-KPNA2P3 | AC068700.1 | SIRT7 | AC115223.1 | NUDT14 | AC022400.6 |
| OR7E91P | VN1R54P | LINC01151 | TSTD1 | AL023284.4 | ANKRD19P | AC116533.1 | TBX1 | WASH9P |
| TMEM191C | TMA7 | MEX3A | ATAD3C | FTLP14 | C3orf62 | CEBPA | UQCR10 | SAMD12-AS1 |
| DIO1 | DUX4L50 | SNHG9 | AC110619.1 | AC003102.1 | SBK1 | AC016065.1 | LRTOMT | GTF2IP12 |
| PRR22 | PHGR1 | NBPF11 | CCNL2 | CASC22 | FBLL1 | H2AJ | SNRNP35 | AC140479.5 |
| LINC01089 | MROH3P | RBP3 | NSUN5P1 | LINC01311 | RPL14 | SNHG10 | COLEC10 | AC013268.4 |
| RNF208 | LINC01518 | AC132812.1 | CPTP | SPINT1-AS1 | ALKBH2 | AL359220.1 | CIB1 | AC073648.6 |
| CRYGS | AC011294.1 | ZNF488 | NUDT19 | AC010168.2 | SP6 | TMED2-DT | ANO9 | BX255925.3 |
| RPL7AP50 | RPS15P4 | AC103810.3 | ZNF726 | LINC01355 | MYBPC1 | MTND4P12 | TNFAIP2 | AC092306.1 |
| RPL17P50 | LNCTAM34A | MAP3K14-AS1 | PTOV1-AS2 | SMG1P7 | KIAA0895L | ABHD14A | MAGEA11 | AC084024.4 |
| RPL13AP7 | MAPKAPK5-AS1 | TBX2-AS1 | RAB11B-AS1 | SSTR5-AS1 | MYT1 | ZNF436-AS1 | ZNF74 | AC083855.2 |
| PPM1N | MAGIX | CEBPA-DT | ZNF285 | AC087741.1 | S100A4 | CASC9 | CDK10 | Z83745.1 |
| AL590062.1 | FAM174B | HOXC13-AS | AC104532.2 | AC110285.2 | S100A5 | AC024560.2 | TMEM105 | AC027601.5 |

| | | | | | | | | |
|---|---|---|---|---|---|---|---|---|
| ALOX12P2 | NPIPB15 | | | | | | | |
| **1086 exclusive downregulated DEGs for F3-T3 fusion-positive BLCA** | | | | | | | | |
| LAP3 | USP13 | IRAK3 | PLEKHA4 | DSE | IGSF21 | SEPTIN6 | IL15RA | STAT4 |
| TMEM176A | CDK14 | LYZ | LILRB5 | GCNT2 | GBP1 | HS3ST3B1 | DOCK2 | ITGAV |
| KLHL13 | TBXAS1 | CD209 | PIK3CG | ADTRP | CNN3 | TNFSF9 | KLRD1 | PARP9 |
| CYP26B1 | NGFR | CERS4 | TFEC | SERINC1 | F3 | C3 | PIWIL4 | ARHGAP24 |
| CD38 | HIPK2 | WDR7 | CAV2 | NCOA7 | TNFSF4 | NRSN2 | SLC43A3 | HERC5 |
| COPZ2 | OAT | MAP3K20 | CAV1 | RIPOR2 | NRP2 | IFI6 | C5AR2 | RASGEF1B |
| ITGAL | TRAM2 | CDV3 | CPVL | SOD2 | C1orf54 | SGPP1 | DZIP1 | GPAT3 |
| MAP3K14 | ANKRD44 | TGFB2 | COBL | PHACTR1 | ELOVL4 | EVI2A | ETS1 | LEF1 |
| CX3CL1 | ME1 | FMO2 | PTPRZ1 | MDGA1 | CNR1 | AIF1L | RFK | PARVG |
| MATK | CD84 | CREM | GIMAP2 | CD83 | SGIP1 | FAM78A | ANXA1 | CPNE8 |
| CD79B | SPI1 | HIVEP1 | TMEM176B | BACH2 | PLAGL1 | IPPK | CTSL | PRICKLE1 |
| CYTH3 | NAV3 | FKBP5 | BLVRA | BAG2 | TNFAIP3 | HIVEP3 | ADAM19 | CD27 |
| NFIX | ROCK1 | JAK2 | TGFBR1 | VNN2 | RAB32 | MACF1 | HAVCR2 | GLIPR1 |
| IL32 | HDAC4 | IL12RB1 | C5 | PHACTR2 | SGK1 | TNFRSF19 | SDS | SLC46A3 |
| TRAF3IP3 | ACSL4 | ABL1 | AKNA | QKI | MFSD1 | STEAP4 | CD36 | SSTR1 |
| IFFO1 | KCNAB2 | NRP1 | TNFSF8 | DUSP22 | KLF12 | APOL3 | PRR5L | RTN1 |
| STAB1 | RAB27A | PRTFDC1 | DNM1 | CLIC5 | CCND2 | LIF | TESPA1 | GPR65 |
| CD4 | NUCB2 | OSM | RIC1 | LY86 | NEK6 | DOCK4 | STX11 | WARS1 |
| BTK | SMG6 | GGT5 | NCS1 | LOX | GPR68 | DNAJB9 | CCDC102A | PSTPIP1 |
| FYN | FSTL3 | CYTH4 | RGP1 | SPARC | OGFRL1 | CHN1 | SP110 | BCL2A1 |
| ANOS1 | ACTN1 | MFNG | DDX58 | PCDHB5 | IFIT3 | ARHGAP22 | DOCK10 | MFGE8 |
| PLAUR | ACAP1 | LGALS2 | GLIS3 | PCDHB7 | IFIT2 | WDFY4 | USP37 | ITGAX |
| ZBTB32 | ST6GAL1 | HMOX1 | RAPGEF1 | ITK | MOB3B | TMOD2 | SERPINE2 | IGSF6 |
| TYROBP | TEAD2 | GRAP2 | SETX | ARSB | CD274 | MAP7D3 | ALDH1L2 | NLRC5 |
| ELOVL5 | ZNF532 | CSF2RB | RAB11FIP2 | ST8SIA4 | SNX19 | APOE | PLXNC1 | IRF8 |
| CLDN11 | SEMA3A | IL2RB | PALD1 | NR3C1 | KCNJ5 | GADD45G | SLC41A2 | IMPA2 |
| MAMLD1 | ACAT1 | CERK | VSIR | HRH2 | TGFBI | LRCH2 | LRCH1 | OSBPL1A |
| CD6 | ADD2 | GZMH | SPOCK2 | DBN1 | PPP3CC | PXDN | SCRN1 | NPC1 |
| WAS | ARHGAP15 | GZMB | LIPA | CD86 | TNFRSF8 | LSP1 | TNS3 | PIK3R5 |
| WWTR1 | MCAM | SUSD6 | CCL2 | AMOTL2 | TMEM13 | COL5A1 | IL6 | SECTM1 |

| | | | | | | | | |
|---|---|---|---|---|---|---|---|---|
| | | | | | 1L | | | |
| CD74 | APBB1IP | SEC23A | CCL8 | RBP1 | ADCY7 | PRRC2B | AOAH | FKBP10 |
| RUNX3 | JADE1 | SLA2 | ABI3 | HYAL1 | CCR2 | THEMIS 2 | MYO1G | VAV1 |
| SERPINB1 | CST7 | HCK | COL1A1 | GNB4 | TMEM15 6 | MPP1 | BIN1 | IFITM3 |
| OSBPL5 | MAP2 | SAMHD1 | DUSP3 | FRMD4B | SASH3 | PRRG1 | MYC | COL6A1 |
| GRAMD1 B | LAMP3 | PPP1R16 B | FAM20A | PLSCR4 | LY9 | LILRB2 | CCN3 | APP |
| BIRC3 | FAP | LAMA1 | VTN | KCNIP3 | WIPF3 | RFTN1 | IL33 | EMP3 |
| PLEKHO1 | EDN1 | RNF125 | KLHL5 | ZAP70 | FKBP9 | GFPT2 | TLN1 | MYO1F |
| EHD2 | P2RY10 | TLR8 | JHY | CYTIP | RAMP3 | CHSY1 | CCL21 | SIGLEC1 0 |
| RRAGD | OSBPL6 | PLS3 | HSPA8 | FNDC4 | TWIST1 | TRIM22 | SIT1 | RCN3 |
| TYMP | SP140 | RENBP | MS4A6A | ITGA4 | GLIPR2 | RAMP1 | DNAJB5 | C1orf21 6 |
| VIM | EPB41L2 | RUBCNL | MS4A4A | IFIH1 | ACO1 | CLEC10A | CD72 | C1orf16 2 |
| FAS | CXCL2 | TNFSF13 B | TRIM3 | LOXL3 | SRGN | ZBED3 | GMPR | CD53 |
| SLAMF7 | OSTM1 | KATNAL 1 | PANX1 | EFEMP1 | CHST3 | ALOX5A P | IRF4 | GPR161 |
| BTN3A1 | PTPRC | CORO1A | IL10RA | FN1 | BICC1 | TMCC2 | TUBB2A | FCGR2A |
| TNFRSF1 B | STK17B | CPPED1 | GALNT18 | STAT1 | RASSF8 | DCLK1 | TUBB2B | ECM1 |
| RABEP1 | PCDHB4 | IL21R | CBL | GNLY | ADGRE5 | EPSTI1 | MYO7A | ADAMTS L4 |
| ARHGAP3 1 | FYB1 | ZNF106 | SLC15A3 | IL18R1 | ARHGAP 9 | POSTN | SLCO2B1 | DYRK3 |
| MFAP3 | ERC1 | ATP8B4 | CD5 | IL18RAP | NCKAP1 L | PLAAT4 | RAB30 | DEGS1 |
| TLL1 | CYLD | SLC30A4 | HPS5 | WIPF1 | SARDH | GIMAP4 | PI15 | WNT9A |
| VCAN | SMAP2 | TRPS1 | POU2AF1 | PLEK | TNFAIP6 | ARNTL | SULF1 | PTPN7 |
| MSR1 | COL16A1 | SLC39A1 4 | SELPLG | PRRX1 | BATF3 | MICAL2 | SDCBP | FBLN7 |
| TNC | CD82 | MAP4K1 | CORO1C | RNF19B | PREX1 | RRAS2 | DDX60 | OSBPL10 |
| LCP2 | FCN1 | CD37 | BIN2 | DNAJC6 | TOX2 | RNF122 | CASP1 | STAC |
| CNTLN | PILRA | LILRB1 | ELK3 | NCF2 | ZNF831 | ADAMD EC1 | PLCB2 | CAND2 |
| HDAC9 | LAT2 | IL27RA | PARP11 | CD2 | SNAI1 | CD180 | STRA6 | PTPRG |
| KITLG | ADAMTS 2 | RASAL3 | SH2B3 | RIMS3 | ZBP1 | PTPN22 | IFI44L | IL17RD |
| LAMA3 | CASS4 | NUMBL | CREBL2 | AKT3 | BTN2A2 | NOTCH2 | ARHGAP 29 | LRIG1 |
| SYNE2 | SIGLEC1 | EBI3 | ALDH2 | ST6GALN | COL21A | SPIRE1 | IFI44 | NXPE3 |

| | | | | | | | | |
|---|---|---|---|---|---|---|---|---|
| | | | | AC5 | 1 | | | |
| FAM168A | P2RX7 | DENND3 | MGP | SLAMF1 | ATXN1 | RSAD2 | ADCY3 | PLA1A |
| TRAF1 | LAG3 | NKG7 | STX2 | CD48 | AHNAK | CMPK2 | ENTPD1 | ADPRH |
| PRDM1 | RGS1 | CD33 | COL12A1 | PADI2 | MT2A | IL2RA | RBP4 | ATP10D |
| TMCC3 | ICAM1 | PLA2G4C | BTN3A3 | RGS4 | IRF1 | ECHDC3 | ADAMTS14 | GREM1 |
| SNCA | LRP12 | ANO4 | LY96 | PRXL2C | CD3G | PYHIN1 | FNDC1 | EVA1C |
| SETD7 | CDKN2B | FLI1 | NCAM2 | DZIP1L | CCR5 | IFI16 | PSIP1 | MEI1 |
| OSMR | FBXO10 | WWC2 | L3MBTL4 | MRAS | FCRL3 | EIF5A2 | SYK | PBX3 |
| PLK2 | STOM | SAV1 | PLCL2 | TENT5B | HK3 | CTLA4 | NFIL3 | PRRX2 |
| GZMA | ITPRIP | SACS | KLF10 | CD1D | ITGA5 | ICOS | ALDH1A1 | SNX20 |
| IQGAP2 | ADAM12 | BEND6 | PIP4P2 | NCF1 | SCIMP | PPM1K | CYBB | ACAA2 |
| PAM | SERPING1 | TCF7L1 | HSPA13 | SLAMF8 | ADCY9 | CXCL3 | NCF1C | AXL |
| ARHGAP26 | NPAT | NMT2 | SAMSN1 | ADAMTS4 | PDPN | ABLIM2 | OTUD1 | LAIR1 |
| FGD2 | P4HA3 | CAMK4 | SLC7A7 | FCER1G | SDC3 | SGMS2 | PGM2L1 | CD300C |
| PNRC1 | JPH2 | MBNL1 | PIK3AP1 | STC1 | MXRA8 | SLC9B2 | FOLR2 | CD300A |
| SCML4 | KIAA1755 | IFIT5 | VSIG4 | C1QC | GBP4 | ANXA5 | BEND7 | TMC8 |
| SLC22A3 | FERMT3 | DAB2 | ELMO1 | C1R | GFI1 | IL15 | CASP7 | ABCA3 |
| CREB5 | ITGB1 | CD8A | SLA | ADGRG5 | NLRP3 | STARD4 | AMOTL1 | NLRC3 |
| TMEM140 | TMCO3 | JAZF1 | CLIC2 | SPON2 | SLAMF6 | ANKRD33B | HTRA1 | RAB3IL1 |
| SH3KBP1 | LATS2 | MCOLN2 | CD109 | CARMIL2 | C1orf115 | GPX8 | GPR176 | BEST1 |
| MSN | FAM124A | ANKRD29 | UBE2L6 | KALRN | DISC1 | TLR3 | CLMP | TRANK1 |
| SLC16A2 | VEGFC | TBCEL | FBXO32 | UBASH3A | SLC66A3 | CGAS | SMPD1 | GNG4 |
| CHST7 | CCDC102B | ROBO3 | SUSD3 | GAB3 | SGCB | TMEM200A | CLEC4E | PXK |
| GPR174 | CD226 | RETREG1 | IL34 | ITGB2 | CTSS | KIAA0895 | RIMKLB | TAP1 |
| IL2RG | IL18 | ANGPT1 | ERG | GPSM1 | TNFAIP8L2 | BMPER | RRAD | RHOH |
| DOCK11 | PIP4K2A | UCHL1 | MX1 | SIK3 | ARHGAP25 | CDCA7L | B2M | RAB31 |
| MFHAS1 | CACNA2D4 | ENAH | PALM2AKAP2 | JAML | FSTL1 | SYTL3 | NNMT | COL3A1 |
| DOK2 | FER | GBP5 | FMNL2 | PCSK7 | MNDA | TAGAP | ANPEP | STAT3 |
| IL7R | CRTAP | PRNP | C1QB | CSRP2 | KCTD12 | TIGIT | TNFAIP8L3 | SOCS1 |
| COL4A3 | DENND5B | SYNPO | C1QA | TBC1D10C | ZHX2 | CHST15 | ZNF438 | TCN2 |
| ROR2 | CD14 | KLF11 | PEAK1 | LRRC25 | CD300LB | P2RY8 | GPR1 | MAPK11 |

| | | | | | | | | |
|---|---|---|---|---|---|---|---|---|
| AFAP1L2 | SERPINB 9 | LRRC15 | HEG1 | TTLL11 | MSC | HHIPL1 | RFLNB | GPRIN3 |
| MN1 | CDH2 | CD8B | CD7 | TUBB6 | GIMAP7 | UBE2E2 | CCR4 | AHNAK2 |
| RNASE6 | HTRA3 | CCL11 | GOLIM4 | NUPR1 | HLA-DQB1 | AP1S2 | KIRREL1 | OLFML2 A |
| CD52 | GSTA4 | MALT1 | C1QTNF1 | SYNE3 | CIITA | C1S | ARSI | IFIT1 |
| CLIC4 | OSCAR | ISG20 | NRROS | PNMA1 | LACC1 | CSF1R | SH2D1A | KCNQ5 |
| GPR183 | FPR2 | CXCR6 | SLC25A30 | SMCR8 | SEPTIN1 | NDN | DENND5 A | IKZF1 |
| CCDC8 | FPR1 | CLEC7A | TLR10 | SAMD9L | MCFD2 | NTM | CSF1 | EVI2B |
| ANTXR1 | GIMAP8 | IL16 | TLR1 | CAVIN1 | GAS1 | LCK | NELL2 | IFITM1 |
| MT1E | CHST11 | RASGRP 1 | TLR6 | CD163 | SSTR2 | COL18A 1 | ATL3 | RTN4RL1 |
| CSGALNA CT2 | LRRC8C | KLHL6 | ZHX3 | FAM20C | PRF1 | ADGRG3 | TRARG1 | RASA3 |
| PCDH7 | PIK3CD | THEMIS | CMKLR1 | CALHM5 | TMEM64 | GJC1 | PRR16 | AIDA |
| ITGAM | GPR34 | SLFN11 | SLCO2A1 | ZC3H12D | ARSJ | GAS6 | FMNL1 | BCL9L |
| TM4SF1 | RASGRP 4 | GXYLT2 | SH3PXD2 B | POGLUT3 | CCL13 | ARHGEF 37 | SORCS2 | BTLA |
| KLF13 | COL8A2 | HPSE | GPR171 | CD28 | ZNF467 | GPR132 | TMEM10 6A | BACE1 |
| SIMC1 | C3AR1 | PARP15 | ZDHHC14 | MAF | RELL1 | TENT5C | IFITM2 | CD300E |
| GNG2 | PEAR1 | PTPRT | C5AR1 | DPP4 | PAPSS2 | MAP3K3 | HLA-DOA | HLA-E |
| BTN3A2 | CLEC17A | HLA-DRB1 | STMN3 | PDCD1LG 2 | ECI2 | L1CAM | HLA-DMA | HLA-F |
| ARHGAP3 0 | ARL4C | GREB1 | COL13A1 | PARVA | APCDD1 L | MAGEE1 | PSMB8-AS1 | SNX2 |
| CXCR3 | MAPK12 | FUT4 | SPN | SGTB | RCSD1 | TGM2 | HLA-DRA | MT1A |
| LILRB4 | PLSCR1 | HLA-DQA1 | IRF1-AS1 | SIRPA | ALPK2 | CR1 | COL15A1 | MT1M |
| TRABD2A | PDCD1 | ARID5A | GZMM | CHSY3 | CD247 | FCGR3A | LAYN | CFI |
| LILRA5 | DPYD | PDLIM7 | ENPP1 | CSF2RA | SELENO M | INPP5B | CARD16 | SAMD9 |
| GCNT1 | PARVB | FLNA | CPLANE1 | TXNRD1 | TOX | MAFB | AIF1 | DENND1 C |
| FPR3 | HACD4 | GAL3ST4 | MFAP5 | HLA-DRB5 | CD3E | TMEM2 73 | LST1 | ATP10A |
| TRPV2 | NUGGC | FITM2 | MPEG1 | KLHL9 | SFMBT2 | ZDBF2 | LILRB3 | HCP5 |
| HLA-H | GIMAP1 | ZNF469 | ZEB1-AS1 | LINC0109 4 | GJA5 | IGLV7-46 | PLXNA4 | LINC007 02 |
| TMEM20 0C | TSPAN4 | MIAT | LINC0167 9 | CCDC71L | AC00468 7.1 | TRBC1 | HLA-DPB1 | LINC004 72 |
| HLA-A | C19orf38 | HLA-DRB6 | PSMB9 | GVINP1 | S1PR2 | IGHA1 | LINC0113 3 | HLA-B |
| DIO2 | NEURL1 | HLA- | PLEKHO2 | AC09055 | MIR222 | IGHG1 | C4B | MSC-AS1 |

| | | | | | | | | |
|---|---|---|---|---|---|---|---|---|
| | B | DPA1 | | 9.1 | HG | | | |
| IGKC | MORF4L 1P1 | LINC-PINT | KCTD7 | SMIM3 | CCL5 | LINC026 93 | PELATON | NFAM1 |
| IGLV8-61 | TENM3 | HLA-DQB2 | DUBR | CHURC1 | LIX1L | PCED1B-AS1 | LINC0066 7 | AC10453 0.1 |
| C4A | MT1L | TRAC | LILRA6 | AL365361 .1 | AL66284 4.4 | RAB7B | CCL4 | GAS6-DT |
| RASSF8-AS1 | CCPG1 | PRR33 | SCARF2 | MRC1 | | | | |
| **1494 exclusive upregulated DEGs for F3-T3 fusion-negative BLCA** | | | | | | | | |
| C1orf112 | PSMA3 | PCGF1 | COLGALT 1 | TRIM65 | XDH | PHB | PARP10 | HLA-DRB5 |
| LAS1L | AHSA1 | AUP1 | KLHDC7B | SECTM1 | FANCC | SNRPD1 | ERFE | GPN1 |
| MYH16 | ERH | EVA1A | HELZ2 | P3H4 | ABHD3 | SLC27A4 | C5orf46 | WDHD1 |
| LAP3 | MTHFD1 | MRPL19 | ADRM1 | TPGS1 | ESYT3 | DOLPP1 | RMI1 | PPIAP22 |
| CD38 | VRK1 | STAT1 | EXOSC2 | IFITM3 | H2BC5 | NOD2 | ALOXE3 | PPP1R14C |
| MCUB | PSMB5 | CCT4 | UCK1 | TRPM2 | H4C8 | RRM1 | CALR | MSRB1 |
| ITGA3 | PSMA6 | TXNDC9 | SHFL | PSMB6 | MITD1 | SPATA33 | LDLRAD3 | C1orf68 |
| RALA | PSME2 | GNLY | DNMT1 | SIGLEC10 | FAM86C1 P | RHEBL1 | HLA-DQB1 | LINC0128 5 |
| DBF4 | RABGGT A | IL1R2 | DKC1 | SLC2A5 | TAGLN2 | TUBA1C | CIITA | FCGR3A |
| CCDC124 | MMP9 | HPCAL1 | LILRB2 | MYOM3 | SLAMF8 | LAIR2 | APOBEC3 B | SPRR2E |
| PAX6 | SLA2 | NOL10 | EMC8 | IL22RA1 | DUSP23 | YIF1B | MRGPRX3 | PLPP4 |
| E2F2 | RAB5IF | SLC1A4 | IDO1 | FCN3 | CDA | KLK5 | AKAP5 | H2BC18 |
| PSMB1 | TPD52L2 | DHCR24 | VPS25 | PSMA5 | B4GALT3 | KLK6 | TMEM150 B | MAFB |
| CYB561 | PSMA7 | CACYBP | DIAPH1 | UCK2 | FCER1G | OTUB1 | FGD6 | HLA-DMA |
| IL32 | SLC17A9 | MRPL37 | KREMEN2 | XCL1 | TOMM40 L | CD300A | CCDC43 | PSMB8-AS1 |
| ETV7 | CDC25B | FBXO6 | BARX1 | FCGR2A | FBXW5 | SDHAF2 | FZD2 | PSMB8 |
| DPF1 | HM13 | MAD2L2 | KRT34 | HDGF | GART | POLR2G | BHLHA15 | TAP2 |
| ANLN | SAMHD1 | NCF2 | PODNL1 | LHX9 | C1QC | SLC3A2 | H2AC6 | HLA-DRA |
| TYROBP | NOP56 | UCHL5 | FBXW9 | ANP32E | ATP5MC1 | FADD | H2BC4 | NELFE |
| SLC38A5 | AHCY | TFAP2E | MATN3 | MRPL9 | SNF8 | BATF2 | SLC47A2 | LSM2 |
| SLC11A1 | FAM83D | MFAP2 | TRIM21 | INTS7 | IGF2BP1 | DDIT4 | PRF1 | VARS1 |
| CD74 | LIPG | ACTL8 | RAN | SMYD2 | CBR1 | GNG4 | PSMG4 | AIF1 |
| SERPINB1 | CSTF2 | GBP1 | COQ3 | ADAM15 | CBR3 | MPLKIP | HEPHL1 | MICB |
| GRAMD1B | TLR8 | GPR89A | SEC61G | S100A8 | PSMD4 | TAP1 | DDX60L | HLA-C |
| DERA | PLP2 | EBNA1BP 2 | FIGNL1 | TPM3 | PSMB4 | MLKL | TMEM45A | CCHCR1 |
| TYMP | PGK1 | ATP6V0B | TRIM47 | S100A7 | TCHH | RFWD3 | MRPS23 | PSORS1C 1 |
| CD44 | SMS | B4GALT2 | XAF1 | EFNA3 | MED8 | STIP1 | MACIR | C6orf15 |
| SH2D2A | PHEX | FASLG | ERAL1 | C1orf131 | SPRR2G | RAB31 | GINS3 | LILRB3 |
| ARNTL2 | UBL4A | STMN1 | PCNA | ACP1 | PGLYRP3 | TMEM20 8 | CHST15 | TRIM15 |

| | | | | | | | | |
|---|---|---|---|---|---|---|---|---|
| BAK1 | PBDC1 | POLR1G | ACY3 | DEGS1 | LYPD5 | WFDC12 | IDH2 | HLA-G |
| CENPQ | GLA | MREG | TESMIN | CNIH4 | WDR4 | SIGLEC7 | OGFRP1 | HLA-F |
| EIPR1 | TNFSF13B | MYB | POMP | GUK1 | PDXK | KLK7 | DEXI | SPDYC |
| ABCF2 | ACP5 | VAMP8 | EPSTI1 | MBOAT2 | ITGB2 | ACTBL2 | NOP10 | MROH6 |
| UBA6 | SLC25A15 | DCLRE1B | RFC3 | PTPN7 | FAM207A | PCSK9 | MRPL41 | IGFL2 |
| TMSB10 | ACOD1 | SPP1 | EPHB2 | PDIA6 | VAV2 | APEX2 | EXT1 | MZT1 |
| FAM136A | SLC38A7 | CPSF3 | SLF1 | ATP6V1C2 | TOR2A | RGS14 | UBE2E2 | FAM83A-AS1 |
| RPL26L1 | COTL1 | ABITRAM | WDR74 | CHAC2 | MED27 | RAB24 | NLRP10 | PCDHGA1 |
| HOXC8 | HAGHL | DCAF4 | PLAAT4 | SNRPG | RUSC1 | PRELID1 | KPNA2 | C4orf46 |
| MSR1 | SLC7A5 | IFI27L2 | C12orf29 | MRPS5 | GPATCH4 | CXCL10 | EPHB3 | PSMB10 |
| BEST2 | MEFV | FLVCR2 | RRAS2 | TMEM177 | HK3 | CXCL11 | SKA2 | SAMD9 |
| SPDL1 | IL21R | OGFRL1 | LOXL2 | HSPD1 | LY6K | GPRIN1 | ANXA2 | KRT6A |
| RTN4R | AQP9 | ELOVL3 | CDK7 | ABCA12 | CPSF4 | SPRR1B | HCAR2 | KRT81 |
| PSMA4 | HOMER2 | IFIT3 | FKBP11 | MARCHF4 | LY6E | BNC1 | VMO1 | SERPINB4 |
| CAPG | CA2 | IFIT2 | TMEM106C | SRPRB | LRWD1 | CHRNA5 | TCHHL1 | DUXAP8 |
| BARX2 | RIPK2 | MOB3B | YWHAQ | TMEM44 | MFSD12 | AVEN | PYCR1 | HCP5 |
| ZIC2 | POP1 | CD274 | RSAD2 | TM4SF19 | NAGS | KRT78 | NKX2-5 | HLA-H |
| HSPA5 | CCN4 | MTHFD1L | CMPK2 | LYAR | CXCL16 | KRT75 | GBP6 | HLA-A |
| ROS1 | GSDMD | PCDHB8 | LDHA | SRD5A1 | SENP3 | CD14 | PSMG1 | HACD2 |
| MRPS10 | PYCR3 | TP53AIP1 | SAA2 | MYO10 | CCDC78 | KRT6C | RP1L1 | IGHG1 |
| TNFRSF9 | MCM4 | PDZD11 | TIMM17A | OSMR | CLPB | RIOX1 | ALYREF | IGHG3 |
| CLPTM1L | KLC3 | NUDCD1 | RBM17 | PLK2 | TTC9C | SERPINB9 | UPP1 | TCTEX1D2 |
| NFE2L3 | AMH | MASTL | IL2RA | GZMA | RPS6KA4 | STAT2 | FOXL2 | TRIM59 |
| MCUR1 | IL4I1 | CCDC77 | IL15RA | TBC1D7 | PDZK1IP1 | COMMD5 | LIN9 | ARHGEF35 |
| TNIP3 | LILRB1 | TGFBI | TMEM241 | GFOD1 | COA7 | GTSF1 | ARSI | DNAJC9 |
| PRSS8 | IL27RA | TMPO | KLRD1 | PLA2G7 | DRAXIN | TRIAP1 | ADAP2 | VDAC1 |
| MRTO4 | CCNP | SLC35B1 | KLRC1 | DOK3 | LAPTM5 | OSCAR | SPRR4 | SLC35F6 |
| EIF2AK2 | PLIN3 | ADCY7 | ACP2 | PPP1R18 | GBP4 | C11orf24 | IRAK1 | LTB4R |
| SERPINB3 | MRPL4 | CD80 | YARS1 | PRIM2 | GFI1 | NETO2 | CMSS1 | IRF9 |
| LAMC2 | NKG7 | DEPDC7 | DSC2 | NFKBIE | SLAMF9 | SHCBP1 | S100A7A | MRPL23 |
| NDC1 | CNFN | PILRB | DSG3 | TPBG | IGSF8 | CHST11 | PKP3 | IFRD2 |
| PARP12 | CDK6 | TNFSF10 | DSC3 | MMS22L | MT2P1 | CLCN5 | XPOT | SMTNL1 |
| OGFR | TFPI2 | GPSM2 | TIMM10 | EGFR | KIF26B | KRT9 | H2BC21 | MIR17HG |
| GNA15 | DNAH11 | PAEP | POGLUT2 | SSC4D | KIAA1841 | LRRC8D | FMNL1 | FAM72D |
| COL11A1 | STEAP1B | MRPS2 | NREP | CCT6A | DPY30 | ECEL1 | USP18 | TNFRSF25 |
| BCAT1 | MPP6 | CBX3 | C9orf40 | PSPH | WDCP | BCL2L1 | TNFAIP8L1 | AL031777.1 |
| MRPS35 | TFEC | INHBA | CTSL | STRA8 | SMC6 | P2RY6 | KRT6B | AL139095.2 |

| | | | | | | | | |
|---|---|---|---|---|---|---|---|---|
| POLD1 | GARS1 | NT5C3A | PSAT1 | ZC3HAV1L | VSNL1 | RGS19 | STAC3 | KIR2DS4 |
| INTS13 | CHCHD2 | CCZ1 | HAVCR2 | NCAPG2 | EN1 | NINJ2 | IRF7 | APOL6 |
| RFXANK | EIF3B | CLTA | OASL | SH3KBP1 | TNFAIP8L2 | C3AR1 | PSMD13 | FADS3 |
| SBNO2 | SNX8 | PLAU | HRK | MSN | SCNM1 | TEFM | KRT79 | CYTOR |
| MCM10 | AIMP2 | PRDX4 | BICDL1 | IL2RG | LCE3D | MRPL13 | IFIT1 | C10orf55 |
| KARS1 | IMPDH1 | NLN | KHDC1 | DOK2 | SPRR2D | ISG20 | WDR53 | AL441992.1 |
| MTHFD2 | LSM5 | OPTN | NAT10 | RGS20 | PGLYRP4 | CLEC7A | IFITM1 | LINC01615 |
| SPI1 | AP1S1 | CDK2 | SEMA4F | GINS4 | S100A9 | COPRS | LCE3E | IFITM3P2 |
| MSANTD3 | RPA3 | NFE2 | CCT7 | FAM83A | S100A12 | PRSS27 | CD300LF | PELATON |
| MTFR1 | POLD2 | TUBA1B | KCNK1 | GSDMC | TGFA | CLP1 | KRT5 | RPL29P19 |
| PFKP | YKT6 | COL10A1 | SP110 | NAPRT | ALPP | PPP1CA | KRT10 | PSME2P2 |
| STOML1 | LIMK1 | NMI | TTLL4 | HAUS6 | CCT3 | RASGRP1 | NKRF | SLC9A3-AS1 |
| PKM | FSD1L | PI3 | ARHGEF4 | CDKN2A | CHCHD4 | CFL1 | KRT3 | HSPA7 |
| IFI35 | SEC61B | SLPI | SLC41A2 | IFNK | SGO2 | MYEOV | CEACAM19 | GBP1P1 |
| FTSJ1 | PLGRKT | KCNS1 | CKAP2 | SIGMAR1 | NUAK2 | HPSE | SPIN4 | PROSER2-AS1 |
| TTC7A | DDX58 | CSE1L | BORA | POLE3 | MNDA | COQ2 | LILRB4 | MIAT |
| GAL | EXOSC3 | PMEPA1 | SCEL | REXO4 | IFI16 | HSPA6 | TNFRSF4 | NOL7 |
| HES2 | TWNK | ZBP1 | NUDT15 | GTF3C5 | AIM2 | RHOD | KRT16 | LGALS17A |
| FSTL3 | NPM3 | XG | LCP1 | NTMT1 | RPL22L1 | MTX1 | KRT14 | KRT16P6 |
| RAD18 | GIT1 | WRNIP1 | IGF2BP3 | PTGES | CTLA4 | PARP14 | TNFRSF18 | NUP50-DT |
| MAP4K4 | PSMD3 | RRP36 | TBRG4 | PROSER2 | GMPS | CKS1B | ESPN | SLC16A1-AS1 |
| HSD17B10 | GALK1 | SNRPC | DDX56 | PDSS1 | CFAP20DC | SYT12 | CMC1 | HSBP1L1 |
| NDE1 | PFN1 | XPO5 | CCM2 | TAF5 | CXCL5 | C1QB | MT1X | AC006460.1 |
| PVR | PSMD11 | NQO2 | MYO1G | ADM | CXCL1 | C1QA | GCNT1 | SNHG26 |
| MCM2 | LGALS3BP | RPS10 | TACO1 | DGKZ | TOPBP1 | OLR1 | FPR3 | LINC02577 |
| IGF2BP2 | CCL7 | H2BC11 | RTP4 | SSRP1 | CDCP1 | NAA20 | HSPA14 | SPRR2C |
| ENO1 | DHX58 | TREM1 | ACTL6A | SERPINH1 | CCR1 | SAA1 | NHLRC1 | HLA-DRB6 |
| MYDGF | LRRC59 | RPP40 | MRPL47 | ST14 | DTX3L | ZNRD2 | DNAH17 | MYOSLID |
| TIPIN | ALOX12 | DEK | IL36G | B3GAT3 | YEATS2 | PC | CENPP | CXCR2P1 |
| NUP37 | EFTUD2 | EREG | TXN | CHEK1 | RPN1 | LRFN4 | PLSCR1 | MAGEA4-AS1 |
| VPS9D1 | FAM20A | MT2A | SMC2 | DSN1 | SLBP | RCE1 | PRR19 | TRPM2-AS |
| FSCN1 | SLC9A3R1 | IRF1 | DPM2 | PLCB3 | MELTF | AP000769.1 | ZP3 | PRRT3-AS1 |
| MCM6 | TMEM104 | SOX9 | PSMB7 | HMGA2 | H2AZ1 | CD7 | PDCD1 | SERPINB9P1 |
| SLC25A43 | NMU | NUP85 | ANP32B | FCGR1A | CCDC51 | CNP | KRTDAP | AL445933.1 |

| | | | | | | | | |
|---|---|---|---|---|---|---|---|---|
| UBE2A | NSD2 | SLC25A1 9 | CTSV | LYPD1 | DUSP7 | HAP1 | FAM83G | LNCARO D |
| LAMP3 | HTATIP2 | DDX31 | LMX1B | PPP1R1C | HMGB2 | RNF213 | NDOR1 | MANCR |
| FAP | CTSC | CLPP | ARPC5L | CCT5 | IL15 | CBX2 | FAM72B | LINC0259 5 |
| TP73 | FOXRED1 | CD70 | PPIL1 | TIMM8B | STARD4 | XXYLT1 | S100A16 | LINC0201 5 |
| SLC1A3 | CCDC86 | RBCK1 | TFAP2A | DHX37 | GRPEL2 | FBXO45 | CALHM6 | DLEU2 |
| SP140 | PRPF19 | AP5S1 | HMGA1 | C12orf45 | GPX8 | TLR6 | KLRG2 | KRT17P3 |
| COL5A3 | PANX1 | SNRPB2 | TCF19 | TEX30 | TERT | ZDHHC2 4 | GJB3 | TAPBP |
| PUM3 | COMMD9 | ITPA | NRM | KCTD14 | CGAS | CTU2 | NELFB | CT69 |
| MOK | SLC15A3 | MCM8 | TLR2 | ADAMTS1 2 | DCBLD1 | C11orf45 | SBSN | DCST1-AS1 |
| RBL1 | NAA40 | EIF2S2 | MYO7A | CDC123 | SFXN1 | SLC26A9 | ADAT2 | MCTS1 |
| PKP1 | SLC35F2 | PSMB2 | IL18BP | NCAPD3 | IL22RA2 | MRPL11 | KIR2DL4 | EMSLR |
| DUSP12 | TCIRG1 | FFAR2 | GGH | ARL11 | IL31RA | IL20RB | RNF222 | TNF |
| AACS | PSMD9 | FRMD8 | DDX60 | PSTPIP2 | KIAA0895 | CMKLR1 | CGB5 | SNHG15 |
| IL12RB2 | MAGOHB | IFI6 | DCUN1D5 | HAUS1 | STK17A | BRMS1 | FAM111B | LINC0046 0 |
| FYB1 | OAS3 | EMG1 | MMP13 | KCNK13 | RELL2 | ODAPH | RRP7A | AL138789 .1 |
| MRPL22 | OAS2 | DLGAP5 | CASP1 | XRCC4 | CTSB | ADGRE1 | S100A14 | HLA-B |
| PALB2 | RASAL1 | L3HYPD H | CASP5 | SUV39H2 | CTHRC1 | YIF1A | TOGARA M2 | MIR155H G |
| HAL | IFNG | TIMM8A | SQOR | TRIM36 | SHOC1 | CNIH2 | SH2D5 | AC007879 .3 |
| CAD | NUP107 | IPPK | BCAR3 | IFIT5 | ZNF367 | CD164L2 | IL1RAP | C12orf75 |
| CD82 | MRPL51 | TRAF2 | IFI44L | PLOD2 | STOML2 | PSMD2 | HLA-DRB1 | LINC0215 4 |
| SNX10 | GAPDH | YEATS4 | IFI44 | CARHSP1 | MELK | CKAP5 | SIRPB2 | ELFN1-AS1 |
| MRPL28 | TPI1 | LRRC61 | PNPT1 | CYRIB | GJB2 | CCNE2 | TUBB | LINC0219 5 |
| BAX | SLCO1B3 | ADGRE2 | PREB | CCSAP | LRR1 | GRAMD2 A | PPIA | LINC0118 6 |
| SH3BP2 | GOLT1B | SYNGR3 | SLC5A6 | GBP5 | TMEM63 C | BANF1 | STK31 | PRKAR1B -AS1 |
| SULT2B1 | DSE | TUBA4A | PARP9 | BUB3 | NUDT5 | PTPN2 | WDR5 | BX276092 .7 |
| DNMT3B | TPD52L1 | PTPN12 | HERC6 | ELOC | QSOX2 | CLTB | ARL9 | FOXD2-AS1 |
| PDRG1 | FBXO5 | PAICS | HERC5 | SLFN13 | TMEM52 B | LRRC25 | SLC6A9 | LINC0085 7 |
| SIGLEC1 | FANCE | PPAT | CXCL9 | USP43 | ENTR1 | DRAP1 | SULF2 | PYY2 |
| DYNLL1 | MCM3 | ADM2 | ANXA3 | FBXL18 | HPRT1 | RPS6KB2 | MMP1 | MAP3K20 -AS1 |
| SIRPG | RNF8 | CDC42EP 1 | SLC39A8 | MOV10 | ISCA2 | RMI2 | RABL6 | TMEM250 |
| FXYD5 | E2F3 | APOL3 | YARS2 | PPM1J | PACSIN3 | RUVBL1 | HLA-DQA1 | HMGN2P 17 |
| LAG3 | WASF1 | APOL2 | SLC7A1 | SLC16A1 | PSMC3 | SFN | COL27A1 | AP001469. 3 |
| ICAM1 | VNN1 | RAC2 | GPR84 | HEATR3 | IFI27L1 | TUBB6 | S100A2 | PSMB9 |
| DNAJB11 | PERP | KRT17 | GALNT6 | SLC7A7 | IFI27 | ZNF683 | H2AC11 | SMKR1 |

| | | | | | | | | |
|---|---|---|---|---|---|---|---|---|
| PUS7 | SLC16A10 | SPECC1 | RHOF | PIK3AP1 | SMCO4 | SPHK1 | ADA | KIR3DL2 |
| PSME1 | BYSL | CPA4 | DENR | RAET1L | TRMT61A | RTTN | CASP4 | LINC00973 |
| GEMIN2 | POLR3G | STRIP2 | DIAPH3 | ADK | CCT2 | LMNB2 | LAGE3 | CDKN2B-AS1 |
| TGM1 | TARS1 | MYO1B | ZIC5 | SH3RF2 | C2 | CDK5R1 | SLC6A17 | HLA-DOB |
| WDR76 | BRIX1 | HOXD11 | WARS1 | MTERF3 | RAG1 | TYMS | SERPINA1 | RPSAP52 |
| UNC13D | NUP155 | CHAC1 | PSTPIP1 | FBXO43 | SERPINB7 | FAM210A | IL27 | SPRR2A |
| VNN3 | SMC4 | CLN6 | BCL2A1 | CD109 | CENPN | MAN1B1 | VEPH1 | PRR34-AS1 |
| TREM2 | CD86 | PSMA1 | PML | UBE2L6 | AKIP1 | SAMD9L | SHPK | HLA-DMB |
| ITPR3 | PDCD10 | CDKN2D | HAPLN3 | TBC1D31 | CCDC68 | ZBED2 | SLC28A3 | AP5Z1 |
| DSP | COL7A1 | SIGLEC9 | RHCG | NTAQ1 | CLEC4E | SLC25A22 | DPP4 | AC082651.3 |
| IL12RB1 | ECT2 | KLK8 | ITGAX | ATAD2 | CDH16 | ATOX1 | PDCD1LG2 | CFB |
| ACOT7 | TFG | PARP2 | IGSF6 | MALSU1 | HSP90B1 | AGTRAP | S100A10 | TMEM199 |
| HSD3B7 | GNB4 | NEDD8 | NLRC5 | RPUSD3 | B2M | POLR2L | H2BC12 | LINC01322 |
| CYTH4 | RRP9 | RHBDF2 | KIFC3 | TMEM171 | SAAL1 | SOX12 | TEAD4 | DUXAP10 |
| SLC5A1 | RTKN | DOHH | LRRC46 | LRP8 | CIAO2A | KCNJ10 | SIRPA | LILRA6 |
| IL2RB | SF3B6 | SAT1 | PTRH2 | SUSD3 | C18orf54 | PCDHB9 | PRIM1 | AC108676.1 |
| RANGAP1 | TP53I3 | APOE | PRELID3A | MX1 | PATL1 | ODF3B | NUP62CL | FCGR2C |
| GZMH | PSMD14 | APOC1 | C18orf21 | FMNL2 | MRPL16 | ZNF114 | TFDP1 | CASC8 |
| GZMB | PPM1G | BST2 | ZMYND15 | MRPL17 | C15orf48 | CALML5 | HYLS1 | SBF2-AS1 |
| GNPNAT1 | IFIH1 | NSUN5 | SLC16A3 | GALNT14 | MS4A14 | BEND3 | MT1F | PGAM5 |
| MIR210HG | AC125807.2 | PINX1 | CLEC5A | AP005233.2 | SNRPGP2 | MIR924HG | AC016394.1 | CCL18 |
| LUCAT1 | LINC01094 | SIGLEC12 | LINC00520 | AL031058.1 | AC138207.4 | DNAH17-AS1 | KLHDC7B-DT | H2BC9 |
| AC022126.1 | FOXD1 | CASP1P2 | AC004816.1 | LINC00165 | ANXA8 | CSAG3 | AC019069.1 | PICSAR |
| AC093895.1 | AC100801.1 | DPP3 | LINC02345 | H2BC20P | TIMM23 | CCL5 | AP001033.2 | ORAI1 |
| LINC00964 | CCDC71L | CARD17 | LINC02323 | LINC00514 | RPL17 | AC026740.1 | LHX1 | RN7SL1 |
| PVT1 | GASAL1 | PRECSIT | AC015660.1 | MMP12 | LINC01910 | AL136162.1 | H2BC8 | H4C9 |
| AC034213.1 | AC007991.2 | LINC02446 | AC004943.2 | AC005722.3 | AC008105.3 | AL138724.1 | H3C8 | AC245884.12 |
| AP002784.1 | LYN | HMBS | LINC01882 | ANXA8L1 | SCAT1 | AL021807.1 | AL161431.1 | AC245041.2 |
| AC002401.4 | BISPR | AL139351.3 | AC018978.1 | AC026401.3 | LCAL1 | H3C10 | AC116407.2 | AC010198.2 |
| AC008760.2 | BX119927.1 | AL021978.1 | C2orf27A | SH3PXD2A-AS1 | AC018865.2 | AL353763.2 | CCL3 | BX470102.2 |
| H2AC8 | LINC02009 | AC083862.2 | AL732437.2 | LINC01127 | AL162253.2 | F8A1 | NKILA | AC108925.1 |
| PIGW | SCO2 | AL353807.5 | AC008875.3 | BLACAT1 | AL137800.1 | LHX1-DT | AL451123.1 | AC103718.1 |
| **1210 exclusive downregulated DEGs for F3-T3 fusion-negative BLCA** | | | | | | | | |
| TSPAN6 | SEMA6A | CNTN3 | MRAP2 | ANKRD50 | KLF15 | TCEAL1 | FAM174B | TMEM170 |

| | | | | | | | | |
|---|---|---|---|---|---|---|---|---|
| | | | | | | | | B |
| GCLC | SEMA4G | BCL6 | ELF5 | EPS8 | LIPH | RND1 | PBX1 | NYNRIN |
| CFTR | CRTAC1 | WNT5A | PAN2 | TMEM45B | BDH2 | ZMAT3 | NTF3 | SERPINB11 |
| ICA1 | TPSD1 | ABCC5 | CAPN9 | SLC25A4 | PITX2 | COL6A5 | POU3F1 | CFAP44 |
| TFPI | CRISP3 | VIPR1 | ITM2C | SERP2 | HPGD | FADS6 | CYP4F12 | LNP1 |
| SARM1 | SYDE2 | ZBTB47 | TSGA10 | AP1S3 | SPINK1 | OVOL1 | ZNF501 | VGLL3 |
| CAMKK1 | PCSK5 | TNNC1 | GCC2 | TLCD4 | RHOBTB3 | ADH6 | INSIG1 | MT-RNR2 |
| TSPOAP1 | CIRBP | FOXP1 | STAB2 | TMEM163 | SERINC5 | ACSM6 | ANKRD20A5P | GPX3 |
| CROT | SERPIND1 | DNAJC27 | VILL | PDE3B | CMYA5 | ABLIM3 | CYP4F8 | Z99572.1 |
| KMT2E | CECR2 | ST3GAL5 | RCBTB2 | GUCY1A2 | FOXQ1 | MMRN2 | TPCN1 | ZNF254 |
| TAC1 | P2RX6 | SLC9A2 | ALPK3 | JMY | SHROOM1 | STOX2 | PPARA | LTC4S |
| ARHGAP44 | SLC7A4 | MLPH | SKIL | GPD1L | LEAP2 | DAB1 | EMID1 | GSTM2 |
| SCIN | TRIOBP | SLC5A7 | RALGPS1 | GJA1 | GPR85 | THAP2 | PLCD1 | ZNF888 |
| ARSD | TTC28 | CD207 | GABBR2 | PRDM8 | SLC29A4 | AGR3 | ENTPD5 | ZNF321P |
| PROM1 | SUN2 | QSOX1 | ENPP2 | BMP3 | HEY1 | PTPRM | ZNF546 | PLAC9P1 |
| BAIAP3 | SYNGR1 | RHOU | TMEM8B | PART1 | ZNF704 | NUDT4 | SESTD1 | EEF1A1P12 |
| MYLIP | CACNA1I | PRDM2 | DENND4C | ACOXL | LMTK2 | GPR160 | LUZP2 | COLCA2 |
| CDKL5 | PVALB | MYCL | PIM1 | MERTK | BAALC | SLCO4C1 | KCNJ11 | LRRC37A4P |
| TFAP2B | UPK3A | SIPA1L2 | FOXF2 | SLC25A27 | RASEF | PIFO | IFT140 | PLIN5 |
| PLEKHG6 | FAM118A | ETV3 | SYTL2 | ADGRF1 | GKAP1 | CTSF | AGMO | EML6 |
| RHOBTB2 | EP300 | ID3 | SORL1 | LGI4 | FBP1 | SNX31 | SAMD11 | FMO9P |
| NISCH | CHADL | PIK3R3 | TMPRSS4 | DDAH1 | PGAP4 | ZBTB4 | C2orf88 | AC110619.1 |
| IDS | TRIM9 | SLC19A2 | C11orf1 | MSI2 | PTCHD1 | RALGAPA1 | DMBT1 | FAM228B |
| ALOX5 | CCDC198 | MAN1C1 | LRRC49 | SEMA3D | PIGA | GTF2IRD2B | WNT7B | MTMR9LP |
| ZMYND11 | CHGA | SLC8A2 | SMAD6 | CHST9 | CLDN3 | MFSD4A | COL4A5 | NPTXR |
| CLCA4 | BDKRB1 | B4GALT6 | SPTBN5 | CDH12 | ATP7A | GOLT1A | NHS | LINC02593 |
| VSIG2 | DHRS2 | IRAG2 | GCOM1 | ABCA9 | AQP7 | TMEM51-AS1 | ZNF626 | OSER1-DT |
| PLEKHB1 | CPNE6 | CYP20A1 | BTBD16 | ABCA6 | ARHGAP12 | ABO | SMTNL2 | SNHG14 |
| NRXN3 | SGK2 | GDA | PLCE1 | ABCA5 | FAT3 | FAM131A | ZC3H6 | BCAR3-AS1 |
| ALAS1 | BMP7 | TJP2 | CARF | C4orf19 | PKNOX2 | SCUBE2 | ZNF559 | HAGLR |
| IKZF2 | HRH3 | SLC46A2 | IDH1 | DISP1 | OXGR1 | DPP10 | HBA2 | LINC00885 |
| LRRC7 | EEF1A2 | ALDH6A1 | SLC40A1 | TDH | NDRG2 | LONRF3 | RALGAPA2 | SH3BP5-AS1 |
| MYOC | CCM2L | RHOQ | SENP7 | OBSCN | ARHGEF40 | GPX2 | FSIP2 | NRSN2-AS1 |
| CASR | TNNC2 | PPP1R3C | SECISBP2L | LONRF1 | PPP1R36 | CIDEA | FAM78B | TRAM2-AS1 |

| | | | | | | | | |
|---|---|---|---|---|---|---|---|---|
| TUBG2 | VSIG1 | PYROXD2 | APH1B | SH3RF1 | BTNL9 | ZNF135 | ASTL | SGMS1-AS1 |
| TG | ZC3H12B | HOXB8 | FAM13A | CHODL | SALL2 | PRR15 | C9orf152 | NPY6R |
| ADAM28 | PCYT1B | HOXB3 | PRKG2 | XPC | CACNB2 | FOXL1 | APOD | AC005082.1 |
| ADRB1 | SRPX2 | MSX2 | KIAA1109 | FGD5 | TCP11L2 | TCIM | CLDN4 | LINC02672 |
| PREX2 | BEX4 | PLEKHG1 | ERP27 | PIEZO2 | CKB | GCNT4 | BTBD8 | CYP4F29P |
| FAM214A | NALCN | CYSTM1 | SLC38A4 | SPAG17 | LARP6 | ACER2 | MAOA | SDAD1P1 |
| ARHGAP6 | CAB39L | SOHLH2 | PTPRQ | DEPTOR | BORCS7 | ANO6 | FHIT | EEF1A1P11 |
| LMO3 | VWA8 | KBTBD7 | SUOX | PPARGC1B | CYB5A | RIMKLA | MYBPC1 | EMX2OS |
| COL9A2 | DGKH | PLS1 | MAP3K12 | PSD3 | USP54 | CASKIN2 | SPOCK3 | HSPD1P6 |
| H6PD | NFAT5 | CLU | SLAIN1 | FAM161B | ATP9B | ERICH5 | MYT1 | PSMG3-AS1 |
| NEDD4L | NDRG4 | TBX2 | TTC6 | WIF1 | PPFIBP2 | RPRM | ACADSB | AC005077.4 |
| FSTL4 | FA2H | FAM117A | RAB15 | CLDN8 | MOGAT2 | SAMD12 | STK40 | LINC02541 |
| NNAT | LMF1 | ZNF211 | ESR2 | TIAM1 | CENPV | CRACR2B | CTSE | KLF3-AS1 |
| NRIP2 | SALL1 | ADGRB2 | DUOXA2 | PPP2R2B | ATF7IP2 | NAALADL2 | EEF1A1P5 | LINC00689 |
| KCNQ1 | TOX3 | ACVR2A | DUOX2 | KAT6B | GOLM2 | DPY19L2 | SRGAP3 | FAHD2CP |
| HHAT | RBL2 | EEPD1 | DNAJA4 | RAB11FIP1 | ACSM1 | GRAMD1C | ZNF471 | AL157786.1 |
| PLEKHH1 | SYT17 | NUDT10 | ARRDC4 | MPV17L | ZNF667-AS1 | GALNT11 | GTF2IRD2 | CYP1B1-AS1 |
| ATP9A | HERC1 | BHLHE41 | CYP1A1 | SST | KIF7 | WDR6 | NLGN3 | MIR3936HG |
| KCNH2 | CTSH | MMP19 | ABHD2 | ZFYVE9 | PLIN1 | P4HTM | ZNF781 | DIRC3-AS1 |
| CYFIP2 | FAM189A1 | BAZ2B | CDR2 | RBPMS | PEX11A | CA8 | ZNF774 | EEF1A1P6 |
| GYG2 | ZDHHC2 | PLA2G12A | ZFHX3 | STEAP2 | SCNN1G | EPM2AIP1 | ARMCX4 | CYP2T1P |
| EIF4B | NIPAL2 | OBSL1 | ZNF287 | CACNA1D | ANKDD1A | ERN1 | ESRRG | LINC02765 |
| LIMCH1 | ZC2HC1A | CHD6 | TOB1 | KIT | TAC3 | CSRNP3 | TCEAL3 | AC005064.1 |
| BCAS1 | SH2D4A | IL9R | ARSG | PWWP3B | CHP2 | FAM219B | TPK1 | TSPY26P |
| GLP2R | NEFM | ATP8A1 | GAREM1 | C2CD2 | MTMR10 | DPY19L3 | SPTSSB | ITGA9-AS1 |
| TLE2 | ASAH1 | POF1B | SLC14A1 | BRAF | SGSM1 | ZNF552 | ZNF846 | AC019117.2 |
| FAM107B | DMPK | TRERF1 | RAB40B | CABP1 | UGT1A6 | TMTC2 | UGT2B15 | TRHDE-AS1 |
| TBC1D1 | NOVA2 | CLYBL | IGFBP4 | SLC30A2 | TBC1D2B | FUCA1 | ZNF136 | ZNF853 |
| MPPED2 | TJP3 | ABCC4 | TMEM50B | TRIM63 | IGF2 | GATA2 | ALKAL1 | MIR600HG |
| PDK3 | NOP53 | SRMS | RERE | GRHL3 | DPEP2 | ZBTB18 | MAML3 | LINC02814 |
| INPP5A | SULT2A1 | FNDC11 | PADI3 | SLC13A3 | CA4 | ARL14 | MVB12B | NR2F1-AS1 |
| MAST4 | MEGF8 | AMOT | NBPF3 | SHROOM4 | TMC4 | GIPC3 | ZFP28 | RNF223 |

| | | | | | | | | |
|---|---|---|---|---|---|---|---|---|
| ADGRF5 | CAPS | MCF2L | CYP4B1 | NRG2 | CDC42EP5 | TDRP | KPNA5 | ZNF737 |
| PLA2G10 | BCAT2 | DACH2 | SLC44A3 | PPP1R9A | PPP1R14A | SLC9A4 | ZNF429 | C10orf143 |
| MNT | RAB3A | HSPA2 | RGS5 | DYNC1I1 | PSCA | ZNF816 | ZNF470 | SHISA9 |
| ST6GALNAC1 | UPK1A | PTPRB | CASQ1 | NPM2 | PLIN4 | ZFP3 | SERTAD1 | AC108058.1 |
| RPS6KA6 | TSPAN12 | CASD1 | CGN | VWA5B1 | CYB5D2 | CXCR2 | SIGLEC15 | BX470102.1 |
| LNX1 | HOXA13 | KDR | MINDY1 | EPB41 | GGT6 | RPH3AL | PGAP1 | KLHL41 |
| TRHDE | EVX1 | MGAT3 | SELENBP1 | SV2A | HID1 | SLC25A42 | ACSL5 | AMY2B |
| CRMP1 | EPHB6 | SNRPN | SUSD4 | CLIC6 | TMEM88 | TNFSF15 | NOL4L | ZNF542P |
| SIDT1 | CYP3A5 | ALDH1A2 | SCCPDH | ALDH4A1 | VWCE | ZBTB20 | FAM110D | AL450384.2 |
| FRY | ZNF862 | VPS13C | CNIH3 | ZFYVE28 | ENTPD3 | DIPK2A | EIF4BP6 | NSUN6 |
| NTN4 | ANKMY2 | MTUS1 | EPHX1 | ABR | LTBP3 | AMIGO1 | GATA3-AS1 | UGT1A1 |
| ARHGEF10L | AGR2 | FOXA1 | PLEKHA6 | ZNF222 | SCARA3 | TMEM30B | PELI1 | PEG10 |
| TXK | AHR | ARHGEF6 | OSR1 | GNE | LMBRD1 | CREB3L2 | FBXL22 | UGT1A8 |
| WSCD2 | CLIP2 | GAMT | GDF7 | FNDC5 | KIF5C | TSHZ2 | SLC22A5 | UGT1A10 |
| TACR2 | NPDC1 | KIF1A | KANSL1L | CPAMD8 | PCMTD1 | SATB1 | SLC2A10 | CCDC169 |
| SEMA3C | APBA1 | PGPEP1 | SPAG16 | CILP2 | DEGS2 | TRAK1 | COL4A6 | STON1 |
| FOSL2 | ABHD17B | IQCN | ACKR3 | TFF3 | SCN11A | PLCB1 | INKA2 | UPK3B |
| RGS11 | GATA3 | JUND | GASK1A | PDE9A | KLHL30 | TTC3 | ADAMTSL2 | HOTTIP |
| MBNL3 | ATRNL1 | HRC | ZNF660 | ZNF66 | SCNN1B | IGIP | ZNF677 | GSTA1 |
| RAP1GAP | DNMBP | PNCK | ALDH1L1 | ADAMTS13 | MTCL1 | RGS6 | GOLGA6L9 | IFITM10 |
| TOP2B | ATE1 | ZNF331 | ILDR1 | AZGP1 | SLC16A4 | TMEM198B | ZNF334 | WNT5A-AS1 |
| CAPN6 | CDH23 | COX4I2 | VWA5B2 | SQSTM1 | GSTM4 | PLCXD3 | SLC29A3 | HBB |
| NAALAD2 | DKK1 | GGT7 | TBCK | FBXO27 | CXXC4 | ZNF662 | SH3BGRL2 | AP000866.1 |
| MCCC1 | ZMIZ1 | EDA2R | USP53 | ACOX1 | NSG1 | ZNF320 | ZNF808 | CEBPA |
| ZCWPW1 | PBLD | SH3BGRL | NKD2 | PLCD3 | ATOH8 | PCP4 | ANKRD35 | ALDH1L1-AS2 |
| PCM1 | RAPGEFL1 | SH3BP5 | BHMT | ADCY10P1 | PLA2G4F | ABAT | CNGA1 | AP002026.1 |
| TNRC6C | EFNB3 | RAMP2 | CXCL14 | HS3ST6 | CA5B | CAMK1D | LRBA | LINC00900 |
| CDC14A | TMEM97 | AOC2 | KLHL3 | ZG16B | STK32A | IGSF5 | FAM3D | UBAP1L |
| LIPE | MAPK10 | TMEM204 | TNFRSF21 | BEND5 | SCN9A | BEGAIN | MT-ND6 | LINC00926 |
| CRYBG3 | WFS1 | PPP1R1B | SCUBE3 | ZSWIM5 | RASSF6 | CTNNA3 | CEP290 | RRN3P1 |
| KCNN2 | ANXA10 | KHDRBS3 | SDK1 | DHRS3 | SDC2 | BCOR | MT-CYB | ADH1C |
| ARG2 | CLCN3 | FMO5 | TMEM168 | BRINP3 | GKN1 | CBX6 | F5 | FAM198B-AS1 |
| PHLPP1 | TRIM2 | ZNF132 | NLGN4X | CNST | NRG4 | B3GALT5 | MT-ND5 | PCP4L1 |
| ATP8B1 | RAPGEF2 | SPATA6 | TMEM47 | PM20D1 | TACR3 | ZNF703 | MUC2 | SRP14-AS1 |

| | | | | | | | | |
|---|---|---|---|---|---|---|---|---|
| SMARCD3 | KLF3 | PPARG | ITGB1BP2 | PIGR | SLC16A5 | FAM3B | SCAMP5 | SMIM31 |
| C1QTNF3 | PPARGC1A | KDM6B | ZNF711 | REL | FABP4 | TOB2 | MT-CO1 | AL691482.3 |
| COBLL1 | SC5D | DLG4 | RADX | CAPN13 | EMX2 | SCN5A | MT-ND3 | AL033397.2 |
| DOP1A | B3GAT1 | MTSS2 | GPC3 | KCNF1 | GPRC5C | HTR7P1 | MT-ND1 | AC026369.1 |
| EFR3B | ST3GAL4 | BTBD3 | SNTG1 | LRATD1 | NPAS2 | TMPRSS2 | DZIP3 | HS3ST5 |
| MAPRE3 | UPK2 | ATP1A4 | SHC3 | FRZB | LONRF2 | ADRA2C | SOWAHA | DNAH10OS |
| MECOM | P3H3 | CASP9 | ASTN2 | DUSP19 | TMC7 | GPR173 | MFAP3L | FZD10-AS1 |
| OVGP1 | LIN7A | MTUS2 | ADD3 | SLC16A14 | SLC26A5 | PRKD1 | CAPN8 | SNHG18 |
| HSD17B2 | PPM1H | PIK3C2B | INA | NOSTRIN | RASA4B | CCSER1 | MIR29B2CHG | GMDS-DT |
| HSD17B14 | FZD10 | RNF128 | CAPN5 | INHBB | KIAA0232 | SRPK3 | SAMD5 | ZNF345 |
| PIR | TBC1D30 | TBC1D8B | PLCH2 | MSX1 | PKHD1 | PLA2G6 | RBM20 | AC107884.1 |
| RFX2 | CHPT1 | LARGE1 | ESAM | BNIPL | PKIA | MAML2 | LINC00963 | AC104825.1 |
| ASAP3 | SASH1 | MYO18B | DUSP15 | NIPAL1 | INSR | NCMAP | FOXO6 | TRNP1 |
| FER1L4 | COL9A1 | SLC2A11 | TM7SF2 | DAPL1 | ZNF439 | SLITRK6 | CHIC1 | AC022893.1 |
| EPB41L1 | GPLD1 | SARAF | FXYD4 | LMOD3 | ZNF440 | KRBA2 | NHSL2 | CLDN23 |
| PPP1R13B | ALDH5A1 | DGLUCY | CTF1 | SLC15A2 | KRT13 | B3GALT5-AS1 | NOTCH4 | AC100810.1 |
| C20orf194 | ARFGEF3 | MEIS2 | HNMT | ELF3 | KRT20 | ZBTB7C | PRRT1 | KRTAP5-9 |
| TESC | PACRG | GSTM3 | PDCD4 | IHH | COL24A1 | DRD1 | SLC44A4 | AF131215.5 |
| SLC23A2 | BICRAL | HMGCS2 | FREM2 | PPM1L | MAP6 | ARMCX2 | ACOXL-AS1 | B3GAT1-DT |
| CHGB | ENPP5 | SORT1 | THRB | ICA1L | LGALS4 | TMEM121 | TRIM31 | CYP2A6 |
| KCNH4 | SEMA5A | ERN2 | ANK3 | WDFY3 | CYP4F22 | MROH2A | ZNF425 | ZNF10 |
| ANKRD24 | CERT1 | CYP2J2 | DIP2C | PTPN13 | MAL | MFSD6L | SPIRE2 | ZNF486 |
| RAB11FIP3 | RASGRF2 | CEMIP2 | MAGI1 | SMIM14 | FGGY | NOTUM | TRIQK | OR13A1 |
| NLRP1 | CDH6 | TBX3 | MIPOL1 | FYCO1 | IL17D | HS6ST3 | ZBTB10 | CYP2B7P |
| JPH4 | FAM172A | CCDC146 | ME3 | DNALI1 | FUT9 | METTL7A | LGR4 | CAPNS2 |
| AC008014.1 | AL158206.1 | LRRC37A16P | ERVV-1 | AL035661.1 | AC084759.3 | INAFM2 | LOXL1-AS1 | AC011487.2 |
| AC068888.1 | TMEM178B | ERVE-1 | IPO5P1 | ZNF658 | AL158175.1 | CTXND1 | LINC01569 | MTRNR2L12 |
| RHOXF1-AS1 | AC021016.1 | TBX2-AS1 | AF131215.6 | FCGBP | AP000547.3 | IQCH-AS1 | AC090617.5 | AC024075.2 |
| LINC00930 | AP003119.3 | CEBPA-DT | ATP2B1-AS1 | HNF1B | AC068587.4 | CCDC187 | MYZAP | TRABD2B |
| AL133370.1 | GAN | NBPF12 | AL691432.2 | AC007998.4 | AC092329.4 | LINC02562 | MSMB | AC011503.1 |
| SMIM6 | AC036108.3 | AC010329.1 | AC067750.1 | SRCIN1 | AC105233.4 | AC091544.4 | LINC00672 | CYP4F23P |
| AL670729.3 | AC003681.1 | AC023421.2 | SAP30-DT | AC007906.2 | AC092053.3 | ERVK13-1 | TXNIP | AC011455.8 |
| AC006230. | AP002800. | AC133530 | AL391121.1 | SRD5A2 | AP003498. | AL121839 | BAHCC1 | AL499602 |

| | | | | | | | | |
|---|---|---|---|---|---|---|---|---|
| 1 | 1 | .1 | | | 2 | .2 | | .1 |
| AC127526.5 | AL603750.1 | AL136169.1 | AC008124.1 | AC010998.3 | AC055839.2 | LINC01963 | MYO15B | LINC01451 |
| AC108519.1 | AC027290.2 | AC079848.2 | AL135999.3 | | | | | |

**Supplementary Table S3. The identified significant GO terms of the genes of modules in each dataset.**

| Enriched GO terms of upregulated DEGs of module in fusion-positive BLCA samples | | | | |
|---|---|---|---|---|
| | Category | Term | Genes | PValue |
| **Module1** | GOTERM_BP_DIRECT | GO:0042776~mitochondrial ATP synthesis | NDUFA13, NDUFS5, NDUFA3, NDUFA1, ATP5F1D, ATP5ME | 4.43E-11 |
| | GOTERM_BP_DIRECT | GO:0009060~aerobic respiration | NDUFA13, NDUFS5, NDUFA3, NDUFA1, ATP5F1D | 1.69E-08 |
| | GOTERM_BP_DIRECT | GO:1902600~hydrogen ion transmembrane transport | UQCR10, COX5B, ATP5F1D, ATP5ME, COX6B1 | 3.23E-07 |
| | GOTERM_BP_DIRECT | GO:0032981~mitochondrial respiratory chain complex I | NDUFA13, NDUFS5, NDUFA3, NDUFA1 | 2.94E-06 |
| | GOTERM_BP_DIRECT | GO:0045333~cellular respiration | UQCR10, COX5B, COX6B1 | 1.62E-04 |
| | GOTERM_CC_DIRECT | GO:0070469~respiratory chain | NDUFS5, NDUFA1 | 0.014429418 |
| | GOTERM_CC_DIRECT | GO:0005753~mitochondrial proton-transporting | ATP5F1D, ATP5ME | 0.010120396 |
| | GOTERM_CC_DIRECT | GO:0005747~mitochondrial respiratory complex I | NDUFA13, NDUFS5, NDUFA3, NDUFA1 | 1.58E-06 |
| | GOTERM_CC_DIRECT | GO:0005751~mitochondrial respiratory complex IV | NDUFA4L2, COX5B, COX6B1 | 6.29E-05 |
| | GOTERM_CC_DIRECT | GO:0031966~mitochondrial membrane | NDUFA13, NDUFA1, COX5B, COX6B1 | 6.30E-05 |
| | GOTERM_MF_DIRECT | GO:0004129~cytochrome-c oxidase activity | COX5B, COX6B1 | 0.009447757 |
| | GOTERM_MF_DIRECT | GO:0046933~proton-transporting ATP synthase activity | ATP5F1D, ATP5ME | 0.008977261 |
| | GOTERM_ MF _DIRECT | GO:0008137~NADH dehydrogenase (ubiquinone) activity | NDUFA13, NDUFS5, NDUFA3, NDUFA1 | 1.04E-06 |
| **Module2** | GOTERM_ BP_DIRECT | GO:0002181~cytoplasmic translation | RPS15, RPL14, RPL37, RPL18, RPL29, RPL28 | 1.20E-11 |
| | GOTERM_ BP_DIRECT | GO:0006412~translation | RPS15, RPL14, RPL37, RPL18, RPL29, RPL28 | 1.10E-09 |
| | GOTERM_ BP_DIRECT | GO:1901798~positive regulation of signal transduction | RPS15, RPL37 | 0.002771788 |
| | GOTERM_ BP_DIRECT | GO:0006364~rRNA processing | RPS15, RPL14 | 0.041802634 |
| | GOTERM_ | GO:0005829~cytosol | RPS15, KCNS3, RPL14, | 4.08E-04 |

| | | | | |
|---|---|---|---|---|
| | CC_DIRECT | | RPL37, RPL18, RPL29, RPL28 | |
| | GOTERM_ CC_DIRECT | GO:0022626~cytosolic ribosome | RPS15, RPL14, RPL37, RPL18, RPL29, RPL28 | 9.91E-12 |
| | GOTERM_ CC _DIRECT | GO:0005840~ribosome | RPS15, RPL14, RPL37, RPL18, RPL29, RPL28 | 2.83E-10 |
| | GOTERM_ CC_DIRECT | GO:0022625~cytosolic large ribosomal subunit | RPL14, RPL37, RPL18, RPL29, RPL28 | 1.10E-09 |
| | GOTERM_ CC_DIRECT | GO:0005737~cytoplasm | RPS15, RPL14, RPL37, RPL18, RPL29, RPL28 | 0.007980592 |
| | GOTERM_ MF _DIRECT | GO:0003723~RNA binding | RPS15, RPL14, RPL37, RPL18, RPL29, RPL28 | 1.63E-05 |
| | GOTERM_ MF _DIRECT | GO:0003735~structural constituent of ribosome | RPS15, RPL14, RPL37, RPL18, RPL29, RPL28 | 5.84E-10 |
| | GOTERM_ MF _DIRECT | GO:1990948~ubiquitin ligase inhibitor activity | RPS15, RPL37 | 0.002842702 |
| | GOTERM_ MF _DIRECT | GO:0097371~MDM2/MDM4 family protein binding | RPS15, RPL37 | 0.003788771 |
| **Module3** | GOTERM_ BP_DIRECT | GO:0036148~phosphatidylglycerol acyl-chain remodeling | PLA2G2F, PLA2G4F | 0.002464444 |
| | GOTERM_ BP_DIRECT | GO:0036150~phosphatidylserine acyl-chain remodeling | PLA2G2F, PLA2G4F | 0.002926076 |
| | GOTERM_ BP_DIRECT | GO:0036152~phosphatidylethanolamine acyl-chain remodeling | PLA2G2F, PLA2G4F | 0.003233752 |
| | GOTERM_ BP_DIRECT | GO:0036151~phosphatidylcholine acyl-chain remodeling | PLA2G2F, PLA2G4F | 0.003541364 |
| | GOTERM_ BP_DIRECT | GO:0050482~arachidonic acid secretion | PLA2G2F, PLA2G4F | 0.004156399 |
| | GOTERM_ MF _DIRECT | GO:0047498~calcium-dependent phospholipase A2 activity | PLA2G2F, PLA2G4F | 0.002843151 |
| | GOTERM_ MF _DIRECT | GO:0004623~phospholipase A2 activity | PLA2G2F, PLA2G4F | 0.005523396 |
| | GOTERM_ MF _DIRECT | GO:0016705~oxidoreductase activity | CYP2J2, CYP4F8 | 0.009613298 |
| | GOTERM_ MF _DIRECT | GO:0004497~monooxygenase activity | CYP2J2, CYP4F8 | 0.01243819 |
| | GOTERM_ MF _DIRECT | GO:0005506~iron ion binding | CYP2J2, CYP4F8 | 0.022750164 |
| **Enriched GO terms of downregulated DEGs of module in fusion-positive BLCA samples** | | | | |
| **Module1** | GOTERM_ BP_DIRECT | GO:0006955~immune response | CD86, CD274, IL15, GZMA, CXCR6, CD1D, PDCD1LG2, IL2RG, TNFSF13B, ZAP70, FCGR3A, IL2RA, CCL5, CXCR3, CCL4, FAS, CCL2, CTLA4, JAK2, CCR5, IL7R, ICOS, B2M, CCR2 | 1.62E-21 |

| | | | | |
|---|---|---|---|---|
| | GOTERM_ BP_DIRECT | GO:0007166~cell surface receptor signaling | CD86, CD274, LAG3, PDCD1LG2, CD3E, TNFSF13B, CD2, FCGR3A, IL2RA, CXCR3, CD28, CD27, CCL2, KLRD1, CD247, CCR5, IL7R | 1.80E-15 |
| | GOTERM_ BP_DIRECT | GO:0002250~adaptive immune response | CD86, CD274, LAG3, PDCD1LG2, PRDM1, CD3E, ZAP70, BTLA, CTLA4, KLRD1, PDCD1, CD247, JAK2, HAVCR2, SLAMF1 | 6.44E-11 |
| | GOTERM_ BP_DIRECT | GO:0006954~inflammatory response | CSF1R, IL15, ITGB2, CXCR6, ITGAL, IL6, IL2RA, CCL5, CXCR3, CCL4, CCL2, CCR5, HAVCR2, CCR2 | 4.76E-10 |
| | GOTERM_ BP_DIRECT | GO:0007165~signal transduction | CD274, CSF1R, CD83, IL15, STAT1, ITGAL, IL2RG, TNFSF13B, TYROBP, IL2RB, CCL4, FAS, CCL2, CD38, CD226, JAK2, CCR5, IL7R | 1.99E-07 |
| | GOTERM_ CC_DIRECT | GO:0005886~plasma membrane | CD86, CD274, CSF1R, CD83, ITGAM, ITGB2, PRF1, CD1D, CXCR6, CD3E, ITGAL, IL2RG, TNFSF13B, ICAM1, FCGR3A, CXCR3, BTLA, ITGAX, CTLA4, CD38, JAK2, CCR5, ICOS, TIGIT, B2M, CCR2, CD52, ENTPD1, LAG3, ITGA4, PDCD1LG2, CD2, ZAP70, TYROBP, CD5, LCK, IL2RA, IL2RB, CD28, FAS, CD27, CD48, CD226, KLRD1, PDCD1, CD247, IL7R | 1.71E-16 |
| | GOTERM_ CC_DIRECT | GO:0016021~integral component of membrane | CD86, CD274, CSF1R, CD83, ITGAM, PRF1, CD1D, CXCR6, CD3E, ITGAL, IL2RG, TNFSF13B, ICAM1, FCGR3A, CXCR3, CTLA4, CD38, CCR5, ICOS, TIGIT, HAVCR2, CCR2, SLAMF1, CD52, ENTPD1, LAG3, ITGA4, PDCD1LG2, CD2, TYROBP, CD5, IL2RA, IL2RB, FAS, CD48, CD226, KLRD1, PDCD1, CD247, IL7R | 2.33E-10 |
| | GOTERM_ CC_DIRECT | GO:0009897~external side of plasma membrane | CD86, CD274, CD83, ITGAM, ITGB2, CD1D, CXCR6, CD3E, ITGAL, IL2RG, | 1.19E-38 |

| | | | | |
|---|---|---|---|---|
| | | | ICAM1, FCGR3A, CXCR3, ITGAX, CTLA4, CCR5, B2M, CCR2, SLAMF1, LAG3, ITGA4, PDCD1LG2, CD2, CD5, IL2RA, IL2RB, CD28, FAS, CD27, CD48, CD226, KLRD1, PDCD1, IL7R | |
| | GOTERM_ CC_DIRECT | GO:0009986~cell surface | CD86, CSF1R, ITGAM, ITGA4, IL15, ITGB2, CD1D, ITGAL, IL2RG, ICAM1, CD2, TYROBP, CXCR3, IL2RB, ITGAX, CD28, FAS, CD38, CD226, TIGIT, CCR5, HAVCR2, SLAMF1 | 2.33E-18 |
| | GOTERM_ CC_DIRECT | GO:0016020~membrane | CD52, ENTPD1, ITGA4, ITGB2, PRF1, GZMB, ITGAL, IL2RG, TNFSF13B, ICAM1, TYROBP, IRF4, CD5, IL2RB, ITGAX, CD28, FAS, CD38, CD48, JAK2, B2M, CCR2 | 8.92E-04 |
| | GOTERM_ MF_DIRECT | GO:0005515~protein binding | CD86, CD83, ITGAM, ITGB2, PRF1, PRDM1, CD3E, ITGAL, TNFSF13B, ICAM1, FCGR3A, ITGAX, CTLA4, JAK2, CCR5, B2M, HAVCR2, CCR2, ENTPD1, LAG3, ITGA4, IL15, PDCD1LG2, ZAP70, TYROBP, IRF4, LCK, CD226, CD48, CSF1R, CD274, CD1D, IL2RG, CXCR3, CCL5, CCL4, BTLA, STAT4, CCL2, ICOS, TIGIT, SLAMF1, STAT1, GZMA, GZMB, CD2, IL6, CD5, IL2RA, IL2RB, CD28, CD27, FAS, KLRD1, PDCD1, CD247, IL7R | 2.69E-08 |
| | GOTERM_ MF_DIRECT | GO:0038023~signaling receptor activity | CD2, CD86, CD5, CXCR3, ITGB2, BTLA, ITGAX, FAS, CD48, ICAM1, SLAMF1 | 2.09E-09 |
| | GOTERM_ MF_DIRECT | GO:0004888~transmembrane signaling receptor | FCGR3A, LAG3, FAS, CD27, KLRD1, CD247, CD3E, ICAM1, SLAMF1 | 1.68E-07 |
| | GOTERM_ MF_DIRECT | GO:0042802~identical protein binding | STAT1, PRF1, CD3E, CD2, TYROBP, LCK, CCL5, CCL4, CD28, STAT4, FAS, CD38, CD226, CD247, TIGIT, JAK2, CCR5, B2M, CCR2, SLAMF1 | 4.89E-07 |

| | | | | |
|---|---|---|---|---|
| | GOTERM_ MF_DIRECT | GO:0005102~receptor binding | CD2, ZAP70, TYROBP, LCK, CCL2, TIGIT, JAK2, TNFSF13B | 2.50E-04 |
| **Module 2** | GOTERM_ BP_DIRECT | GO:0006955~immune response | CXCL2, CX3CL1, CTSS, SPN, CD36, HLA-DPA1, CD74, CCL21, IL18, HLA-B, HLA-A, LILRB2, TNFRSF1B, HLA-E, TLR1, CD4, CD8B, CD8A, CD209, TNFSF4, CD7, PXDN, IRF8, HLA-DRA, LCP2, HLA-DRB1 | 6.59E-23 |
| | GOTERM_ BP_DIRECT | GO:0002250~adaptive immune response | ITK, SYK, SH2D1A, HLA-B, LILRB1, CD3G, HLA-A, LILRB2, CTSS, HLA-E, CD4, CD6, CD8B, CD209, CD8A, CD7, BTK, HLA-DRA, SLAMF7, HLA-DRB1, HLA-DPA1 | 2.46E-17 |
| | GOTERM_ BP_DIRECT | GO:0006954~inflammatory response | CSF1, CCL21, STAT3, IL18, CYBB, NKG7, TNFRSF1B, AIF1, CXCL2, CX3CL1, TLR1, CLEC7A, TNFSF4, C3AR1, TLR8, NLRP3, SIGLEC1, TLR3 | 6.09E-14 |
| | GOTERM_ BP_DIRECT | GO:0045087~innate immune response | FCER1G, SYK, CSF1, SH2D1A, HLA-B, CYBB, HLA-A, HLA-E, TLR1, CD6, CD209, BTK, TLR8, NLRP3, SLAMF6, TLR3 | 2.46E-09 |
| | GOTERM_ BP_DIRECT | GO:0007165~signal transduction | ITK, ANXA5, STAT3, LILRB1, LILRB2, SPN, TLR1, CD4, TNFSF4, CASP1, TNFRSF8, NLRP3, IL12RB1, CD33, HLA-DRB1, TLR3 | 2.50E-05 |
| | GOTERM_ CC_DIRECT | GO:0005886~plasma membrane | ITK, CSF1, CD3G, CX3CL1, SPN, MRC1, CASP1, C3AR1, TNFRSF8, SIRPA, CD36, CD33, HLA-DPA1, IL15RA, FCER1G, SYK, HLA-B, CYBB, HLA-A, TNFRSF1B, HLA-E, TLR1, CD8B, CD8A, BTK, TLR8, TLR3, CLEC7A, IL21R, SLAMF7, SLAMF6, IL12RB1, CD74, CD163, IL10RA, STAT3, FN1, NKG7, LILRB1, LILRB2, CD4, PTPRC, FCGR2A, CD6, CD209, TNFSF4, CD7, HLA-DRA, | 6.82E-16 |

| | | | | |
|---|---|---|---|---|
| | | | SIGLEC1, HLA-DRB1 | |
| | GOTERM_ CC_DIRECT | GO:0016021~integral component of membrane | SPI1, CSF1, CD3G, CX3CL1, SPN, CLEC7A, MRC1, C3AR1, IL21R, SIRPA, SLAMF7, TNFRSF8, SLAMF6, CD36, CD33, HLA-DPA1, CD74, IL15RA, IL33, CD163, FCER1G, IL10RA, HLA-B, CYBB, NKG7, LILRB1, HLA-A, LILRB2, TNFRSF1B, HLA-E, TLR1, CD4, PTPRC, FCGR2A, CD6, CD8B, CD8A, CD209, TNFSF4, CD7, TLR8, HLA-DRA, SIGLEC1, HLA-DRB1, TLR3 | 1.06E-11 |
| | GOTERM_ CC_DIRECT | GO:0009897~external side of plasma membrane | CD74, CD163, FCER1G, ANXA5, HLA-B, LILRB1, CD3G, HLA-A, HLA-E, SPN, CD4, PTPRC, CD6, CLEC7A, CD209, CD8A, IL21R, TLR8, SLAMF7, SLAMF6, CD36, IL12RB1, CD33, HLA-DRB1 | 2.50E-21 |
| | GOTERM_ CC_DIRECT | GO:0005887~integral component of cell membrane | CD3G, SPN, MRC1, C3AR1, SIRPA, CD36, CD33, HLA-DPA1, CD163, FCER1G, IL10RA, HLA-B, CYBB, NKG7, HLA-A, LILRB2, TLR1, CD4, PTPRC, FCGR2A, CD6, CD8B, CD8A, TNFSF4, HLA-DRA, HLA-DRB1, TLR3 | 9.36E-14 |
| | GOTERM_ CC_DIRECT | GO:0016020~membrane | CSF1, CX3CL1, SPN, CLEC7A, IL21R, SIRPA, NLRP3, CD36, CD74, CD163, ANXA5, HLA-B, HLA-A, LILRB2, TNFRSF1B, GZMH, HLA-E, TLR1, CD4, PTPRC, CD6, CD209, CD7, SIGLEC1, HLA-DRB1, TLR3 | 9.65E-05 |
| | GOTERM_ MF_DIRECT | GO:0005515~protein binding | ITK, SPI1, CSF1, CD3G, IKZF1, CXCL2, CX3CL1, SPN, GNLY, MRC1, CASP1, CD36, CD33, IL15RA, FCER1G, SYK, ANXA5, IL18, HLA-B, CYBB, HLA-A, TNFRSF1B, HLA-E, TLR1, CD8B, CD8A, IRF1, BTK, IRF8, LCP2, TLR3, AIF1, CLEC7A, IL21R, NLRP3, | 0.001994507 |

| | | | | |
|---|---|---|---|---|
| | | | SLAMF6, CD74, IL33, CD163, CCL21, IL10RA, SH2D1A, STAT3, FN1, NKG7, LILRB1, LILRB2, CD4, PTPRC, FCGR2A, CD6, CD209, TNFSF4, CD7, HLA-DRA, HLA-DRB1 | |
| | GOTERM_ MF_DIRECT | GO:0004888~transmembrane signaling activity | SPN, TLR1, CD4, FCGR2A, MRC1, IL21R, TNFRSF8, TLR8, CD3G, TLR3 | 3.08E-08 |
| | GOTERM_ MF_DIRECT | GO:0038023~signaling receptor activity | TLR1, CD4, IL10RA, MRC1, CD7, TLR8, CD33, TLR3 | 1.66E-05 |
| | GOTERM_ MF_DIRECT | GO:0005102~receptor binding | PTPRC, SYK, TNFSF4, STAT3, HLA-B, FN1, HLA-A, CX3CL1, HLA-E | 8.81E-05 |
| | GOTERM_ MF_DIRECT | GO:0042802~identical protein binding | CD74, FCER1G, CSF1, STAT3, FN1, CD3G, IKZF1, TLR1, CD4, CD6, CLEC7A, CASP1, BTK, TLR8, NLRP3, SLAMF7, TLR3 | 2.33E-04 |
| **Module 3** | GOTERM_ BP_DIRECT | GO:0051607~defense response to virus | IFITM3, GBP5, IFITM1, IFITM2, RSAD2, IFI6, IFIT5, IFIT1, DDX60, PARP9, IFIT3, IFI44L, IFIT2, ISG20, HERC5, PLSCR1, IFI16, GBP1, TRIM22 | 7.25E-22 |
| | GOTERM_ BP_DIRECT | GO:0009615~response to virus | IFITM3, IFITM1, CCL11, IFITM2, RSAD2, IFI44, IFIT1, DDX60, IFIT3, IFIT2, ISG20, CCL8, TRIM22 | 1.24E-16 |
| | GOTERM_ BP_DIRECT | GO:0045087~innate immune response | C1QA, RSAD2, SAMD9, NCF2, IFI6, IFIT5, LY86, LY96, UBE2L6, DDX60, PARP9, IFIT3, IFIT2, ISG20, HERC5, HCK, IFI16, FYN, GBP1, TRIM22 | 1.86E-15 |
| | GOTERM_ BP_DIRECT | GO:0006955~immune response | IFITM3, CIITA, HLA-DRB5, CCL11, IFITM2, C5AR1, IFI6, IFI44, VAV1, IFI44L, CD79B, CCL8, HLA-DPB1, IL18R1, TRIM22, HLA-DQB2 | 4.07E-12 |
| | GOTERM_ BP_DIRECT | GO:0006954~inflammatory response | GBP5, HCK, CIITA, CCL8, CCL11, IFI16, C5AR1, LY86, LY96, IL18R1, FUT4 | 2.37E-07 |
| | GOTERM_ CC_DIRECT | GO:0048471~perinuclear region of cytoplasm | HERC5, ITGB1, IFITM3, GBP5, PLSCR1, HMOX1, FYN, PTPN22, GBP4 | 7.14E-04 |

| | | | | |
|---|---|---|---|---|
| | GOTERM_ CC_DIRECT | GO:0005886~plasma membrane | ITGB1, IFITM3, IFITM1, IFITM2, NCF2, C5AR1, PLEK, IFIT5, IFI6, LY96, CD79B, FYN, APOE, GBP1, GBP4, CD53, MSR1, HLA-DRB5, CD300A, VAV1, HCK, KITLG, PLSCR1, HLA-DPB1, IL18R1, HLA-DQB2 | 0.001302 17 |
| | GOTERM_ CC_DIRECT | GO:0005829~cytosol | CIITA, SAMD9, NCF2, PLEK, IFIT5, PTPN22, UBE2L6, IFIT1, DDX60, IFIT3, IFIT2, CD79B, HERC5, SOCS1, IFI16, HMOX1, FYN, GBP1, GBP4, TRIM22, MSR1, PARP9, VAV1, HCK, PLSCR1, MNDA | 0.002083 735 |
| | GOTERM_ CC_DIRECT | GO:0005737~cytoplasm | ITGB1, SAMD9L, SAMD9, PLEK, PTPN22, UBE2L6, IFIT1, DDX60, IFI44L, IFIT3, IFIT2, HERC5, SOCS1, FYN, APOE, GBP1, TRIM22, GBP5, IFI44, PARP9, VAV1, ISG20, HCK, KITLG, PLSCR1, CMPK2 | 0.003216 603 |
| | GOTERM_ CC_DIRECT | GO:0005794~Golgi apparatus | GBP5, HCK, PLSCR1, RSAD2, APOE, GBP1, GBP4, TRIM22, FUT4 | 0.009427 568 |
| | GOTERM_ MF_DIRECT | GO:0005515~protein binding | ITGB1, IFITM3, C1QA, IFITM1, SAMD9L, CIITA, CCL11, SAMD9, NCF2, PLEK, IFIT5, IFI6, LY96, PTPN22, UBE2L6, IFIT1, DDX60, IFIT3, IFIT2, CD79B, HERC5, SOCS1, CCL8, IFI16, MYC, HMOX1, FYN, APOE, GBP1, GBP4, TRIM22, CD53, MSR1, GBP5, RSAD2, CD300A, LY86, IFI44, PARP9, VAV1, HCK, KITLG, PLSCR1, HLA-DPB1, MNDA, IL18R1 | 0.002396 4 |
| | GOTERM_ MF_DIRECT | GO:0019899~enzyme binding | PLSCR1, HMOX1, FYN, APOE, PARP9, GBP1 | 0.004469 11 |
| | GOTERM_ MF_DIRECT | GO:0042803~protein homodimerization activity | GBP5, PLEK, HMOX1, APOE, GBP1, GBP4, TRIM22 | 0.017598 996 |
| | GOTERM_ MF_DIRECT | GO:0042802~identical protein binding | CD53, CD79B, GBP5, IFI16, HMOX1, FYN, APOE, GBP1, | 0.020527 197 |

| | | | | |
|---|---|---|---|---|
| | | | GBP4, TRIM22, IFIT3 | |
| | GOTERM_ MF_DIRECT | GO:0005525~GTP binding | GBP5, CIITA, GBP1, GBP4, IFI44L | 0.026762 004 |
| Enriched GO terms of upregulated DEGs of module in fusion-negative BLCA samples | | | | |
| **Modul e1** | GOTERM_BP_DI RECT | GO:0045087~innate immune response | SAMD9, IFIT5, IFI6, UBE2L6, IFI35, DDX60, IFIT3, IFIT2, OASL, IFIH1, HERC5, IFI16, DHX58, GBP1, TRIM21, RSAD2, MX1, EIF2AK2, PARP9, PARP14, ISG20, BST2, IFI27, OAS2, OAS3, IRF7 | 3.20E-25 |
| | GOTERM_BP_DI RECT | GO:0060337~type I interferon signaling pathway | IFIH1, IFITM3, IFITM1, IFI27, STAT1, OAS2, STAT2, IRF1, IRF7, IRF9 | 8.66E-15 |
| | GOTERM_BP_DI RECT | GO:0032728~positive regulation of interferon-beta production | IFIH1, OAS2, IRF1, OAS3, DHX58, IRF7 | 5.93E-08 |
| | GOTERM_BP_DI RECT | GO:0045944~positive regulation of transcription from RNA polymerase II promoter | HELZ2, CXCL10, PLSCR1, IFI16, STAT1, STAT2, IRF1, IRF7, IRF9 | 0.008709 067 |
| | GOTERM_BP_DI RECT | GO:0006915~apoptotic process | PLSCR1, IFI27, IRF1, MX1, IFI6, XAF1 | 0.016448 213 |
| | GOTERM_CC_DI RECT | GO:0005737~cytoplasm | RTP4, SAMD9L, SAMD9, UBE2L6, IFI35, IFIT1, DDX60, USP18, IFI44L, IFIT3, IFIT2, OASL, HELZ2, IFIH1, HERC5, DHX58, GBP1, TRIM21, HERC6, GBP5, STAT1, STAT2, MX1, IFI44, EIF2AK2, PARP9, PARP14, ISG20, BST2, PLSCR1, OAS2, IRF1, OAS3, IRF7, CMPK2, XAF1, IRF9 | 1.94E-11 |
| | GOTERM_CC_DI RECT | GO:0005829~cytosol | SAMD9, IFIT5, UBE2L6, IFI35, IFIT1, DDX60, USP18, IFIT3, IFIT2, OASL, HELZ2, IFIH1, HERC5, IFI16, GBP1, TRIM21, GBP4, HERC6, STAT1, STAT2, MX1, EIF2AK2, PARP9, PARP14, BST2, PLSCR1, OAS2, IRF1, OAS3, IRF7, XAF1, IRF9 | 7.10E-08 |
| | GOTERM_CC_DI RECT | GO:0005654~nucleoplasm | SP110, STAT1, STAT2, UBE2L6, IFI35, PARP9, OASL, HELZ2, ISG20, PLSCR1, IFI16, OAS2, IRF1, | 0.001200 121 |

| | | | | |
|---|---|---|---|---|
| | | | OAS3, IRF7, CMPK2, XAF1, TRIM21, IRF9, HERC6 | |
| | GOTERM_CC_DIRECT | GO:0005739~mitochondrion | IFIH1, SAMD9L, IFI27, RSAD2, IFI6, CMPK2, PARP9, XAF1, IFIT3 | 0.018779297 |
| | GOTERM_CC_DIRECT | GO:0016020~membrane | RTP4, GBP5, IFITM1, MX1, EIF2AK2, IFI35, PARP14, PARP9, OASL, HELZ2, BST2, PLSCR1, IFI16, OAS2, OAS3 | 0.037462301 |
| | GOTERM_MF_DIRECT | GO:0003723~RNA binding | IFIT5, EIF2AK2, IFIT1, DDX60, IFIT3, PARP12, IFIT2, OASL, HELZ2, IFIH1, BST2, HERC5, IFI16, TRIM21 | 5.56E-05 |
| | GOTERM_MF_DIRECT | GO:0042802~identical protein binding | GBP5, STAT1, STAT2, MX1, EIF2AK2, IFI35, IFIT3, IFIH1, BST2, IFI16, IFI27, GBP1, TRIM21, GBP4 | 2.75E-04 |
| | GOTERM_MF_DIRECT | GO:0005515~protein binding | IFITM3, RTP4, IFITM1, SAMD9L, SAMD9, IFIT5, IFI6, UBE2L6, IFI35, IFIT1, DDX60, USP18, IFIT3, IFIT2, OASL, HELZ2, IFIH1, HERC5, IFI16, DHX58, GBP1, TRIM21, GBP4, GBP5, RSAD2, STAT1, STAT2, MX1, IFI44, EIF2AK2, PARP9, PARP14, BST2, CXCL10, PLSCR1, IFI27, OAS2, IRF1, OAS3, IRF7, XAF1, IRF9, LY6E | 9.20E-04 |
| | GOTERM_MF_DIRECT | GO:0003677~DNA binding | HELZ2, IFIH1, PLSCR1, STAT1, SP110, STAT2, IRF1, DHX58, IRF7, TRIM21, OASL | 0.001710644 |
| | GOTERM_MF_DIRECT | GO:0005524~ATP binding | HELZ2, IFIH1, OAS2, OAS3, DHX58, CMPK2, EIF2AK2, UBE2L6, DDX60 | 0.038001426 |
| **Module2** | GOTERM_BP_DIRECT | GO:0043161~proteasome-mediated ubiquitin-dependent protein catabolic process | PSMD11, PSMD14, PSMD13, ADRM1, PSMA7, PSMA5, PSMB6, PSMA6, PSMB7, PSMA3, PSMB4, PSMA4, PSMB5, PSMC3, PSMD4, PSMA1, PSMB2, PSMD2, PSMD3, PSMB1 | 6.63E-23 |
| | GOTERM_BP_DIRECT | GO:0033209~tumor necrosis factor-mediated signaling pathway | TNFRSF18, TRAF2, PSMB10, PSMA7, PSMA5, PSMB6, PSMA6, PSMB7, PSMA3, PSMB4, PSMA4, PSMB5, | 1.12E-23 |

| | | | | |
|---|---|---|---|---|
| | | | PSMA1, PSMB2, PSMB1, TNFRSF4 | |
| | GOTERM_BP_DIRECT | GO:0016579~protein deubiquitination | PSMD14, PSMB10, UCHL5, PSMA7, PSMA5, PSMB6, PSMA6, PSMB7, PSMA3, PSMB4, PSMA4, PSMB5, PSMA1, PSMB2, PSMB1 | 1.74E-18 |
| | GOTERM_BP_DIRECT | GO:0006511~ubiquitin-dependent protein catabolic process | PSMD11, PSMD14, PSMD13, ADRM1, NEDD8, UCHL5, PSMA7, PSMD9, PSMA6, PSMA3, PSMA4, PSMA1, PSMD3, ELOC | 2.99E-12 |
| | GOTERM_BP_DIRECT | GO:0043687~post-translational protein modification | PSMB10, PSMA7, PSMA5, PSMB6, PSMA6, PSMB7, PSMA3, PSMB4, PSMA4, PSMB5, PSMA1, PSMB2, PSMB1, CDK2 | 5.79E-20 |
| | GOTERM_CC_DIRECT | GO:0005654~nucleoplasm | CD274, PSMD11, PSMD14, PSMD13, NEDD8, SAMHD1, PSMB10, PSMA7, CKS1B, PSMB6, PSMD9, PSMB7, PSMB4, PSMB5, PSMD4, PSMB2, PSMD2, PSMD3, PSMB1, ELOC, FBXO5, IL15, ADRM1, TRAF2, UCHL5, PSMA5, PSMA6, PSMA3, PSMA4, PSMC3, CCNE2, PSMA1, CDK2, PSME1, PSME2, NFKBIE, TP73 | 4.20E-12 |
| | GOTERM_CC_DIRECT | GO:0005829~cytosol | PSMD11, PSMD14, PSMD13, POMP, PRF1, NEDD8, PSMB10, PSMA7, PSMB6, PSMD9, PSMB7, PSMB4, PSMB5, PSMD4, PSMB2, PSMD2, PSMD3, PSMB1, CASP1, ELOC, FBXO5, IL15, ADRM1, TRAF2, UCHL5, PSMA5, PSMA6, PSMA3, PSMA4, PSMC3, CCNE2, PSMA1, CDK2, PSME1, PSME2, NFKBIE, GAPDH, CD44, IDO1, TP73 | 5.26E-10 |
| | GOTERM_CC_DIRECT | GO:0070062~extracellular exosome | CD274, NEDD8, FASLG, MMP9, PSMA7, ICAM1, PSMA5, PSMB6, PSMA6, PSMA3, PSMB4, PSMA4, | 8.88E-08 |

| | | | | |
|---|---|---|---|---|
| | | | PSMB5, PSMA1, PSMB2, PSMD2, PSMD3, PSMB1, PSME1, CD38, PSME2, GAPDH, CD44 | |
| | GOTERM_CC_DIRECT | GO:0005634~nucleus | PSMD11, PSMD14, PSMD13, POMP, NEDD8, FASLG, SAMHD1, PSMB10, PSMA7, PSMB6, PSMD9, PSMB7, PSMB4, PSMB5, PSMD4, PSMB2, PSMD2, PSMD3, PSMB1, CD38, FBXO5, BATF2, ADRM1, UCHL5, PSMA5, PSMA6, PSMA3, PSMA4, PSMC3, CCNE2, PSMA1, CDK2, GAPDH, TP73 | 2.10E-05 |
| | GOTERM_CC_DIRECT | GO:0005737~cytoplasm | POMP, NEDD8, PSMA7, PSMB6, PSMD9, PSMB7, PSMB4, PSMB5, PSMB2, PSMB1, CASP1, FBXO5, CCR1, IL15, ADRM1, UCHL5, PSMA5, PSMA6, PSMA3, PSMA4, PSMC3, CCNE2, PSMA1, CDK2, PSME1, PSME2, NFKBIE, GAPDH, IDO1, TP73 | 4.69E-04 |
| | GOTERM_MF_DIRECT | GO:0004298~threonine-type endopeptidase activity | PSMB6, PSMB7, PSMB4, PSMA4, PSMB5, PSMB2, PSMB1, PSMB10 | 6.54E-15 |
| | GOTERM_MF_DIRECT | GO:0004175~endopeptidase activity | PSMB6, PSMA6, PSMB7, PSMB5, CASP1, MMP9, PSMB10 | 1.62E-07 |
| | GOTERM_MF_DIRECT | GO:0005515~protein binding | CXCL9, PRF1, FASLG, CXCL1, CKS1B, ICAM1, PSMD9, PSMD4, PSMD2, PSMD3, CASP1, ELOC, FBXO5, TNFRSF4, BATF2, LAG3, IL15, TNFRSF18, TRAF2, PDCD1LG2, MMP9, UCHL5, PSMA5, PSMA6, PSMA3, PSMA4, CCNE2, PSMA1, PSME1, PSME2, GAPDH, CD44, CD274, PSMD11, PSMD14, POMP, NEDD8, SAMHD1, PSMA7, PSMB10, PSMB6, PSMB7, PSMB4, PSMB5, PSMB2, PSMB1, | 1.34E-06 |

| | | | | |
|---|---|---|---|---|
| | | | KLRC1, CCR1, ADRM1, CXCL11, FCGR2A, PSMC3, IL2RA, CDK2, NFKBIE, TP73 | |
| | GOTERM_MF_DIRECT | GO:0004888~transmembrane signaling receptor activity | LAG3, FCGR2A, KLRC1, CD44, ICAM1 | 0.00352907 |
| | GOTERM_MF_DIRECT | GO:0042802~identical protein binding | PSMC3, PSMD4, CASP1, PRF1, CD38, PSME2, TRAF2, SAMHD1, GAPDH, MMP9, PSMA7, TP73 | 0.016919787 |
| **Module3** | GOTERM_BP_DIRECT | GO:0006955~immune response | CD86, CD74, CD70, CD80, GZMA, HLA-B, HLA-C, HLA-A, IL2RG, HLA-G, CXCL5, FCGR3A, AIM2, IFNG, CCL3, HLA-DRA, CTLA4, TLR6, FCGR1A, B2M, HLA-DQB1, TLR2 | 4.95E-21 |
| | GOTERM_BP_DIRECT | GO:0045087~innate immune response | C1QA, FCER1G, HLA-B, NLRC5, HLA-C, CGAS, HLA-A, NOD2, TREM1, AIM2, IRAK1, TLR8, TLR6, FCGR1A, B2M, HAVCR2, TLR2 | 1.17E-12 |
| | GOTERM_BP_DIRECT | GO:0002250~adaptive immune response | CD86, HLA-B, HLA-C, TAP1, LILRB1, HLA-A, NOD2, TREM1, IFNG, HLA-DRA, CTLA4, KLRD1, PDCD1, HAVCR2, HLA-DQB1 | 2.96E-12 |
| | GOTERM_BP_DIRECT | GO:0006954~inflammatory response | AIM2, CLEC7A, ITGB2, CCL3, C3AR1, XCL1, TLR8, TLR6, AIF1, CXCL5, HAVCR2, TLR2 | 6.74E-09 |
| | GOTERM_BP_DIRECT | GO:0007165~signal transduction | TYROBP, CD70, IL2RB, XCL1, LILRB1, TLR6, FCGR1A, IL2RG, IL12RB1, EGFR, CXCL5, TLR2 | 2.75E-04 |
| | GOTERM_CC_DIRECT | GO:0009897~external side of plasma membrane | CD86, CD74, FCER1G, CD80, TNFRSF9, ITGB2, HLA-B, HLA-C, LILRB1, HLA-A, IL2RG, HLA-G, FCGR3A, CLEC7A, IL2RB, ITGAX, IL21R, TLR8, CTLA4, KLRD1, PDCD1, FCGR1A, IL12RB1, B2M | 3.34E-25 |
| | GOTERM_CC_DIRECT | GO:0009986~cell surface | CD86, CD74, IL15RA, FCER1G, CD80, ITGB2, HLA-B, HLA-C, HLA-A, NOD2, IL2RG, TREM1, EGFR, TYROBP, IRAK1, CLEC7A, IL2RB, ITGAX, HLA-DRA, | 2.57E-19 |

| | | | | |
|---|---|---|---|---|
| | | | HAVCR2, HLA-DQB1, TLR2 | |
| | GOTERM_CC_DIRECT | GO:0005886~plasma membrane | CD86, CD80, ITGB2, CGAS, NOD2, IL2RG, TREM1, EGFR, FCGR3A, IRAK1, CLEC7A, C3AR1, ITGAX, IL21R, CTLA4, IL12RB1, FCGR1A, B2M, CD74, IL15RA, FCER1G, CD70, TNFRSF9, HLA-B, HLA-C, LILRB1, HLA-A, HLA-G, TYROBP, IL2RB, TLR8, HLA-DRA, KLRD1, PDCD1, TLR6, HLA-DQB1, TLR2 | 2.86E-12 |
| | GOTERM_CC_DIRECT | GO:0005887~integral component of plasma membrane | FCER1G, CD70, TNFRSF9, HLA-B, HLA-C, TAP1, HLA-A, IL2RG, EGFR, FCGR3A, TYROBP, IL2RB, C3AR1, HLA-DRA, CTLA4, KLRD1, TLR6, FCGR1A, TLR2 | 1.56E-09 |
| | GOTERM_CC_DIRECT | GO:0016021~integral component of membrane | CD86, CD80, IL2RG, TREM1, EGFR, FCGR3A, CLEC7A, C3AR1, IL21R, CTLA4, FCGR1A, HAVCR2, CD74, IL15RA, FCER1G, CD70, TNFRSF9, HLA-B, HLA-C, TAP1, LILRB1, HLA-A, HLA-G, TYROBP, IL2RB, TLR8, HLA-DRA, KLRD1, PDCD1, TLR6, HLA-DQB1, TLR2 | 3.75E-08 |
| | GOTERM_MF_DIRECT | GO:0004888~transmembrane signaling receptor activity | FCGR3A, IL21R, TLR8, KLRD1, TLR6, FCGR1A, TREM1, EGFR, TLR2 | 3.62E-08 |
| | GOTERM_MF_DIRECT | GO:0038023~signaling receptor activity | CD86, TNFRSF9, ITGB2, ITGAX, TLR8, TLR6, TREM1, TLR2 | 1.90E-06 |
| | GOTERM_MF_DIRECT | GO:0042803~protein homodimerization activity | TYROBP, FCER1G, IRAK1, GZMA, XCL1, TAP1, CGAS, LILRB1, B2M, HLA-G | 9.32E-05 |
| | GOTERM_MF_DIRECT | GO:0005515~protein binding | CD86, C1QA, CD80, ITGB2, NLRC5, CGAS, NOD2, IL2RG, AIF1, EGFR, CXCL5, FCGR3A, IRAK1, CLEC7A, CCL3, ITGAX, IL21R, CTLA4, FCGR1A, B2M, HAVCR2, CD74, IL15RA, FCER1G, CD70, TNFRSF9, GZMA, HLA-B, HLA-C, TAP1, GZMB, | 5.17E-05 |

| | | | | |
|---|---|---|---|---|
| | | | LILRB1, HLA-A, HLA-G, TYROBP, AIM2, IFNG, IL2RB, XCL1, HLA-DRA, KLRD1, PDCD1, TLR6, HLA-DQB1, TLR2 | |
| | GOTERM_MF_DIRECT | GO:0042802~identical protein binding | CD74, FCER1G, HLA-G, EGFR, CXCL5, AIM2, TYROBP, IRAK1, CLEC7A, CCL3, TLR8, TLR6, B2M, TLR2 | 2.75E-04 |
| **Enriched GO terms of downregulated DEGs of module in fusion-negative BLCA samples** | | | | |
| **Module 1** | GOTERM_BP_DIRECT | GO:0006805~xenobiotic metabolic process | UGT1A10, CYP2A6, UGT1A1, EPHX1, GSTA1, CYP1A1, UGT1A8, CYP3A5, UGT1A6, SULT2A1 | 3.47E-14 |
| | GOTERM_BP_DIRECT | GO:0008202~steroid metabolic process | CYP2A6, UGT1A1, CYP1A1, UGT1A8, CYP3A5, SULT2A1 | 1.38E-08 |
| | GOTERM_BP_DIRECT | GO:0042178~xenobiotic catabolic process | GSTM4, GSTM3, GSTM2, CYP2A6, CYP3A5 | 2.55E-08 |
| | GOTERM_BP_DIRECT | GO:0006631~fatty acid metabolic process | ACOX1, CYP1A1, PPARG, PPARA, UGT1A8 | 1.01E-05 |
| | GOTERM_BP_DIRECT | GO:0006629~lipid metabolic process | UGT1A10, ACOX1, CIDEA, PPARG, PLIN1 | 2.31E-04 |
| | GOTERM_CC_DIRECT | GO:0045171~intercellular bridge | GSTM4, GSTM3, GSTM2, GPX2 | 2.96E-04 |
| | GOTERM_CC_DIRECT | GO:0005811~lipid particle | LIPE, FABP4, CIDEA, PLIN1 | 3.61E-04 |
| | GOTERM_CC_DIRECT | GO:0005789~endoplasmic reticulum membrane | UGT1A10, CYP2A6, UGT1A1, EPHX1, CYP1A1, UGT1A8, CYP3A5, UGT1A6 | 4.99E-04 |
| | GOTERM_CC_DIRECT | GO:0043231~intracellular membrane-bounded organelle | CEBPA, CYP2A6, CYP1A1, PPARG, CYP3A5, PPARGC1A, UGT1A6 | 0.001685551 |
| | GOTERM_CC_DIRECT | GO:0005829~cytosol | GSTM4, GSTM3, GSTM2, GPX2, ADH1C, CIDEA, LIPE, GCLC, FABP4, ACOX1, GSTA1, PPARG, PLIN1, PPARGC1A, SULT2A1 | 0.005725493 |
| | GOTERM_MF_DIRECT | GO:0019899~enzyme binding | GSTM4, GSTM3, UGT1A10, GSTM2, CYP2A6, UGT1A1, CYP1A1, PPARG, UGT1A8, UGT1A6 | 1.93E-09 |
| | GOTERM_MF_DIRECT | GO:0005504~fatty acid binding | GSTM2, FABP4, ACOX1, GSTA1, UGT1A8 | 2.08E-07 |
| | GOTERM_MF_DIRECT | GO:0042803~protein homodimerization activity | GSTM4, GSTM3, CEBPA, UGT1A10, GSTM2, UGT1A1, ACOX1, CIDEA, UGT1A8, | 5.48E-07 |

| | | | | |
|---|---|---|---|---|
| | | | UGT1A6 | |
| | GOTERM_MF_DIRECT | GO:0015020~glucuronosyltransferase activity | UGT1A10, UGT1A1, UGT1A8, UGT1A6 | 1.02E-05 |
| | GOTERM_MF_DIRECT | GO:0046982~protein heterodimerization activity | UGT1A10, UGT1A1, UGT1A8, UGT1A6 | 0.016436854 |
| **Module2** | GOTERM_BP_DIRECT | GO:0009060~aerobic respiration | MT-ND6, MT-ND5, MT-CO1, MT-ND3, MT-ND1 | 4.73E-09 |
| | GOTERM_BP_DIRECT | GO:0006120~mitochondrial electron transport, NADH to ubiquinone | MT-ND6, MT-ND5, MT-ND3, MT-ND1 | 4.30E-07 |
| | GOTERM_BP_DIRECT | GO:0042776~mitochondrial ATP synthesis coupled proton transport | MT-ND6, MT-ND5, MT-ND3, MT-ND1 | 1.23E-06 |
| | GOTERM_BP_DIRECT | GO:0045333~cellular respiration | MT-CO1, COX4I2, MT-CYB | 9.48E-05 |
| | GOTERM_BP_DIRECT | GO:0032981~mitochondrial respiratory chain complex I assembly | MT-ND6, MT-ND5, MT-ND1 | 2.28E-04 |
| | GOTERM_CC_DIRECT | GO:0005743~mitochondrial inner membrane | MT-ND6, MT-ND5, MT-CO1, MT-CYB, MT-ND3, MT-ND1 | 4.04E-07 |
| | GOTERM_CC_DIRECT | GO:0005747~mitochondrial respiratory chain complex I | MT-ND6, MT-ND5, MT-ND3, MT-ND1 | 7.41E-07 |
| | GOTERM_CC_DIRECT | GO:0005739~mitochondrion | MT-ND6, H6PD, MT-ND5, MT-CO1, MT-CYB, MT-ND3, MT-ND1 | 3.14E-06 |
| | GOTERM_CC_DIRECT | GO:0031966~mitochondrial membrane | MT-CO1, MT-ND3, MT-ND1 | 0.001824362 |
| | GOTERM_CC_DIRECT | GO:0016021~integral component of membrane | MT-ND6, MT-ND5, MT-CO1, COX4I2, MT-CYB, MT-ND3, MT-ND1 | 0.005868953 |
| | GOTERM_MF_DIRECT | GO:0008137~NADH dehydrogenase (ubiquinone) activity | MT-ND6, MT-ND5, MT-CO1, MT-ND3, MT-ND1 | 9.61E-10 |
| | GOTERM_MF_DIRECT | GO:0004129~cytochrome-c oxidase activity | MT-CO1, COX4I2, MT-CYB | 2.21E-05 |
| | GOTERM_MF_DIRECT | GO:0003954~NADH dehydrogenase activity | MT-ND5, MT-ND1 | 0.004418952 |
| **Module3** | GOTERM_BP_DIRECT | GO:0030198~extracellular matrix organization | COL24A1, COL9A1, COL4A6, COL4A5, COL9A2, COL6A5 | 5.01E-10 |
| | GOTERM_BP_DIRECT | GO:0030199~collagen fibril organization | COL24A1, COL9A1, COL4A6, COL4A5 | 1.18E-06 |
| | GOTERM_BP_DIRECT | GO:0038063~collagen-activated tyrosine kinase receptor signaling pathway | COL4A6, COL4A5 | 0.00338687 |
| | GOTERM_CC_DIRECT | GO:0005581~collagen trimer | COL24A1, COL9A1, COL4A6, COL4A5, COL9A2, COL6A5 | 1.29E-11 |

| | | | | |
|---|---|---|---|---|
| | GOTERM_CC_DIRECT | GO:0031012~extracellular matrix | COL24A1, COL9A1, COL4A6, COL4A5, COL9A2 | 3.55E-07 |
| | GOTERM_CC_DIRECT | GO:0005788~endoplasmic reticulum lumen | COL24A1, COL9A1, COL4A6, COL4A5, COL9A2 | 7.38E-07 |
| | GOTERM_CC_DIRECT | GO:0005576~extracellular region | COL24A1, COL9A1, COL4A6, COL4A5, COL9A2, COL6A5 | 7.68E-05 |
| | GOTERM_CC_DIRECT | GO:0005615~extracellular space | COL24A1, COL9A1, COL4A6, COL4A5, COL9A2 | 0.001095384 |
| | GOTERM_MF_DIRECT | GO:0030020~extracellular matrix structural constituent conferring tensile strength | COL24A1, COL9A1, COL4A6, COL4A5, COL9A2, COL6A5 | 3.57E-13 |
| | GOTERM_MF_DIRECT | GO:0005201~extracellular matrix structural constituent | COL24A1, COL9A1, COL4A6, COL4A5, COL9A2 | 4.09E-08 |

### Supplementary Table S4. GSEA Result of fusion-positive vs fusion-negative cohorts.

| GENESET | NES | FDR |
|---|---|---|
| gene_sets.gmt#KEGG_METABOLISM_OF_XENOBIOTICS_BY_CYTOCHROME_P450 | 1.5777 | 0.9826 |
| gene_sets.gmt#KEGG_PEROXISOME | 1.5141 | 0.8387 |
| gene_sets.gmt#KEGG_GLUTATHIONE_METABOLISM | 1.4997 | |
| gene_sets.gmt#KEGG_OXIDATIVE_PHOSPHORYLATION | 1.4944 | 0.4822 |
| gene_sets.gmt#KEGG_SPLICEOSOME | 1.4924 | 0.3916 |
| gene_sets.gmt#KEGG_GLYCOSYLPHOSPHATIDYLINOSITOL_GPI_ANCHOR_BIOSYNTHESIS | 1.4869 | 0.3403 |
| gene_sets.gmt#KEGG_PHENYLALANINE_METABOLISM | 1.4354 | 0.4131 |
| gene_sets.gmt#KEGG_DRUG_METABOLISM_CYTOCHROME_P450 | 1.4221 | 0.3924 |
| gene_sets.gmt#KEGG_PARKINSONS_DISEASE | 1.3892 | 0.4306 |
| gene_sets.gmt#KEGG_HUNTINGTONS_DISEASE | 1.3887 | 0.3881 |
| gene_sets.gmt#KEGG_GLYCEROPHOSPHOLIPID_METABOLISM | 1.3546 | 0.43 |
| gene_sets.gmt#KEGG_PENTOSE_AND_GLUCURONATE_INTERCONVERSIONS | 1.3342 | 0.4384 |
| gene_sets.gmt#KEGG_STEROID_HORMONE_BIOSYNTHESIS | 1.3316 | 0.4102 |
| gene_sets.gmt#KEGG_LINOLEIC_ACID_METABOLISM | 1.3194 | 0.407 |
| gene_sets.gmt#KEGG_NOTCH_SIGNALING_PATHWAY | 1.3074 | 0.4077 |
| gene_sets.gmt#KEGG_TYROSINE_METABOLISM | 1.3012 | 0.3942 |
| gene_sets.gmt#KEGG_ETHER_LIPID_METABOLISM | 1.2986 | 0.3753 |
| gene_sets.gmt#KEGG_RNA_POLYMERASE | 1.261 | 0.4284 |
| gene_sets.gmt#KEGG_RETINOL_METABOLISM | 1.2563 | 0.4159 |
| gene_sets.gmt#KEGG_RNA_DEGRADATION | 1.215 | 0.4723 |
| gene_sets.gmt#KEGG_ALANINE_ASPARTATE_AND_GLUTAMATE_METABOLISM | 1.2095 | 0.4609 |
| gene_sets.gmt#KEGG_RIBOSOME | 1.2086 | 0.4419 |
| gene_sets.gmt#KEGG_ALPHA_LINOLENIC_ACID_METABOLISM | 1.2078 | 0.424 |
| gene_sets.gmt#KEGG_FRUCTOSE_AND_MANNOSE_METABOLISM | 1.1804 | 0.4546 |
| gene_sets.gmt#KEGG_ALZHEIMERS_DISEASE | 1.1523 | 0.4899 |
| gene_sets.gmt#KEGG_BASE_EXCISION_REPAIR | 1.1356 | 0.5008 |
| gene_sets.gmt#KEGG_VALINE_LEUCINE_AND_ISOLEUCINE_DEGRADATION | 1.1354 | 0.4825 |
| gene_sets.gmt#KEGG_AMINOACYL_TRNA_BIOSYNTHESIS | 1.1296 | 0.477 |
| gene_sets.gmt#KEGG_ARACHIDONIC_ACID_METABOLISM | 1.1219 | 0.474 |
| gene_sets.gmt#KEGG_UBIQUITIN_MEDIATED_PROTEOLYSIS | 1.1135 | 0.4729 |
| gene_sets.gmt#KEGG_PORPHYRIN_AND_CHLOROPHYLL_METABOLISM | 1.0642 | 0.5465 |
| gene_sets.gmt#KEGG_ASCORBATE_AND_ALDARATE_METABOLISM | 1.053 | 0.551 |
| gene_sets.gmt#KEGG_BUTANOATE_METABOLISM | 1.0428 | 0.5525 |
| gene_sets.gmt#KEGG_STEROID_BIOSYNTHESIS | 0.9876 | 0.6399 |
| gene_sets.gmt#KEGG_DRUG_METABOLISM_OTHER_ENZYMES | 0.9872 | 0.6225 |
| gene_sets.gmt#KEGG_PENTOSE_PHOSPHATE_PATHWAY | 0.9542 | 0.6677 |
| gene_sets.gmt#KEGG_HISTIDINE_METABOLISM | 0.9388 | 0.6793 |

| | | |
|---|---|---|
| gene_sets.gmt#KEGG_STARCH_AND_SUCROSE_METABOLISM | 0.9364 | 0.666 |
| gene_sets.gmt#KEGG_HOMOLOGOUS_RECOMBINATION | 0.9276 | 0.6661 |
| gene_sets.gmt#KEGG_OTHER_GLYCAN_DEGRADATION | 0.9032 | 0.6938 |
| gene_sets.gmt#KEGG_GLYCOLYSIS_GLUCONEOGENESIS | 0.8756 | 0.7264 |
| gene_sets.gmt#KEGG_SELENOAMINO_ACID_METABOLISM | 0.8667 | 0.7251 |
| gene_sets.gmt#KEGG_N_GLYCAN_BIOSYNTHESIS | 0.8526 | 0.734 |
| gene_sets.gmt#KEGG_BASAL_TRANSCRIPTION_FACTORS | 0.8225 | 0.7688 |
| gene_sets.gmt#KEGG_TERPENOID_BACKBONE_BIOSYNTHESIS | 0.8198 | 0.7566 |
| gene_sets.gmt#KEGG_GALACTOSE_METABOLISM | 0.7922 | 0.7852 |
| gene_sets.gmt#KEGG_MISMATCH_REPAIR | 0.7574 | 0.8217 |
| gene_sets.gmt#KEGG_PROTEIN_EXPORT | 0.7034 | 0.8784 |
| gene_sets.gmt#KEGG_NUCLEOTIDE_EXCISION_REPAIR | 0.6299 | 0.9305 |
| gene_sets.gmt#KEGG_PROTEASOME | -0.4072 | 0.9978 |
| gene_sets.gmt#KEGG_DNA_REPLICATION | -0.7521 | 0.7987 |
| gene_sets.gmt#KEGG_ONE_CARBON_POOL_BY_FOLATE | -0.7576 | 0.7971 |
| gene_sets.gmt#KEGG_CITRATE_CYCLE_TCA_CYCLE | -0.7704 | 0.7854 |
| gene_sets.gmt#KEGG_CELL_CYCLE | -0.8192 | 0.7175 |
| gene_sets.gmt#KEGG_MATURITY_ONSET_DIABETES_OF_THE_YOUNG | -0.8464 | 0.6798 |
| gene_sets.gmt#KEGG_NITROGEN_METABOLISM | -0.8548 | 0.6717 |
| gene_sets.gmt#KEGG_AMINO_SUGAR_AND_NUCLEOTIDE_SUGAR_METABOLISM | -0.8591 | 0.6705 |
| gene_sets.gmt#KEGG_OLFACTORY_TRANSDUCTION | -0.8868 | 0.6328 |
| gene_sets.gmt#KEGG_P53_SIGNALING_PATHWAY | -0.9095 | 0.6016 |
| gene_sets.gmt#KEGG_PYRIMIDINE_METABOLISM | -0.9273 | 0.5791 |
| gene_sets.gmt#KEGG_PROPANOATE_METABOLISM | -0.9373 | 0.5688 |
| gene_sets.gmt#KEGG_CARDIAC_MUSCLE_CONTRACTION | -0.9502 | 0.5549 |
| gene_sets.gmt#KEGG_REGULATION_OF_AUTOPHAGY | -0.9638 | 0.5394 |
| gene_sets.gmt#KEGG_FATTY_ACID_METABOLISM | -0.972 | 0.532 |
| gene_sets.gmt#KEGG_PROXIMAL_TUBULE_BICARBONATE_RECLAMATION | -0.9928 | 0.5041 |
| gene_sets.gmt#KEGG_PPAR_SIGNALING_PATHWAY | -1.0094 | 0.484 |
| gene_sets.gmt#KEGG_TASTE_TRANSDUCTION | -1.016 | 0.4785 |
| gene_sets.gmt#KEGG_BIOSYNTHESIS_OF_UNSATURATED_FATTY_ACIDS | -1.0321 | 0.4594 |
| gene_sets.gmt#KEGG_VASOPRESSIN_REGULATED_WATER_REABSORPTION | -1.0645 | 0.4181 |
| gene_sets.gmt#KEGG_GLYCEROLIPID_METABOLISM | -1.0779 | 0.4041 |
| gene_sets.gmt#KEGG_GLYCINE_SERINE_AND_THREONINE_METABOLISM | -1.0829 | 0.4007 |
| gene_sets.gmt#KEGG_LYSINE_DEGRADATION | -1.1057 | 0.3739 |
| gene_sets.gmt#KEGG_ARGININE_AND_PROLINE_METABOLISM | -1.1218 | 0.3556 |
| gene_sets.gmt#KEGG_CYSTEINE_AND_METHIONINE_METABOLISM | -1.1258 | 0.3536 |
| gene_sets.gmt#KEGG_BETA_ALANINE_METABOLISM | -1.1394 | 0.3392 |
| gene_sets.gmt#KEGG_PYRUVATE_METABOLISM | -1.1491 | 0.3294 |
| gene_sets.gmt#KEGG_ERBB_SIGNALING_PATHWAY | -1.1539 | 0.3264 |
| gene_sets.gmt#KEGG_VIBRIO_CHOLERAE_INFECTION | -1.2025 | 0.2718 |

| | | |
|---|---|---|
| gene_sets.gmt#KEGG_AMYOTROPHIC_LATERAL_SCLEROSIS_ALS | -1.2063 | 0.2702 |
| gene_sets.gmt#KEGG_GLYCOSAMINOGLYCAN_BIOSYNTHESIS_HEPARAN_SULFATE | -1.2136 | 0.2646 |
| gene_sets.gmt#KEGG_INOSITOL_PHOSPHATE_METABOLISM | -1.2177 | 0.2623 |
| gene_sets.gmt#KEGG_ABC_TRANSPORTERS | -1.223 | 0.2589 |
| gene_sets.gmt#KEGG_GLYOXYLATE_AND_DICARBOXYLATE_METABOLISM | -1.2293 | 0.2549 |
| gene_sets.gmt#KEGG_TRYPTOPHAN_METABOLISM | -1.2345 | 0.252 |
| gene_sets.gmt#KEGG_INSULIN_SIGNALING_PATHWAY | -1.2369 | 0.2517 |
| gene_sets.gmt#KEGG_THYROID_CANCER | -1.2393 | 0.2516 |
| gene_sets.gmt#KEGG_MTOR_SIGNALING_PATHWAY | -1.2521 | 0.2403 |
| gene_sets.gmt#KEGG_BLADDER_CANCER | -1.2642 | 0.2297 |
| gene_sets.gmt#KEGG_NICOTINATE_AND_NICOTINAMIDE_METABOLISM | -1.2728 | 0.2239 |
| gene_sets.gmt#KEGG_LYSOSOME | -1.2842 | 0.2153 |
| gene_sets.gmt#KEGG_ENDOCYTOSIS | -1.3141 | 0.1894 |
| gene_sets.gmt#KEGG_ALDOSTERONE_REGULATED_SODIUM_REABSORPTION | -1.3328 | 0.1739 |
| gene_sets.gmt#KEGG_RENAL_CELL_CARCINOMA | -1.3365 | 0.1726 |
| gene_sets.gmt#KEGG_TIGHT_JUNCTION | -1.3373 | 0.174 |
| gene_sets.gmt#KEGG_EPITHELIAL_CELL_SIGNALING_IN_HELICOBACTER_PYLORI_INFECTION | -1.3418 | 0.1721 |
| gene_sets.gmt#KEGG_ADHERENS_JUNCTION | -1.3436 | 0.1723 |
| gene_sets.gmt#KEGG_OOCYTE_MEIOSIS | -1.3805 | 0.1418 |
| gene_sets.gmt#KEGG_PHOSPHATIDYLINOSITOL_SIGNALING_SYSTEM | -1.3814 | 0.1429 |
| gene_sets.gmt#KEGG_TYPE_II_DIABETES_MELLITUS | -1.3861 | 0.1409 |
| gene_sets.gmt#KEGG_VEGF_SIGNALING_PATHWAY | -1.3991 | 0.1317 |
| gene_sets.gmt#KEGG_SNARE_INTERACTIONS_IN_VESICULAR_TRANSPORT | -1.4123 | 0.1232 |
| gene_sets.gmt#KEGG_RIG_I_LIKE_RECEPTOR_SIGNALING_PATHWAY | -1.4161 | 0.1218 |
| gene_sets.gmt#KEGG_CYTOSOLIC_DNA_SENSING_PATHWAY | -1.4205 | 0.1199 |
| gene_sets.gmt#KEGG_TGF_BETA_SIGNALING_PATHWAY | -1.4267 | 0.117 |
| gene_sets.gmt#KEGG_GLYCOSPHINGOLIPID_BIOSYNTHESIS_LACTO_AND_NEOLACTO_SERIES | -1.4357 | 0.1119 |
| gene_sets.gmt#KEGG_NON_SMALL_CELL_LUNG_CANCER | -1.4369 | 0.1127 |
| gene_sets.gmt#KEGG_PANTOTHENATE_AND_COA_BIOSYNTHESIS | -1.4375 | 0.1136 |
| gene_sets.gmt#KEGG_HEDGEHOG_SIGNALING_PATHWAY | -1.4388 | 0.1144 |
| gene_sets.gmt#KEGG_GLYCOSAMINOGLYCAN_DEGRADATION | -1.4402 | 0.115 |
| gene_sets.gmt#KEGG_ENDOMETRIAL_CANCER | -1.4556 | 0.1049 |
| gene_sets.gmt#KEGG_CHRONIC_MYELOID_LEUKEMIA | -1.4593 | 0.1036 |
| gene_sets.gmt#KEGG_SPHINGOLIPID_METABOLISM | -1.4636 | 0.1022 |
| gene_sets.gmt#KEGG_RENIN_ANGIOTENSIN_SYSTEM | -1.4743 | 0.0962 |
| gene_sets.gmt#KEGG_GLYCOSAMINOGLYCAN_BIOSYNTHESIS_KERATAN_SULFATE | -1.4841 | 0.0915 |
| gene_sets.gmt#KEGG_APOPTOSIS | -1.4931 | 0.0869 |
| gene_sets.gmt#KEGG_PANCREATIC_CANCER | -1.5261 | 0.0683 |
| gene_sets.gmt#KEGG_PURINE_METABOLISM | -1.5294 | 0.0677 |
| gene_sets.gmt#KEGG_LONG_TERM_POTENTIATION | -1.5371 | 0.0648 |
| gene_sets.gmt#KEGG_DORSO_VENTRAL_AXIS_FORMATION | -1.5434 | 0.0627 |

| | | |
|---|---|---|
| gene_sets.gmt#KEGG_GLYCOSPHINGOLIPID_BIOSYNTHESIS_GANGLIO_SERIES | -1.5664 | 0.0523 |
| gene_sets.gmt#KEGG_ADIPOCYTOKINE_SIGNALING_PATHWAY | -1.5714 | 0.0508 |
| gene_sets.gmt#KEGG_GNRH_SIGNALING_PATHWAY | -1.5726 | 0.0511 |
| gene_sets.gmt#KEGG_FC_EPSILON_RI_SIGNALING_PATHWAY | -1.5751 | 0.0508 |
| gene_sets.gmt#KEGG_PRIMARY_IMMUNODEFICIENCY | -1.5811 | 0.0487 |
| gene_sets.gmt#KEGG_NEUROTROPHIN_SIGNALING_PATHWAY | -1.5819 | 0.0493 |
| gene_sets.gmt#KEGG_B_CELL_RECEPTOR_SIGNALING_PATHWAY | -1.5823 | 0.0501 |
| gene_sets.gmt#KEGG_RIBOFLAVIN_METABOLISM | -1.5942 | 0.0462 |
| gene_sets.gmt#KEGG_BASAL_CELL_CARCINOMA | -1.5957 | 0.0466 |
| gene_sets.gmt#KEGG_AXON_GUIDANCE | -1.597 | 0.0471 |
| gene_sets.gmt#KEGG_NATURAL_KILLER_CELL_MEDIATED_CYTOTOXICITY | -1.5982 | 0.0477 |
| gene_sets.gmt#KEGG_GLIOMA | -1.6098 | 0.0432 |
| gene_sets.gmt#KEGG_ANTIGEN_PROCESSING_AND_PRESENTATION | -1.6113 | 0.0434 |
| gene_sets.gmt#KEGG_PROGESTERONE_MEDIATED_OOCYTE_MATURATION | -1.6117 | 0.0442 |
| gene_sets.gmt#KEGG_SMALL_CELL_LUNG_CANCER | -1.6202 | 0.0412 |
| gene_sets.gmt#KEGG_WNT_SIGNALING_PATHWAY | -1.6217 | 0.0416 |
| gene_sets.gmt#KEGG_PATHOGENIC_ESCHERICHIA_COLI_INFECTION | -1.63 | 0.0394 |
| gene_sets.gmt#KEGG_GRAFT_VERSUS_HOST_DISEASE | -1.6306 | 0.0401 |
| gene_sets.gmt#KEGG_COLORECTAL_CANCER | -1.6374 | 0.0383 |
| gene_sets.gmt#KEGG_SYSTEMIC_LUPUS_ERYTHEMATOSUS | -1.6419 | 0.0372 |
| gene_sets.gmt#KEGG_PROSTATE_CANCER | -1.6426 | 0.038 |
| gene_sets.gmt#KEGG_TYPE_I_DIABETES_MELLITUS | -1.6521 | 0.0356 |
| gene_sets.gmt#KEGG_LONG_TERM_DEPRESSION | -1.6747 | 0.0277 |
| gene_sets.gmt#KEGG_ALLOGRAFT_REJECTION | -1.6865 | 0.0242 |
| gene_sets.gmt#KEGG_T_CELL_RECEPTOR_SIGNALING_PATHWAY | -1.6986 | 0.0217 |
| gene_sets.gmt#KEGG_GLYCOSAMINOGLYCAN_BIOSYNTHESIS_CHONDROITIN_SULFATE | -1.7042 | 0.0208 |
| gene_sets.gmt#KEGG_PRIMARY_BILE_ACID_BIOSYNTHESIS | -1.709 | 0.0201 |
| gene_sets.gmt#KEGG_FC_GAMMA_R_MEDIATED_PHAGOCYTOSIS | -1.7112 | 0.0202 |
| gene_sets.gmt#KEGG_PATHWAYS_IN_CANCER | -1.7152 | 0.0197 |
| gene_sets.gmt#KEGG_INTESTINAL_IMMUNE_NETWORK_FOR_IGA_PRODUCTION | -1.7232 | 0.0187 |
| gene_sets.gmt#KEGG_NOD_LIKE_RECEPTOR_SIGNALING_PATHWAY | -1.7248 | 0.0191 |
| gene_sets.gmt#KEGG_O_GLYCAN_BIOSYNTHESIS | -1.7293 | 0.0185 |
| gene_sets.gmt#KEGG_VASCULAR_SMOOTH_MUSCLE_CONTRACTION | -1.7357 | 0.0174 |
| gene_sets.gmt#KEGG_TOLL_LIKE_RECEPTOR_SIGNALING_PATHWAY | -1.7378 | 0.0176 |
| gene_sets.gmt#KEGG_ARRHYTHMOGENIC_RIGHT_VENTRICULAR_CARDIOMYOPATHY_ARVC | -1.7447 | 0.0167 |
| gene_sets.gmt#KEGG_ASTHMA | -1.7486 | 0.0165 |
| gene_sets.gmt#KEGG_COMPLEMENT_AND_COAGULATION_CASCADES | -1.7493 | 0.0171 |
| gene_sets.gmt#KEGG_MELANOMA | -1.7523 | 0.0171 |
| gene_sets.gmt#KEGG_CALCIUM_SIGNALING_PATHWAY | -1.761 | 0.0157 |
| gene_sets.gmt#KEGG_ACUTE_MYELOID_LEUKEMIA | -1.7676 | 0.0147 |
| gene_sets.gmt#KEGG_REGULATION_OF_ACTIN_CYTOSKELETON | -1.784 | 0.0122 |

| | | |
|---|---|---|
| gene_sets.gmt#KEGG_MAPK_SIGNALING_PATHWAY | -1.8006 | 0.0099 |
| gene_sets.gmt#KEGG_ECM_RECEPTOR_INTERACTION | -1.8096 | 0.0089 |
| gene_sets.gmt#KEGG_JAK_STAT_SIGNALING_PATHWAY | -1.8146 | 0.0086 |
| gene_sets.gmt#KEGG_PRION_DISEASES | -1.819 | 0.0081 |
| gene_sets.gmt#KEGG_FOCAL_ADHESION | -1.8231 | 0.0077 |
| gene_sets.gmt#KEGG_NEUROACTIVE_LIGAND_RECEPTOR_INTERACTION | -1.8245 | 0.0081 |
| gene_sets.gmt#KEGG_HEMATOPOIETIC_CELL_LINEAGE | -1.8258 | 0.0084 |
| gene_sets.gmt#KEGG_AUTOIMMUNE_THYROID_DISEASE | -1.8287 | 0.0088 |
| gene_sets.gmt#KEGG_LEISHMANIA_INFECTION | -1.8651 | 0.0047 |
| gene_sets.gmt#KEGG_GAP_JUNCTION | -1.8658 | 0.0052 |
| gene_sets.gmt#KEGG_MELANOGENESIS | -1.8732 | 0.0052 |
| gene_sets.gmt#KEGG_CYTOKINE_CYTOKINE_RECEPTOR_INTERACTION | -1.8825 | 0.0045 |
| gene_sets.gmt#KEGG_HYPERTROPHIC_CARDIOMYOPATHY_HCM | -1.8888 | 0.0043 |
| gene_sets.gmt#KEGG_LEUKOCYTE_TRANSENDOTHELIAL_MIGRATION | -1.8947 | 0.0045 |
| gene_sets.gmt#KEGG_CHEMOKINE_SIGNALING_PATHWAY | -1.9166 | 0.003 |
| gene_sets.gmt#KEGG_DILATED_CARDIOMYOPATHY | -1.9303 | 0.0022 |
| gene_sets.gmt#KEGG_VIRAL_MYOCARDITIS | -1.9305 | 0.0033 |
| gene_sets.gmt#KEGG_CELL_ADHESION_MOLECULES_CAMS | -1.9783 | 0.003 |

**Supplementary Table S6. Exclusive set of identified significant DEMs in Fusion-Positive and Fusion-Negative BLCA datasets.**

| Exclusive upregulated DEMs in fusion-positive BLCA | Exclusive downregulated DEMs in fusion-positive BLCA | Exclusive upregulated DEMs in fusion-negative BLCA | Exclusive downregulated DEMs in fusion-negative BLCA |
|---|---|---|---|
| hsa-mir-652 | hsa-mir-150 | hsa-mir-16-2 | hsa-mir-184 |
| hsa-mir-744 | hsa-mir-214 | hsa-mir-766 | hsa-mir-552 |
| hsa-mir-651 | hsa-mir-1245a | hsa-mir-3074 | hsa-mir-585 |
| hsa-mir-151a | hsa-mir-146b | hsa-mir-4772 | hsa-mir-218-1 |
| hsa-mir-191 | hsa-mir-142 | hsa-mir-92a-2 | hsa-mir-218-2 |
| hsa-mir-1226 | hsa-mir-155 | hsa-mir-92a-1 | hsa-mir-508 |
| hsa-mir-4728 | hsa-mir-199b | hsa-mir-2355 | hsa-mir-514a-3 |
| hsa-mir-149 | hsa-mir-199a-2 | hsa-mir-629 | hsa-mir-514a-2 |
| hsa-mir-452 | hsa-mir-223 | hsa-mir-34c | hsa-mir-23b |
| hsa-mir-3200 | hsa-mir-376b | hsa-mir-130a | hsa-mir-1468 |
| hsa-mir-301b | hsa-mir-4772 | hsa-mir-1270 | hsa-mir-664a |
| hsa-mir-7706 | hsa-mir-146a | hsa-mir-137 | hsa-mir-10a |
| hsa-mir-6715b | hsa-mir-212 | hsa-mir-21 | hsa-mir-27b |
| hsa-mir-187 | hsa-mir-221 | hsa-mir-196a-2 | hsa-mir-29c |
| hsa-mir-200c | hsa-mir-148a | hsa-mir-146b | hsa-mir-144 |
| hsa-mir-200b | hsa-mir-193a | hsa-mir-155 | hsa-mir-101-2 |
| hsa-mir-1251 | hsa-mir-887 | hsa-mir-4724 | hsa-mir-30d |
| hsa-mir-200a | hsa-mir-511 | hsa-mir-196a-1 | hsa-mir-664b |
| hsa-mir-429 | hsa-mir-222 | hsa-mir-455 | hsa-mir-101-1 |
| hsa-mir-934 | hsa-mir-130a | hsa-mir-193b | hsa-let-7e |
| hsa-mir-6499 | hsa-mir-323a | hsa-mir-34b | hsa-mir-125a |
| hsa-mir-944 | hsa-mir-338 | hsa-mir-7702 | hsa-mir-328 |
|  | hsa-mir-29a | hsa-mir-3934 | hsa-mir-30c-2 |
|  | hsa-mir-132 | hsa-mir-142 | hsa-mir-598 |
|  | hsa-mir-92b | hsa-mir-7-2 | hsa-mir-1287 |
|  | hsa-let-7i | hsa-mir-3614 | hsa-mir-30e |
|  | hsa-mir-374b | hsa-mir-1910 | hsa-mir-378a |
|  | hsa-mir-26a-2 | hsa-mir-3662 | hsa-mir-326 |
|  | hsa-mir-26a-1 | hsa-mir-4652 |  |
|  |  | hsa-mir-577 |  |
|  |  | hsa-mir-3651 |  |
|  |  | hsa-mir-522 |  |
|  |  | hsa-mir-31 |  |
|  |  | hsa-mir-147b |  |
|  |  | hsa-mir-1293 |  |

**Supplementary Table S8. Immune related genes retrieved from Immunome and IMMPORT databases.**

| | | | | | | | |
|---|---|---|---|---|---|---|---|
| CD1A | HSPA5 | DEFB133 | GDF5 | C4BPB | IGKV1-9 | OAS1 | FCGRT |
| CD1B | HSPA6 | S100Z | GDF6 | C7 | IGKV1D-12 | AGER | PDIA3 |
| CD1C | HSPA8 | MAVS | GDF7 | CFHR1 | IGKV1D-13 | UNC93B1 | HLA-DQA2 |
| CD1D | HSP90AA1 | TMSB4XP8 | GDF9 | CFHR2 | IGKV1D-16 | NOS1 | HLA-DRB1 |
| CD1E | HSP90AB1 | S100A14 | GDNF | CFHR3 | IGKV1D-17 | ACTG1 | HLA-G |
| CD8A | IFNA2 | LCN10 | INHA | CFHR4 | IGKV1D-33 | ACTA1 | HLA-H |
| MRC1 | IFNA4 | S100A16 | INHBA | CFHR5 | IGKV1D-37 | ACO1 | HSPA1A |
| CLEC4M | IFNA5 | DEFB136 | INHBB | CLU | IGKV1D-39 | SERPINA3 | HSPA1B |
| CD209 | IFNA6 | DEFB135 | INHBC | CR1L | IGKV1D-42 | CXCR1 | HSPA1L |
| PSMB10 | IFNA7 | DEFB117 | INHBE | NCR2 | IGKV1D-43 | ACKR2 | HSPA2 |
| PSMB5 | IFNA8 | ZC3HAV1L | LEFTY1 | SLAMF7 | IGKV1D-8 | ACKR4 | HSPA4 |
| TAPBP | IFNA10 | S100A7L2 | LEFTY2 | TLR4 | IGKV2-24 | TAFA5 | PTK2B |
| CANX | IFNA13 | MBL3P | NODAL | TLR5 | IGKV2-28 | TAFA3 | CDH1 |
| PSMB9 | IFNA14 | DEFB4B | TGFB2 | TLR9 | IGKV2-30 | TAFA4 | LTBP1 |
| PSMB8 | IFNA16 | BPIFB4 | TGFB3 | LYST | IGKV2-40 | TAFA1 | PPARG |
| HLA-DMA | IFNA17 | IFNAR1 | TNFRSF11B | STX11 | IGKV2D-24 | TAFA2 | FGR |
| HLA-DMB | IFNA21 | AZU1 | TNFSF18 | TCIRG1 | IGKV2D-28 | CCL15-CCL14 | JAK2 |
| HLA-A | KIR2DS3 | DEFB131B | TNFSF9 | CD2 | IGKV2D-29 | PTGDR | PTK2 |
| HLA-B | KLRC3 | DEFA1A3 | IL36RN | PSTPIP1 | IGKV2D-30 | CD38 | ACVR1C |
| HLA-C | NFYA | LCN1P1 | IL36A | CD2BP2 | IGKV2D-40 | ITGA2B | ACVR2A |
| HLA-DOA | NFYB | S100G | IL37 | CD22 | IGKV3-11 | GP9 | ACVR2B |
| HLA-DOB | NFYC | DEFA7P | IL36B | CD27 | IGKV3-15 | GP1BA | ACVRL1 |
| HLA-DPA1 | LGMN | DEFB130B | IL36G | CD28 | IGKV3-20 | GP1BB | AMHR2 |
| HLA-DPB1 | PSMC1 | DEFB108F | IFNL3 | FCGR2A | IGKV3-7 | GP5 | BMPR1A |
| HLA-DQA1 | PSMC2 | DEFB131C | IL33 | FCGR2B | IGKV3D-11 | CD44 | BMPR1B |
| HLA-DQB1 | PSMC3 | TCHHL1 | IL34 | CD37 | IGKV3D-15 | PTPRC | BMPR2 |
| HLA-DQB2 | PSMC4 | TINAGL1 | TXLNA | ENTPD1 | IGKV3D-20 | CD47 | TGFBR1 |
| HLA-DRA | PSMC5 | SLC22A17 | IFNE | CD53 | IGKV3D-7 | CD48 | TGFBR2 |
| HLA-DRB3 | PSMC6 | WFIKKN1 | IFNK | CD58 | IGKV4-1 | ITGA2 | TGFBR3 |
| HLA-DRB4 | PSMD1 | WFDC2 | IFNW1 | CD72 | IGKV5-2 | ITGA4 | TNFRSF13C |
| HLA-DRB5 | PSMD2 | UMODL1 | ADM | ST6GAL1 | IGKV6-21 | ITGA5 | TNFRSF19 |
| HLA-E | PSMD3 | TGFB1 | ADM2 | CD83 | IGKV6D-21 | ITGA6 | TNFRSF21 |
| HLA-F | PSMD4 | PF4V1 | AGRP | KLRD1 | IGKV6D-41 | ICAM3 | TNFRSF25 |
| MR1 | PSMD5 | MMP9 | AGT | IGSF2 | IGL | ITGAV | TNFRSF6B |
| TAP1 | PSMD7 | ANOS1 | AMBN | CSF1R | IGLC1 | CD52 | CXCR5 |
| PSME3 | PSMD8 | SPAG11B | AMELX | IL7R | IGLC2 | ICAM1 | ACKR3 |
| CADM2 | PSMD10 | CTSL | AMH | CSF2RB | IGLC3 | NCAM1 | CYSLTR1 |

| | | | | | | | |
|---|---|---|---|---|---|---|---|
| B2M | PSMD11 | NFKB1 | ANGPTL5 | SLAMF1 | IGLC6 | CD59 | CYSLTR2 |
| PSMB6 | PSMD13 | APOBEC3G | ANGPTL7 | BST1 | IGLC7 | ITGB3 | ACKR1 |
| PSMB7 | SLC10A2 | FABP6 | APLN | KLRC1 | IGLJ | SELL | EDNRA |
| PSME1 | THBS1 | SFTPA1 | AREG | KLRB1 | IGLJ1 | SELP | EDNRB |
| PSME2 | SEM1 | RBP1 | MANF | ALCAM | IGLJ2 | CD63 | FPR1 |
| TAP2 | KLRC4 | SLC40A1 | CDNF | CXCR3 | IGLJ3 | FCGR1A | FPR2 |
| ULBP1 | AP3B1 | PLAU | ARTN | CD226 | IGLJ4 | CEACAM1 | GPR17 |
| PSMF1 | PSMD6 | PAEP | AVP | LY9 | IGLJ5 | CEACAM8 | GPR32 |
| CTSC | PSMD14 | HJV | BDNF | XCL1 | IGLJ6 | CEACAM6 | GPR33 |
| IFNGR2 | IFI30 | MUC5AC | BTC | MASP2 | IGLJ7 | CEACAM3 | PTGDR2 |
| PTGFRN | ADRM1 | OBP2A | MYDGF | IL2 | IGLV | CEACAM5 | C5AR2 |
| CD14 | ECPAS | PLTP | CALCA | DEFA3 | IGLV1-36 | PSG1 | CXCR2 |
| FCGR3A | TRPC4AP | DDX58 | CALCB | DEFA5 | IGLV1-40 | CD68 | LTB4R2 |
| FCGR3B | UBXN1 | IFNL1 | CAT | DEFA6 | IGLV1-44 | CD69 | PLXNA1 |
| ITGB2 | ERAP1 | IRF3 | CCK | FADD | IGLV1-47 | TFRC | PLXNA2 |
| FCER2 | TAPBPL | SFTPA2 | ADA2 | KLRK1 | IGLV1-50 | NT5E | PLXNA3 |
| CD40 | ERAP2 | LPA | CER1 | TNFRSF17 | IGLV1-51 | CD74 | PLXNA4 |
| CD40LG | ULBP3 | LBP | CGA | ADA | IGLV10-54 | CD79A | PLXNB1 |
| SELE | ULBP2 | RBP4 | CGB3 | RFXANK | IGLV11-55 | CD79B | PLXNB2 |
| C5AR1 | RAET1E | NOX4 | CGB1 | AIRE | IGLV2-11 | CD81 | PLXNB3 |
| CD97 | RAET1L | RBP5 | CGB2 | BLM | IGLV2-14 | CD82 | PLXND1 |
| IL1R1 | UBR1 | FABP7 | CGB5 | BATF | IGLV2-18 | CD84 | ROBO1 |
| IL1R2 | RAET1G | FABP5 | CGB7 | SH2B2 | IGLV2-23 | LILRB1 | ROBO2 |
| IL6R | PDIA2 | FABP3 | CGB8 | IFI44L | IGLV2-33 | PLAUR | RXFP3 |
| IL8RA | PTPN6 | FABP2 | CHGA | DOCK2 | IGLV2-8 | FCAR | IL20RB |
| IL8RB | PTPN11 | FABP4 | CHGB | ICOS | IGLV3-1 | THY1 | ST2 |
| IL6ST | VAV3 | R3HDML | CLEC11A | PILRA | IGLV3-10 | LRP1 | ADCYAP1R1 |
| SIGLEC1 | VAV1 | BPIFA3 | CNTF | SH2D1A | IGLV3-12 | SLC44A1 | ADIPOR1 |
| CD180 | VAV2 | BPIFB1 | CORT | MX1 | IGLV3-16 | CD96 | ADIPOR2 |
| CXCR4 | RAC1 | OASL | CRH | NFKBIA | IGLV3-19 | SLC3A2 | ADRB1 |
| CCR5 | RAC2 | CRABP2 | CSH1 | POU2AF1 | IGLV3-21 | SLC7A5 | ADRB2 |
| CCR7 | RAC3 | CRABP1 | CSH2 | LAX1 | IGLV3-22 | CD99 | AGTR1 |
| PROCR | PAK1 | RBP7 | CSHL1 | STAT3 | IGLV3-25 | SEMA4D | AGTR2 |
| LY75 | TYROBP | DUOX1 | CSPG5 | STAT4 | IGLV3-27 | ICAM2 | ANGPT1 |
| IL12RB1 | LCK | OBP2B | CTF1 | STAT5B | IGLV3-32 | ITGAE | ANGPT4 |
| CCR1 | ZAP70 | RBP2 | CCN2 | TAL1 | IGLV3-9 | ITGB4 | ANGPTL1 |
| CCR2 | SYK | LCN15 | DKK1 | HSP90B1 | IGLV4-3 | ENG | ANGPTL2 |
| CCR3 | LCP2 | CETP | EBI3 | FCAMR | IGLV4-60 | VCAM1 | ANGPTL3 |
| CCL1 | LAT | FABP12 | EGF | BST2 | IGLV4-69 | LAMP1 | ANGPTL4 |
| CCL2 | PLCG1 | FABP9 | EPGN | BTLA | IGLV5-37 | LAMP2 | ANGPTL6 |
| CCL3 | PLCG2 | BPIFA1 | EPO | CD274 | IGLV5-39 | SEMA7A | APLNR |
| CCL4 | SH3BP2 | LCNL1 | EREG | CD276 | IGLV5-45 | CD109 | AR |
| CCL5 | PIK3CA | SPAG11A | ESM1 | FCGR2C | IGLV5-48 | PVRL1 | AVPR1A |

| | | | | | | | |
|---|---|---|---|---|---|---|---|
| CCL7 | PIK3CB | PI15 | FAM3B | FCRL5 | IGLV5-52 | PVRL2 | AVPR1B |
| CCL8 | PIK3CD | NOX1 | FAM3C | IL27 | IGLV6-57 | KIT | AVPR2 |
| CCL11 | PIK3R5 | PMP2 | FAM3D | PDCD1LG2 | IGLV7-43 | PROM1 | BRD8 |
| CCL13 | PIK3R1 | APOD | FGF1 | PDCD1 | IGLV7-46 | FLT3 | CALCR |
| CCL15 | PIK3R2 | ORM2 | FGF11 | STAT6 | IGLV8-61 | SDC1 | CALCRL |
| CCL16 | PIK3R3 | ORM1 | FGF12 | IGHG2 | IGLV9-49 | PDGFRA | CNTFR |
| CCL17 | FYN | CTSG | FGF13 | IGHM | ITK | PDGFRB | CRHR1 |
| CCL18 | SHC2 | PRTN3 | FGF14 | BLNK | TEC | THBD | CRHR2 |
| CCL19 | SHC4 | MAPK1 | FGF16 | CCRN4L | NCK1 | ACE | CRIM1 |
| CCL20 | SHC3 | PML | FGF17 | CIITA | NCK2 | CDH5 | EGFR |
| CCL21 | SHC1 | AEN | FGF18 | RFX5 | GRAP2 | MCAM | EPOR |
| CCL22 | GRB2 | BPIFA2 | FGF19 | RFXAP | PAK2 | BSG | ESR1 |
| CCL23 | SOS1 | ISG20 | FGF20 | ATM | PAK3 | PTPRJ | ESR2 |
| CCL24 | SOS2 | BCL3 | FGF21 | STAT2 | PAK4 | CD151 | ESRRA |
| CCL25 | HRAS | ISG20L2 | FGF22 | NFATC2 | PAK6 | CTLA4 | ESRRB |
| CCL26 | KRAS | NOX5 | FGF23 | FOXP3 | PAK5 | SELPLG | ESRRG |
| CXCL1 | NRAS | NOX3 | FGF3 | TCF3 | RHOA | CD163 | FGFR2 |
| CXCL2 | ARAF | DUOX2 | FGF4 | EBF1 | CDC42 | CD164 | FGFRL1 |
| CXCL3 | BRAF | IFIH1 | FGF5 | RELA | MAP3K8 | DDR1 | FLT1 |
| CXCL5 | RAF1 | TRIM5 | FGF6 | NFATC1 | MAP3K14 | HMMR | FLT4 |
| CXCL6 | HCST | IDO1 | FGF7 | IRF9 | CBLC | L1CAM | FSHR |
| IL8 | PPP3CA | GDF15 | FGF8 | STAT1 | CBL | FUT3 | GALR2 |
| CXCL9 | PPP3CB | NEDD4 | FGF9 | IRF8 | CBLB | VPREB1 | GALR3 |
| CXCL10 | PPP3CC | ADIPOQ | VEGFD | IRF1 | CDK4 | IGLL1 | GCGR |
| CXCL11 | CHP1 | IFNL2 | FIGNL2 | IRF2 | RASGRP1 | CD200 | GHR |
| CXCL12 | PPP3R1 | SOCS3 | FLT3LG | RFX1 | PDK1 | TEK | GHRHR |
| CXCL13 | PPP3R2 | SEMG1 | FSHB | TCF7 | PRKCQ | MSR1 | GHSR |
| CX3CL1 | CHP2 | SOCS1 | GAL | TBX21 | TRAC | CD207 | GIPR |
| SCYE1 | NFAT5 | RNASEL | GALP | IRF5 | TRAJ1 | LAMP3 | GLP1R |
| CXCL14 | PRKCA | APOBEC3F | GAST | RELB | TRAJ2 | INSR | GLP2R |
| IL9 | PRKCB | PLAAT4 | GCG | FOXN1 | TRAJ3 | IGF1R | GNRHR |
| CSF3 | PRKCG | CHIT1 | GH1 | CD3EAP | TRAJ4 | IGF2R | GPER1 |
| NOS2A | SH2D1B | IFNA1 | GH2 | FOXK2 | TRAJ5 | LAG3 | HNF4A |
| TIRAP | CASP3 | TLR7 | GHRH | ZEB1 | TRAJ6 | MUC1 | HNF4G |
| IRAK1 | BID | HFE | GHRL | FANCD2 | TRAJ7 | MFI2 | HTR3A |
| LY96 | LYN | ZYX | GIP | GFI1 | TRAJ8 | PRNP | HTR3B |
| HRH4 | BTK | NLRX1 | GKN1 | SP110 | TRAJ9 | TSPAN7 | HTR3C |
| CYBB | FOS | PGC | GMFB | CR2 | TRAJ10 | PLXNC1 | HTR3D |
| IL12B | CARD11 | VEGFA | GMFG | CR1 | TRAJ11 | SLC4A1 | HTR3E |
| CAMP | BCL10 | IKBKE | GNRH1 | CD46 | TRAJ12 | DARC | LEPR |
| TLR1 | MALT1 | ISG15 | GNRH2 | CD55 | TRAJ13 | GYPA | LGR4 |
| TLR2 | CHUK | DHX58 | GPHA2 | CD93 | TRAJ14 | GYPB | LGR5 |
| C4A | IKBKB | TNFAIP3 | GPHB5 | C2 | TRAJ15 | GYPC | LGR6 |

| | | | | | | | |
|---|---|---|---|---|---|---|---|
| MPO | IKBKG | TFR2 | GPI | CFB | TRAJ16 | KEL | LHCGR |
| LYZ | NFKBIB | FCN2 | GREM1 | MBL2 | TRAJ17 | BCAM | MC1R |
| LTF | NFKBIE | MUC4 | GREM2 | MASP1 | TRAJ18 | RHCE | MC2R |
| TIMP1 | AKT3 | F2R | GRP | C1QA | TRAJ19 | RHD | MC3R |
| YWHAZ | AKT1 | ELN | GUCA2A | C1QB | TRAJ20 | RHAG | MC4R |
| PLAA | AKT2 | MAPT | HBEGF | C8A | TRAJ21 | ICAM4 | MCHR1 |
| PLA2R1 | GSK3B | LEP | HDGF | C8B | TRAJ22 | ABCB1 | MCHR2 |
| MARCO | INPP5D | CYLD | HDGFL3 | C8G | TRAJ23 | CD244 | MET |
| PAFAH1B1 | RASGRP3 | KLKB1 | IAPP | SERPING1 | TRAJ24 | ALK | MLNR |
| PAFAH1B2 | IGH | CST4 | IGF1 | CFD | TRAJ25 | CD247 | MTNR1A |
| PAFAH1B3 | IGHA1 | CSRP1 | IGF2 | CFI | TRAJ26 | KIR2DL1 | MTNR1B |
| PLA2G7 | IGHA2 | JUN | INS | CFH | TRAJ27 | KIR2DL3 | NGFR |
| NFATC3 | IGHD | TLR8 | INS-IGF2 | CFP | TRAJ28 | KIR2DS1 | NMBR |
| NFATC4 | IGHD1-1 | EIF2AK2 | INSL3 | C4B | TRAJ29 | KIR2DS2 | NPR1 |
| C3 | IGHD1-14 | APOM | INSL4 | XRCC5 | TRAJ30 | KIR2DS5 | NPR3 |
| IFNB1 | IGHD1-20 | CACYBP | INSL5 | C1QBP | TRAJ31 | KIR3DL1 | NR0B1 |
| PAFAH2 | IGHD1-26 | NOD1 | INSL6 | C1QC | TRAJ32 | SLAMF6 | NR0B2 |
| CASP10 | IGHD1-7 | MAPK8 | JAG1 | C1R | TRAJ33 | ADAM10 | NR1D1 |
| PRG2 | IGHD2-15 | MAPK3 | JAG2 | C6 | TRAJ34 | CD164L2 | NR1D2 |
| TNF | IGHD2-2 | BPHL | FGF7P6 | C9 | TRAJ35 | CD200R1 | NR1H2 |
| CCR4 | IGHD2-21 | PLA2G2A | FGF7P3 | C1QL1 | TRAJ36 | CD200R2 | NR1H3 |
| FCER1A | IGHD2-8 | GRN | KITLG | C1QL2 | TRAJ37 | CD248 | NR1H4 |
| HRH2 | IGHD3-10 | NEWENTRY | KL | C1QL3 | TRAJ38 | CD24 | NR1I2 |
| IL1A | IGHD3-16 | GNAI1 | LACRT | C1QL4 | TRAJ39 | CD300E | NR1I3 |
| IL1B | IGHD3-22 | WNT5A | LHB | C1QTNF2 | TRAJ40 | CD300LB | NR2C1 |
| IL5 | IGHD3-3 | FURIN | LRSAM1 | C1QTNF3 | TRAJ41 | CD300LF | NR2C2 |
| IL13 | IGHD3-9 | ADAR | LTBP2 | F12 | TRAJ42 | CD300LG | NR2E1 |
| MIF | IGHD4-11 | TYK2 | LTBP3 | LTBR | TRAJ43 | CD302 | NR2E3 |
| PTAFR | IGHD4-17 | NOS2 | LTBP4 | IL2RA | TRAJ44 | CD320 | NR2F1 |
| IL18R1 | IGHD4-23 | TRAF3 | MDK | CD70 | TRAJ45 | SIGLEC15 | NR2F2 |
| CRP | IGHD4-4 | TPT1 | MIA | TNFRSF8 | TRAJ46 | CD99L2 | NR2F6 |
| IL1F5 | IGHD5-12 | TPM2 | MLN | TNFSF8 | TRAJ47 | FGFR1 | NR3C1 |
| IL1F6 | IGHD5-18 | NEO1 | MSTN | CD80 | TRAJ48 | FGFR3 | NR3C2 |
| IL1RN | IGHD5-24 | AHNAK | NAMPT | CD86 | TRAJ49 | FGFR4 | NR4A1 |
| MYD88 | IGHD5-5 | TK2 | NDP | FAS | TRAJ50 | IFITM1 | NR4A2 |
| IL20 | IGHD6-13 | PRDX2 | NENF | FASLG | TRAJ52 | ITGA1 | NR4A3 |
| IL22 | IGHD6-19 | MX2 | NGF | MPL | TRAJ53 | ITGAD | NR5A1 |
| TOLLIP | IGHD6-25 | FGF2 | NMB | CSF3R | TRAJ54 | KDR | NR5A2 |
| IL1F9 | IGHD6-6 | FGA | CCN3 | IFNGR1 | TRAJ56 | KIR2DL2 | NR6A1 |
| S100A8 | IGHD7-27 | TCF7L2 | NPFF | TNFRSF1A | TRAJ57 | KIR2DL4 | NRP1 |
| CCL3L1 | IGHE | F2RL1 | NPPA | TNFRSF1B | TRAJ58 | KIR2DL5A | NRP2 |
| IL25 | IGHG1 | TKFC | NPPB | IL2RB | TRAJ59 | KIR2DS4 | OGFR |
| C5 | IGHG3 | NFKBIZ | NPPC | IL4R | TRAJ61 | KIR3DL2 | OPRD1 |

| | | | | | | | |
|---|---|---|---|---|---|---|---|
| IL1F10 | IGHG4 | LMBR1 | NPY | IL5RA | TRAV1-1 | KIR3DL3 | OPRK1 |
| FCRLA | IGHJ1 | EPPIN | NRG1 | IL2RG | TRAV1-2 | KLRC2 | OPRL1 |
| LY86 | IGHJ2 | SRC | NRG2 | TNFSF4 | TRAV2 | LAIR1 | OPRM1 |
| ABCF1 | IGHJ3 | ELAVL1 | NRG3 | TNFRSF4 | TRAV3 | LAIR2 | OSMR |
| NOD2 | IGHJ4 | ROBO3 | NRG4 | MST1R | TRAV4 | LIFR | OXTR |
| CASP1 | IGHJ5 | SP1 | NRTN | TNFRSF9 | TRAV5 | LILRA1 | PGR |
| CCRL2 | IGHJ6 | SOD1 | NTF3 | F3 | TRAV7 | LILRA2 | PGRMC2 |
| CKLF | IGHV1-18 | PDF | NTF4 | IL13RA1 | TRAV8-1 | LILRA3 | PPARA |
| LTB4R | IGHV1-2 | DLL4 | NTS | IL13RA2 | TRAV8-2 | LILRA4 | PPARD |
| MAPK14 | IGHV1-24 | ECD | NUDT6 | IL17RA | TRAV8-3 | LILRA5 | PRLHR |
| NCR3 | IGHV1-3 | SLC11A1 | OGN | BLR1 | TRAV8-4 | LILRA6 | PRLR |
| XCR1 | IGHV1-45 | DMBT1 | OSGIN1 | CCR8 | TRAV8-6 | LILRB3 | PTGER1 |
| ELA2 | IGHV1-46 | STING1 | OSM | CCBP2 | TRAV8-7 | LILRB5 | PTGER2 |
| CD3D | IGHV1-58 | SKIV2L | OSTN | CCR10 | TRAV9-1 | MRC2 | PTGER3 |
| CD3E | IGHV1-69 | SEMG2 | OXT | CCL14 | TRAV9-2 | SIVA1 | PTGER4 |
| CD3G | IGHV1-8 | DES | ENDOU | CCL27 | TRAV10 | TNFRSF10A | PTGFR |
| CD4 | IGHV1-38-4 | DCK | PDGFA | CCL28 | TRAV12-1 | TNFRSF10B | PTH1R |
| CD5 | IGHV1-69-2 | DAXX | PDGFB | PF4 | TRAV12-2 | TNFRSF10C | PTH2R |
| CD5L | IGHV2-26 | EED | PDGFC | PPBP | TRAV12-3 | TNFRSF10D | RARA |
| CD7 | IGHV2-5 | LIMS1 | PDGFD | XCL2 | TRAV13-1 | TNFRSF11A | RARB |
| CD8B | IGHV2-70 | LALBA | PDGFRL | CXCL16 | TRAV13-2 | TNFRSF12A | RARG |
| ITGAL | IGHV3-11 | APOBEC3H | PGF | IL3 | TRAV14DV4 | TNFRSF18 | RORA |
| ITGAM | IGHV3-13 | TMPRSS6 | PMCH | IL11 | TRAV16 | TNFSF10 | RORB |
| ITGAX | IGHV3-15 | SPINK5 | PNOC | LIF | TRAV17 | TNFSF11 | RORC |
| CD19 | IGHV3-16 | BECN1 | POMC | IL10 | TRAV18 | PI3 | RXFP1 |
| ITGB1 | IGHV3-20 | KNG1 | PPBPP2 | LTA | TRAV19 | DEFB4A | RXFP2 |
| SPN | IGHV3-21 | CSK | PPY | LTB | TRAV20 | REG3G | RXRA |
| CD160 | IGHV3-23 | KCNH2 | PRL | TNFSF13 | TRAV21 | SLPI | RXRB |
| CCR6 | IGHV3-30 | JUND | PRLH | TNFSF13B | TRAV22 | CXCL8 | RXRG |
| CCR9 | IGHV3-30-3 | JAK1 | PROK1 | TNFRSF13B | TRAV23DV6 | DEFB103B | S1PR1 |
| IL4 | IGHV3-30-5 | CREB1 | PSPN | TRADD | TRAV24 | ELANE | S1PR2 |
| PRF1 | IGHV3-33 | CLDN4 | PTH | IFNAR2 | TRAV25 | TMSB10 | SCTR |
| GZMA | IGHV3-35 | RNASE3 | PTH2 | IFNG | TRAV26-1 | LCN2 | SDC2 |
| GZMB | IGHV3-38 | RN7SL1 | PTHLH | TNFSF14 | TRAV26-2 | LCN1 | SDC3 |
| GZMK | IGHV3-43 | IRF7 | PTN | TNFRSF14 | TRAV27 | COLEC10 | SDC4 |
| GZMM | IGHV3-48 | IREB2 | PYY | CSF1 | TRAV29DV5 | BPI | SORT1 |
| LPO | IGHV3-49 | ILK | QRFP | NPTN | TRAV30 | S100A9 | SSTR1 |
| CD300C | IGHV3-53 | APOBEC3A | RABEP1 | ILF3 | TRAV34 | LCN6 | SSTR2 |
| NCF2 | IGHV3-64 | TRIM27 | RABEP2 | IFI16 | TRAV35 | S100A12 | SSTR5 |
| GNLY | IGHV3-66 | PTX3 | REG1A | IFI27 | TRAV36DV7 | LCN8 | TACR1 |
| NCR1 | IGHV3-7 | IFN1@ | RETN | IFI35 | TRAV38-1 | DEFA1B | THRA |

| | | | | | | | |
|---|---|---|---|---|---|---|---|
| MICA | IGHV3-72 | SYTL1 | RETNLB | IFIT2 | TRAV38-2DV8 | CELA1 | THRB |
| MICB | IGHV3-73 | APOBEC3C | RLN1 | IFIT1 | TRAV39 | PENK | TIE1 |
| CSF2 | IGHV3-74 | DDX17 | RLN2 | IFIT3 | TRAV40 | BPIFC | TRHR |
| PRSS16 | IGHV3-9 | PTGS2 | RLN3 | IL15 | TRAV41 | MMP12 | TSHR |
| CLEC10A | IGHV3-38-3 | HTR1A | SCG2 | IL16 | TRBC1 | BPIFB6 | TUBB3 |
| PGLYRP3 | IGHV3-69-1 | SEPTIN7 | SCGB3A1 | IL17A | TRBC2 | SFTPD | VIPR1 |
| LEAP2 | IGHV4-28 | PROC | SCT | IL18 | TRBD1 | LCN9 | VIPR2 |
| DCD | IGHV4-30-1 | MAP2K2 | AIMP1 | IL17B | TRBD2 | BPIFB2 | CCL3P1 |
| WFDC12 | IGHV4-30-2 | MAP2K1 | SECTM1 | IL24 | TRBJ1-1 | PTGDS | CMA1 |
| A2ML1 | IGHV4-30-4 | HRG | SLURP1 | IL23R | TRBJ1-2 | TMSB4X | CXCL17 |
| CTSS | IGHV4-31 | NDRG1 | SPP1 | IL1F8 | TRBJ1-3 | PGLYRP1 | CCN1 |
| CLEC12A | IGHV4-34 | TRIM22 | SST | IL19 | TRBJ1-4 | ZC3HAV1 | EDN1 |
| DEFA1 | IGHV4-39 | LANCL1 | STC1 | IL6 | TRBJ1-5 | TMSB15A | EDN2 |
| DEFA4 | IGHV4-4 | PPP4C | STC2 | IL7 | TRBJ1-6 | S100B | EDN3 |
| DEFB1 | IGHV4-59 | HMOX1 | TAC1 | IL12A | TRBJ2-1 | S100A13 | FGF10 |
| A2M | IGHV4-61 | HMGB1 | TDGF1 | IL27RA | TRBJ2-2 | S100A6 | LECT2 |
| CLEC5A | IGHV4-38-2 | ABCC4 | TDGF1P3 | CCL3L3 | TRBJ2-3 | DEFB107A | PPBPP1 |
| DEFB105A | IGHV5-51 | HGF | TG | CCL4L1 | TRBJ2-4 | SERPIND1 | PROK2 |
| DEFB106A | IGHV5-10-1 | HDAC1 | TGFA | CCL4L2 | TRBJ2-5 | DEFB129 | SAA1 |
| DEFB119 | IGHV6-1 | IFNLR1 | THPO | CCRL1 | TRBJ2-6 | DEFB127 | SAA2 |
| DEFB123 | IGHV7-4-1 | PLSCR1 | TOR2A | CLCF1 | TRBJ2-7 | S100P | SBDS |
| LYG2 | IGHV7-81 | BACH2 | TRH | CMKLR1 | TRBV2 | S100A7 | SEMA3A |
| CLEC4E | IGK | TANK | TSHB | CXCR7 | TRBV3-1 | DEFB104A | SEMA3B |
| HTN3 | IGKC | PIK3CG | TSLP | CMTM1 | TRBV4-1 | DEFB126 | SEMA3C |
| CLEC4D | IGKDEL | ARRB1 | UCN | CMTM2 | TRBV4-2 | DEFB106B | SEMA3D |
| NCF1C | IGKJ | RSAD2 | UCN2 | CMTM3 | TRBV4-3 | DEFB104B | SEMA3E |
| NCF4 | IGKJ1 | STAB2 | UCN3 | CMTM4 | TRBV5-1 | DEFB107B | SEMA3F |
| CLEC4A | IGKJ2 | TBK1 | UTS2 | CMTM5 | TRBV5-4 | PGLYRP2 | SEMA3G |
| PPIA | IGKJ3 | PDYN | UTS2B | CMTM6 | TRBV5-5 | S100A10 | SEMA4A |
| DEFB103A | IGKJ4 | PCSK2 | VEGFB | CMTM7 | TRBV5-6 | S100A2 | SEMA4B |
| HAMP | IGKJ5 | PCSK1 | VEGFC | CMTM8 | TRBV5-7 | DEFB125 | SEMA4C |
| CLEC7A | IGKV | ARG2 | VGF | CRLF1 | TRBV5-8 | DEFB105B | SEMA4F |
| COLEC12 | IGKV1-12 | AQP9 | VIP | CRLF2 | TRBV6-1 | DEFB132 | SEMA4G |
| RNASE7 | IGKV1-13 | APOH | VDR | CRLF3 | TRBV6-2 | BPIFB3 | SEMA5A |
| C1QTNF4 | IGKV1-16 | BIRC5 | OLR1 | CX3CR1 | TRBV6-3 | LCN12 | SEMA5B |
| C1QTNF5 | IGKV1-17 | ANXA6 | GRK2 | CXCR6 | TRBV6-4 | PGLYRP4 | SEMA6A |
| C1QTNF6 | IGKV1-27 | VTN | TXK | CYTL1 | TRBV6-5 | S100A11 | SEMA6B |
| C1QTNF7 | IGKV1-33 | VIM | RNASE2 | CSF2RA | TRBV6-6 | S100A5 | SEMA6C |
| C1RL | IGKV1-37 | PRDX1 | AZGP1 | IL10RA | TRBV6-7 | S100A3 | SEMA6D |
| C1S | IGKV1-39 | GFAP | CALR | IL10RB | TRBV6-8 | S100A1 | SLIT1 |

| | | | | | | | |
|---|---|---|---|---|---|---|---|
| C3AR1 | IGKV1-5 | GBP2 | CTSB | IL11RA | TRBV6-9 | DEFB128 | SLIT2 |
| C4BPA | IGKV1-6 | ALB | CTSE | IL12RB2 | TRBV7-2 | DEFB108B | TNC |
| TRDJ3 | IGKV1-8 | SLC29A3 | FCER1G | IL15RA | TRBV7-3 | HTN1 | TYMP |
| IL28RA | TRBV25-1 | MS4A5 | TRGV2 | IL17C | TRBV7-4 | LMBR1L | BMP1 |
| IL29 | TRBV27 | DPP4 | TRGJP2 | IL17D | TRBV7-6 | S100A7A | BMP10 |
| IL31 | TRBV28 | PECAM1 | TRGJP1 | IL17F | TRBV7-7 | DEFB118 | BMP15 |
| IL31RA | TRBV29-1 | CD33 | TRGJP | IL17RB | TRBV7-8 | TMSB4Y | BMP2 |
| IL32 | TRBV30 | SIGLEC6 | TRGJ2 | IL17RC | TRBV7-9 | DEFB131A | BMP3 |
| IL3RA | TRDC | SIGLEC5 | TRGJ1 | IL17RD | TRBV9 | DEFB134 | BMP4 |
| IL4I1 | TRDD1 | CD34 | TRGC2 | IL17RE | TRBV10-1 | DEFB130A | BMP5 |
| IL9R | TRDD2 | CD36 | TRGC1 | IL18BP | TRBV10-2 | DEFB124 | BMP6 |
| ILF2 | TRDD3 | SCARB1 | TRAV6 | IL18RAP | TRBV10-3 | DEFB121 | BMP7 |
| SIGIRR | TRDJ1 | SCARB2 | ACVR1B | IL1F7 | TRBV11-1 | DEFB116 | BMP8A |
| TICAM2 | TRDJ2 | TNFSF12 | TRDJ4 | IL1RAP | TRBV11-2 | DEFB115 | BMP8B |
| IL21R | TRBV15 | TSPYL2 | TRDV2 | IL1RAPL1 | TRBV11-3 | DEFB114 | GDF1 |
| IL22RA1 | TRBV16 | CD6 | TRDV3 | IL1RAPL2 | TRBV12-3 | DEFB113 | GDF10 |
| IL22RA2 | TRBV17 | CD9 | TRGV9 | IL1RL1 | TRBV12-4 | DEFB112 | GDF11 |
| IL23A | TRBV18 | MME | TRGV8 | IL1RL2 | TRBV12-5 | DEFB110 | GDF2 |
| IL26 | TRBV19 | ANPEP | TRGV5 | IL20RA | TRBV13 | TMSB15B | GDF3 |
| IL28A | TRBV20-1 | MS4A1 | TRGV4 | IL21 | TRBV14 | TNFSF15 | TRDV1 |
| IL28B | TRBV24-1 | MS4A3 | TRGV3 | TLR3 | HCK | | |
